# Evolutionary conservation and divergence of CXCL17 orthologs: functional evidence in reptiles and loss in the avian lineage

**DOI:** 10.64898/2026.05.18.725876

**Authors:** Jie Yu, Hao-Zheng Li, Juan-Juan Wang, Ya-Li Liu, Zhan-Yun Guo

## Abstract

The mucosal chemoattractant C-X-C motif chemokine ligand 17 (CXCL17) was recently identified as a ligand for the orphan G protein-coupled receptor 25 (GPR25). Although our recent studies have identified non-mammalian CXCL17 orthologs in fishes and amphibians, their presence in reptiles and birds remains unclear. In this study, we employed bioinformatic searches based on gene synteny and sequence features to identify reptilian and avian CXCL17 orthologs in public databases. We identified intact CXCL17 orthologs in 46 reptilian species, including lizards, snakes, turtles, and alligators. In contrast, we found only *CXCL17* gene relics in 22 bird species, suggesting the avian lineage lost functional CXCL17 during evolution. A recombinant reptilian CXCL17 from the loggerhead turtle (*Caretta caretta*), termed Cc-CXCL17, directly bound to and efficiently activated its corresponding receptor, Cc-GPR25, in a C-terminal fragment-dependent manner. Activation of Cc-GPR25 by recombinant Cc-CXCL17 also induced chemotactic movement of transfected human embryonic kidney (HEK) 293T cells. In cross-species activity assays, recombinant CXCL17s from other species had almost no activity towards Cc-GPR25; Cc-CXCL17 and human CXCL17 had low activity towards mallard GPR25, but all tested CXCL17s had no detectable activity towards chicken GPR25. Our findings demonstrate the presence of functional CXCL17 orthologs in extant reptiles and provide evidence for their evolutionary loss in the avian lineage, offering new insights into the phylogenetic distribution of the newly identified CXCL17−GPR25 signaling system. −

## 1. Introduction

C-X-C motif chemokine ligand 17 (CXCL17) is a mucosal chemoattractant that recruits several leukocyte populations, including T cells, monocytes, macrophages, and dendritic cells [1−9]. CXCL17 has also been implicated in the progression of multiple tumors, likely through modulation of tumor-associated immune responses [10−16]. Although the chemoattractant function of CXCL17 is expected to be mediated by a cell-surface receptor, its identity has remained controversial for years [17−20]. While CXCL17 was recently reported as a modulator of the chemokine receptor CXCR4 [20], this observation lacks independent validation. Additional studies reported that CXCL17 acts as a weak agonist of several MAS-related receptors, including MRGPRX2, MRGPRX1, and MAS1 [21,22]. However, this activity does not depend on the conserved C-terminal fragment of CXCL17 [22], suggesting that these MAS-related receptors are unlikely to represent evolutionarily conserved receptors for CXCL17. Two recent independent studies, including our own, demonstrated that CXCL17 activates the poorly characterized orphan G protein-coupled receptor 25 (GPR25) primarily through its conserved C-terminal region [23,24]. A recent study further confirmed their interaction [25], strongly supporting GPR25 as a bona fide receptor for CXCL17. Despite its designation as a C-X-C motif chemokine, CXCL17 does not appear to be a classical chemokine because of its highly flexible conformation [26,27] and its distinct mode of interaction with GPR25 [23−25].

GPR25 orthologs are widely distributed across vertebrates, from fishes to mammals, whereas CXCL17 orthologs initially appeared to be restricted to the mammalian lineage [28], raising questions regarding the evolutionary origin of CXCL17 and its functional pairing with receptor GPR25. To address this issue, we recently searched public databases and identified CXCL17 orthologs in diverse fish and amphibian species based on characteristic amino acid sequence features and conserved gene synteny [29−32]. These findings suggest that CXCL17 has served as an evolutionarily conserved ligand for GPR25 since emergence in ancient fish ancestors. The CXCL17 orthologs identified in fishes and amphibians exhibit little overall sequence similarity to mammalian CXCL17s and therefore cannot be recognized using conventional homology-based approaches [28]. Consequently, they were previously uncharacterized proteins or entirely unannotated in public databases. According to our recent findings [29−32], some public databases, such as National Center for Biotechnology Information (NCBI), Zebrafish Information Network (ZFIN), and Xenbase, have updated their annotations for some fish and amphibian CXCL17 orthologs and paralogs.

To clarify the phylogenetic distribution of the CXCL17−GPR25 signaling system across vertebrate lineages, we searched public databases for reptilian and avian CXCL17 orthologs using our previously established strategy in this study. We identified intact CXCL17 orthologs in 46 reptilian species, including lizards, snakes, turtles, and alligators, whereas only nonfunctional *CXCL17* gene relics, rather than intact CXCL17 orthologs, were detected in 22 bird species. Following recombinant expression, a representative reptilian CXCL17 from the loggerhead turtle (*Caretta caretta*), termed Cc-CXCL17, exhibited strong activity toward its corresponding receptor, Cc-GPR25, in cell-based functional assays. However, deletion of the three C-terminal residues markedly reduced its activity. We also evaluated the cross-species activities of fish, amphibian, and human CXCL17 orthologs toward reptilian and avian GPR25 receptors. Our findings support the presence of a functional CXCL17−GPR25 signaling system in extant reptiles and suggest that this signaling pathway was likely lost during avian evolution.

## 2. Materials and methods

### 2.1. Recombinant preparation of Cc-CXCL17 proteins

The DNA fragment encoding an N-terminally 6×His-tagged Cc-CXCL17 was chemically synthesized using *Escherichia coli*-biased codons and ligated into a pET vector via Gibson assembly, resulting in the bacterial expression construct pET/6×His-Cc-CXCL17 (Fig. S1). The expression construct pET/6×His-Cc-[desC3]CXCL17 for the C-terminally truncated version was generated via QuikChange approach using pET/6×His-Cc-CXCL17 as the mutagenesis template. The expression construct pET/6×His-SmBiT-Cc-CXCL17 for the small NanoLuc fragment SmBiT-fused version was generated via polymerase chain reaction (PCR) amplification and subsequent cloning into the expression vector (Fig. S1). Thereafter, these Cc-CXCL17 proteins were recombinantly prepared via bacterial overexpression, purification, and *in vitro* refolding according to our previous procedures developed for other CXCL17 orthologs [24,29−32]. After the refolded Cc-CXCL17 proteins were purified by high performance liquid chromatography (HPLC) and lyophilized, they were dissolved in 1.0 mM aqueous hydrochloride (pH 3.0) and quantified by ultra-violet absorbance at 280 nm according to their theoretical extinction coefficient (5875 M^-1^ cm^-1^ for wild-type or C-terminally truncated version; 7365 M^-1^ cm^-1^ for the SmBiT-fused version). During purification, samples at different preparation steps were analyzed by sodium dodecyl sulfate-polyacrylamide gel electrophoresis (SDS-PAGE).

### 2.2. Generation of expression constructs for reptilian and avian GPR25s

The coding region of Cc-GPR25 from loggerhead turtle (*C. caretta*), Ap-GPR25 from mallard (*Anas platyrhynchos*), and Gg-GPR25 from chicken (*Gallus gallus*) were chemically synthesized at CWbio (Taizhou, Jiangsu province, China) according to their NCBI reference sequence with or without codon optimization and subsequently ligated into a pcDNA3.1 vector (Table S1 and Fig. S2). The NCBI reference sequence of Gg-GPR25 (Gene ID: 770582; mRNA ID: XM_015275387; protein ID: XP_015130873) was probably incorrectly annotated because it carries a long extracellular N-terminal domain without a cleavable signal peptide. Compared to other GPR25 orthologs, it seemed that Gg-GPR25 should be translated from the second methionine residue (Fig. S3), thus our synthetic Gg-GPR25 is a short form with 365 amino acids (Table S1 and Fig. S2). Subsequently, the synthetic reptilian or avian GPR25 genes were PCR amplified and ligated into appropriate doxycycline (Dox)-responsive expression vectors for functional assays (Table S1 and Fig. S2). The expression constructs in pTRE3G-BI vector coexpresses a C-terminally large NanoLuc fragment LgBiT-fused receptor (Cc-GPR25, Ap-GPR25, or Gg-GPR25) and an N-terminally SmBiT-fused human β-arrestin 2 (SmBiT-ARRB2) for NanoLuc Binary Technology (NanoBiT)-based β-arrestin recruitment assays; the construct PB-TRE/sLgBiT-Cc-GPR25 encodes an N-terminally secretory LgBiT (sLgBiT)-fused Cc-GPR25 for NanoBiT-based ligand−receptor binding assays; the construct PB-TRE/Cc-GPR25 encodes an untagged receptor for chemotaxis assays.

### 2.3. The NanoBiT-based β-arrestin recruitment assays

The NanoBiT-based β-arrestin recruitment assays for Cc-GPR25, Ap-GPR25, and Gg-GPR25 were conducted on transiently transfected human embryonic kidney (HEK) 293T cells according to our previous procedure developed for other receptors [24,29−32]. After cotransfection with the β-arrestin recruitment assay construct and the expression control vector pCMV-Tet3G (Clontech, Mountain View, CA, USA), HEK293T cells were seeded into white opaque 96-well plates and cultured in complete Dulbecco’s modified Eagle medium (DMEM) containing 1.0 ng/mL of Dox for ∼24 h to ∼90% confluence. To conduct the activation assays, the induction medium was removed, and NanoLuc substrate-containing activation solution (serum-free DMEM plus 1% bovine serum albumin) was added (40 μL/well, containing 1.0 μL of substrate stock from Promega, Madison, WI, USA). Thereafter, bioluminescence data were collected on a SpectraMax iD3 plate reader (Molecular Devices, Sunnyvale, CA, USA) for ∼4 min with an interval of 11 s. Thereafter, indicated CXCL17 proteins were added (10 μL/well, diluted in activation solution), and bioluminescence data were continuously collected for ∼10 min. The measured absolute bioluminescence data were corrected for inter well variability by forcing all curves after addition of NanoLuc substrate to same level and plotted using the SigmaPlot 10.0 software (SYSTAT software, Chicago, IL, USA). To obtain the dose-response curve, the highest bioluminescence data were plotted with the agonist concentrations using the SigmaPlot 10.0 software (SYSTAT software,).

### 2.4. The NanoBiT-based ligand−receptor binding assays

The NanoBiT-based binding assays for Cc-GPR25 were conducted on transiently transfected HEK293T cells using recombinant 6×His-SmBiT-Cc-CXCL17 as a tracer according to our previous procedure developed for other receptors [29−32]. After transient transfection with the expression construct PB-TRE/sLgBiT-Cc-GPR25 with or without cotransfection with the tyrosylprotein sulfotransferase expression construct pTRE3G-BI/TPST1:TPST2, HEK293T cells were seeded into white opaque 96-well plates and cultured in complete DMEM containing 20 ng/mL of Dox for ∼24 h to ∼90% confluence. To conduct the binding assays, the induction medium was removed, and pre-warmed binding solution (serum-free DMEM plus 0.1% bovine serum albumin and 0.01% Tween-20) was added (50 μL/well). The binding solution either contained varied concentrations of the tracer (for saturation binding) or contained a constant concentration of the tracer and varied concentrations of a competitor (for competition binding). After incubation at room temperature for ∼1 h, diluted NanoLuc substrate (30-fold dilution using the binding solution) was added (10 μL/well), and bioluminescence data were immediately measured on a SpectraMax iD3 plate reader (Molecular Devices). The measured bioluminescence data were expressed as mean ± standard deviation (SD, *n* = 3) and fitted to one-site binding model using the SigmaPlot 10.0 software (SYSTAT software).

### 2.5. Chemotaxis assays

The chemotaxis assays were conducted on HEK293T cells transiently transfected with the expression construct PB-TRE/Cc-GPR25 according to our previous procedure developed for other receptors [24,29−32]. After being cultured in complete DMEM containing 1.0 ng/mL of Dox for ∼24 h, the transfected cells were trypsinized, suspended in serum-free DMEM at a density of ∼5×10^5^ cells/mL, and seeded into polyethylene terephthalate membrane (8 μm pore size)-coated permeable transwell inserts that were put into a 24-well plate containing chemotactic agent (wild-type or truncated Cc-CXCL17 diluted in serum-free DMEM plus 0.2% bovine serum albumin, 500 μL/well). After incubation at 37°C for ∼5 h, solution in the inserts was removed and cells on the upper face of the permeable membrane were wiped off using cotton swaps, and cells on the lower face of the permeable membrane were fixed with 4% paraformaldehyde solution, stained with crystal violet solution (Beyotime Technology), and observed under a microscope. The migrated cells were quantitated using the ImageJ software and the results were expressed as mean ± SD (*n* = 3).

## 3. Results

### 3.1. Searching for reptilian and avian CXCL17 orthologs from public databases

In our previous studies, CXCL17 orthologs were identified in diverse fish and amphibian species based on conserved gene synteny (adjacent to *LIPE*), genomic architecture (typically four exons), and characteristic amino acid sequence features, including secretory proteins of fewer than 200 amino acids, 4−8 cysteine residues in mature peptide, and a highly conserved Xaa-Pro-Yaa motif at the C-terminus [29−32]. Using the same strategy, we searched public databases for reptilian and avian CXCL17 orthologs in the present study. A total of 35 putative reptilian CXCL17 orthologs, including 15 annotated and 21 unannotated sequences, were identified in the NCBI Gene database (Table S2). Among the annotated reptilian orthologs, only two sequences (gene symbols: LOC309376218 and LOC121915120) from *Podarcis melisellensis* and *Sceloporus undulatus*, respectively, were annotated as “C-X-C motif chemokine 17-like”. One ortholog from *Anolis carolinensis* (gene symbol: LOC134293725) was incorrectly annotated as “synapsin-2-like”, whereas the remaining annotated sequences were classified as “uncharacterized proteins” (Table S2). The 21 unannotated reptilian CXCL17 orthologs were reconstructed by manually assembling their exon sequences based on RNA-sequencing data available in the NCBI Gene database (Table S2 and Fig. S4−S24).

Using sequence BLAST searches based on these reptilian CXCL17 orthologs, we identified nine additional reptilian homologs, including one sequence (KYO44589) from the American alligator (*Alligator mississippiensis*) (Table S2). The alligator protein KYO44589 was derived from a previous whole-genome shotgun sequence (GenBank: AKHW03000708.1) and is located adjacent to the *ZNF225* (also known as *ZNF420*) gene; however, it lacks an N-terminal signal peptide (Table S2). To verify its identity, we further examined the American alligator reference genome (NC_081838.1) and identified an unannotated gene adjacent to *ZNF420* (LOC106737775) (Fig. S25). Manual assembly of its four exons revealed a full-length CXCL17 ortholog containing an intact N-terminal signal peptide (Table S2 and Fig. S25). The American alligator *CXCL17* gene is located approximately 2.8 Mb away from *LIPE* on chromosome 15, making it difficult to identify solely based on its gene synteny. In the genomic region adjacent to the *zinc finger protein 420-like* gene (LOC102385260) of the Chinese alligator (*Alligator sinensis*), an unannotated full-length CXCL17 ortholog was identified (Table S2 and Fig. S26). In several reptilian species, two CXCL17 isoforms were detected, likely resulting from alternative splicing or gene polymorphisms (Table S2). Collectively, we identified CXCL17 orthologs in 46 reptilian species, including lizards, snakes, turtles, and alligators, indicating that CXCL17 is widely present among extant reptiles.

However, no potential avian CXCL17 orthologs were identified in the NCBI databases using the same search strategy, despite wide presence of GPR25 orthologs in extant bird species (Table S3 and Fig. S27). We also screened secretory avian proteins of fewer than 200 amino acids from the UniProt database, but no convincing CXCL17 candidates were detected. In known *CXCL17* genes, the first exon typically encodes the signal peptide region of approximately 20 residues, whereas the final exon encodes the mature peptide region of approximately 40 residues. Therefore, potential CXCL17-related sequences can be identified from genomic DNA by searching all possible translation frames for an N-terminal fragment with signal peptide characteristics together with a positively charged C-terminal fragment containing an Xaa-Pro-Yaa motif at the end. Based on these criteria, we analyzed genomic regions adjacent to *LIPE* and identified putative *CXCL17* gene relics in 22 extant bird species (Table S4). These avian *CXCL17* gene relics encode a signal peptide-like region together with a positively charged mature peptide fragment containing an LPL or LPI motif at the C-terminus (Table S4). However, they no longer appear to be transcribed or properly spliced according to RNA-sequencing data from the reference genomes and frequently contain frameshift or nonsense mutations (Table S4). In addition, these avian *CXCL17* gene relics are only 300−600 bp in length (Table S4), but functional CXCL17 genes from other vertebrate lineages typically span thousands base pairs in genome and contain three introns, suggesting these *CXCL17* gene relics lost most internal DNA sequence, including most introns and some exons. Consequently, these avian *CXCL17* gene relics are unlikely to produce intact functional CXCL17 proteins, suggesting that the avian lineage has probably lost functional CXCL17 during evolution.

All identified reptilian CXCL17 orthologs contain an N-terminal signal peptide and a conserved Xaa-Pro-Yaa motif at the C-terminus (Fig. 1A and Table S2). Most mature peptides possess six cysteine residues arranged in a characteristic CXC-CXC-C-C pattern. However, these reptilian CXCL17 orthologs exhibit little overall sequence similarity to known mammalian, amphibian, or fish CXCL17 proteins (Fig. 1A and Table S5), which likely explains why most remained previously unrecognized. Phylogenetic analysis further showed that reptilian CXCL17 orthologs from snakes, turtles, alligators, and lizards cluster into four distinct branches (Fig. 1B).

**Fig. 1.**
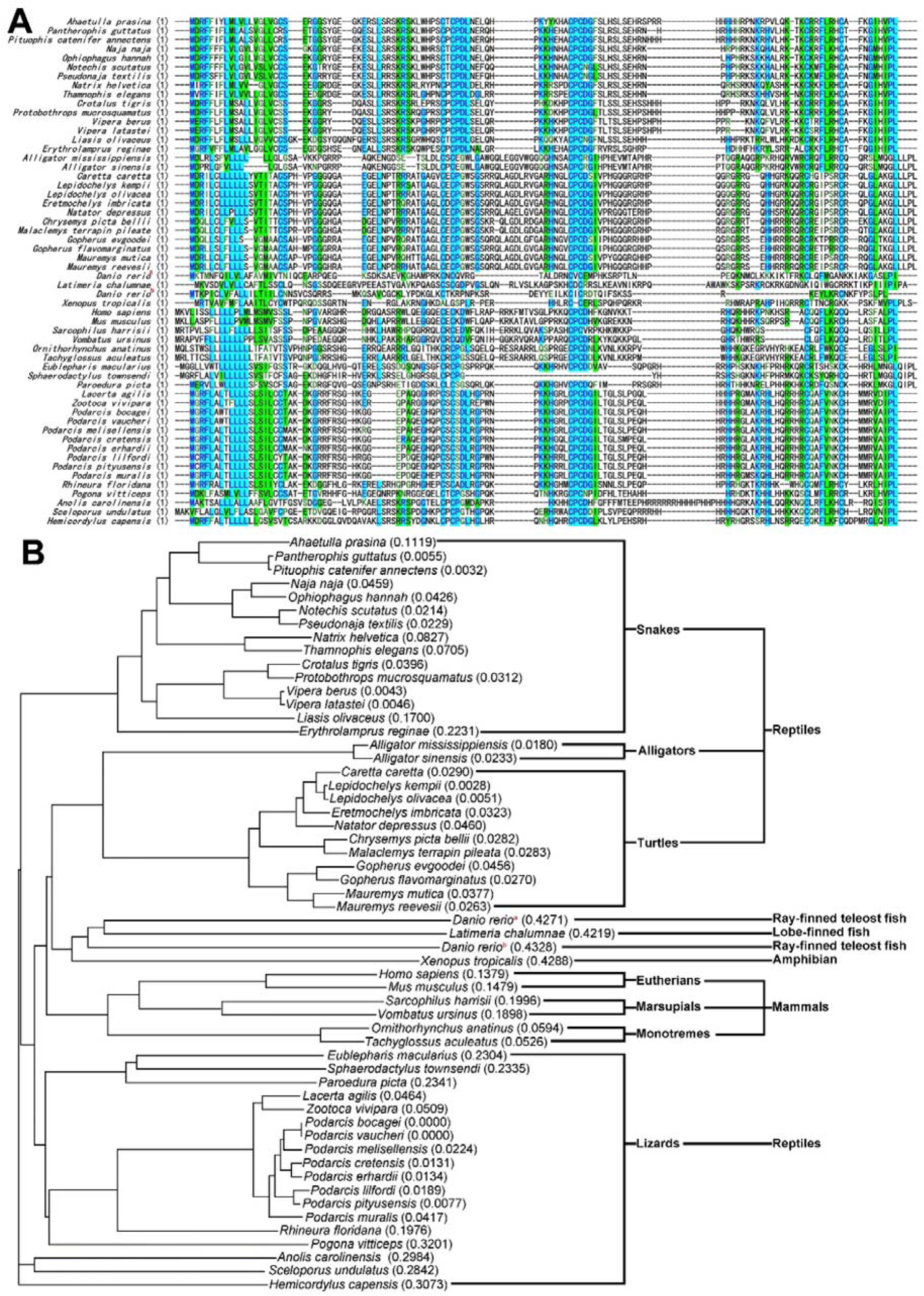
Identification of reptilian CXCL17 orthologs. (**A**) Amino acid sequence alignment of CXCL17 orthologs from reptiles and some other vertebrates. The detailed information is listed in supplementary Table S2 and S5. (**B**) Phylogenetic tree of the CXCL17 orthologs aligned in panel A. These sequences were aligned via AlignX algorithm using the Vector NTI 11.5.1 software. ^a^ zebrafish CXCL17, ^b^ zebrafish CXCL17-like.

Despite their substantial sequence diversity, mature reptilian CXCL17 proteins are consistently highly basic, with calculated pI values of approximately 10 or higher (Table S2). This strong basicity is likely important for their chemoattractant activity, as it may facilitate binding to negatively charged cell-surface glycosaminoglycans and promote the formation of local concentration gradients required for immune-cell recruitment. In addition, the highly basic nature of these proteins may contribute to their interaction with GPR25 through electrostatic interactions with negatively charged residues within the extracellular domain of the receptor [25].

### 3.2. Preparation of Cc-CXCL17 by bacterial overexpression

To obtain recombinant Cc-CXCL17 for functional assays, bacterial overexpression was performed. Following induction of transformed *E. coli* cells with isopropyl-β-D-thiogalactopyranoside (IPTG), a prominent protein band of approximately 12 kDa was detected by SDS-PAGE, indicating successful overexpression of 6×His-Cc-CXCL17 (Fig. 2A). After solubilization of inclusion bodies using an *S*-sulfonation strategy, 6×His-Cc-CXCL17 was purified by immobilized metal ion affinity chromatography. SDS-PAGE analysis of the eluted fractions revealed a predominant monomeric band (indicated by an asterisk) together with several weaker oligomeric bands (Fig. 2A), suggesting that Cc-CXCL17 is prone to intermolecular cross-linking, a phenomenon previously observed for other CXCL17 orthologs [24,29−32].

**Fig. 2.**
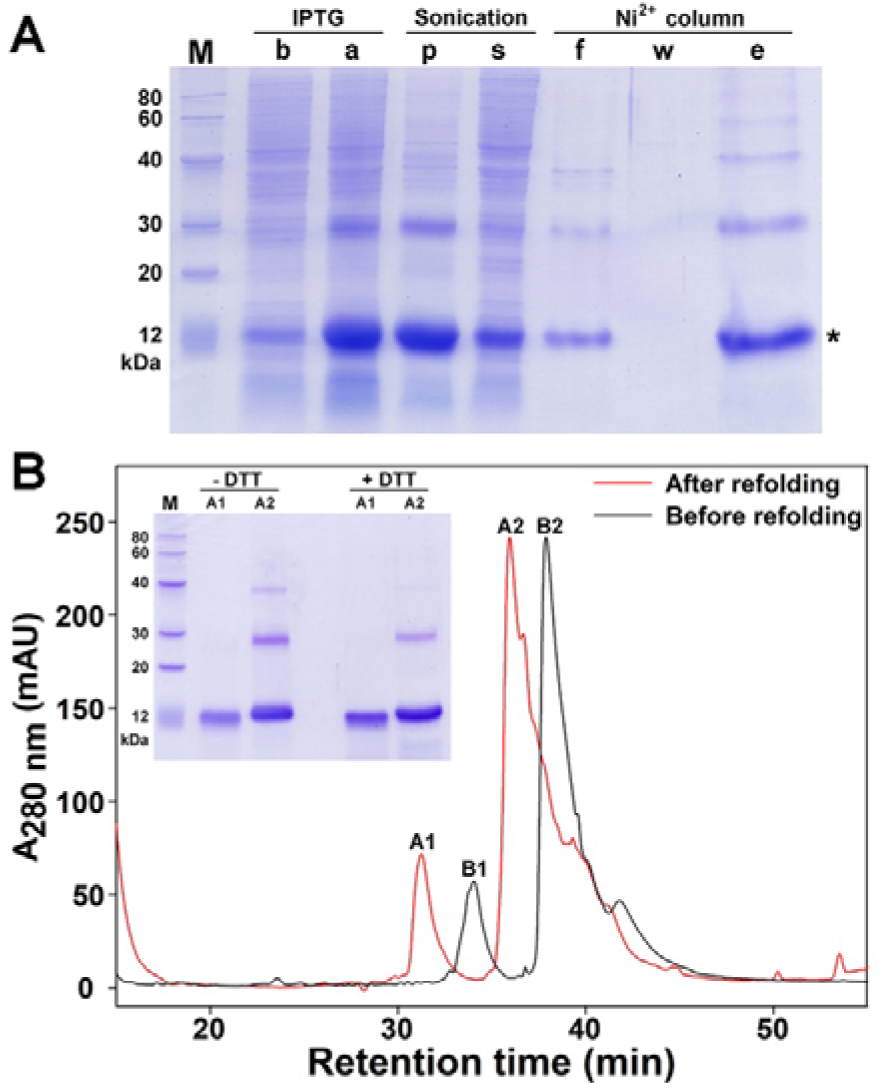
Preparation of Cc-CXCL17 via bacterial overexpression. (**A**) SDS-PAGE analysis of the recombinant 6×His-Cc-CXCL17. Samples were analyzed by SDS-PAGE and visualized by Coomassie brilliant blue R250 staining after electrophoresis. Lane (M), protein ladder; lane (b), before IPTG induction; lane (a), after IPTG induction; lane (p), pellet after sonication; lane (s), supernatant after sonication; lane (f), flowthrough from the Ni^2+^ column; lane (w), washing fraction by 30 mM imidazole; lane (e), eluted fraction by 250 mM imidazole. Band of the monomeric 6×His-Cc-CXCL17 is indicated by an asterisk. (**B**) HPLC analysis of the recombinant 6×His-Cc-CXCL17. The eluted fraction from the Ni^2+^ column was subjected to HPLC analysis either before or after *in vitro* refolding. **Inner panel**, SDS-PAGE analysis of the refolded samples either with or without dithiothreitol (DTT) treatment. The eluted peak A1 and A2 were manually collected, lyophilized, and analyzed by SDS-PAGE.

Subsequently, 6×His-Cc-CXCL17 was subjected to *in vitro* refolding to facilitate the formation of intramolecular disulfide bonds. HPLC analysis using a C_18_ reverse-phase column revealed two elution peaks both before and after the refolding procedure (Fig. 2B). SDS-PAGE analysis further showed that the minor peak, A1, migrated slightly faster than the major peak, A2 (Fig. 2B, inner panel), suggesting that A1 was likely a degradation product. Therefore, the major peak A2 was selected for subsequent activity assays, despite containing a small proportion of dimeric and trimeric species (Fig. 2B, inner panel).

### 3.3. Efficient activation of Cc-GPR25 by recombinant Cc-CXCL17

To determine whether Cc-CXCL17 can activate its corresponding receptor, Cc-GPR25, we established a NanoBiT-based β-arrestin recruitment assay by coexpressing a C-terminally LgBiT-fused Cc-GPR25 (Cc-GPR25-LgBiT) and an N-terminally SmBiT-fused human β-arrestin 2 (SmBiT-ARRB2) in transiently transfected HEK293T cells. Upon agonist stimulation, activated Cc-GPR25-LgBiT recruits SmBiT-ARRB2, resulting in increased bioluminescence through proximity-induced complementation between SmBiT and LgBiT.

Following sequential addition of NanoLuc substrate and recombinant 6×His-Cc-CXCL17 to living HEK293T cells coexpressing Cc-GPR25-LgBiT and SmBiT-ARRB2, bioluminescence increased rapidly in a dose-dependent manner (Fig. 3A), with a calculated EC_50_ value of approximately 56 nM (Fig. 3A, inner panel). In contrast, deletion of the three C-terminal residues markedly reduced activity, as the truncated mutant 6×His-Cc-[desC3]CXCL17 produced only minimal responses even at concentrations up to 1000 nM (Fig. 3B). These results demonstrate that recombinant Cc-CXCL17 efficiently activates Cc-GPR25 in a C-terminal fragment-dependent manner, supporting the existence of a functional CXCL17−GPR25 signaling system in the loggerhead turtle and other reptilian species.

**Fig. 3.**
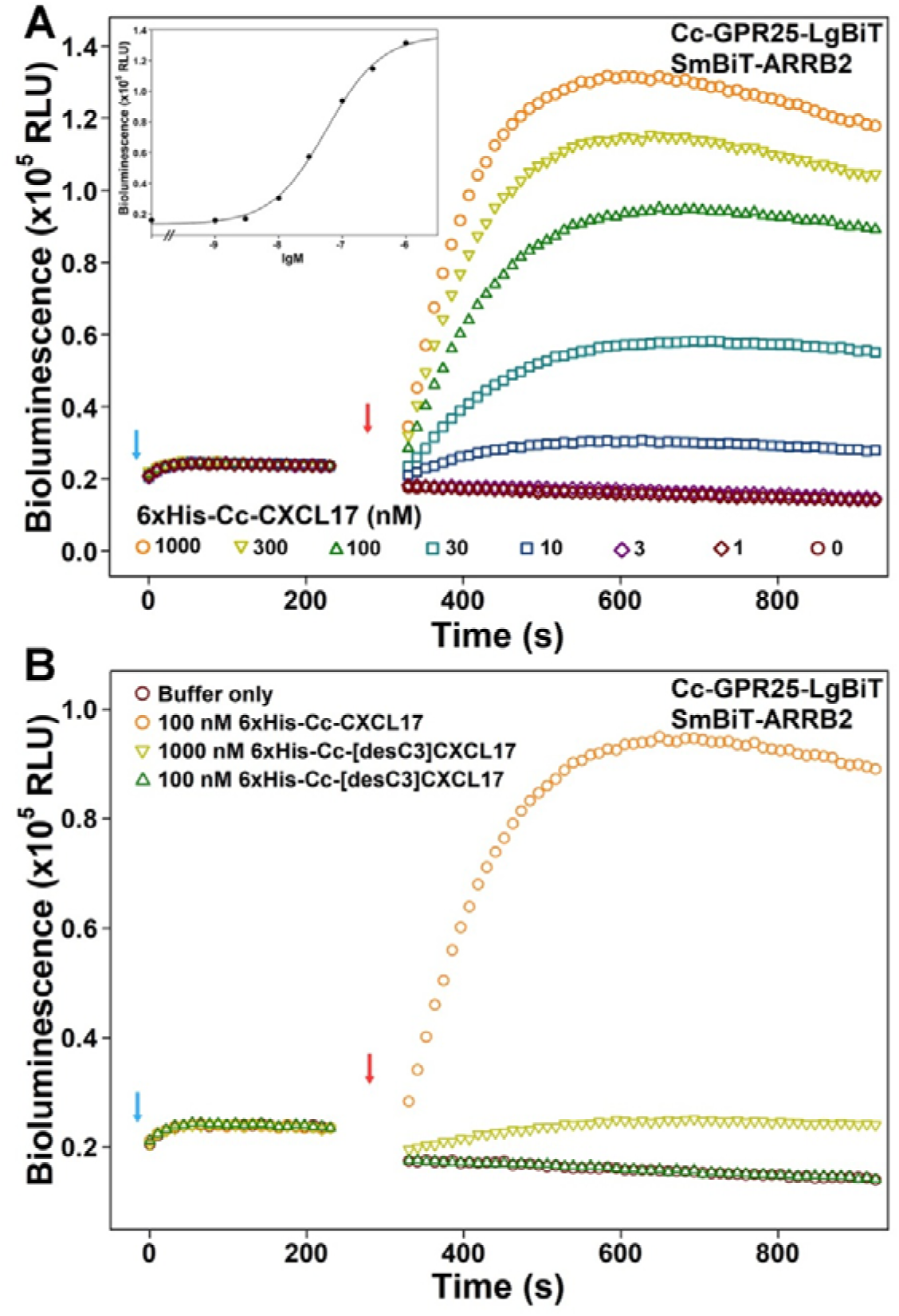
NanoBiT-based β-arrestin recruitment assays for Cc-GPR25. (**A**) Bioluminescence change induced by recombinant 6×His-Cc-CXCL17. **Inner panel**, dose response curve. (**B**) Bioluminescence change induced by the C-terminally truncated mutant. In these NanoBiT-based β-arrestin recruitment assays, NanoLuc substrate and recombinant wild-type or mutant Cc-CXCL17 were sequentially added to living HEK293T cells coexpressing Cc-GPR25-LgBiT and SmBiT-ARRB2, and bioluminescence was continuously measured on a plate reader. Blue arrows indicate the addition of NanoLuc substrate, and red arrows indicate the addition of recombinant peptide.

### 3.4. Direct binding of recombinant Cc-CXCL17 to Cc-GPR25

To examine the direct interaction between Cc-CXCL17 and its corresponding receptor Cc-GPR25, we established a NanoBiT-based homogeneous binding assay by fusing sLgBiT tag to the extracellular N-terminus of Cc-GPR25 and a low-affinity SmBiT tag to the N-terminus of Cc-CXCL17. Upon ligand–receptor binding, the proximity effect between SmBiT and sLgBiT induces NanoBiT complementation and emit bioluminescence.

Recombinant 6×His-SmBiT-Cc-CXCL17 retained the ability to activate Cc-GPR25 in the NanoBiT-based β-arrestin recruitment assay (Fig. 4A); however, its measured EC_50_ value (∼230 nM) was approximately fourfold higher than that of 6×His-Cc-CXCL17 (∼56 nM), suggesting that the SmBiT tag exerted a modest detrimental effect on activity. Addition of 6×His-SmBiT-Cc-CXCL17 to living HEK293T cells overexpressing sLgBiT-Cc-GPR25 induced a concentration-dependent increase in bioluminescence, although the resulting binding curve was nearly linear. The bioluminescence signal was markedly reduced in the presence of 1.0 μM 6×His-Cc-CXCL17 (Fig. 4B), indicating that the binding affinity between the tagged ligand and receptor was relatively low.

**Fig. 4.**
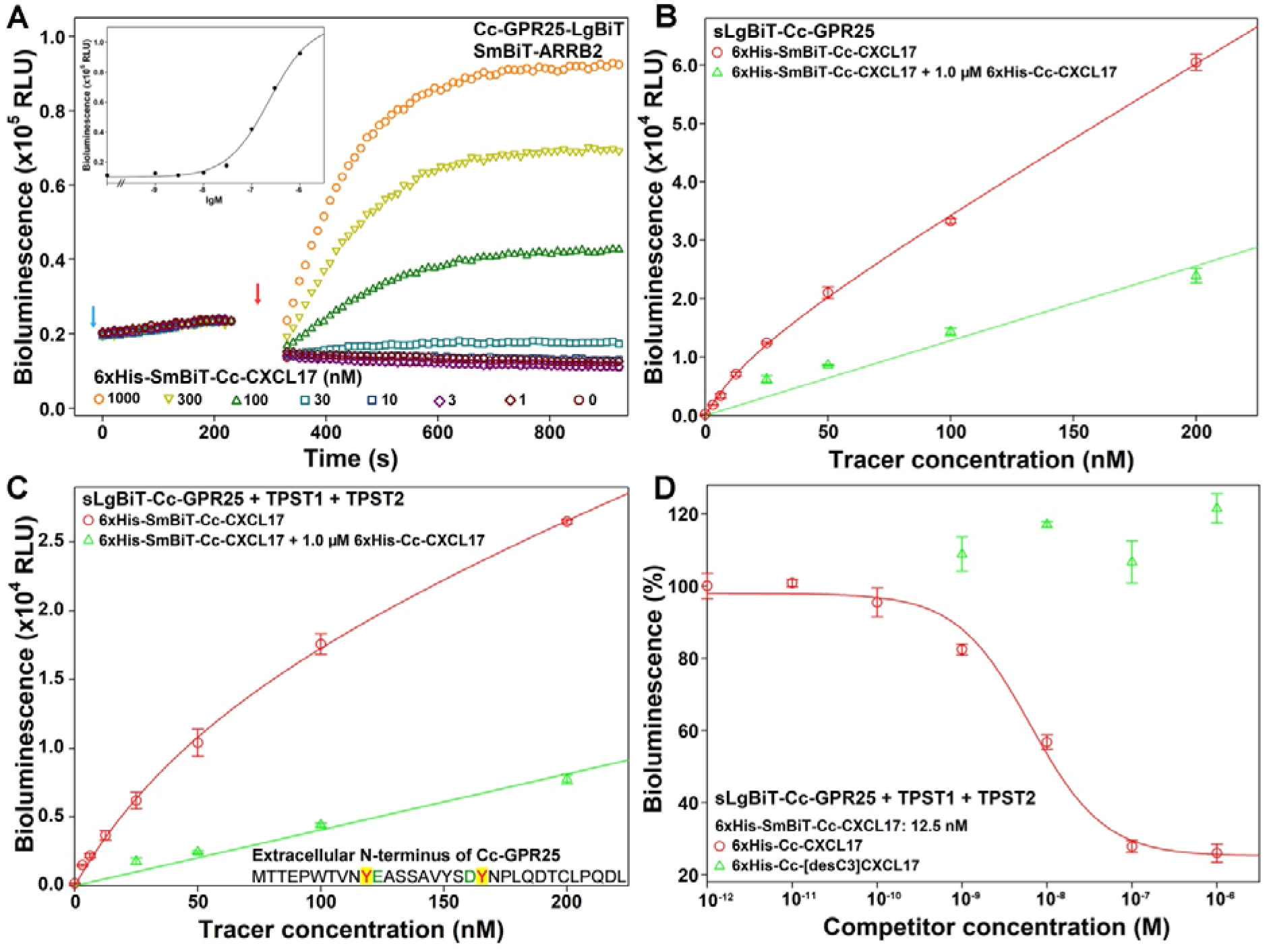
NanoBiT-based ligand-binding assays for Cc-GPR25. (**A**) Activity measurement of the recombinant 6×His-SmBiT-Cc-CXCL17 tracer via the NanoBiT-based β-arrestin recruitment assay. Blue arrow indicates the addition of NanoLuc substrate, and red arrow indicates the addition of tracer. **Inner panel**, dose response curve. (**B,C**) The NanoBiT-based saturation binding assays without (B) or with (C) coexpression of human tyrosylprotein sulfotransferases. (**D**) The NanoBiT-based competition binding assays. In these NanoBiT-based binding assays, the recombinant 6×His-SmBiT-Cc-CXCL17 tracer with or without competitor was added to living HEK293T cells overexpressing sLgBiT-Cc-GPR25. After incubation, NanoLuc substrate was added and bioluminescence was immediately measured on a plate reader. The measured bioluminescence data are expressed as mean ± SD (*n* = 3) and plotted using the SigmaPlot10.0 software. Total binding data (red circles) in panel B,C were fitted with the function of Y = B_max_X/(K_d_+X) + k_non_X, the non-specific binding data (green triangles) in panel B,C were fitted with linear curves, the competition binding data in panel D were fitted with sigmoidal curves using the SigmaPlot10.0 software.

The extracellular N-terminal region of Cc-GPR25 contains two tyrosine (Y) residues predicted to undergo sulfation because they are adjacent to negatively charged residues (Fig. 4C, inner panel). Following coexpression of the tyrosylprotein sulfotransferases TPST1 and TPST2, the binding curve became predominantly hyperbolic (Fig. 4C), yielding a calculated K_d_ value of 78 ± 25 nM (*n* = 3), suggesting that tyrosine sulfation enhances the interaction between Cc-GPR25 and Cc-CXCL17. In competition binding assays, 6×His-Cc-CXCL17 reduced bioluminescence in a sigmoidal manner (Fig. 4D), with a calculated IC_50_ value of 6.3 ± 0.8 nM (*n* = 3). In contrast, the C-terminally truncated mutant 6×His-Cc-[desC3]CXCL17 showed no detectable inhibitory effect, indicating loss of binding to Cc-GPR25. These NanoBiT-based assays demonstrate direct binding of Cc-CXCL17 to Cc-GPR25 in a C-terminal fragment-dependent manner.

### 3.5. Chemotactic migration of transfected HEK293T cells induced by Cc-GPR25 activation

To evaluate the cellular response triggered by Cc-GPR25 activation, we performed chemotaxis assays using transiently transfected HEK293T cells. In HEK293T cells expressing Cc-GPR25, recombinant 6×His-Cc-CXCL17 induced cell migration in a dose-dependent manner, with significant chemotactic activity observed at concentrations as low as 10 nM (Fig. 5A,B). In contrast, the C-terminally truncated mutant 6×His-Cc-[desC3]CXCL17 exhibited only minimal activity even at concentrations up to 1000 nM (Fig. 5A,B). Moreover, 6×His-Cc-CXCL17 produced little effect on non-transfected HEK293T cells, even at the concentration of 1000 nM (Fig. 5A). These findings indicate that activation of Cc-GPR25 by Cc-CXCL17 promotes chemotactic migration of transfected cells and further support the presence of a functional CXCL17−GPR25 signaling system in reptiles.

**Fig. 5.**
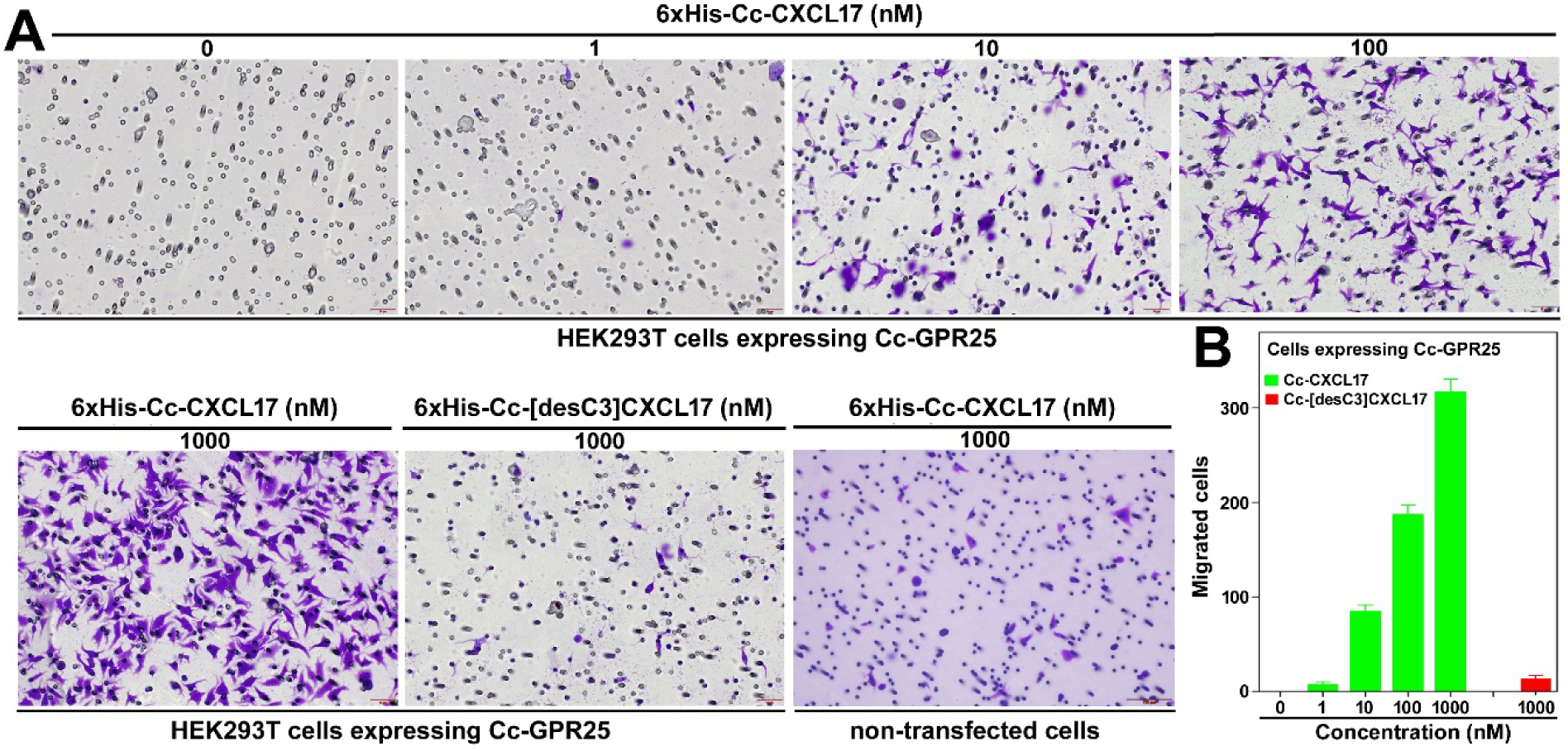
Chemotaxis assays of non-transfected or transfected HEK293T cells expressing Cc-GPR25. (**A**) Representative images of the migrated HEK293T cells induced by wild-type or C-terminally truncated Cc-CXCL17. The scale bar in the image of non-transfected cells is 100 μm, and those in the images of transfected cells is 50 μm. (**B**) Quantitative analysis of the migrated HEK293T cells overexpressing Cc-GPR25. For the chemotaxis assays, non-transfected or transfected HEK293T cells were seeded into permeable membrane-coated inserts, and induced by chemotactic solution containing indicated concentrations of wild-type or mutant Cc-CXCL17. After being cultured at 37°C for ∼5 h, cells on the upper face of the permeable membrane were wiped off, and cells on the lower face were fixed, stained, and observed under a microscope. Three images were analyzed using the ImageJ software and the calculated cell numbers are expressed as mean ± SD (*n* = 3).

### 3.6. Cross-species activity of CXCL17 orthologs toward reptilian and avian GPR25 receptors

The cross-species activities of several recombinant CXCL17 orthologs (Fig. 6A) toward the reptilian receptor Cc-GPR25 and two avian GPR25 orthologs, Ap-GPR25 from mallard (*Anas platyrhynchos*) and Gg-GPR25 from chicken (*Gallus gallus*), were evaluated using NanoBiT-based β-arrestin recruitment assays. Recombinant Cc-CXCL17 could activate Cc-GPR25 efficiently, but CXCL17s from other species had almost no activity toward this reptilian receptor (Fig. 6B). For the mallard receptor Ap-GPR25, only Cc-CXCL17 and human CXCL17 (Hs-CXCL17) produced a weak but detectable response (Fig. 6C). No measurable activation was observed for the chicken receptor Gg-GPR25 with any of the tested CXCL17 orthologs (Fig. 6D). These results indicate that CXCL17 orthologs exhibit substantial receptor selectivity across different vertebrate species.

**Fig. 6.**
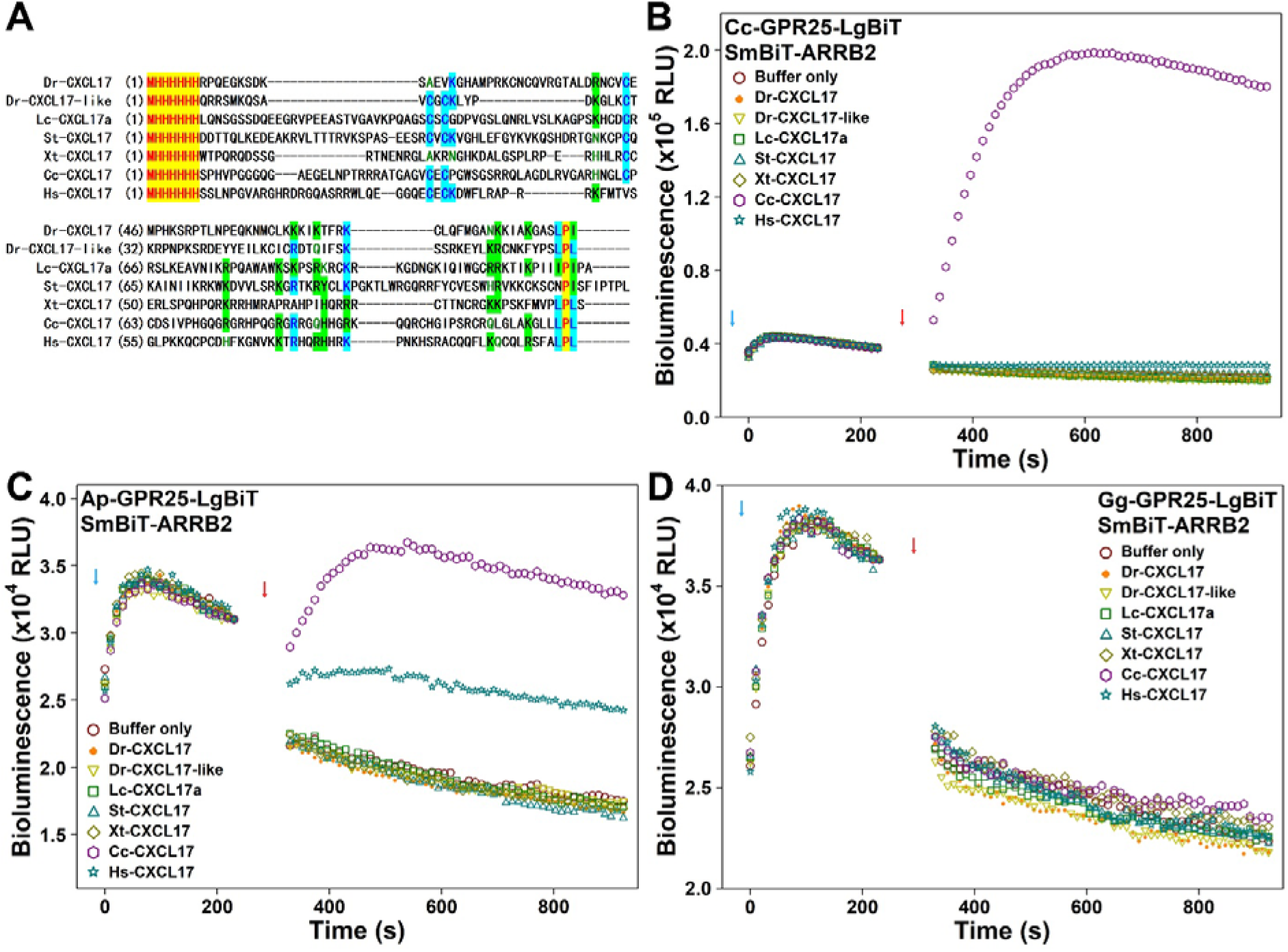
Cross-species activity of CXCL17 orthologs towards reptilian or avian GPR25s. (**A**) Amino acid sequence alignment of recombinant CXCL17s used for cross-species activity assays. (**B**) Cross-species activity of CXCL17s towards reptilian Cc-GPR25 measured via the NanoBiT-based β-arrestin recruitment assays. (**C,D**) Cross-species activity of CXCL17s towards avian Ap-GPR25 (C) and Gg-GPR25 (D) measured via the NanoBiT-based β-arrestin recruitment assays. In these assays, NanoLuc substrate and 1.0 μM of indicated CXCL17s were sequentially added to living HEK293T cells coexpressing indicated C-terminally LgBiT-fused GPR25 and SmBiT-ARRB2, and bioluminescence was continuously measured on a plate reader. Blue arrows indicate the addition of NanoLuc substrate, and red arrows indicate the addition of recombinant peptide.

## 4. Discussion

In the present study, we identified CXCL17 orthologs from multiple reptilian species and demonstrated the functionality of the representative ortholog Cc-CXCL17 using cell-based binding and receptor activation assays. According to the NCBI database, GPR25 orthologs are widely present in various reptiles (Table S3) and display quite high amino acid sequence homology (Fig. S27). Thus, it is reasonable to deduce that functional CXCL17−GPR25 signaling system is widely present in extant reptiles. Together with our previous findings [29−32], these results suggest that the CXCL17−GPR25 signaling axis originated in ancient fish ancestors and has been conserved throughout the evolution of extant fishes, amphibians, reptiles, and mammals, implying important biological roles across these vertebrate lineages.

In contrast, we were unable to identify intact avian CXCL17 orthologs in public databases based on either conserved gene synteny or characteristic amino acid sequence features. Transcriptomic sequencing of mallard and chicken lung tissues also failed to detect any potential CXCL17 candidates in the assembled datasets (our unpublished data). Nevertheless, we identified putative *CXCL17* gene relics in 22 extant bird species within the NCBI Gene database (Table S4). These relics appear to be transcriptionally inactive and frequently contain frameshift or nonsense mutations, indicating that they are unlikely to encode functional CXCL17 proteins. These findings suggest that the avian lineage has probably lost CXCL17 during evolution. According to a previous study, the avian lineage has lost hundreds of protein coding genes that are present in most other vertebrate lineages [33]. *CXCL17* is probably another missing gene in the avian lineage. However, GPR25 orthologs are present in most bird species according to the NCBI database (Table S3). Most avian GPR25 orthologs display highly sequence homology, but the orthologs from chicken and red-crested turaco (*Tauraco erythrolophus*) are quite different from other avian GPR25 orthologs (Fig. S27). In future, more studies will be required to definitively determine the evolutionary status and potential remnants of CXCL17 in the avian lineage.

CXCL17 orthologs from mammals, reptiles, amphibians, and fishes exhibit remarkable sequence diversity; however, they share several conserved characteristics, including an N-terminal signal peptide, a C-terminal Xaa-Pro-Yaa motif (where Xaa and Yaa are typically large aliphatic residues such as Leu, Ile, Met, Val, or Thr), a highly basic mature peptide with pI values generally around or above 10, an overall length of fewer than 200 amino acids, and the presence of 4−8 cysteine residues within the mature peptide region. These conserved features provide valuable criteria for the identification of additional CXCL17 orthologs in future studies. Moreover, CXCL17 orthologs display relatively conserved gene synteny, often located adjacent to *LIPE*, as well as a conserved genomic architecture consisting typically of four exons and occasionally five exons. These genomic characteristics may further facilitate the identification of previously unrecognized CXCL17 orthologs.

The conserved C-terminal Xaa-Pro-Yaa motif appears to be critical for the binding and activation of GPR25 by CXCL17 orthologs. In AlphaFold3-predicted binding models, the relatively conserved C-terminal region of CXCL17 consistently inserts into the orthosteric ligand-binding pocket of the corresponding GPR25 receptor with ipTM values of approximately 0.6. In contrast, the long N-terminal region of CXCL17 orthologs is highly variable in sequence but consistently enriched in positively charged Arg (R) and Lys (K) residues, which likely mediate electrostatic interactions with negatively charged residues in the extracellular N-terminal domain of GPR25 [25]. Further structural studies at high resolution will be necessary to elucidate the precise molecular mechanism underlying CXCL17–GPR25 recognition and activation.

## CRediT authorship contribution statement

Jie Yu: Investigation, Methodology, Visualization. Hao-Zheng Li: Investigation, Methodology. Juan-Juan Wang: Investigation, Methodology. Ya-Li Liu: Formal analysis, Project administration. Zhan-Yun Guo: Supervision, Conceptualization, Writing - Original Draft, Writing - Review & Editing, Funding acquisition.

## Acknowledgments

This work was supported by grant from the National Natural Science Foundation of China (31971193).

## Appendix A. Supplementary data

Supplementary data to this article can be found online

## Data availability

The data of this study are available in this manuscript, as well as the associated supplementary information.

## References

[1] M.T. Pisabarro, B. Leung, M. Kwong, R. Corpuz, G.D. Frantz, N. Chiang, R. Vandlen, L.J. Diehl, N. Skelton, H.S. Kim, D. Eaton, K.N. Schmidt, Cutting edge: novel human dendritic cell- and monocyte-attracting chemokine-like protein identified by fold recognition methods, J. Immunol. 176 (2006) 2069−2073, 10.4049/jimmunol.176.4.2069.

[2] A.M. Burkhardt, K.P. Tai, J.P. Flores-Guiterrez, N. Vilches-Cisneros, K. Kamdar, O. Barbosa-Quintana, R. Valle-Rios, P.A. Hevezi, J. Zuñiga, M. Selman, A.J. Ouellette, A. Zlotnik, CXCL17 is a mucosal chemokine elevated in idiopathic pulmonary fibrosis that exhibits broad antimicrobial activity, J. Immunol. 188 (2012) 6399−6406, 10.4049/jimmunol.1102903.

[3] A.M. Burkhardt, J.L. Maravillas-Montero, C.D. Carnevale, N. Vilches-Cisneros, J.P. Flores, P.A. Hevezi, A. Zlotnik, CXCL17 is a major chemotactic factor for lung macrophages, J. Immunol. 193 (2014) 1468−1474, 10.4049/jimmunol.1400551.

[4] T. Oka, M. Sugaya, N. Takahashi, T. Takahashi, S. Shibata, T. Miyagaki, Y. Asano, S. Sato, CXCL17 Attenuates Imiquimod-Induced Psoriasis-like Skin Inflammation by Recruiting Myeloid-Derived Suppressor Cells and Regulatory T Cells, J. Immunol. 198 (2017) 3897−3908, 10.4049/jimmunol.1601607.

[5] R. Srivastava, M. Hernández-Ruiz, A.A. Khan, M.A. Fouladi, G.J. Kim, V.T. Ly, T. Yamada, C. Lam, S.A.B. Sarain, U. Boldbaatar, A. Zlotnik, E. Bahraoui, L. BenMohamed, CXCL17 Chemokine-Dependent Mobilization of CXCR8(+)CD8(+) Effector Memory and Tissue-Resident Memory T Cells in the Vaginal Mucosa Is Associated with Protection against Genital Herpes, J. Immunol. 200 (2018) 2915−2926, 10.4049/jimmunol.1701474.

[6] M. Hernández-Ruiz, S. Othy, C. Herrera, H.T. Nguyen, G. Arrevillaga-Boni, J. Catalan-Dibene, M.D. Cahalan, A. Zlotnik, Cxcl17(-/-) mice develop exacerbated disease in a T cell-dependent autoimmune model, J. Leukoc. Biol. 105 (2019) 1027−1039, 10.1002/JLB.3A0918-345RR.

[7] E. Lowry, R.C. Chellappa, B. Penaranda, K.V. Sawant, M. Wakamiya, R.P. Garofalo, K. Rajarathnam, CXCL17 is a proinflammatory chemokine and promotes neutrophil trafficking, J. Leukoc. Biol. 115 (2024) 1177−1182, 10.1093/jleuko/qiae028.

[8] Y. Yin, C. Mu, J. Wang, Y. Wang, W. Hu, W. Zhu, X. Yu, W. Hao, Y. Zheng, Q. Li, W. Han, CXCL17 Attenuates Diesel Exhaust Emissions Exposure-Induced Lung Damage by Regulating Macrophage Function, Toxics 11 (2023) 646, 10.3390/toxics11080646.

[9] S.V. Silver, K.J. Tucker, R.E. Vickman, N.A. Lanman, O.J. Semmes, N.S. Alvarez, P. Popovics, Characterization of prostate macrophage heterogeneity, foam cell markers, and CXCL17 upregulation in a mouse model of steroid hormone imbalance, Sci. Rep. 14 (2024) 21029, 10.1038/s41598-024-71137-4.

[10] E.J. Weinstein, R. Head, D.W. Griggs, D. Sun, R.J. Evans, M.L. Swearingen, M.M. Westlin, R. Mazzarella, VCC-1, a novel chemokine, promotes tumor growth, Biochem. Biophys. Res. Commun. 350 (2006) 74−81, 10.1016/j.bbrc.2006.08.194.

[11] N. Hiraoka, R. Yamazaki-Itoh, Y. Ino, Y. Mizuguchi, T. Yamada, S. Hirohashi, Y. Kanai, CXCL17 and ICAM2 are associated with a potential anti-tumor immune response in early intraepithelial stages of human pancreatic carcinogenesis, Gastroenterology 140 (2011) 310−121, 10.1053/j.gastro.2010.10.009.

[12] E. Koni, I. Congur, Z. Tokcaer Keskin, Overexpression of CXCL17 increases migration and invasion of A549 lung adenocarcinoma cells, Front. Pharmacol. 15 (2024) 1306273, 10.3389/fphar.2024.1306273.

[13] B.H. Su, C.T. Wang, J.M. Chang, H.Y. Chen, T.H. Huang, Y.T. Yen, Y.L. Tseng, M.Y. Chang, C.H. Lee, L.H. Cheng, Y.C. Wu, C.L. Wu, P. Ling, A.L. Shiau, OCT4 promotes lung cancer progression through upregulation of VEGF-correlated chemokine-1, Int. J. Med. Sci. 22 (2025) 680−695, 10.7150/ijms.102505.

[14] J. Zhai, T. Pang, Y. An, J. Wang, X. Zhang, L. Zhang, L. Zhang, L. Dong, Clinical significance of CXCL17 in cervical cancer, Medicine (Baltimore) 105 (2026) e46178, 10.1097/MD.0000000000046178.

[15] Z. Ma, M. Dong, H. Pan, J. Xu, T. Luo, W. Xie, M. Xiao, X. Wen, J. Hua, D. Qian, Q. Meng, Y. Wang, D. Wu, X. Zhang, T. Yang, Z. Song, W. Wang, J. Xu, X. Yu, S. Shi, ENPP1-Regulated Extracellular Purine Metabolism Drives Pancreatitis-Mediated Pancreatic Cancer, Gastroenterology 170 (2026) 735−752, 10.1053/j.gastro.2025.09.032.

[16] X. Kong, L. Lei, L. Jin, C. Ren, T. Mi, Q. Wang, D. He, SNRPC promotes chemoresistance in Wilms tumor via the NF-kappaB-CXCL17 axis regulating M2-Type TAMs infiltration and targeted nanotherapy research, J. Exp. Clin. Cancer Res. 45 (2026) 97, 10.1186/s13046-026-03680-z.

[17] J.L. Maravillas-Montero, A.M. Burkhardt, P.A. Hevezi, C.D. Carnevale, M.J. Smit, A. Zlotnik, Cutting edge: GPR35/CXCR8 is the receptor of the mucosal chemokine CXCL17, J. Immunol. 194 (2015) 29−33, 10.4049/jimmunol.1401704.

[18] S.J. Park, S.J. Lee, S.Y. Nam, D.S. Im, GPR35 mediates lodoxamide-induced migration inhibitory response but not CXCL17-induced migration stimulatory response in THP-1 cells; is GPR35 a receptor for CXCL17? Br. J. Pharmacol. 175 (2018) 154−161, 10.1111/bph.14082.

[19] N.A.S. Binti Mohd Amir, A.E. Mackenzie, L. Jenkins, K. Boustani, M.C. Hillier, T. Tsuchiya, G. Milligan, J.E. Pease, Evidence for the Existence of a CXCL17 Receptor Distinct from GPR35, J. Immunol. 201 (2018) 714−724, 10.4049/jimmunol.1700884.

[20] C.W. White, S. Platt, L.E. Kilpatrick, N. Dale, R.S. Abhayawardana, S. Dekkers, N.D. Kindon, B. Kellam, M.J. Stocks, K.D.G. Pfleger, S.J. Hill, CXCL17 is an allosteric inhibitor of CXCR4 through a mechanism of action involving glycosaminoglycans, Sci. Signal. 17 (2024) eabl3758, 10.1126/scisignal.abl3758.

[21] J. Ding, C. Hillig, C.W. White, N.A. Fernandopulle, H. Anderton, J.S. Kern, M.P. Menden, G.A. Mackay, CXCL17 induces activation of human mast cells via MRGPRX2, Allergy 79 (2024) 1609−1612, 10.1111/all.16036.

[22] W.F. Hu, J.J. Wang, J. Yu, Y.L. Liu, Z.G. Xu, Z.Y. Guo, CXCL17 activates three MAS-related G protein-coupled receptors independently of its conserved C-terminal fragment, Arch. Biochem. Biophys. 775 (2026) 110666, 10.1016/j.abb.2025.110666.

[23] B. Ocón, M. Xiang, Y. Bi, S. Tan, K. Brulois, A. Ayesha, M. Kunte, C. Zhou, M. LaJevic, N. Lazarus, F. Mengoni, T. Sharma, S. Montgomery, J.E. Hooper, M. Huang, T. Handel, J.R.D. Dawson, I. Kufareva, B.A. Zabel, J. Pan, E.C. Butcher, A lymphocyte chemoaffinity axis for lung, non-intestinal mucosae and CNS, Nature 635 (2024) 736−745, 10.1038/s41586-024-08043-2.

[24] W.F. Hu, J. Yu, J.J. Wang, R.J. Sun, Y.S. Zheng, T. Zhang, Y.L. Liu, Z.G. Xu, Z.Y. Guo, Identification of orphan GPR25 as a receptor for the chemokine CXCL17, FEBS J. 292 (2025) 5998−6015, 10.1111/febs.70117.

[25] W. Yang, S.P. Giblin, J.E. Pease, Evidence for a Two-Step Model for Activation of GPR25 by the Chemoattractant CXCL17, Basic Clin. Pharmacol. Toxicol. 139 (2026) e70255, 10.1111/bcpt.70255.

[26] S.S. Denisov, CXCL17: The Black Sheep in the Chemokine Flock, Front. Immunol. 12 (2021) 712897, 10.3389/fimmu.2021.712897.

[27] S.P Giblin, J.E. Pease, What defines a chemokine? - The curious case of CXCL17, Cytokine 168 (2023) 156224, 10.1016/j.cyto.2023.156224.

[28] A. Aleotti, M. Goulty, C. Lewis, F. Giorgini, R. Feuda, The origin, evolution, and molecular diversity of the chemokine system, Life Sci. Alliance 7 (2024) e202302471, 10.26508/lsa.202302471.

[29] J. Yu, W.F. Hu, J.J. Wang, Y.L. Liu, Z.G. Xu, Z.Y. Guo, Identification and functional characterization of two CXCL17 paralogs from zebrafish, Biochem. J. 483 (2026) 135−147, 10.1042/BCJ20253418.

[30] J. Yu, J.J. Wang, W.F. Hu, Y.L. Liu, Z.G. Xu, Z.Y. Guo, Ancestral Origin of the CXCL17–GPR25 System Traced to the Lobe-Finned Fish *Latimeria chalumnae*, Biochimie 245 (2026) 168−177, 10.1016/j.biochi.2026.03.011.

[31] J. Yu, H.Z. Li, J.J. Wang, J.J. Yao, W.F. Hu, Y.L. Liu, Z.Y. Guo, Identification and functional characterization of CXCL17 orthologs in amphibians, Arch. Biochem. Biophys. 780 (2026) 110797, 10.1016/j.abb.2026.110797.

[32] J. Yu, J.J. Wang, H.Z. Li, Y.L. Liu, Z.Y. Guo, Identification and functional characterization of CXCL17 in cartilaginous fishes reveals an ancient origin of the CXCL17-GPR25 signaling pathway, Preprint at BioRxiv, https://www.biorxiv.org/cgi/content/short/2026.03.04.709523v1

[33] P.V. Lovell, M. Wirthlin, L. Wilhelm, P. Minx, N.H. Lazar, L. Carbone, W.C. Warren, C.V. Mello, Conserved syntenic clusters of protein coding genes are missing in birds, Genome Biol. 15 (2014) 565, 10.1186/s13059-014-0565-1.

