## Supplemental Table S1-S5 and Fig. S1-S27 for "Evolutionary conservation and divergence of CXCL17 orthologs: functional evidence in reptiles and loss in the avian lineage"

**Contents**

**Table S1.** Information for generating expression constructs for reptilian and avian GPR25s.
**Table S2.** Reptilian CXCL17 orthologs identified in present study.
**Table S3.** Avian and reptilian and GPR25 orthologs.
**Table S4.** Possible *CXCL17* gene relics nearby *LIPE* gene in genome of some bird species.
**Table S5.** Other CXCL17 orthologs used in this study.
**Fig. S1.** The nucleotide and amino acid sequence of Cc-CXCL17 proteins in pET expression vector.
**Fig. S2.** The nucleotide and amino acid sequence of Cc-GPR25, Ap-GPR25, and Gg-GPR25 in different expression constructs.
**Fig. S3.** Amino acid sequence alignment of chicken GPR25 with other GPR25 orthologs from mammals, birds, reptiles, amphibians, and fishes.
**Fig. S4.** Gene synteny and architecture (**A**), genomic DNA sequence (**B**), cDNA sequence (**C**), and encoded protein sequence (**D**) of the unannotated *cxcl17* gene in the NCBI reference genome of *Ahaetulla prasina*.
**Fig. S5.** Gene synteny and architecture (**A**), genomic DNA sequence (**B**), cDNA sequence (**C**), and encoded protein sequence (**D**) of the unannotated *cxcl17* gene in the NCBI reference genome of *Crotalus tigris* (Tiger rattlesnake).
**Fig. S6.** Gene synteny and architecture (**A**), genomic DNA sequence (**B**), cDNA sequence (**C**), and encoded protein sequence (**D**) of the unannotated *cxcl17* gene in the NCBI reference genome of *Erythrolamprus reginae* (royal ground snake).
**Fig. S7.** Gene synteny and architecture (**A**), genomic DNA sequence (**B**), cDNA sequence (**C**), and encoded protein sequence (**D**) of the unannotated *cxcl17* gene in the NCBI reference genome of *Notechis scutatus* (mainland tiger snake).

**Fig. S8.** Gene synteny and architecture (**A**), genomic DNA sequence (**B**), cDNA sequence (**C**), and encoded protein sequence (**D**) of the unannotated *cxcl17* gene in the NCBI reference genome of *Pantherophis guttatus*.

**Fig. S9.** Gene synteny and architecture (**A**), genomic DNA sequence (**B**), cDNA sequence (**C**), and encoded protein sequence (**D**) of the unannotated *cxcl17* gene in the NCBI reference genome of *Pituophis catenifer annectens*.

**Fig. S10.** Gene synteny and architecture (**A**), genomic DNA sequence (**B**), cDNA sequence (**C**), and encoded protein sequence (**D**) of the unannotated *cxcl17* gene in the NCBI reference genome of *Protobothrops mucrosquamatus* (brown spotted pit viper).

**Fig. S11.** Gene synteny and architecture (**A**), genomic DNA sequence (**B**), cDNA sequence (**C**), and encoded protein sequence (**D**) of the unannotated *cxcl17* gene in the NCBI reference genome of *Pseudonaja textilis*.

**Fig. S12.** Gene synteny and architecture (**A**), genomic DNA sequence (**B**), cDNA sequence (**C**), and encoded protein sequence (**D**) of the unannotated *cxcl17* gene in the NCBI reference genome of *Thamnophis elegans* (Western terrestrial garter snake).

**Fig. S13.** Gene synteny and architecture (**A**), genomic DNA sequence (**B**), cDNA sequence (**C**), and encoded protein sequence (**D**) of the unannotated *cxcl17* gene in the NCBI reference genome of *Eublepharis macularius*.

**Fig. S14.** Gene synteny and architecture (**A**), genomic DNA sequence (**B**), cDNA sequence (**C**), and encoded protein sequence (**D**) of the unannotated *cxcl17* gene in the NCBI reference genome of *Hemicordylus capensis* (graceful crag lizard).

**Fig. S15.** Gene synteny and architecture (**A**), genomic DNA sequence (**B**), cDNA sequence (**C**), and encoded protein sequence (**D**) of the unannotated *cxcl17* gene in the NCBI reference genome of *Lacerta agilis* (Sand lizard).

**Fig. S16.** Gene synteny and architecture (**A**), genomic DNA sequence (**B**), cDNA sequence (**C**), and encoded protein sequence (**D**) of the unannotated *cxcl17* gene in the NCBI reference genome of *Rhineura floridana*.

**Fig. S17.** Gene synteny and architecture (**A**), genomic DNA sequence (**B**), cDNA sequence (**C**), and encoded protein sequence (**D**) of the unannotated *cxcl17* gene in the NCBI reference genome of *Sphaerodactylus townsendi*.

**Fig. S18.** Gene synteny and architecture (A), genomic DNA sequence (B), cDNA sequence (C), and encoded protein sequence (D) of the unannotated *cxcl17* gene in the NCBI reference genome of *Zootoca vivipara* (common lizard).

**Fig. S19.** Gene synteny and architecture (**A**), genomic DNA sequence (**B**), cDNA sequence (**C**), and encoded protein sequence (**D**) of the unannotated *cxcl17* gene in the NCBI reference genome of *Chrysemys picta bellii* (western painted turtle).

**Fig. S20.** Gene synteny and architecture (**A**), genomic DNA sequence (**B**), cDNA sequence (**C**), and encoded protein sequence (**D**) of the unannotated *cxcl17* gene in the NCBI reference genome of *Gopherus evgoodei* (Goodes thornscrub tortoise).

**Fig. S21.** Gene synteny and architecture (**A**), genomic DNA sequence (**B**), cDNA sequence (**C**), and encoded protein sequence (**D**) of the unannotated *cxcl17* gene in the NCBI reference genome of *Gopherus flavomarginatus* (Mexican gopher tortoise).

**Fig. S22.** Gene synteny and architecture (**A**), genomic DNA sequence (**B**), cDNA sequence (**C**), and encoded protein sequence (**D**) of the unannotated *cxcl17* gene in the NCBI reference genome of *Mauremys mutica* (yellowpond turtle).

**Fig. S23.** Gene synteny and architecture (**A**), genomic DNA sequence (**B**), cDNA sequence (**C**), and encoded protein sequence (**D**) of the unannotated *cxcl17* gene in the NCBI reference genome of *Mauremys reevesii* (Reeves's turtle).

**Fig. S24.** Gene synteny and architecture (**A**), genomic DNA sequence (**B**), cDNA sequence (**C**), and encoded protein sequence (**D**) of the unannotated *cxcl17* gene in the NCBI reference genome of *Malaclemys terrapin pileata*.

**Fig. S25.** Gene synteny and architecture (**A**), genomic DNA sequence (**B**), cDNA sequence (**C**), and encoded protein sequence (**D**) of the unannotated *cxcl17* gene in the NCBI reference genome of *Alligator mississippiensis* (American alligator).

**Fig. S26.** Gene synteny and architecture (**A**), genomic DNA sequence (**B**), cDNA sequence (**C**), and encoded protein sequence (**D**) of the unannotated *cxcl17* gene in the NCBI reference genome of *Alligator sinensis* (Chinese alligator).

**Fig. S27.** Amino acid sequence alignment of avian and reptilian GPR25 orthologs.

**Table S1.** Information for generating expression constructs for reptilian and avian GPR25s.

| Expression constructs | Vectors for cloning | Restriction enzymes cleaving the vector | Primers for PCR amplification (5' to 3') | Template for PCR amplification | Approach for construct generation |
|---|---|---|---|---|---|
| pcDNA3.1/ Cc-GPR25 | pcDNA3.1(+) | NheI; NotI | Chemical synthesis of Cc-GPR25 coding region (XM_048826666) at CWBio (Taizhou, Jiangsu province, China) | No | Gibson assembly |
| pTRE3G-BI/ Cc-GPR25-LgBiT: SmBiT-ARRB2 | pTRE3G-BI/ MRGPRX2-LgBiT: SmBiT-ARRB2 | NheI; AgeI (removing MRGPRX2) | Forward: CCG TCA GAT CGC CTG GAG AAT TCG GGG AGA CCC AAG CTG GCT AGC<br>Reverse: GCT AGA CCC TCC GCC GGT ACC GAC CGG TGG CAT CTC CGG CCG AAT GGA | pcDNA3.1/ Cc-GPR25 | Gibson assembly |
| PB-TRE/ sLgBiT-Cc-GPR25 | PB-TRE/ sLgBiT-GPR182 | KpnI; PmeI (removing GPR182) | Forward: GGT GGC AGC GGC GGT GGT ACC ACA ACT GAA CCC TGG ACT GT<br>Reverse: GAG GCT GAT CAG CGG GT TTA CAT CTC CGG CCG AAT GGA | pcDNA3.1/ Cc-GPR25 | Gibson assembly |
| PB-TRE/ Cc-GPR25 | PB-TRE/ dCas9-VPR | NheI; PmeI (removing dCas9-VPR) | Forward: TTC CTA CCC TCG TAA AGG TCT AGA G CTC ACT ATA GGG AGA CCC AAG CT<br>Reverse: GTT TCA GTT AGC CTC CCC CGT TT C ACA GTC GAG GCT GAT CAG CGG | pcDNA3.1/ Cc-GPR25 | Gibson assembly |
| pcDNA3.1/ Ap-GPR25 | pcDNA3.1(+) | NheI; NotI | Chemical synthesis of Ap-GPR25 coding region (XM_072028639) at CWBio (Taizhou, Jiangsu province, China) using optimized codons | No | Gibson assembly |
| pTRE3G-BI/ Ap-GPR25-LgBiT: SmBiT-ARRB2 | pTRE3G-BI/ MRGPRX2-LgBiT: SmBiT-ARRB2 | NheI; AgeI (removing MRGPRX2) | Forward: CCG TCA GAT CGC CTG GAG AAT TCG GGG AGA CCC AAG CTG GCT AGC<br>Reverse: CCC TCC GCC GGT ACC GAC CGG TGG TGT TCC AGG TCT AGG GTT | pcDNA3.1/ Ap-GPR25 | Gibson assembly |
| pcDNA3.1/ Gg-GPR25 | pcDNA3.1(+) | NheI; NotI | Chemical synthesis of Gg-GPR25 coding region (XM_015275387) of C-terminal 365 amino acids at CWBio (Taizhou, Jiangsu province, China) using optimized codons | No | Gibson assembly |
| pTRE3G-BI/ Gg-GPR25-LgBiT: SmBiT-ARRB2 | pTRE3G-BI/ MRGPRX2-LgBiT: SmBiT-ARRB2 | NheI; AgeI (removing MRGPRX2) | Forward: CCG TCA GAT CGC CTG GAG AAT TCG GGG AGA CCC AAG CTG GCT AGC<br>Reverse: CCC TCC GCC GGT ACC GAC CGG TGG CAC CTT GGT GCC CCC CGA GGA | pcDNA3.1/ Gg-GPR25 | Gibson assembly |

PB-TRE/sLgBiT-GPR182 was generated in our laboratory based on PB-TRE/dCas9-VPR (Addgene cat#: 63800) via removal of the dCas9-VPR fragment by NheI and PmeI cleavage and then ligation of sLgBiT-GPR182 fragment, unpublished data.

pTRE3G-BI/MRGPRX2-LgBiT:SmBiT-ARRB2 was generated in our previous study.

For oligo primers, the sequence pairing with vector for Gibson assembly is highlighted in yellow, the sequence pairing with template for PCR amplification is underlined.

**Table S2.** Reptilian CXCL17 orthologs identified in present study. Information about these reptilian CXCL17 orthologs was retrieved from NCBI databases. Their signal peptide was predicted via the SignalP6.0 algorithm (https://services.healthtech.dtu.dk/services/SignalP-6.0/). The pI value of mature peptide was predicted via an online server (https://www.chinapeptides.net/tool.aspx).

| Group | Species | Information | Amino acid sequence (signal peptide shaded) | pI |
|---|---|---|---|---|
| Snakes | *Ahaetulla prasina* | Unannotated, adjacent to *LIPE* (Fig. S4)<br>Genomic Sequence: NC_080548.1; Chromosome 10; Reference ASM2864084v1; Position: 29,248,800-29,253,200; Complementary strand | MDRFFIYLMLVLLVGLVGCSERGGSYGEGKERSLS<br>RSKRSKLWHPSCTCPDLNELQHPKYYKHACPCDG<br>FSLHSLSEHRSPRRHHHHHRPNKRPVLQKTKCRRF<br>LRHCAFKGIHVPL | 10.2 |
| | *Crotalus tigris*<br>(Tiger rattlesnake) | Unannotated, adjacent to *LIPE* (Fig. S5)<br>Genomic Sequence: NW_024096990.1; Unplaced Scaffold; Reference ASM1654583v1; Position: 1,413,400-1,420,200; Complementary strand | MDRFFLFLMSALLVGLVCSSEKGGRSDQASLLSRS<br>KRSRLEPRSCPCPDLSELQYPHKDKHPCPCDGFTLS<br>SLSEHSSHHHHPPRKNKQLVLHKKKCRRFLRHCAF<br>KGIHIPL | 9.9 |
| | *Erythrolamprus reginae* (royal ground snake) | Unannotated, adjacent to *LIPE* (Fig. S6)<br>Genomic Sequence: NC_091960.1; Chromosome 11; Reference rEryReg1.hap1; Position: 4,809,500-4,817,500 | MGRFVFFLMLAVLGGLVCCSEGDGSHSEGNEALLS<br>RSKRSKQWPESCPCPDLNALGRLKKHEHACPCDG<br>FRVRSLSGHRPHDHHFHKRYLSRRALCKKFLRHCT<br>FQGIHIPL | 9.6 |
| | *Liasis olivaceus* | Annotated, NCBI BLAST<br>Protein ID: KAM6428849 | MDRFFLFLMSLLLVGVVCCSGKEKDGSYGQGNFQ<br>ERSLSRSKRSGRWHQSCTCPDLGELQNPKQQKYAC<br>PCNGFNLSSLSEHHHHHHPRKTRRLWLHKRDKCR<br>KFLRHCAFKGIHIPL | 10.1 |
| | *Naja naja* | Annotated, NCBI BLAST<br>Protein ID: KAG8143504 | MDRFFFFLVLGVLVGLVCCSGKGGRYGEEKESLLS<br>RSKRSKLWHRSCTCPDLNELQNPKKHNHACPCDG<br>FSLHSLSEHRKHRPHRKSKQHALRKAKCKRFLKH<br>CAFNGMHIPL | 10.2 |
| | *Natrix helvetica* | Annotated, adjacent to *LIPE*<br>Gene symbol: LOC309397096; Gene description: uncharacterized protein; Gene ID: 309397096; mRNA ID:XM_081962819; Protein ID: XP_081818945 | MIRFFIFLMLVVGLVGCSEEGGRDGEGKESLLRRSK<br>RSRLRYPNCPCPDLDELRHPKRRSPDCPCDGFSLRS<br>LSEHRNQRHSRKNKQYASHKKKCRKFLRHCAFQG<br>IHVPL | 10.1 |
| | *Notechis scutatus* (mainland tiger snake) | Unannotated, adjacent to *LIPE* (Fig. S7)<br>Genomic Sequence: NW_020716766.1; Unplaced Scaffold; Reference TS10Xv2-PRI; Position: 1,675,900-1,682,500; Complementary strand | MDRFFFFLVLGVLVSLVCCSEKGTRYGEEKESLLRR<br>SKRSKLWHPSCTCPDLNEFQQLKKHNHACPCNGL<br>SLHSLSEHRNPHPHRKSKKHALHKKKCKMFLRHC<br>AFNGMHIPL | 9.9 |
| | *Ophiophagus hannah* | Annotated, NCBI BLAST<br>Protein ID: CAP9673002 | MDRFFFFLVLGILVGLVCCSEKGGRYGEEKESLLSR<br>SKRSKLWHRSCTCPDLNELQHPKKHNHACPCDGF<br>SLHSLSEHRNLHPHRKSKKHALRKTKCKRFLRHCA<br>FNGMHVPL | 10.0 |
| | *Pantherophis guttatus* | Unannotated, adjacent to *LIPE* (Fig. S8)<br>Genomic Sequence: NW_026844053.1; Unplaced Scaffold; Reference CU_Pguttatus_1; Position: 4,374,600-4,380,200 | MDRFFIFLMLALSVGLLCRSETGGSYGEGQESLLS<br>RSKRSKLWHPSCPCPDLNERQHPKKHEHACPCDGF<br>SLRSLSEHRNHHRHHHRKNKRHVLHKTKCRRFLR<br>HCTFKGIHVPL | 10.1 |
| | *Pituophis catenifer annectens* | Unannotated, adjacent to *LIPE* (Fig. S9)<br>Genomic Sequence: NW_027256889; Unplaced Scaffold; Reference rPitCat1.0.hap2; Position: 4,505,500-4,511,500 | MDRFFIFLMLALSVGLLCRSETGGSYGEGKESLLS<br>RSKRSKLWHPSCPCPDLNERQHPKKHEHACPCDGF<br>SLRSLSEHRNHHHHRHHHRKNKRHVLHKTKCRRF<br>LRHCTFKGIHVPL | 10.1 |
| | *Protobothrops mucrosquamatus* (brown spotted pit viper) | Unannotated, adjacent to *LIPE* (Fig. S10)<br>Genomic Sequence: NW_015387026.1; Unplaced Scaffold; Reference P.Mucros_1.0 Primary Assembly; Position: 292,200-297,200; Complementary strand | MDRFFLFLMSALLVGLVCSSEKGGRYDQASLLSRS<br>KRSRQGPRSCPCPDLSELQYPKKDKHPCPCDGLTL<br>SSLSEHPSHHHHHPPRKNKQLVLRKKKCRRFLRHC<br>AFKGIHIPL | 10.1 |
| | *Pseudonaja textilis* | Unannotated, adjacent to *LIPE* (Fig. S11)<br>Genomic Sequence: NW_020769385.1; Unplaced Scaffold; Reference EBS10Xv2-PRI; Position: 3,279,800-3,285,800 | MDRFSFFLVLGVLVSLVCCSEKGRRYGEEKESLLR<br>RSKRSKLWHPSCTCPDLNEFQHLKKHNHACPCNG<br>LSLHSLSEHHNPHPHRKSKKHALRKKKCKMFLRH<br>CAFNGMHIPL | 10.1 |

| | | | | |
|---|---|---|---|---|
| | *Thamnophis elegans* (Western terrestrial garter snake) | Unannotated, adjacent to *LIPE* (Fig. S12)<br>Genomic Sequence: NC_045552.1; Chromosome 12; Reference rThaEle1.pri; Position: 4,545,200-4,549,200 | MVRFFIFLVLVVLLGLVGCSEEDGRDGEGKESLLR<br>RSKRSRLGHPNCPCPDLDGLRRPKRRSPECPCDGFS<br>LRSLSEHHNQRHSRKSKHHVSHKNKCRRFLKHCA<br>FKGIHVPL | 10.1 |
| | *Vipera berus* (adder) | Annotated, adjacent to *LIPE*<br>Gene symbol: LOC308672102; Gene description: uncharacterized protein; Gene ID: 308672102; mRNA ID: XM_081334721; Protein ID: XP_081190847 | MERFFLFLMSALLIGLVCSSEKGGRYDQESLLSRSK<br>RSKPGHRPCPCPDLSELQRPKKHKHPCPCDGFTLTS<br>LSEHPSHPHPPRKNKQFVLRKKKCTRFLRHCAFKG<br>IHIPL | 10.1 |
| | *Vipera latastei* | Annotated, NCBI BLAST<br>Protein ID: KAM3829549 | MERFFLFLMSALLIGLVCSSEKGGRYDQESLLSRSK<br>RSKPGHRPCPCPDLSELQRPKKHKHPCPCDGFTLTS<br>LSEHPSHPHPPRKNKQFVLRKKKCSRFLRHCAFKG<br>IHIPL | 10.1 |
| Lizards | *Anolis carolinensis* (green anole) | Annotated, adjacent to *LIPE*<br>Gene symbol: LOC134293725; Gene description: synapsin-2-like; Gene ID: 134293725; mRNA ID: XM_062962179; Protein ID: XP_062818249 (isoform 1) | MAKTSALLLALLAAFLGVTFGSVSDGGEGLVLPKA<br>ELSRSKRSPGGTELCPCPGMDAPKRHKHHCPCDHF<br>GFFFMTEEPHRRRRRRHHHHPHHPHHHHGRKAKH<br>HLQQRKNQCLRFLKHCLRRQISIPL | 10.7 |
| | | Annotated, adjacent to *LIPE*<br>Gene symbol: LOC134293725; Gene description: synapsin-2-like; Gene ID: 134293725; mRNA ID: XM_062962178; Protein ID: XP_062818248 (isoform 2) | MAKTSALLLALLAAFLGVTFGSGERPVSDGGEGLV<br>LPKAELSRSKRSPGGTELCPCPGMDAPKRHKHHCP<br>CDHFGFFFMTEEPHRRRRRRHHHHPHHPHHHHGR<br>KAKHHLQQRKNQCLRFLKHCLRRQISIPL | 10.6 |
| | *Eublepharis macularius* | Unannotated, adjacent to *LIPE* (Fig. S13)<br>Genomic Sequence: NC_072804.1; Chromosome 15; Reference MPM_Emac_v1.0 Primary Assembly; Position: 52,267,800-52,274,500; Complementary strand | MGGLLVWTLLLLLLSVSIFGSSTRGKDQGLHVGQT<br>ERELSGSDQSQNGSGFCRCPGSPRPQKQKKHRHVC<br>PCDDVAVSQPGRHRRHPHSKNKHFPRHRKQRCRTS<br>HSKCKHRHLMNGLQIPL | 10.7 |
| | *Hemicordylus capensis* (graceful crag lizard) | Unannotated, adjacent to *LIPE* (Fig. S14)<br>Genomic Sequence: NC_069663.1; Chromosome 7; Reference rHemCap1.1.pri; Position: 24,291,500-24,296,500; Complementary strand | MDRFFALTLLLLLLQSVSVTCSARKKDGGLQVDQ<br>AVAKLSRSRRSYDGNKLCPCPGLHGLHRQNHHQH<br>RCPCDGLKLYLPEHSRHHRYRHGRSSRHLNSRRQE<br>CQKFLKFCQDPMRGLQIPL | 10.4 |
| | *Lacerta agilis* (Sand lizard) | Unannotated, adjacent to *LIPE* (Fig. S15)<br>Genomic Sequence: NC_046319.1; Chromosome 8; Reference rLacAgi1.pri Primary Assembly; Position: 74,732,500-74,741,500; Complementary strand | MGRFLALTLLLLSLSILCCTAKDKGRRFRSGHKEGE<br>PAQGGHRPCSCSDLHGPRNPKKHGRLCPCDGILTG<br>LSLPGQLHHHPRGMAKRHLHQRRHRCQAFVNKC<br>HMMRVDIPL | 10.6 |
| | *Paroedura picta* | Annotated, adjacent to *LIPE*<br>Gene symbol: LOC143837164; Gene description: uncharacterized LOC143837164; Gene ID: 143837164<br>mRNA ID: XM_077336691; Protein ID: XP_077192806 | MERVLLWLLLLLSFSVSCFSAGEKDRGFQVGQSEG<br>NPSRHETIGDGSKLCLCSGSQRLQKQKKHGHVCPC<br>DQFIMPRSGRHHRHTHHKNRELPHHRKHKCRDFQ<br>SNCKHRKLMKGLQIPL | 10.1 |
| | *Podarcis bocagei* | Annotated, adjacent to *LIPE*<br>Gene symbol: LOC308592568; Gene description: uncharacterized protein; Gene ID: 308592568; mRNA ID: XM_081204126; Protein ID: XP_081060252.1 | MGRFLAWTLLLLSLSILCCTAKDKGRRFRSGHKGG<br>EPAQEGHQPCSCSDLRGPRNPKKHGRLCPCDGILT<br>GLSLPEQHHRHHRGLAKRHLHQRRHRCQAFVNKC<br>HMMRVAIPL | 10.9 |
| | *Podarcis cretensis* | Annotated, adjacent to *LIPE*<br>Gene symbol: LOC307221370; Gene description: uncharacterized protein; Gene ID: 307221370; mRNA ID: XM_079985840; Protein ID: XP_079841966 | MGRFLALTLLLLSLSILCCMAKDKGRRFRSGHKGG<br>ERAQEGHQPCSCSDLRGPRNPKKHGQLCPCDGILT<br>GLSMPEQLHRHHRGLAKRHLHQRRHRCQAFVNK<br>CHMMRVAIPL | 10.9 |
| | *Podarcis erhardii* | Annotated, adjacent to *LIPE*<br>Gene symbol: LOC309346154; Gene description: uncharacterized protein; Gene ID: 309346154; mRNA ID: XM_081869884; Protein ID: XP_081726010 | MGRFLALTLLLLSLSILCCMAKNKGRRFRSGHKGG<br>EPAQEGHQPCSCSDLRGPRNPKKHGQLCPCDGILT<br>GLSLPEQLHRHHRGLAKRHLHQRRHRCQAFVNKC<br>HMMRVAIPL | 10.9 |
| | *Podarcis lilfordi* | Annotated, NCBI BLAST<br>Protein ID: CAI5780994 | MCRFLALTLLLLSLSILCYTAKDKGRRFRSGHRGG<br>EPDQEGHQPCSCSDLRGPWNPKKHRRLCPCDGILT<br>GLSLPEQHHRHHRGLAKRHLHQRRHRCQAFVNKC<br>HMMRVAIPL | 10.8 |
| | *Podarcis melisellensis* | Annotated, adjacent to *LIPE* | MGRFLALTLLLLSLSILCCMAKDKGRRFRSGHKRG<br>EPAHEGHQPCSCSDLRGPRNPKKHGRLCPCDGILT | 11.3 |

| | | | | |
|---|---|---|---|---|
| | | Gene symbol: LOC309376218; Gene description: C-X-C motif chemokine 17-like; Gene ID: 309376218; mRNA ID: XM_081922491; Protein ID: XP_081778617 | GLSLPEQHHRHHRGLAKRHLHQRRRRCQAFVNKCHMMRVAIPL | |
| | *Podarcis muralis* (Common wall lizard) | Annotated, adjacent to *LIPE*<br>Gene symbol: LOC144328615; Gene description: uncharacterized; Gene ID: 144328615; mRNA ID: XM_077932619; Protein ID: XP_077788745 | MGRFLALTLLLLSLSILCCTAKDKGRRFRSGHKGGEPDQEGHEPCSCSDLHGPRNPKKHGRLCPCDGILTGLSLPEQLHRHHRGMSKQHLHQRRHRCQAFVNKCHMMRVAIPL | 10.0 |
| | *Podarcis pityusensis* | Annotated, adjacent to *LIPE*<br>Gene symbol: LOC308484494; Gene description: uncharacterized protein; Gene ID: 308484494; mRNA ID: XM_081019507; Protein ID: XP_080875633 | MCRFLALTLLLLSLSILCCTAKDKGRRFRSGHKGGEPDQEGHQPCSCSDLRGPRNPKKHRRLCPCDGILTGLSLPEQHHRHHRGLAKRHLHQRRHRCQAFVNKCHMMRVAIPL | 10.9 |
| | *Podarcis vaucheri* | Annotated, adjacent to *LIPE*<br>Gene symbol: LOC309536991; Gene description: uncharacterized; Gene ID: 309536991; mRNA ID: XM_082231425; Protein ID: XP_082087551 | MGRFLAWTLLLLSLSILCCTAKDKGRRFRSGHKGGEPAQEGHQPCSCSDLRGPRNPKKHGRLCPCDGILTGLSLPEQHHRHHRGLAKRHLHQRRHRCQAFVNKCHMMRVAIPL | 10.9 |
| | *Pogona vitticeps* (central bearded dragon) | Annotated, adjacent to *LIPE*<br>Gene symbol: LOC140701947; Gene description: uncharacterized LOC140701947; Gene ID: 140701947; mRNA ID: XM_072979500; Protein ID: XP_072835601 | MDKLFASMLVLLFFSVLCCSATETGVRHHFGHAEGEQNRPKNRGGEQELCTCPGSHGPQKQTNHKRGCPCNPIDFHLTEHAHHHRHDKTKHHLHHKKQSCLMFLRRCRLKRLHVPL | 9.8 |
| | *Rhineura floridana* | Unannotated, adjacent to *LIPE* (Fig. S16)<br>Genomic Sequence: NC_084494.1; Chromosome 15; Reference rRhiFlo1.hap2; Position: 31,807,800-31,813,500; Complementary strand | MHRFRALTLLLLSLSIIYCLAKEKDGGFHLGRKERELSRHQPGRHGHEPCSCADLRGPQKQKKHGHMCPCDGISNNLSLPEQPHPHRHGKTKRHHKWHQCQMFLRHCYLKRLHIPL | 9.9 |
| | *Sceloporus undulatus* (fence lizard) | Annotated, adjacent to *LIPE*<br>Gene symbol: LOC121915120; Gene description: C-X-C motif chemokine 17-like; Gene ID: 121915120<br>mRNA ID (isoform 1): XM_042438999; XM_042438997; XM_042438998; XM_042438992; XM_042439000; XM_042438995; XM_042438993; XM_042438994; XM_042438991;<br>Protein ID: XP_042294933; XP_042294931; XP_042294932; XP_042294926; XP_042294934; XP_042294929; XP_042294927; XP_042294928; XP_042294925<br>Isoform 1 | MAKVFLALGLVLFLASLGAVFCPGEETDVGQEIGRPGGRLSRSKRSPDGHKPCPCPGTHGPQKQERHRWACPCDDIPLSVPEQPRRRHHHHHHGGKTKRHLHHKKKQCQRFLRHCHLRQVNIPL | 10.3 |
| | | Gene symbol: LOC121915120<br>Gene description: C-X-C motif chemokine 17-like; Gene ID: 121915120; mRNA ID (isoform 1): XM_042439001; Protein ID: XP_042294935<br>Isoform 2 | MAKVFLALGLVLFLASLGAVFCPDVGQEIGRPGGRLSRSKRSPDGHKPCPCPGTHGPQKQERHRWACPCDDIPLSVPEQPRRRHHHHHHGGKTKRHLHHKKKQCQRFLRHCHLRQVNIPL | 10.7 |
| | *Sphaerodactylus townsendi* | Unannotated, adjacent to *LIPE* (Fig. S17)<br>Genomic Sequence: NC_059430.1; Chromosome LG06; Reference MPM_Stown_v2.3; Position: 125,500,500-125,503,700; Complementary strand | MGRFLALVILLLLLSVSTFCFSAGEKDHGFHIRHVERKLSRSELGHGRSGLCPCPGYHRSHSHGKNKHFPHRMKQKCKSYQRHCQHRTLMGGLQIPL | 10.3 |
| | *Zootoca vivipara* (common lizard) | Unannotated, adjacent to *LIPE* (Fig. S18)<br>Genomic Sequence: NC_083281.1; Chromosome 6; Reference rZooViv1.1; Position: 21,533,600-21,541,200 | MGRFLALTFLLLSLSILCCTAKDKGRRFRSGHKERQPARGGHQPCSCSDLREPWNPKKHGRPCPCDGILTGLSLPEQLHHHHRGMAKRHLHQRRHRCQAFVNKCHMMRVDIPL | 10.6 |
| Turtles | *Caretta caretta* (loggerhead turtle) | Annotated in INSDC annotation, adjacent to *LIPE*<br>Gene ID: ACLRTB_LOCUS24568; Protein ID: CAM4711560 | MDRILCLLLLLLYVTITACSPHVPGGGQGAEGELNPTRRRATGAGVCECPGWSGSRRQLAGDLRVGARHNGLCPCDSIVPHGQGRGRHPQGRGRRGQHHGRKQQRCHGIPSRCRQLGLAKGLLLPL | 11.5 |
| | | Unannotated in NCBI RefSeq, adjacent to *LIPE*<br>Genomic Sequence: NC_064494.1; Chromosome 22; GSC_CCare_1.0; Position: 424,500-427,100 | MDRILCLLLLLLYVTITACSPHVPGGGQGAEGELNPTRRRATGAGLCECPGWSGSRRQLAGDLRVGARHNGLCPCDGIVPHGQGRGRHPQGRGRRGQHHGRKQQRCHGIPSRCRQLGLAKGLLLPL | 11.5 |

| | | | | |
|---|---|---|---|---|
| | *Chrysemys picta bellii* (western painted turtle) | Unannotated, adjacent to *LIPE* (Fig. S19)<br>Genomic Sequence: NC_088810.1; Chromosome 20; Reference ASM1138683v2; Position:732,100-734,000; Complementary strand | MDQLLCLFVLLSVTITTCSPHVPGGGHGADGELNP<br>VRRRATGAGLCECPGWSGSKRQLGDLRDGARHN<br>GLCPCDGIVPHGQGRGRHPQGRGRRTQHHRRRRQ<br>RCREIPSRCREKGLAKGLLLPL | 11.4 |
| | *Eretmochelys imbricata* | Annotated, NCBI BLAST<br>Protein ID: CAM5176759 | MDRILCLLLLLLSVTITACSPHVPGGGQGAEGELNP<br>TRQRATGAGLCDCPGWSGSQRQLAGDLGVGARH<br>NGLCPCDGIVPHGQGRGRHPQGRGRRGQHHGRRQ<br>QRCRGIPSRCRQPGLAKGLLLPL | 11.6 |
| | | Annotated, NCBI BLAST<br>Protein ID: CAM5176396 | MDRILCLLPLLLSVTITACSPHVPGGGQGAEGELNP<br>TRQRATGAGLCDCPGWSGSRRQLAGDLGVGARH<br>NGLCPCDGIVPHGQGRGRHPQGRGRRGQHHGRRQ<br>QRCRGIPSRCRQPGLAKGLLLPL | 12.2 |
| | *Gopherus evgoodei* (Goodes thornscrub tortoise) | Unannotated, adjacent to *LIPE* (Fig. S20)<br>Genomic Sequence: NW_022060097.1; Unplaced Scaffold; Reference rGopEvg1_v1.p Primary Assembly; Position: 78,300-80,100; Complementary strand | MDQLLSLFLLSVGMAACSAHVPGGGRRAEGELNP<br>VRQRATGAGLCECPGWSGSQRQLAGDLGFGARHN<br>GVCPCDSIIPHGQGRRHHPQSRGRRSEHHRRRRQR<br>CRKTPRRCRRQGLTKGLLLPL | 11.8 |
| | *Gopherus flavomarginatus* (Mexican gopher tortoise) | Unannotated, adjacent to *LIPE* (Fig. S21)<br>Genomic Sequence: NC_066636.1; Chromosome 20; Reference rGopFla2.mat.asm; Position: 435,700-437,300 | MDRLLSLFLLLSVGMAACSAHMPGGGRRAEGELN<br>PVRQRATGAGLCECPGWSGSRRQLAGDLGVGARH<br>NGVCPCDGIIPHGQGRGRHPQSRGRRSEHHRRRRQ<br>RCRETPRRCRRQGLTKGLLLPL | 11.8 |
| | *Lepidochelys olivacea* | Annotated, NCBI BLAST<br>Protein ID: CAM4679888 | MDRILCLLLLLLSVTITACSPHVPGGGQGAEGELNP<br>TRRRATGAGLCECPGWSGSRRQLAGDLRVGARHN<br>GLCPCDGIVPHGQGRGRHPQGQGRRGQHHGRKQ<br>QRCRGIPSRCRQLGLAKGLLLPL | 11.4 |
| | | Annotated, NCBI BLAST<br>Protein ID: CAM4697903 | MDRILCLLLLLLSVTITACSPHVPGGGQGAEEELNP<br>TRRRATGAGLCECPGWSGSRRQLAGDLRVGARHN<br>GLCPCDGIVPHGQGRGRHPQGWGRRGQHHGRKQ<br>QRCRGIPSQCRQLGLAKGLLLPL | 11.7 |
| | *Lepidochelys kempii* | Annotated, NCBI BLAST<br>Protein ID: CAM4625820 | MDRILCLLLLLLSVTITACSPHVPGGGQGAEGELNP<br>TRRSATGAGLCECPGWSGSRRQLAGDLRVGARHN<br>GLCPCDGIVPHGQGRGRHPQGQGRRGQHHGRKQ<br>QRCRGIPSRCRQLGLAKGLLLPL | 11.2 |
| | *Mauremys mutica* (yellowpond turtle) | Unannotated, adjacent to *LIPE* (Fig. S22)<br>Genomic Sequence: NW_025423358.1; Unplaced Scaffold; Reference ASM2049712v1 Primary Assembly; Position: 140,500-138,500; Complementary strand | MDRLLCLFLLLSVGMAACSAPVPGGGHGAEGELN<br>PVRQHTTGAGLCDCPGWSGSQRQLAGDLGVGAR<br>HNGLCPCDSIVPYGQGRGHHPQGRGRRSEHHRRR<br>RQRCREIPSRCRRQGLAKGLLLPL | 10.9 |
| | *Mauremys reevesii* (Reeves's turtle) | Unannotated, adjacent to *LIPE* (Fig. S23)<br>Genomic Sequence: NC_052646.1; Chromosome 24; Reference ASM1616193v1 Primary Assembly; Position: 21,690,400-21,691,950; Complementary strand | MDRLLCLFLLLSVGMAACSAPVPGGGHRAEGELN<br>PDRRHATGAGLCECPGWSGSQRQLAGDLGVGARH<br>NGLCPCDGIVPHGQGRGHHPQSRGRRSEHHRRRR<br>QRCREIPSRCRRQGLAKGLLLPL | 11.3 |
| | *Malaclemys terrapin pileata* | Unannotated, adjacent to *LIPE* (Fig. S24)<br>Genomic Sequence: NC_071525.1; Chromosome 21;<br>Reference rMalTer1.hap1; Position: 21,082,400-21,085,100; Complementary strand | MDQLLCLFLLLSVTITTCSPRVPGGGHGADGELNP<br>VRRRVTGAGLCECPGWSGSKRQLGDLGDGARHN<br>GLCPCDGIVPHGQGRGRHPRGRGRRTQHHRRRRQ<br>RCREIPSRCRQTGLAKGLLLPL | 11.6 |
| | *Natator depressus* (flatback sea turtle) | Unannotated in NCBI RefSeq, adjacent to *LIPE*<br>Annotated in INSDC annotation, adjacent to *LIPE*<br>Gene symbol: ACLRS4_LOCUS15510; Protein ID: CAM5135525 | MDRILCLLPLLLSVTITACSPHVPGGGQGAERELNP<br>TRRRATGAGLCECPGWSGSRRQLAGELGVGARHN<br>GLCPCDGIVPRGQGTERHPQGRGRRGQHHSRKQQ<br>RCRGIPSRCRQLGLAKGLLLPL | 11.1 |
| Alligators | *Alligator mississippiensis* (American alligator) | Annotated, NCBI BLAST<br>Protein ID: KYO44589<br>(lacking signal peptide) | MPARCLDLPRRPAQKENGDSETSLDLCSCEGWLG<br>AWGQLEGGVWGGQGHNSACPCRGIHPHEVMTAP<br>HRPTQGRAQGRPKRHQRVWRCRQFLRRCQQRSL<br>MQGLLLPL | 10.4 |
| | | Unannotated, adjacent to *ZNF225 (LOC106737775)* (Fig. S25) | MDLRLSFVLLLLLLGLGSAVKKPGRRPAQKENGDS<br>ETSLDLCSCEGWLGAWGQLEGGVWGGQGHNSAC<br>PCRGIHPHEVMTAPHRPTQGRAQGRPKRHQRVWR | 10.9 |

| | | | | |
|---|---|---|---|---|
| | | Genomic Sequence: NC_081838.1; Chromosome 15; Reference rAllMis1 Primary Assembly; Position: 28,186,800-28,188,800 | CRQFLRRCQQRSLMQGLLLPL | |
| | *Alligator sinensis* (Chinese alligator) | Unannotated, adjacent to *LOC102385260* (Fig. S26) Genomic Sequence: NW_005843744; Unplaced Scaffold Reference ASM45574v1 Primary Assembly; Position: 34,140-35,400; Complementary strand | MDPRLGFVLLLLGLGSAVKNPGRRPAQKENGGSE TSLDLCPCEGWLGAWGQLEGGVWGGQGHNSACP CRGIHPHEVMTAPHRPTQGRAQGRPKRHQRVWRC RQFLRRCQQRSLMQGLLLPL | 11.1 |

**Table S3.** Avian and reptilian and GPR25 orthologs. Information about these orthologs was downloaded from the NCBI database.

| Group | Species | NCBI information | Amino acid sequence |
|---|---|---|---|
| Birds | *Anas platyrhynchos* (mallard) | Gene ID: 139999625<br>mRNA ID: XM_072028639<br>Protein ID: XP_071884740 | MAPEATEEPEASADYEYPAGGTAPGSPEQLFTTVVIPVLYSFIFLLGFAGNLFVIVLLAKRSGGRRAVDTFVLNLAVADVVFVCTLPFWVAAGARGNRWLLGEGLCKASSYAISVNRCSSILFLTALSVERYLVIWKVLDTRRGGSRRHVPLACGLIWAVSLLLGAPALWYRRLDGDDCWDEDGEDFSLAMVFLTFLLPLGVISFCYCSIYCRLQRHVRLGRGVRRSHRVIVAIVAAFICSWLPLNACKVLLFLLAKGTLALPAGQEGTLRWVVATSTCLAFANSCVNPVIYALMDPRLRCPPALGARTPPSTTASSPSPGSGTKLPAVPNPRPGT |
| | *Anser anser* (Domestic goose) | Gene ID: 310887258<br>mRNA ID: XM_083813007<br>Protein ID: XP_083669133 | MAPEATEEPEASADYEYLAGSNATGSPEQLFTTVFIPVLYSFIFLLGFAGNLFVIALLAKRSGGRRAVDTFVLNLAVADVVFVCTLPLWVAAGARGNRWLLGEGLCKASSYAISVNRCSSILFLTALSVERYLVIWKVLDTRRAGSRRHVLLACGLIWAVSLLLGAPALWYRRLDGDDCWDEDGEDFSLAMVFLTFLLPLGVISFCYCSIYCRLQRHVRLGRGVRRSHRAIVAIIAAFVCSWLPLNACKVLLFLLAKGTLALPAGQEGTLRWVVAASTCLAFANSCVNPLIYALMDPRLRCPPPLGARAPPSTTASSPSPGSGTKPPPAPSPRPGT |
| | *Anser brachyrhynchus* (Pink-footed goose) | Gene ID: 308183337<br>mRNA ID: XM_080569246<br>Protein ID: XP_080425372 | MAPEATEEPEASADYEYLAGSNATGSPEQLFTTVFIPVLYSFIFLLGFAGNLFVIALLAKRSGGRRAVDTFVLNLAVADVVFVCTLPLWVAAGARGNRWLLGEGLCKASSYAISVNRCSSILFLTALSVERYLVIWKVLDTRRAGSRRHVLLACGLIWAVSLLLGAPALWYRRLDGDDCWDEDGEDFSLAMVFLTFLLPLGVISFCYCSIYCRLQRHVRLGRGVRRSHRAIVAIVAAFVCSWLPLNACKVLLFLLAKGTLALPAGQEGTLRWVVAASTCLAFANSCVNPLIYALMDPRLRCPPPLGARAPPSTTASSPSPGSGTKPPPAPSSRPGT |
| | *Apteryx mantelli* (North Island brown kiwi) | Gene ID: 106491482<br>mRNA ID: XM_013951099<br>Protein ID: XP_013806553 | MGPEAAEDLTVSADYYYPEGDNSTGTEEHSSRWESSYMTIFIPILYSFIFLLGFVGNLFVIVLMAKKRGNKRMVDTFVLNLAVADMIFVCTLPLWVVAGACGNRWQFGEGLCKVSSYAINVNRCSSILFLTALSVERYLVIRKVLDTKMTGSKRHIFVTCGLIWVVSLFLGAPSLEYRQLDGDDCWDKDGEDFSLAMVFLTFILPLVVILFCYCSIYYQLHRHIRLGKGMRNSHRIIFTIIGAFICSWLPLNVCKVLLFFIAKEMLVLSEVEEEALRWVISSSTCLAFMNSCINPIIYTLMDRHFRRQLLRPGLRVLCLSPAAKGRNVATTSSVTESSLIFGVRTKPPTSPGHRPAA |
| | *Apteryx rowi* (Okarito brown kiwi) | Gene ID: 112961546<br>mRNA ID: XM_026056851<br>Protein ID: XP_025912636 | MGPEAAEDLTVSADYYYPEGDNSTGTEEHSSRWESSYMTIFIPILYSFIFLLGFVGNLFVIVLMAKKRGNKRMVDTFVLNLAVADMIFVCTLPLWVVAGACGNRWQFGEGLCKVSSYAINVNRCSSILFLTALSVERYLVVRKVLDTKMTGSKRHIFVTCGLIWVVSLFLGAPSLEYRQLDGDDCWDKDGEDFSLAMVFLTFILPLVVILFCYCSIYYQLHRHIRLGKGMRNSHRIIFTIIGAFICSWLPLNVCKVLLFFIAKEMLVLSEVEEEALRWVISSSTCLAFMNSCINPIIYTLMDRHFRRQLLRPGLRVLCLSPAAKGRNVATTSSVTESSLIFGVRTKPPTSLGHQPAA |
| | *Aquila chrysaetos* chrysaetos | Gene ID: 115335183<br>mRNA ID: XM_030000529<br>Protein ID: XP_029856389 | MAPEELAGSADYDYPPVTNVTGNEEVFSRWEVFFTTVFVPILYSFIFLLGLVGNLFVIVLMAKKSGGKRMVDTFVLNLAVADVIFICTLPFWVVAGARGNRWLLGEGLCKVSSYAIAVNRCSSILFLTALSVERYLVIRKVLDTKMMGSQRHVHVTCGIIWTVSLLLGAPSLVYRQLDGDDCWDEDGEDFSLAMVFLTFLLPLGVISFCYCSIYCQLQRHVRLGRGVRRSHRAIVTIVAAFLCSWLPLNTCKVLLFFLAKGTLVLSQGQEVALRWVVAGSTCLAFVNSCINPLVYALMDRRCRPRCPFGPRAPGTGPGAATPSSATDSSLLFGGRIRTKTPTGPRRRSGTELQPALGVPRGHPRVSPPPSSPASPGATAVPAVPPGPQL |
| | *Astur gentilis* (Northern goshawk) | Gene ID: 126052363<br>mRNA ID: XM_049832916<br>Protein ID: XP_049688873 | MAPEELAGSADYDYPPVTNVTGNQEVFSRWEVFFTTVFIPILYSFIFLLGLVGNLFVIVLMAKKSGGKRMVDTFVLNLAVADVIFICTLPFWVVAGARGNRWLLGEGLCKVSSYAIAVNRCSSILFLTALSVERYLVIRKVLDTKMMGSQRHVHVTCGIIWTVSLLLGAPSLVYRQLDGDDCWDEDGEDFSLAMVFLTFLLPLGVISFCYCSIYCQLQRHVRLGRGVRRSHRAIVTIVAAFLCSWLPLNACKVLLFFLAKGTLVLSQGQEVALRWVVAGSTCLAFVNSCVNPLVYALMDRRCRPQCLFSPRALGTGPGAATPSSATDSSLLFGGWIRTKTPAGPRHRSRTELQLALGVLRDHPRVSPPPSSPASLGATAVPPVPPGPQQ |
| | *Athene cunicularia* (burrowing owl) | Gene ID: 113490765<br>mRNA ID: XM_026867336<br>Protein ID: XP_026723137 | MAPEELAALADYDYSPVSNGTGNWEVVFTTVFIPILYSFIFLLGLVGNLFVIVLMGKKSRGKRMVDTFVLNLAVADVIFVCTLPFWVVAGARGNRWLLGEGLCKVSSYAIAVNRCSSILFLTALSVERYLVIRKVLDTKMVGSQRHIHVTCGIIWAVSLLLGAPSLVYRRLDGDDCWDGDGEDFSLAMVFLTFLLPLGVISFCYCSIYCRLQRHVRLGRGVRRSHRAIVTIVAAFLCSWLPLNACKVLLFFLAKGTLVLSQGQEVALRWVLAGSTCLAFVNSCVNPLVYALMDGRCRPRCPFGPCTPGTGPAVAAPSSTTDSSLLFGAWIWTKT |
| | *Athene noctua* | Gene ID: 141969597<br>mRNA ID: XM_074925543<br>Protein ID: XP_074781644 | MAPEELAALADYDYSPVTNGTGNWEVVFTTVFIPILYSFIFLLGLVGNLFVIVLMGKKSRGKRMVDTFVLNLAVADVIFVCTLPFWVVAGARGNRWLLGEGLCKVSSYAIAVNRCSSILFLTALSVERYLVIRKVLDTKMVGSQRHIRVTCGIIWTVSLLLGAPSLVYRRLDGDDCWDGDGEDFSLAMVFLTFLLPLGVISFCYCSIYCRLQRHVRLGRGVRRSHRAIVTIVAAFICSWLPLNACKVLLFFLAKGTLVLSQGQEVALRWVLAGSTCLAFINSCVNPLVYALMDGRCRPRCPFGPCTPGTGPAVAAPSSTTDSSLLFGAWIWTKTPPGPRRRSRTELRPAPGVSPPGCSAVSLGATAVPTVPPGPQP |
| | *Aythya ferina* (common pochard)s | Gene ID: 311423501<br>mRNA ID: XM_084497549<br>Protein ID: XP_084353675 | MGPGPALPGLCPSLWAWSRPAPSSRGSLCSGCFSRCFSVSKINCGSPRACRHPSTPRTLPRTKGSAEACFEQSSDQQPDCRAPPAPPRHRGGTRGSATPMAPEATEEPEASADYEYLAGSNATGSPEQLFTTVVIPVLYSFIFLLGFAGNLFVIVLLAKRSGGRRAVDTFVLNLAVADVVFVCTLPFWVAAGARGNRWLLGEGLCKASSYAISVNRCSSILFLTALSVERYLVIWKVLDTRRAGSRRHVLFACGLIWAVSLLLGAPALWYRRLDGDDCWDEDGEDFSLAMVFLTFLLPLGVILFCYCSIYCRLQRHVRLGRGVRRSHRVIVAIIAAFVCSWLPLNACKVLLFLLAKGTLVLPAGQEGTLRWVVATSTCLAFANSCVNPVIYALMDPRLRCPPALGARTPPSTTASSPSPGSGTKLPPVPSPRPGT |
| | *Aythya fuligula* (tufted duck) | Gene ID: 116498693<br>mRNA ID: XM_032203033<br>Protein ID: XP_032058924 | MAPEATEEPEASADYEYLAGSNATGSPEQLFTTVAIPVLYSFIFLLGFAGNLFVIVLLAKRSGGRRAVDTFVLNLAVADVVFVCTLPFWVAAGARGNRWLLGEGLCKASSYAISVNRCSSILFLTALSVERYLVIWKVLDTRRAGSRRHVLFACGLIWAVSLLLGAPALWYRRLDGDDCWDEDGEDFSLAMVFLTFLLPLGVILFCYCSIYCRLQRHVRLGRGVRRSHRVIVAIVAAFVCSWLPLNACKVLLFLLAKGMLVLPAGQEGTLRWVVATSTCLAFANSCVNPVIYALMDPRLRCPPALGARTPPSTTASSPSPGSGTKLPPVPSPRPGT |
| | *Aythya marila* (greater scaup) | Gene ID: 309887009<br>mRNA ID: XM_082778387<br>Protein ID: XP_082634513 | MAPEATEEPEASADYEYLAGSNATGSPEQLFTTVAIPVLYSFIFLLGFAGNLFVIVLLAKRSGGRRAVDTFVLNLAVADVVFVCTLPFWVAAGARGNRWLLGEGLCKASSYAISVNRCSSILFLTALSVERYLVIWKVLDTRRAGSRRHVLFACGLIWAVSLLLGAPALWYRRLDGDDCWDEDGEDFSLAMVFLTFLLPLGVILFCYCSIYCRLQRHVRLGRGVRRSHRVIVAIVAAFVCSWLPLNTCKVLLFLLAKGTLVLPAGQEGTLRWVVATSTCLAFANSCVNPVIYALMDPRLRCPPALGARTPPSTTASSPSPGSGTKLPPVPSPRPGT |
| | *Balearica regulorum gibbericeps* (East African grey crowned-crane) | Gene ID: 104640805<br>mRNA ID: XM_075775974<br>Protein ID: XP_075632089 | MAPEELASSADYDYPPVTNVTGNWEVSGWEVFFTTVFIPILYSFIFLLGLLGNLFVIVLMAKKSRGKRMVDTFVLNLAVADVIFICTLPFWVVAGARGNRWPLGEGLCKVSSYAIXVNRCSSILFLTALSVERYLVIRKVLDTKMMGSRRHVRVTCGIIWAASLILGAPSLVYRRLDGDDCWDEDGEDFSLTMVFLTFLLPLGVISFCYCSIYCQLQRHVRLGRGVRRSHRAIVTIVVAFLCSWLPLNACKVLLFFLAKGTLVLSQGQEMALRWVVAGSTCLAFVNSCVNPLIYALMDGHCRPRCPFGPCTPGTSPGAAATSSATDSSLLFGGRIRTKTPPGPRHRSGTELQPAPGIPPGHL |

| | | | |
|---|---|---|---|
| | *Branta canadensis* (Canada goose) | Gene ID: 309429597<br>mRNA ID: XM_082035091<br>Protein ID: XP_081891217 | MAPEATEEPEASADYEYLAGSNATGSPEQLFTTVFIPVLYSFIFLLGFAGNLFVIALLAKRSGGRRAVDTFVLNLAVADV<br>VFVCTLPLWVAAGARGNRWLLGEGLCKASSYAISVNRCSSILFLTALSVERYLVIWKVLDTRRAGSRRHVLLACGLIWAV<br>SLLLGAPALWYRRLDGDDCWDEDGEDFSLAMVFLTFLLPLGVISFCYCSIYCRLQRHVRLGRGVRRSHRVIIAIVAAFVC<br>SWLPLNACKVLLFLLAKGTLALPAGQEGTLRWVVAASTCLAFANSCVNPLIYALMDPRLRCPPPLGARAPPSTTASSPSP<br>GSGTKPPPAPSPRPGT |
| | *Caloenas nicobarica* (Nicobar pigeon) | Gene ID: 135996933<br>mRNA ID: XM_065649250<br>Protein ID: XP_065505322 | MAPEELVASADYEYPPVTSVTENQEVSSRWELFFTTIFIPILYSIIFLLGLVGNLFVIVLMAKKSGGKRMVDTFVLNLAV<br>ADVVFVCTLPFWVAAGVRGNRWLLGEGLCKLSSYVIAVNRCSSVLFLTALSVERYLVIRKVLDTKVMGSQRHVRVTCGVI<br>WAVSLLLGAPTLVYRRLDGDDCWDEDGEDFSLAMVFLTFLLPLGVISFCYCSIYCRLQRHVRLGRGVRRSHRAIVTIVTA<br>FLCSWLPLNACKVLLFFLAKGTLVLSQGQEVALRWLVATSTCLAFVNSCVNPLVYALMDGRCRPRCPLGPRVPGTGPRAA<br>TPSSASDSSLLFGVRTRSRHRTRHQPAPGGSPLSPPPSSPASPGATAVPPGPQL |
| | *Calonectris borealis* (Cory's shearwater) | Gene ID: 142093095<br>mRNA ID: XM_075173966<br>Protein ID: XP_075030067 | MAPEELAGSADYDYPPVTTVMENQEVSSSWEVFFTSVFIPILYSFIFLLGLMGNLFVIVLMAKKSGGKRMVDTFVLNLAV<br>ADVIFVCTLPFWVVAGARGNRWLLGEGLCKVSSYAIAVNRCSSILFLTALSVERYLVIRKVLDTKMMGSQRHVHVTCGII<br>WAASLLLGAPSLAYRRLDGDDCWDEDGEDFSLAMVFLTFLLPLGVISFCYCSIYCRLQRHVRLGRGVRRSHRAIVTIVAA<br>FLCSWLPLNACKVLLFFLAKGTLVLSQGQEVALRWVVAGSTCLAFVNSCVNPLVYALMDGRCRPRCPFGPRTPGTGPGAA<br>APSSATDSSLLFGGRIRTKNPPGPRRRSGTELQPAPGVPRGHPGVSPLPSSPASPGATAVPAVPPCPQP |
| | *Caprimulgus europaeus* (Eurasian nightjar) | Gene ID: 306729370<br>mRNA ID: XM_079208642<br>Protein ID: XP_079064768 | MTPEELTTLADYDYPLVTNVTGNQEVSSRWEVFFTTIFIPIFYSFIFLLGLVGNLFVIVLMAKKSGGKRMVDTFVLNLAV<br>ADVIFVCTLPLWVVAGARGNRWLLGEGLCKVSSYVIAVNRCSSILFLTALSVERYLVIRKVLDTKVMGSRRHIRVTCGII<br>WAVSLLLGAPSLLYRRLDGDDCWDEDGEDFSLAMVFLTFLLPLGVISFCYCSIYCRLQRHVRLGRGVRRSHRAIVTIVTA<br>FLCSWLPLNTCKVLLFFLAKGTLVLSQGQEVALRWVVAASTCLAFVNSCVNPLVYALMDGRCRPRCPFGPGALGTGPGVV<br>TSSSTTDSSLFFGGLIRTRTPHGPRRRNGTQLHPVPGTPRGHPGCHAVPCRALITTITIVVAIVAWRRAGAAARGAAWAR<br>ALPVPAAREGGCSPAPPLRLKSKPVPVARAEQSRQRRRAPR |
| | *Chaetura pelagica* (chimney swift) | Gene ID: 104391103<br>mRNA ID: XM_010000329<br>Protein ID: XP_009998631 | MAPEELPGSADYDYPPVTNVTGNQELPSSWEVFFTTVFIPTLYSFIFLLGLVGNLFVIALMAQRSGGKRVVDTFVLNLAV<br>ADVIFVCTLPLWVLAGARGHRWVLGEGLCKLSSYIIAINRCSSILFLTALSVERYLVIRKVLDTKVTGPRWHVRVTCGVI<br>WVVSLLLGAPSLVYRQLDGEDCWDEDGEDFSLAMVFLTFLLPLGIISFCYCSIYCRLQRHVRLGRGVRRSHRAIVTIVTA<br>FLCSWLPLNACKVLLFFLAKGTLVLSQGQEVALSPTVFLYEQDLAPLRYQRLVASHSRIVRTPSLKDYPPGGRGLSKATV<br>TEELKSWHQRARLRGTRPHSLDRQGAFRGPRGGTTRDLPIDRGVLPWAQ |
| | *Charadrius vociferus* (killdeer) | Gene ID: 104284692<br>mRNA ID: XM_009883052<br>Protein ID: XP_009881354 | MATDSHEGLIQPRKVPNPMRESRTHREMHRELLFSHSRGILPRHKAELPRVLESRRLRQLKEELRGPPSDLEQQLRKRHQ<br>RLEEVCAVPLSPVPLAPLERKCEMLFLSRTQINNPTAGCSQHHPGHRGGHPATAMAPEELSGSADYDYPPVTNVTGNREV<br>FSRWEVFFTTVFIPVFYSFIFLLGLVGNLFVIVLMAKKRRGKRMVDTFVLNLAVADVIFVCTLPFWVVAGARGNRWPLGE<br>GLCKVSSYAIAVNRCSSILFLTALSVERYLVIRKVLDTKTTGSQQHVRVTCGVIWAASLLLGAPSLVYRRLDGDDCWDED<br>GEDFSLAMVFLTFLLPLGVISFCYCSIYCRLQRHVRLGRGIRRSHRAIVTIVAAFLCSWLPLNACKVLLFFLAKGTLVLS<br>QGQEVALRWLVAGSTCLAFVNSCVNPLVYALMDGRCRPRCPLGPRAPGDVAVTSSTTDSSVLFGGRMRTSSPP |
| | *Ciconia boyciana* (Oriental stork) | Gene ID: 140643378<br>mRNA ID: XM_072845044<br>Protein ID: XP_072701145 | MAPEEFASSADYDYPPVTNVTGNREVSSGWEVFFTSVFIPILYSFIFLLGLVGNLFVIVLMAKRSGGKRMVDTFVLNLAV<br>ADVIFVCTLPFWVAAGARGNRWLLGEGLCKVSSYAIAVNRCSSILFLTALSVERYLVIRKVLDTKMMGSQRHVRITCGII<br>WAASLLLGAPSLVYRRLDGDDCWDEDGEDFSLAMVFLTFLLPLVVISFCYCSIYCRLQRHVRLGRGVRRSHRAIVTIVAA<br>FLCSWLPLNACKVLLFFLAKGTLVLSQGQEGALRWVVAGSTCLAFVNSCVNPLVYALMDGRCRPRCPFGPCAPGTGPGTA<br>APSSATDSSLLFGGWIWTKTPPGPRRRSGTELQPAPGVPRGHPVVSSPPSSPASPGATAVPAVPPGPQP |
| | *Clangula hyemalis* (long-tailed duck) | Gene ID: 310979603<br>mRNA ID: XM_083962731<br>Protein ID: XP_083818857 | MAPEATEEPEASADYEYLAGSNATGSAEQLFTTVVIPVLYSFIFLLGFAGNLFVIMLLAKRSDGRRAVDTFVLNLAVADV<br>VFVCTLPFWVAAGARGNRWLLGEGLCKASSYAISVNRCSSILFLTALSVERYLVIWKVLDTRRAGSRRHVLLACGLIWAV<br>SLLLGAPALWYRRLDGDDCWDEDGEDFSLAMVFLTFLLPLGVILFCYCSIYCRLQRHVRLGRGVRRSHRVIVAIVAAFVC<br>SWLPLNACKVLLFLLAKGTLALPAGQEGTLRWVVATSTCLAFANSCVNPVIYALMDPRLRCPPALGARTPPSTTASSPSP<br>GSGTKLPPVPSPRPGT |
| | *Colius striatus* (speckled mousebird) | Gene ID: 133627592<br>mRNA ID: XM_062013886<br>Protein ID: XP_061869870 | MAPEELGASADYDYAALPNGTDGPRLISGWELTFTTVFIPTFYSFIFLLGLSGNIFVIVLLARSSGGRRLVDTFVLNLAV<br>ADVVFVLLLPLWVVAGARGQRWPLGEGLCKLSSYVVAVNRCSSIFFLTALSVERYLVMRKALGTKTAPWRRHIGLTCGTI<br>WAASLLLAAPALLYRRLDGDDCWDEDGEDFSLAMVFLTFLLPLGVISFCYCSVSCRLQRHVRLGRGVRRSHRAVVAIVAA<br>FLCSWLPLNACKVLLFFLAKGTLVLSHGQESALRWLVALSTCLAFVNSCVNPVLFALTAGRCRARCPLAPRAPAAGPAAA<br>SSPLTDSSLLLGARIRTSTLPGPRRGFGTELQPAAGVPRAPPGVSPSRPPLSAAASPQ |
| | *Columba livia* (rock pigeon) | Gene ID: 102094681<br>mRNA ID: XM_065038930<br>Protein ID: XP_064895002 | MAPEELVTSDDYEYPPVTSAMENQWVSSRGELFFTTIFIPILYSLIFLLGLVGNLFVIVLMAKKSRGKRMVDTFVLNLAV<br>ADVVFVCTLPFWVAAGAQGNRWLLGEGLCKLSSYVIAVNRCSSILFLTALSVERYLVIRKVLDTKVMGSQRHVRVTCGVI<br>WAVSLLLGAPTLLYRRLDGHDCWDEDGEDFSLAMVFLTFLLPLGVISFCYCSIYCRLQRHVRLGRGVRRSHRAIVTIVTA<br>FLCSWLPLNACKVLLFFLAKGMLVLSQGQEVALRWLVATSTCLAFVNSCVNPLIYALMDGHCRPRCPLGPRAPGTGPRAA<br>APSSTSDSSLLFGVWTRSRCRTRHQPAPGGTPLSPPASSPGATAVPPGPQL |
| | *Cuculus canorus* (common cuckoo) | Gene ID: 104062316<br>mRNA ID: XM_054087959<br>Protein ID: XP_053943934 | MLFFSRSHINNTTAGRGQHHPGHQGGHPATAMAPEELSGSADYDYPLVTNMTVNQEVFSRWEVIFTTIFIPILYSFIFLL<br>GLMGNLFVIVLMAKRRGNKRMVDTFVLNLAVADVIFICTLPFWVAAGARGNRWPLGEGLCKVSSYAIAVNRCSSILFLTA<br>LSMERYLVIKKVLDTKMMGSQRHIRITCGIIWAASLLLGAPSLVYRRLDGDDCWDEDGEDFSLAMVFLTFLLPLGVISFC<br>YCSIYCRLQRHVRLGRGIRRSHRAIVTIVAAFLCSWLPLNTCKVLLFFLAKGTLVLSQGQEVALRWVVAGSTCLAFVNSC<br>VNPLVYALMDGRCRPRCPRAPGTGPGATVPSSTTDSSLLFGVRIWIKPPPGPRRRSGTQLRAAPGVPRDHLGESPQPCSP<br>ASPGATPDPAVPPGP |
| | *Cygnus atratus* (black swan) | Gene ID: 118259551<br>mRNA ID: XM_035569177<br>Protein ID: XP_035425070 | MAPEATEEPEASADYEYLAGSNATGSPEQLFTTVFIPVLYSFIFLLGFAGNLFVIVLLAKRSGGRRAVDTFVLNLAVADV<br>IFVCTLPLWVAAGARGNRWLLGEGLCKASSYAISVNRCSSILFLTALSVERYLVIWKVLDTRRAGSRRHVLLTCGLIWAI<br>SLLLGAPALRYRRLDGDDCWDEDGEDFSLAMVFLTFLLPLGVILFCYCSIYCRLQRHVRLGRGVRRSHRVIVAIVAAFVC<br>SWLPLNACKVLLFLLAKGTLALPAGQEGTLRWVVAASTCLAFANSCVNPLIYALMDPRLRCPPPPGARTPPSTTASSPSP<br>GSGTKPPPAHSPRPGT |
| | *Cygnus columbianus* (tundra swan) | Gene ID: 309910393<br>mRNA ID: XM_082830814<br>Protein ID: XP_082686940 | MAPEATEEPEASADYEYLAGSNATGSPEQLFTTVFIPVLYSFIFLLGFAGNLFVIVLLAKRSGGRRAVDTFVLNLAVADV<br>VFVCTLPLWVAAGARGNRWLLGEGLCKASSYAISVNRCSSILFLTALSVERYLVIWKVLDTRRAGSRRHVLLTCGLIWAI<br>SLLLGTPALWYRRLDGDDCWDEDGEDFSLAMVFLTFLLPLGVILFCYCSIYCRLQRHVRLGRGVRRSHRVIVAIVAAFVC<br>SWLPLNACKVLLFLLAKGMLALPAGQEGTLRWVVAASTCLAFANSCVNPLIYALMDPRLRCPLPPGARTPPSTTASSPSP<br>GSGTKPPPAPSPRPGT |
| | *Cygnus olor* (mute swan) | Gene ID: 121059087<br>mRNA ID: XM_040535442<br>Protein ID: XP_040391376 | MAPEATEEPEASADYEYLAGSNATGSPEQLFTTVFIPVLYSFIFLLGFAGNLFVIVLLAKRSGGRRAVDTFVLNLAVADV<br>VFVCTLPLWVAAGARGNRWLLGEGLCKASSYAISVNRCSSILFLTALSVERYLVIWKVLDTRRAGSRRHVLLTCGLIWAI<br>SLLLGAPALWYRRLDGDDCWDEDGEDFSLAMVFLTFLLPLGVILFCYCSIYCRLQRHVRLGRGVRRSHRVIVAIVAAFVC<br>SWLPLNACKVLLFLLAKGTLALPAGQEGTLRWAVAASTCLAFANSCVNPLIYALMDPRLRCPPPPGARTPPSTTASSPSP<br>GSGTKPPPAHSPRPGT |

| | | | |
|---|---|---|---|
| | *Dromaius novaehollandiae* (emu) | Gene ID: 135323481<br>mRNA ID: XM_064498340<br>Protein ID: XP_064354410 | MGPAAAEDLVPSADYDYAAWANATGTEEPFSRWESSYTTVFIPILYSFIFLLGFAGNLFVIVLMAKKRGSKRMVDTFVLNLAVADMIFVCTLPFWVVAGACGNRWQLGEGLCKLSSYAITVNRCSSILFLTALSVERYLVIKKVLDTKTTGTKKHIFVTCGLIWVASLLLGAPSLEYRRLDGDDCWDEDGEDFSLAMVFLTFILPLAVILFCYCSIYYRLHRHIRLGKGMRNSHRIIFTIIGAFVCSWLPLNVCKVLLFFIAKGMLVLSEAEEEALRWVVTSSTCLAFMNSCINPVIYTLMDRHFRRRLLPPALRALCLSPAEKGLNAATTSSVTESSLMFGTRTRPPAPLGHQPAA |
| | *Dryobates pubescens* (Downy woodpecker) | Gene ID: 104298804<br>mRNA ID: XM_009898901<br>Protein ID: XP_009897203 | MTPEDLLASADYDYLPLPTATGSQELLSGWEVFFTTVFIPVLYSFIFLLGLAGNLFVILLTARRRAGRRLLETLVLNLAVADAVFVCTLPLWVAAGARGNRWLLGEGLCKVSSYVVAVNRCSSILFLTALAVERYLLMRKVRDARLLGARRHVRLACAAIWAASLLLGSPALLYRQLDGEDCWDEDGEDFGLAMVFLSFLLPSAVISFCYCSICCRLRRRRRLGRGVRRSHRTIAAILAAFLCSWLPLNACKVLLFFLAKGTLVLSQGQEVALRWVVAGSTCLAFANSCVSPLLYALMEGHCQPCCPSGTAPRLPAPPSTSSSSLLFSARIWTKTSPDPQRRSRTKPQAAPGTPRGPPFPSPPSPGATSPPWPRAGLRVPL |
| | *Egretta garzetta* (little egret) | Gene ID: 104132711<br>mRNA ID: XM_035901701<br>Protein ID: XP_035757594 | MAPEELAGSADYDYPPVTSVTGNWEVSSGWEVFFTSVFIPILYSFIFLLGLVGNLFVIVLMAKKSGGKRMVDTFVLNLAVADVIFVCTLPFWVAAGVRGNRWPLGEGLCKVSSYVIAVNRCSSILFLTALSVERYLVIRKVLDTKMMGSQRHIHITCGIIWAASLLLGTPSLVYRQLDGDDCWDEDGEDFSLAMVFLTFLLPLGVISFCYCSIYCQLQRHVRLGKGVRRSHRAIITIVAAFLCSWLPLNACKVLLFFLAKGTLVLSQRQEVALRWVVAVSTCLAFINSCINPLIYALMDRRCRPRCPFGPHAPGTGPGTAAPSSTTDSSLLFGGWELTGTLPCEYPLKAGEKPPKVRRRIGAAFKLDETLVLRGADPLSALERDLALQLQIAKAARRLCREENITKAPQYALLGRCTLIVIPRMCPDTSSPVVAPMRMHPKTSSPVHAPLYTIPC |
| | *Falco biarmicus* (lanner falcon) | Gene ID: 130159778<br>mRNA ID: XM_056361775<br>XP_056217750 | MLFLSRYQINNPTAGCGKHHPGHWGGHLATTMAPEELSGSADYDYPPVTNVMVNEEVFSRWEVFFTTVFIPILYTFIFLLGLVGNLFVIMLMAKKSGGKRVVDTFVLNLAVADIIFICTLPFWVVAGAQGNRWLLGEGLCKVSSYAIAVNRCSSILFLTALSMERYLVIRKVLDTKMVGSQRYIRITCAIIWTVSLLLGAPSLVYRRLDGDDCWDEDGEDFSLAMVFLTFLLPLGIITFCYCSIYCRLQRHIRLGRGVQRSHRAIITIVAAFLCSWLPLNTCKVLLFFLAKGMLVLSQGQEVALRWVVASSTCLAFVNSCVNPLVYALMDGRCRPRCPFGPCTLGTGPSTAARSSTTDSSLLFGGRIWTKTPAGPQHRAGTELQLAPAVPRGHPGVNTPPCSPVAPGATAVPAVLPGPQL |
| | *Falco cherrug* (Saker falcon) | Gene ID: 114018094<br>mRNA ID: XM_027816707<br>Protein ID: XP_027672508 | MLFLSRYQINNPTAGCGKHHPGHWGGHLATTMAPEELSGSADYDYPPVTNVMVNEEVFSRWEVFFTTVFIPILYTFIFLLGLVGNLFVIMVMAKKSGGKRVVDTFVLNLAVADIIFICTLPFWVVAGAQGNRWLLGEGLCKVSSYAIAVNRCSSILFLTALSMERYLVIRKVLDTKMVGSQRYIRITCAIIWTVSLLLGAPSLVYRRLDGDDCWDEDGEDFSLAMVFLTFLLPLGIITFCYCSIYCRLQRHIRLGRGVQRSHRAIITIVAAFLCSWLPLNTCKVLLFFLAKGMLVLSQGQEVALRWVVASSTCLAFVNSCVNPLVYALMDGRCRPRCPFGPCTLGTGPSTAARSSTTDSSLLFGGRIWTKTPAGPQHRAGTELQLAPGVPRGHPGVNTPPCSPVAPGATAVPAVLPGPQL |
| | *Falco naumanni* (lesser kestrel) | Gene ID: 121098449<br>mRNA ID: XM_040616407<br>Protein ID: XP_040472341 | MAPEELSGSADYDYPPVTNVMVNEEVFSRWEVFFTTVFIPILYTFIFLLGLVGNLFVIMLMAKKSGGKRVVDTFVLNLAVADIIFICTLPFWVVAGAQGNRWLLGEGLCKVSSYAIAVNRCSSILFLTALSMERYLVIRKVLDTKMVGSQRYIRITCAIIWTVSLLLGAPSLVYRRLDGDDCWDEDGEDFSLAMVFLTFLLPLGIITFCYCSIYCRLQRHIRLGRGVQRSHRAIITIVAAFLCSWLPLNTCKVLLFFLAKGMLVLSQGQEVALRWVVASSTCLAFVNSCVNPLVYALMDGRCRPRCPFCPCALGTGPSTAPSSTTDSSLLFGGRIWTKTPAGPRHRAGTELQLAPGVPRGHPGVNTPPCSPAAPGATAVPAVLPGPQL |
| | *Falco peregrinus* (peregrine falcon) | Gene ID: 101916944<br>mRNA ID: XM_005239686<br>Protein ID: XP_005239743 | MLFLSRYQINNPTAGCGKHHPGHWGGHLATTMAPEELSGSADYDYPPVTNVMVNEEVFSRWEVFFTTVFIPILYTFIFLLGLVGNLFVIMLMAKKSGGKRVVDTFVLNLAVADIIFICTLPFWVVAGAQGNRWLLGEGLCKVSSYAIAVNRCSSILFLTALSMERYLVIRKVLDTKMVGSQRYIRITCAIIWTVSLLLGAPSLVYRRLDGDDCWDEDGEDFSLAMVFLTFLLPLGIITFCYCSIYCRLQRHIRLGRGVQRSHRAIITIVAAFLCSWLPLNTCKVLLFFLAKGMLVLSQGQEVALRWVVASSTCLAFVNSCVNPLVYALMDGRCRPRCPFGPCTLGTGPSTAARSSTTDSSLLFGGRIWTKTPAGPQHRAGTELQLAPGVPRGHPGVNTPPCSPVAPGATAVPAVLPGPQL |
| | *Falco punctatus* | Gene ID: 308054518<br>mRNA ID: XM_080367494<br>Protein ID: XP_080223620 | MLFLSRYQINNPTAGCGKHHPGHWGGHLATAMAPEELSGSADYDYPPVTNVMVNEEVFSRWEVFFTTVFIPILYTFIFLLGLVGNLFVIMLMAKKSGGKRVVDTFVLNLAVADIIFICTLPFWVVAGAQGNRWLLGEGLCKVSSYAIAVNRCSSILFLTALSMERYLVIRKVLDTKMVGSQRYIRITCAIIWTVSLLLGAPSLVYRRLDGDDCWDEDGEDFSLAMVFLTFLLPLGIITFCYCSIYCRLQRHIRLGRGVQRSHRAIITIVAAFLCSWLPLNTCKVLLFFLAKGMLVLSQGQEVALRWVVASSTCLAFVNSCVNPLVYALMDGRCRPRCPFCPCALGTGPSTAAPSSTTDSSLLFGGRIWTKTPAGPRHRAGTELQLAPGVPRGHPGVNTPPCSPAAPGATVVPAVLPGPQL |
| | *Falco rusticolus* (gyrfalcon) | Gene ID: 119158391<br>mRNA ID: XM_037410213<br>Protein ID: XP_037266110 | MAPEELSGSADYDYPPVTNVMVNEEVFSRWEVFFTTVFIPILYTFIFLLGLVGNLFVIMVMAKKSGGKRVVDTFVLNLAVADIIFICTLPFWVVAGAQGNRWLLGEGLCKVSSYAIAVNRCSSILFLTALSMERYLVIRKVLDTKMVGSQRYIRITCAIIWTVSLLLGAPSLVYRRLDGDDCWDEDGEDFSLAMVFLTFLLPLGIITFCYCSIYCRLQRHIRLGRGVQRSHRAIITIVAAFLCSWLPLNTCKVLLFFLAKGMLVLSQGQEVALRWVVASSTCLAFVNSCVNPLVYALMDGRCRPRCPFGPCTLGTGPSTARSSTTDSSLLFGGRIWTKTPAGPQHRAGTELQLAPGVPRGHPGVNTPPCSPVAPGATAVPAVLPGPQL |
| | *Gallinula chloropus* (common moorhen) | Gene ID: 311201422<br>mRNA ID: XM_084275811<br>Protein ID: XP_084131937 | MAPEELTSSGDYDYPPVTNVTGDWEVPSRWELFFTTVFIPIFYSFIFLLGLVGNLFVIVLMAKKSRGKRMVDTFVLNLAVADVIFTCTLPFWVVAGARGNRWPLGEGLCKVSSYVIAVNRCSSILFLTALSVERYLVIRKVLDTKVLGSQRHVRITCGIIWAASLILGAPSLVYRQLDGEDCWDEDGEDFNLTMVFLTFLLPLGVISFCYCSIYCRLQRHVRLGRGVRRSHRAIVTIVAAFLCSWLPLNACKVLLFFLAKGTLVLSQGQEVALRWVVAGSTCLAFVNSCINPLLYALMDRRCRPRCPFGPRAPGTGPGAATTSSTTDSSLFFGGWIWTKPPPAPRCRSGTELQPAPRSPRGHP |
| | *Gallus gallus* (chicken) | Gene ID: 770582<br>mRNA ID: XM_015275387<br>Protein ID: XP_015130873 | MEYAEADYYYGEAEEEEEGNGTACAWQADWEASFALLPALYALVFVLGLAGNALVLLTVWRGPRAKRRSADAYIGNLALADLAFVATLPLWAAYTALRFHWPFGAALCKLSSYLVLLNMFASAFCLGGLSAERYRAVLRAAPPPPRAASARLRRPAALGPLAALWAAAAAAALPALLLREARRDARNRTLCDLRLGGEGGGAERAAAALSLGTTALGFAAPLLLMAVCYCCVGSAVRRHLRPRRAEAAARRRLLRLIAALVGVFAGCWLPFHLLKSLFVLAAAGLLELPCALLGLISRLHPYATCLAYLNSCLNPLLYAFLDGRFRAQCRALLGLRGARPPPAASSTPSAPTQRSELPSSGGTKV |
| | *Grus americana* (Whooping crane) | Gene ID: 129196429<br>mRNA ID: XM_054803886<br>Protein ID: XP_054659861 | MAPEELASSADYDYPPVTNVTGNWEVSGWEVFFTTVFIPILYSFIFLLGLLGNLFVIVLMAKKSGGKRMVDTFVLNLAVADVIFICTLPFWVVAGARGNRWPLGEGLCKVSSYAIAVNRCSSILFLTALSVERYLVIRKVLDTKMMGSRRHVRVTCGIIWAASLILGAPSLVYRQLDGDDCWDEDGEDFSLTMVFLTFVLPLGVISFCYCSIYCRLQRHVRLGRGVRRSHRAIVTIVVAFLCSWLPLNACKVLLFFLAKGTLVLSQGQEMALRWVVAGSTCLAFVNSCVNPLIYALMDGHCRPRCPFGLCTPGTSPGAAATSSTTDSSLLFGGRIRTKTPPGPWHRSGTELQPAPGIPRGHS |
| | *Grus grus* (Eurasian crane) | Gene ID: 308709521<br>mRNA ID: XM_081398999<br>Protein ID: XP_081255125 | MAPEELASSADYDYPPVTNVTGNWEVSGWEVFFTTVFIPILYSFIFLLGLLGNLFVIVLMAKKSGGKRMVDTFVLNLAVADVIFICTLPFWVVAGARGNRWPLGEGLCKVSSYAIAVNRCSSILFLTALSVERYLVIRKVLDTKMMGSRRHVRVTCGIIWAASLILGAPSLVYRRLDGDDCWDEDGEDFSLTMVFLTFVLPLGVISFCYCSIYCRLQRHVRLGRGVRRSHRAIVTIVVAFLCSWLPLNACKVLLFFLAKGTLVLSQGQEMALRWVVAGSTCLAFVNSCVNPLIYALMDGHCRPRCPFGPCTPGTSPGAAATSSATDSSLLFGGRIRTKTPPGPWHRSGTELQPAPGIPRGHS |
| | *Haliaeetus leucocephalus* (bald eagle) | Gene ID: 104842433<br>mRNA ID: XM_010584763<br>Protein ID: XP_010583065 | MAPEELASSADYDYPPVTNVTGNQEVFSRWEVFFTTVFIPILYSFIFFLGLVGNLFVIVLMAKKSGGKRMVDTFVLNLAVADVIFICTLPFWVVAGARGNRWLLGEGLCKVSSYAIAVNRCSSILFLTALSVERYLVIRKVLDTKMMGSQRHVHVTCGIIWTVSLLLGAPSLVYRQLDGDDCWDEDGEDFSLAMVFLTFLLPLGVISFCYCSIYCQLQRHVRLGRGVRRSHRAIVTIVAAFLCSWLPLNACKVLLFFLAKGTLVLSQGQEVALRWVVAGSTCLAFVNSCVNPLVYALMDRRCRPQCPFSARALGTGPGAA |

| | | | |
|---|---|---|---|
| | | | TPSSATDSSLLFGGRIRTKTPTGPQHRSGTELQLALGVLRGHPRVSPPPSSPASPGATPISPVPPGPQQ |
| | *Harpia harpyja* (harpy eagle) | Gene ID: 128154745<br>mRNA ID: XM_052815786<br>Protein ID: XP_052671746 | MAPEELAGSADYDYPPVTNVTGNQEVFSRWEVFFTTVFVPVLYSFIFLLGLVGNLFVIVLMAKKSGGKRMVDTFVLNLAV<br>ADVIFICTLPFWVVAGARGNRWLLGEGLCKVSSYAIAVNRCSSILFLTALSVERYLVIRKVLDTKMMGSQRHVHVTCGII<br>WTVSLLLGAPSLVYRQLDGDDCWDEDGEDFSLAMVFLTFLLPLGVISFCYCSIYCQLQRHVRLGRGVRRSHRAIVTIVAA<br>FLCSWLPLNACKVLLFFLAKGTLVLSQGQEVALRWVVAGSTCLAFVNSCVNPLVYALMDRRCRPQCPFGPRAPGTGPGTA<br>TPSSATDSSLLFGGWIRTKTTTGPRRRSGTELQPALGVLRGHPRVSPPPSSPASPGATAVPPVPPGPQL |
| | *Larus michahellis* (yellow-legged gull) | Gene ID: 141733724<br>mRNA ID: XM_074564507<br>Protein ID: XP_074420608 | MAPEELAASADYDYPPVTNATGNGEVFSRWEVFFTTVFIPVLYSFIFLLGLAGNLFVIALMAKKRRGQRAVDTFVLNLAV<br>ADVIFVCTLPFWVAAGARGNRWPLGEGLCKASSYAIAVNRCSSILFLTALSVERYLVIRKVLDTKRTGSQRHVRLTCGFI<br>WATSLLLGVPALLYRRLDGDDCWDEDGEDFSLAMVFLTFLLPLGVISFCYCSIYCHLQRHVRLGRGVRRSHRAIVTIVAA<br>FLCSWLPLNACKVLLFFLAKGTLALSQGQEVALRWVVAASTCLAFVNSCVNPLVYALMDGRCRPRCPLGPRSSGTAPGVA<br>VTSSTTDSSLGGRIRTKPPPAPGAGPSSGRHRGSRGATPG |
| | *Melanerpes formicivorus* (acorn woodpecker) | Gene ID: 137454565<br>mRNA ID: XM_068164648<br>Protein ID: XP_068020749 | MAPEELLASADYDYPPLPTATGSQELLSGWEVFFTTVFIPVLYSFVFLLGLAGNLFVILLTARRRAGRRLLETLVLNLAV<br>ADAVFVCTLPLWVAAGARGNRWLLGEGLCKVSSYVVAVNRCSSIFFLTALAVERYLLMRKVRDARLLGTRRHVRLTCAAI<br>WAASLLLGSPALLYRQLDGEDCWDEDGEDFGLAMVFLSFLLPSAVISFCYCSICCRLRRRRRLGRGVRRSHRTIAVILAA<br>FLCSWLPLNACKLRQPPPLCPDGRALPALLPLLPLGDGSSPASSSLHLLLQPPLQRPHLDQDLPRPPAQEQDQAPGGTGD<br>PTRSSLPLPNVPRCHQPPVASAGASRPAVIPEAGSGPWDPFPLPPFPLAPLPPCPIPSLPHNPLHPSSPLPIYLELTQIY<br>SQ |
| | *Mycteria americana* (wood stork) | Gene ID: 142419084<br>mRNA ID: XM_075521703<br>Protein ID: XP_075377818 | MAPEEFASSADYDYPPVTNVTGNREVSSGWEVFFTSVFIPILYSFIFLLGLVGNLFVIVLMAKRSGGKRMVDTFVLNLAV<br>ADVIFVCTLPFWVAAGARGNRWLLGEGLCKVSSYAIAVNRCSSILFLTALSVERYLVIRKVLDTKMMGSQRHVRITCGII<br>WAASLLLGAPSLVYRRLDGDDCWDEDGEDFSLAMVFLTFLLPLVVISFCYCSIYCRLQRHVRLGRGVRRSHRAIVTIVAA<br>FLCSWLPLNACKVLLFFLAKGTLVLSQGQEGALRWVVAGSTCLAFVNSCVNPLVYALMDGRCRPRCPFGPCAPGTGPGTA<br>APSSATDSSLLFGGWIWTKTPPGPRRRSRTELQPAPGVPRGHPAVSSPPSSPASPGATAVPAVPPGPQP |
| | *Netta rufina* (red-crested pochard) | Gene ID: 311219015<br>mRNA ID: XM_084301965<br>Protein ID: XP_084158091 | MAPEATEEPEASADYEYLAGSNATGSLEQLFTTVAIPVLYSFIFLLGFAGNLFVIVLLAKRSGGRRAVDTFVLNLAVADV<br>VFVCTLPFWVAAGARGNRWLLGEGLCKASSYAISVNRCSSILFLTALSVERYLVIWKVLDTRRAGSRRHVLLACGLIWAV<br>SLLLGAPALWYRRLDGDDCWDEDGEDFSLAMVFLTFLLPLGVILFCYCSIYCRLQRHVRLGRGVRRSHRVIVAIVAAFVC<br>SWLPLNACKVLLFLLAKGTLVLPAGQEGTLRWVVATSTCLAFANSCVNPVIYALMDPRLRCPPALGARTPPSTTASSPSP<br>GSGTKLPPVPSPRPGTVLCAVLCHAHPCHAVLPHAALSRALQPAGPAAAGPVLLTPLLPCVSSQNTSPQPGRSRAAASAR<br>ARSPATPAPRCGSSHWGN |
| | *Nipponia nippon* (crested ibis) | Gene ID: 104022178<br>mRNA ID: XM_009475891<br>Protein ID: XP_009474166 | MLFLSRFQINNPTAERGEHRPGHRGGHLATAMAPEELVGSADYDYSPVTNVTGNWEVSSRWEVFFTSVFIPIFYSFIFLL<br>GLVGNLFVIVLMAKKGRGKRMVDTFVLNLAVADVIFVCTLPFWVAAGARGNRWLLGEGLCKVSSYAIAVNRCSSILFLTA<br>LSVERYLVIRKMLDTKMMGSRRHVRVTCGIIWAASLLLGAPSLAYRRLDGDDCWDEDGEDFSLVMVFLTFLLPLGVISFC<br>YCSIYCRLQRHVRLGRGVRRSHRAIVTIVAAFLCSWLPLNTCKVLLFFLAKGTLVLSQGQEVALRWVVAGSTCLAFVNSC<br>INPLIYALMDGRCRPRCPFGPRAPGTGPGAATPSSATDSSLLFGSRIRTRTLPGAGVGPSSGRHRGSHGATPG |
| | *Nyctibius grandis* | Gene ID: 137674290<br>mRNA ID: XM_068419537<br>Protein ID: XP_068275638 | MAPEELASSADYDYPPVTDVTGNLEVSSGWEVIFTTVFIPILYSVVFLLGLAGNLFVIVLMAKKSGGKRVVDTFVLNLAV<br>ADVIFVCTLPFWVASGARGNRWPLGEGLCKVSSYAIAVNRCSSILFLTALSVERYLVIRKVLDTKVTGSRRHVRVTCGAI<br>WVASLLLGAPSLLYRRLDGDDCWDEDGEDFSLAMVFLTFLLPLGVISFCYCSIYCRLQRHVRLGRGVRRSHRAIVTIVAA<br>FLCSWLPLNACKVLLFFLAKGTLALSQGQEVALRWLVAGSTCLAFVNSCVNPLVYALMDGRCRPRCPLLSPRGRGTAPAA<br>TSSSATDSSLLCGGRGRTKNPPGPRRRTGAQHQPAPGVPWGHPGVSPPPSSSSPLSPGATKVPVVPPEPRP |
| | *Opisthocomus hoazin* | Gene ID: 104337075<br>mRNA ID: XM_075442898<br>Protein ID: XP_075299013 | MAPEELSGSADYDYPPVTNTTENQELPSKWEFFFTTVFIPILYSFIFLLGLVGNLFVIVLMAKKSGSKRMVDTFVLNLAV<br>ADVIFVCTLPFWVAAGARGNRWLLGEGLCKVSSYAIAVNRCSSILFLTALSVERYLVIRKVLDTKMMGSQRHVRVTCGII<br>WAASLLLGAPSLVYRQLDGDDCWDEDGEDFSLAMVFLTFLLPLGVISFCYCSIYCRLQRHVRLGRGVRRSHRAIVTIVAA<br>FLCSWLPLNACKVLLFFLAKGMLVLSQGQEVALRWVVAGSTCLAFVNSCVNPLVYALMDGRCRPRCPLRPRAPGTSPTTA<br>APSSATDSSLLFGGRNRTKTLPGPRRRSGTELQPAPGVPRGHPGASPPLPSPASPGATAVPAVPPGPRP |
| | *Patagioenas fasciata* (band-tailed pigeon) | Gene ID: 136110692<br>mRNA ID: XM_065854280<br>Protein ID: XP_065710352 | MAPEELITSADYEYPPVTSVMENPWVSSRAELFFTTIFIPILYSIIFLLGLVGNLFVIVLMAKKSRGKRMVDTFVLNLAV<br>ADVVFVCTLPFWVAAGAQGNRWLLGEGLCKLSSYIIAVNRCSSILFLTALSVERYLVIRKVLDTKVMGSQRHVRVTCGVI<br>WAVSLLLGAPTLLYRRLDGDDCWDEDGEDFSLAMVFLTFLLPLGVISFCYCSIYCRLQRHVRLGRGVRRSHRAIVTIVTA<br>FLCSWLPLNACKVLLFFLAKGMLVLSQGQEVALRWLVATSTCLAFVNSCVNPLVYALMDGRCRPRCPLGPRALGTGPGAT<br>APSSTSDSSLLFGVWTRSRCRTRHQPAPWGSPLSPLPSSPGATAVPLGPQL |
| | *Pelecanus crispus* (Dalmatian pelican) | Gene ID: 142595121<br>mRNA ID: XM_075722567<br>Protein ID: XP_075578682 | MGPEELAGSAYYDYPPVTNVTENWEVSSGWEVFFTSVFIPILYSFIFLLGLVGNLFVIVLMAKKSGGKRMVDTFVLNLAV<br>ADVIFVCTLPFWVVAGARGNRWLLGEGLCKVSSYAIAVNRCSSILFLTALSVERYLVIRKVLDTKMTGSQRHIRVTCGLI<br>WAASLLLGAPALAYRRLEGDDCWDEDGEDFSLAMVFLTFLLPLGVISFCYCSIYCRLQRHIRLGRGIRRSHRAIVTIVAA<br>FLCSWLPLNACKVLLFFLAKGTLVLSQGQEVALRWVVAGSTCLAFVNSCINPLVYALMDGRCRPRCPFHPGAPGTGPGAA<br>SPSSATDSSLLFGGWIRTRTVPGPRRRSGTELQPAPGVPRGRPGVSPPRSSPASPGATAVPAVPPGPQP |
| | *Phaenicophaeus curvirostris* | Gene ID: 138730563<br>mRNA ID: XM_069875847<br>Protein ID: XP_069731948 | MLFFSRSHINNPTAGHGQHRPGHQGGHPATSMAPEELSGSADYDYPLVTNTTVNQEVFSKWEVIFTTIFIPILYSFIFLL<br>GLMGNLFVIVLMAKRSGNKRMVDTFVLNLAVADVIFICTLPFWVVAGARGNRWPLGEGLCKVSSYAIAVNRCSSILFLTA<br>LSVERYLVIRKVLDTKMMGSRRHVHITCGIIWAASLLLGAPSLVYRRLDGDDCWDEDGEDFSLAMVFLTFLLPLGVISFC<br>YCSIYCRLQRHVRLGRGVQRSHRAIVTIVTAFLCSWLPLNTCKVLLFFLAKGTLVLSQGQEVALRWVVAGSTCLAFVNSC<br>INPLVYALMDGRCRPRCPRAPGTGPGAAAPSSTTDSSLLFGVRIWTRSPPGPRCRSRTQLRAALGVPRDHPGVSPQPCSE<br>PSPGATSDSAPQP |
| | *Phalacrocorax carbo* (great cormorant) | Gene ID: 135316972<br>mRNA ID: XM_064472453<br>Protein ID: XP_064328523 | MAPEELAGSADYDYPPVTNETGNQEVSSGWEVFFTSVFIPILYSFIFVLGLVGNVFVIVLMAKKGGGKRMVDTFVLNLAV<br>ADVIFVCTLPFWVVAGARGNRWLLGEGLCKVSSYAIAVNRCSSILFLTALSVERYLVIRKVLDTKVMGSRRHIHVTCGII<br>WVASLLLGAPSLVYRRLDGDDCWDEDGEDFSLAMVFLTFLLPLGVISFCYCSIYCQLQRHVRLGRGVQRSHRAIVTIVAA<br>FLCSWLPLNACKVLLFFLAKGTLVLSQGQEVALRWVVAGSTCLAFINSCVNPLIYALMDGRCRPRCPFGPRVPGMGPGAT<br>AHSSATDSSLLFSGWIWTKTTPGPRHRSGTELQLAPGVPRVHSGVSPSTPSPASPGAAAIPAVPPSPQL |
| | *Platalea leucorodia* | Gene ID: 309598907<br>mRNA ID: XM_082329934<br>Protein ID: XP_082186060 | MAPEELVGSADYDYSPVTNVTGNWEVSSRWEVFFTSVFIPIFYSFIFLLGLVGNLFVIVLMAKKGRGKRMVDTFVLNLAV<br>ADVIFVCTLPFWVAAGARGNRWLLGEGLCKVSSYAIAVNRCSSILFLTALSVERYLVIRKMLDTKMMGSRRHVRVTCGII<br>WAASLLLGAPSLAYRRLDGDDCWDEDGEDFSLVMVFLTFLLPLGVISFCYCSIYCRLQRHVRLGRGVRRSHRAIVTIVAA<br>FLCSWLPLNTCKVVLFFLAKGTLVLSQGQEVALRWVVAGSTCLAFVNSCINPLIYALMDGRCRPRCPFGPHAPGTGPGAA<br>TPSSATDSSLLFGGRIRTKTLPGGSGTELRPAPGVPRGHPGVSPPPFLPSVPGCHRRPRRPTRSPAVTQAQPIPARFPRP<br>LPPRALPAASAQAGSDPSIAPLRPIPPPRDALTRDGFGSGNLVLIPI |
| | *Pogoniulus pusillus* (Red-fronted | Gene ID: 135191521<br>mRNA ID: XM_064173873<br>Protein ID: XP_064029943 | MAPEELMASADYDYSLLANATGSQEALSSWDVFFTTVFIPILYSLIFLLGLVGNLFVVALLAVRSGGERAVDTFVLNLAL<br>ADVIFICTLPFWVAAGAQGNRWLLGEGLCKLSSYVIAVNRCSSILFLSALSVERYLVMRRVLDTRLVGARGHVRATCGLI<br>WAASLLLGAPALVYRQLDGEDCWDEDGEDFSLAMVFLTFLLPLVVISFCYCSISCRLQRHVRLGRGVRRSHRAILTIVTA |

| | | | |
|---|---|---|---|
| | tinkerbird) | | FLCSWLPLNACKVLLFLLAKGTLSLSQGQAVALRWVVAGSTCLAFLNSCVNPLLFALMEPRCRQGCARGWPLAPCALGTAPAAPAPSSSSLFFGGWIWTKPPHGRQGGSRTKLRLAPGIPRGPCPCPVSPGAAAPPRPQPAACSPV |
| | *Rhea americana* (greater rhea) | Gene ID: 146596338<br>mRNA ID: XM_085304152<br>Protein ID: XP_085160278 | MGPDAAEELTASADYDYPAEVNSTGTEEHFSRWELSYMTIFIPILYSFIFLLGFVGNLFVIMLMAKKRGNKRMVDTFVLNLAVADMIFVCTLPFWVVAGACGNRWQLGEGLCKLSSYAITVNRCSSILFLTALSVERYLVXRKVLDTKMTGSKRHIFVTCGLIWVVSLLMGAPSLEYRRLDGDDCWDEDGEDFSLAMVFLTFILPLVVILFCYCSIYYRLHRHIRLGKGMRNSHRIIFTIVGAFVCSWLPLNVCKVLLFFIAKGMLLPSEAEEEALRWLVTSSTCLAFMNSCINPIIYTLMDRHFRRQLLPPRLRVLCLSPAEKGHSVATTSSVTESSLMFGVRTKPPAPPGQQPAP |
| | *Streptopelia decaocto* (Eurasian collared-dove) | Gene ID: 309447965<br>mRNA ID: XM_082069108<br>Protein ID: XP_081925234 | MAPEERVTSADYEYPPVTSAMENQWVSSRGELFFTTIFIPILYSIIFLLGLVGNLFVIVLMAKKSRGKRMVDTFVLNLAVADVVFVCTLPFWVAAGAQGNRWLLGEGLCKLSSYVIAVNRCSSILFLTALSVERYLVIRKVLDTKVMGSQRHVRVTCGVIWAVSLLLGAPTLLYRRLDGDDCWDEDGEDFSLAMVFLTFLLPLGVISFCYCSIYCRLQRHVRLGRGVRRSHRAIVTIVTAFLCSWLPLNACKVLLFFLAKGMLVLSQGQEVALRWLVATSTCLAFVNSCVNPLIYALMDGHCQPRCPLGPRAPETGPGAAAPSSTSDSSLLFGVWTRSRCRTRHQPAPGGTPLSPPASSPGATAVPPGPQL |
| | *Strix aluco* (Tawny owl) | Gene ID: 141934628<br>mRNA ID: XM_074850252<br>Protein ID: XP_074706353 | MAPEELAGSADYDYPLVTNGTGNREVSSRWEVFFTTVFIPILYSFIFLLGLVGNLFVIVRMGKKSGGKRMVDTFVLNLAVADVIFVCTLPFWVVAGARGNRWSLGEGLCKVSSYAIAVNRCSSILFLTALSVERYLVIRKVLDTKMVGSRRHIHVTCGIIWTVSLLLGAPSLVYRRLDGDDCWDEDGEDFSLAMVFLTFLLPLGVISFCYCSIYCRLQRHVRLGRGVRRSHRAIVTIVAAFLCSWLPLNACKVLLFFLAKGTLVLSQGQEVALRWVVAGSTCLAFVNSCVNPLVYALMDGRCRPRCPFGPCATGTGPAVAAPSSTTDSSLFGGRIWTKTPPRPPAQERD |
| | *Strix uralensis* (Ural owl) | Gene ID: 141954146<br>mRNA ID: XM_074893389<br>Protein ID: XP_074749490 | MAPEELAGSADYDYPPVTNGTGNREVSSRWEVFFTTVFIPILYSFIFLLGLVGNLFVIVRMGKKSGGKRMVDTFVLNLAVADVIFVCTLPFWVVAGARGNRWSLGEGLCKVSSYAIAVNRCSSILFLTALSVERYLVIRKVLDTKMVGSQRHIHVTCGIIWTVSLLLGAPSLVYRRLDGDDCWDEDGEDFSLAMVFLTFLLPLGVISFCYCSIYCRLQRHVRLGRGVRRSHRAIVTIVAAFLCSWLPLNACKVLLFFLAKGTLVLSQGQEVALRWVVAGSTCLAFVNSCVNPLVYALMDGRCRPRCPFGPCATGTGPAVAAPSSTTDSSLFGGRIWTKTPPGPRRRSGTELRPAPGVSPPLSSSSASPGATAVPAVPPGPQP |
| | *Struthio camelus* (African ostrich) | Gene ID: 104143991<br>mRNA ID: XM_009674745<br>Protein ID: XP_009673040 | MGPQAAEDFVLSADYDYPVGTKSTGTEERFSRWESSYMTVFIPILYSFIFLLGFVGNLFVIVLLAKKRGSKRMVDTFVLNLAVADVIFVCTLPFWVVAGAHGNRWQLGEGLCKVSSYAITVNRCSSILFLTALSVERYLVIRKVLDTKMTGSKRHVFVTCGLIWVVSLLLGAPSLEYRRLDGDDCWDEDGEDFSLAMVFLTFLLPLAVILFCYCSIYYQLHRHIRLGKGMRNSHRIIFTIVGAFICSWLPLNVCKVLLFFIAKGMLVLSEVEEEALRWVVTSSTCLAFMNSCVNPIIYTLMDRHFRRQLLPPALRVLCLSPAEKGHNMATTSSVTESSLMFGVRTKPSAPPGRWPAVQEEKDWGPNE |
| | *Tauraco erythrolophus* (red-crested turaco) | Gene ID: 104375743<br>mRNA ID: XM_009982160<br>Protein ID: XP_009980462 | MRTAWPEMELTQLSPVTTVTFNYDYYYDDNCQYQHLQHMSTFLPILYTAVFLVGIVGNSILIVALVFKRRVQRLIDIFIINLAASDFVFLITLPFWVDKEASDGSWRVGSFLCKASSYVISVNMYCSILLLTCMSADRYLAIMHPSIARRVRTRSYSSGLCICVWLLSCCLGMPTLLSRELKKQYGKTYCTDKAVTEAKQIMSLMLLILAFFFPLLSILTFYCSITRRLCVHYQRAGKHDKKLRKSIKIVFIVVAAFVISWVPFNLFKLMAILLGLLKQPDCLPNTVAQVGMKVSSPLAFANSCANPFIYYCFDNYIRRAMLRCLCPRVKISSSGSNSDTTDTRLSHSLSNFVVGEYAARKRKRSVSL |
| | *Tyto alba* (common barn owl) | Gene ID: 116961340<br>mRNA ID: XM_032992033<br>Protein ID: XP_032847924 | MLFLSRSQINNPTAGRGEHRPGHRRGHPATAMAPEELVGSADYDYPPVTNATGNWEVSSRWEIFFTTVFIPVLYSFIFLLGLVGNLFVIVLMGKKSGGKRMVDTFVLNLAVADVIFVCTLPFWVAAGARGNRWPLGEGLCKVSSYVIAVNRCSSILFLTALSVERYLVIRKVLDTKVVGSQRHVRVTCGIIWTVSLILGAPSLVYRRLDGDDCWDEDGEGFSLAMVFLTFLLPLGVISFCYCSIYCRLQRHVRLGRGVRRSHRAIVTIVAAFLCSWLPLNACKILLFFLAKGTLVLSQGQEVALRWVVAGSTCLAFVNSCVNPLVYALMDGRCRPRCPLGPCALGMGPAVAVPSSATDSSLLFGGWIRTKTPPGPRHRSGTKLPPAPGVSPPCSSPASPGATAVPPGPQP |
| | *Vanellus vanellus* | Gene ID: 309685348<br>mRNA ID: XM_082501124<br>Protein ID: XP_082357250 | MAPEELSGSADYDYPPVTNVTGNREVFSSWEVFFTTVFIPGLYSFIFLLGLVGNLFVIVLMAKKRRGKRMVDTFVLNLAVADVIFVCTLPFWVVAGARGNRWPLGEGLCKVSSYAIAVNRCSSILFLTALSVERYLVIRKVLDTKTMGSQQHVRVTCGVIWAASLLLGAPSLVYRRLDGDDCWDEDGEDFSLAMVFLTFLLPLGVISFCYCSIYCRLQRHVRLGRGVRRSHRAIVTIVAAFLCSWLPLNACKVLLFFLAKGTLVLSQGQEVALRWLVAGSTCLAFVNSCVNPLVYALMDGRCRPPRPFGPRAPGTGPGVALTSSTTDSSGLFSGRIRIKPTPPPPPRHGTWLRPAPGVPRGHPGGGVAPRQAGSDPPPEMPFTRDGCGSRNPV |
| Reptiles | *Ahaetulla prasina* | Gene ID: 131195628<br>mRNA ID: XM_058177705<br>Protein ID: XP_058033688 | MKFAELVTDLAITKNWSRDLESSTVDFFGLGFTSDDFCFMEKLPFASIFISVIYFMIFFIGFFGNLFVILVMVQKRGNQRLVDKFVLNLAIADLVFVCTLLFWALTEASDSWWFGEELCKFSSYAISVNRCSSILFLMGMSVERFLVMNKHWDSRSLGTKRNTNITCSCIWVFSFLLGIPSLLYRKLIPVGDRYACQDEEPSIIYIFTVLCLTFLLPLGVILFCYCSIFSMLQSHVNVGKKTNHALKIIFVIIGAFICCWLPFNTFTAVITYGVVQNIDFSCEALKALRKGVIISACLAFTNSCINPIIYAFMDQQFRQKVLKKFLGLCHPKENVQSSTLFFSSTESSLLFGSKKRFIPATTVPVSSGDDLYASKRQI |
| | *Alligator mississippiensis* (American alligator) | Gene ID: 132244885<br>mRNA ID: XM_059717561<br>Protein ID: XP_059573544 | MTTEAWTTGFEMSGDYELGDLCQHLKLPYAHVYIPILYFLVFFIGFIGNLFVIVLMAVKKGNKRMVDNFVLNLAVADLVFVCTLPFWALSAAHNNHWYFGESLCKVSSYIISVNRCSSILFLTGMGVERYLVVTKRVRSKSSVTKKCIFTTCGCIWAVSLLLGIPSLLYRKLNASDKFCQDEDSSAYQKYSLALLILTFLLPLGVILFCYCSICNKLLSHISVGKGTKNSLKIIFTIITTFICFWLPFNAFKAFLILSSPSELDQEASLSCSAQIAVKWGLSLSTYLAFMNSCINPIIYAFMDHHFRRRVQLHCFRFFGVSQKTTQNWVFSSSAAESSLMFASRNKPSSPMSHQPSM |
| | *Alligator sinensis* (Chinese alligator) | Gene ID: 102371715<br>mRNA ID: XM_006038971<br>Protein ID: XP_006039033 | MTTEAWTTGFEMSGDYELGDLCQHLKLPYAHVYIPILYFFVFFIGFIGNLFVIVLMAMKKGNKRMVDNFVLNLAVADLVFVCTLPFWALSAAHNNHWYFGESLCKVSSYIISVNRCSSILFLTGMGVERYLVVTKRVRSKSSVTKKCIFTTCGCIWAVSLLLGIPSLLYRKLNASDKFCQDEDSSAYQKYSLALLILTFLLPLGVILFCYCSICNKLLSHISVGKGTKNSLKIIFTIISTFICFWLPFNAFKAFLILSSPSELDQEASLSCSAQIAVKWGLSLSTYLAFMNSCINPIIYAFMDHHFRRRVQLHCFRFLGVSQKTTQNWVFSSSAAESSL |
| | *Anolis carolinensis* (green anole) | Gene ID: 100553771<br>mRNA ID: XM_062978417<br>Protein ID: XP_062834487 | MEAAQLATTSAMTENWSSNFEAYPIDYDGLDSMNICHLELPFAYIFIPILYFLSFLIGFFGNLFVILLMARKQGSKRLVDTFVLNLAIADLIFVFTLPFWALSAALDGWLFSEGLCKLSSYTISVNRCSSILFLTGMSMERFLVMTKRWDAKSIGTRKNAITSCGCIWAISLLLGIPSLVYRNLTFKGDSKYSCQGDPPATFSVVMLSLTFLLPLGVILFCYCSIFSKLRSHARLGKRRNNALKIIFTIIGAFVCSWLPFNTFKMIFTMTQNIELSCQALRTLFGGLTITACLAFVNSCINPVIYVFMDQHFRRQVLMYFPGFCKAKGNMQASSMSFSSTESSVLFGSRKKLPPATSVQI |
| | *Anolis sagrei* (brown anole) | Gene ID: 132772666<br>mRNA ID: XM_060771540<br>Protein ID: XP_060627523 | MEAAQLSTTSAMTENWSSNFEEYPTEYDGLDSMNICHLELPFAYIFIPILYFLSFFIGFFGNLFVILLMARKQGSKRLVDTFVLNLAIADLVFVFTLPFWGLSAALDGWLFSEGLCKLSSYTISVNRCSSILFLTGMSVERFLVMTKRWDARSIGTRKNAITSCGCIWAISLLLGIPSLVYRNLAFKGDSKYSCQGDPPATFSVVMLSLTFLLPLGVILFCYCSIFSKLQSHARLGKRRNNALKIIFTIIGAFVCSWLPFNTFKIIFTMAQNIDLSCQALRTLFGGLTITACLAFVNSCINPVIYVFMDQHFRQQVLMYFPGFWKAKGNMQASSMSFSSTESSVLFGSRKKLPPATSVQI |
| | *Candoia aspera* | Gene ID: 134494859<br>mRNA ID: XM_063300105<br>Protein ID: XP_063156175 | MKIAQLVTDSAMTKNWSTDLDSSSEDYYGSGFISHDICLMEKLPFASIFISVIYFMIFFIGFFGNLFVVLVMVQKQGNQRLVDNFILNLAIADLVFVCTLPFWALSEAFDSWWFGETLCKFSSYAISVNRCSSILFLMGMSVERFLVMNKHWDSRFLGTKRNTSMICSCIWVFSLLLGIPSLVYRKLILVGDKFSCQDEISAIYTFTVLFLTFLLPLGVILFCYCSIFCMLQGHVSLGKRRNHVLKIIFVIIGAFISCWLPFNTFIITYSMVKDINLSCEALKALRRGLTISACLAFTNSCINPIIYVFMDQQFRQKVLKKFLGLCRPRETVQNPALSFSSTESSLLFASRKKFIPNKTVPVSSGAGLCASKREIAAIVENIQQNC |

| | | | |
|---|---|---|---|
| | *Caretta caretta* (loggerhead turtle) | Gene ID: 125625072<br>mRNA ID: XM_048826666<br>Protein ID: XP_048682623 | MTTEPWTVNYEASSAVYSDYNPLQDTCLPQDLPYAWCYIPILYFIAFFVGFIGNLLVIVLMAKKKGSKRLVDTFVLNLAV<br>ADLVFVCTLPLWAVSAAYDNKWYFGEGMCKFSSFIISVNRSSSILFLMGMSVERYLVVMKRVDSKSTGTKNCLIIACGCI<br>WAVSLLLGIPSMMYRKLSPSEDTTGELCEDEDSITFQVLSLALLILTFLLPLGVILFCYCSISFKLLNHISLGKGTKNSL<br>QIIFTIIGAFICCWLPFNAFKAFLFFSSTFQGADLSCWALMALRRGLTISACLAFMNSCVNPIIYALMDHHFRRQVLQQR<br>LRVCGAFQETRQTSVLSVSSATESSLMFASRNKPPSPASIRPEM |
| | *Chelonia mydas* (Green sea turtle) | Gene ID: 102942126<br>mRNA ID: XM_027834774<br>Protein ID: XP_027690575 | MTTEAWTVNYEASSAVYSDYNPLQDTCPPQDLPYAWVYIPILYFIAFFVGFMGNLLVIVLMAKKKGSKRLVDTFVLNLAV<br>ADLVFVCTLPLWAVSAAYDNKWYFGEGMCKFSSFIISVNRSSSILFLMGMSVERYLVVMKRVDSKSTGTKNCLIIACGCI<br>WAVSLLLGIPSMMYRKLSHSEDTTGELCEDEDSIAFQVLSLALLILTFLVPLGVILFCYCSISFKLLNHISLGKGTKNSL<br>QIIFTIIGAFICCWLPFNAFKAFLFFSSTFQGADLSCWALMALRRGLTISACLAFMNSCVNPIIYALMDHHFRRQVLQQR<br>LRVCGAFQETQQTSVLSVSSATESSLMFASRNKPPSPASIRPEM |
| | *Chelonoidis abingdonii* (Abingdon island giant tortoise) | Gene ID: 116832686<br>mRNA ID: XM_032793590<br>Protein ID: XP_032649481 | MKTEAWTGSHEASSAVYWDYNPLQDTCPPQDLPYACVYIPMLYFIAFFIGFIGNLLVIVLMAKKRGRKRLVDMFVLNLAV<br>ADLVFVLTLPFWAVSAAHGNKWYFGEGLCKFSSFIISVNRSSSILFLMGMSVERYLVVMKRVDSKSTGTKNCLIIVCGCI<br>WAVSLLLGIPSLMFRKLDTTGELCEDEDSVAFQVLSLALLILTFLLPLGVILFCYCSISFKLLNHISPGKGTKNSLQIIF<br>TIIGAFICCWLPFNAFKAFLFFSFMFQGANLSCRTLMALRRGLTISACLAFMNSCVNPIIYALMDRHFRRQLLPQWLRVF<br>WKVPQTRQTSGLSFSSTTESSLMFASRNKPSSPAFIRPEM |
| | *Chrysemys picta bellii* (western painted turtle) | Gene ID: 101950668<br>mRNA ID: XM_005313112<br>Protein ID: XP_005313169 | MTTEAWTVSYEASSAVYWDYNPLQDTCPPQDLPYACVYIPILYFITFFIGFIGNLLVIVLMAKKRGSKRLVDTFVLNLAV<br>ADLVFVLTLPLWAVSAAHGNKWYFGESLCKFSSFIISVNRSSSILFLMGMSVERYLVVMKRVDSKSTGTKNCLIIACGCI<br>WAVSLLLGIPSLMYRKLSLSEDTTGELCEDEDSVAFQVLSLALLILTFLLPLGVILFCYCSISFKLLNHIRLGKGTKNSL<br>QIIFTIIGAFICCWLPFNAFKAFLFFSSMFQGADLSCWALMALRRGLTISACLAFMNSCVNPIIYALMDRHFRHQVLPQR<br>LRVCGAFQETRQTSGLSFSTTTESSLMFASRNKPSSPASIRPEM |
| | *Crocodylus porosus* (Australian saltwater crocodile) | Gene ID: 109317483<br>mRNA ID: XM_019546190<br>Protein ID: XP_019401735 | MTTEAWTTGFEMSGDYEWGDLCQQQKLHFAQVYIPILYFFAFLIGFIGNLFVIVLMAMKKGNKRMVDNFVLNLAVADLVF<br>VCTLPLWALSAAHNNHWYFGESLCKVSSYIISVNRCSSILFLTGMGVERYLVVTKRVHSKSSMTKKCIFTTCGCIWAVAL<br>LLGIPSLLYRKLNASDKFCQDEDSSVYRKYNLALLVLTFLLPLGVILFCYCSICIKLLSHISMGKGTKNSLKIIFTIIST<br>FICSWLPFNAFKAFLILSSPSELNQEASVSCSAQTFIKWGLSLSTCLAFMNSCINPIIYAFMDHHFRRRVQLYCFHFFGT<br>SRKTSQGLVVSSSAAESSLVFASRNKPSSPICHQPSM |
| | *Crotalus tigris* (Tiger rattlesnake) | Gene ID: 120298940<br>mRNA ID: XM_039323193<br>Protein ID: XP_039179127 | MKISELVTDLAMTKNWNGDLELPTVDYYGSGVTSDAFCLREKLPFASIFISVIYFMIFFIGFFGNLFVVLVMVQKRGNQR<br>LVDNFVLNLAIADLVFVCTLPFWALTEAFDSWWFGEELCKFSSYAISVNRCSSILFLMGMSVERFLVMNKHWDSRSLGTK<br>RNSNITCGCIWVFSFLLGIPSLLYRKLIPVGDRFSCQDEEPSVIYIFTVLFLTFLLPLGVILFCYCSIFAMLQSHVKLGR<br>KTNHALKIIFVVIGAFICCWLPFNTFTAFITYGVVQNIDFSCEVLKALRRGLIISACLAFTNSCINPIIYAFMDQQFRQK<br>VLKNFLGLFHSKENVQSSSLFFSSTDSGLMFGSKKRFIPATTVPVSSGAGLCASKRQI |
| | *Dermochelys coriacea* (leatherback sea turtle) | Gene ID: 119846506<br>mRNA ID: XM_038380289<br>Protein ID: XP_038236217 | MTTETWTVNYEASSAVYSDYNPLXDTCPPQDLPYACVYIPILYFIAFFIGSLLVIVLMAKKKSSKMLVDTFVLNLAVADL<br>VLVYTLPHWAVSAAYDNKWYFGEGMCKFSRFIISVNRSSSILFLMGMSVERYLVVMKCVDSKSTGTKNXIIACGCIWAVS<br>LLLGIPSMMYRKLSPSEDTTGELCEDEDSIAFQVLSLALLILTFLLPLGVILFCYCSXSLKLLNHISLGKGTKNSLQIIF<br>TIISAFICCWLPFNAFKAFLFFSSTFQGADLSCWALMALRQGLTISACLAFMNSCINPIIYALMDHHFRRQVLQQQLLVC<br>DAFQEIRQTLVLSVSSTTESSLMFASQNKPSSPASIRLEM |
| | *Elgaria multicarinata webbii* | Gene ID: 134393744<br>mRNA ID: XM_063118834<br>Protein ID: XP_062974904 | MEAAQLETGSVITESWNSGSESSITDYYDSNITNDDTCILKEMPFAYIFIPTFYIITFLVGFFGNLLVILLMARKQGSKR<br>LVDTFVLNLAIADLVFVCTLPFWAVSEALSYWCFSEGLCKFSSYALSVNRYSSMLFLTGMSVERFLVISKHWDSKSIGTK<br>KSTGISCGCIWLVSLLLGIPSLVYRKLILVGKNKSLCQDEEPSPTFNLVILALTFLLPLGIILFCYCSISSKLRGHVSLG<br>KRRNNALKIIFAIIGAFICSWLPFNTIKAYFIFIITQDIGLSCQALMALKWALPISVCLAFTNSCINPVIYVFMDQHFQQ<br>QVWKNFPSFCRKRENMQSSALSFSSTESSLLFASRKKLKPALSTQIGDPNSRGTGL |
| | *Emydura macquarii macquarii* | Gene ID: 137142441<br>mRNA ID: XM_067531817<br>Protein ID: XP_067387918 | MTTEAWTASYDYPIGDNFCPPQDLAYTYIYIPVLYFIAFFIGFTGNLFVIVLMAKKRGSKRLVDTFVLNLAVADLVFVCT<br>LPLWAMSAAHGNKWHFGEGLCKFSSYIISVNRCSSILFLMGMSVERYLVVMKRVDSKSIGTKNCMIIACGCIWAVSLFLG<br>IPSLIHRKLSPMQDTTGELCEDEDSITFQVLSLVLLILTFLLPLGVILFCYCSISFKLLNHVSLRKGTKNSLHIIFTIIG<br>AFVGCWLPFNAFKAFLFFSSVFQEAVLSCSALMVLRWGLTISVCLAFLNSCINPIIYALMDRHFRRQVLPQRFWVCGASQ<br>ETRQNTVLSFSSATESSLMFASRNKPSSPTSIWPEM |
| | *Emys orbicularis* (European pond turtle) | Gene ID: 135876995<br>mRNA ID: XM_065402312<br>Protein ID: XP_065258384 | MTTEAWTVSYEASSAVYWDNPLQDTCPPQDLPYACVYIPILYFITFFIGFIGNLLVIVLMAKKRGSKRLVDTFVLNLAVA<br>DLVFVLTLPLWAVSAAHGNKWYFGEGLCKFSSFIISVNRSSSILFLMGMSVERYLVVMKRVDSKSTGTKNCLIIACGCIW<br>AVSLLLGIPSLMYRKLSLSEDTTGELCEDEDSVAFQVLSLALLILTFLLPLGVILFCYCSISFKLLNHIRLGKGTKNSLQ<br>IIFTIIGAFICCWLPFNAFKAFLFFSSMFQGADLSCWALMALRRGLTISACLAFMNSCVNPIIYALMDRHFRRQVLPQRL<br>RVCGAFQETRQTSGLSFSSTTESSLMFASRNKPSSPASIQPEM |
| | *Eretmochelys imbricata* (hawksbill sea turtle) | Gene ID: 144278240<br>mRNA ID: XM_077839474<br>Protein ID: XP_077695600 | MTTEPWTVNYEASSAVYSDYNPLQDTCLPQDLPYAWCYIPILYFIAFFVGFMGNLLVIVLMAKKKGSKRLVDTFVLNLAV<br>ADLVFVCTLPLWAVSAAYDNKWYFGEGMCKFSSFIISVNRSSSILFLMGMSVERYLVVMKRVDSKSTGTKNCLIIACGCI<br>WAVSLLLGIPSMMYRKLSPSEDTTGELCEDEDSITFQVLSLALLILTFLLPLGVILFCYCSISFKLLNHISLGKGTKNSL<br>QIIFTIIGAFICCWLPFNAFKAFLFFSSTFQGADLSCWALMALRQGLTISACLAFMNSCVNPIIYALMDHHFRRQVLQQW<br>LRVCGAFQETRQTSVLSVSSATESSLMFASRNKPPSPASIRPEM |
| | *Erythrolamprus reginae* (royal ground snake) | Gene ID: 139165650<br>mRNA ID: XM_070748862<br>Protein ID: XP_070604963 | MKIAELVTDLAITTNWSGDLESFIVDFSDLELTNDNFCLTEKLPFASIFISVIYFMIFFIGFFGNLFVILVMVQKKGNQK<br>LVDNFVLNLAVADLIFVCTLPFWALTEAFDSWWFGEELCKFSSYAISVNRCSSILFLMGMSVERFLVMNKHWVSRSFGTK<br>RNINVTCGCIWVFSFLLGIPSLLYRKLIPVGDRFACQDKNPSTVYIFTVLCLTFLLPLGVILFCYCSIFSMLKSHVNIGR<br>KANHALKIIFVIIGAFICCWLPFNTFTAVATYVKVQNPDFSCEALNALTRGLIISACLAFTNSCINPIIYAFMDQQFRQK<br>VMKNFLGRLCHPKENAQSSTLFLSSTESSLLFGSKKRFIPATTVPVSIETDLYTAKGQI |
| | *Eublepharis macularius* | Gene ID: 129330379<br>mRNA ID: XM_054980407<br>Protein ID: XP_054836382 | MGPAHLEFGSAMIANWSTSIDSFTADYSTVGSDDICLLEDLPYAHIFIPTMNFITFFTGFFGNLYVILLMARKQGSKRLV<br>DTFVLNLAVADLVFVCTLPFWAVSVALGYWCFGEVICKLSSYTISVNRCSSILFLMGMSVERFLVIMKHWDSRSMGTKKT<br>ISMSCGCIWMISLLVGIPSLVYRKLIPMKYTSKSLCQDETPSTTFSLVLLILTFLLPLGVILFCYCSIFSKLRGHVSQGK<br>RRNNALKIIFAIIGAFICFWLPFNTFSAFLIFAVSQNMYLSCPEQTALRWGLTTSACLAFTNSCINPIIYVFMDQHFRQQ<br>ALKYCTSFCNKGGTMQNSALAFSSTESSLLFANKKKFTSVTSNRLEDVKGTGTDVQTSARCLILV |
| | *Euleptes europaea* (tarantolino) | Gene ID: 130473284<br>mRNA ID: XM_056844807<br>Protein ID: XP_056700785 | MCFCQPEHSFGGSSIETGPAHLESGSAMAANWSIGFESFIEDYSDVGSNDSCLLQELPYAHIFMPTLNFITFFTGFFGNL<br>YVILLMARKQGSKRLVDTFVLNLAVADLVFVCTLPFWALSVALGYWCFGEVMCKLSSYTISVNRCSSILFLMGMSVERFL<br>VVTKHWDSRSMGTKKSIGVSCGCIWVVSLLLGIPSLVYRKLIPITDTSKSLCQDESSSPTFSLVLLIVTFLLPLGVILFC<br>YCSIFSKLRGHASRGKRRSNALKIIFAIIGAFICFWLPFNTFNTFLIFAVSQHMDLSCPAQMALRWGLTISACLAFTNSC<br>INPIIYVFMDQHFRQQALKNCPIFCSKGSKMQNSALSFSSTESSLLFANKKKFTSATSTQLEDMKGTGRGV |
| | *Gavialis gangeticus* (Gharial) | Gene ID: 109295412<br>mRNA ID: XM_019514272<br>Protein ID: XP_019369817 | MTTETWITGFEMSGDYELGDLCQQQKLPYAHIYIPILYFFVFLIGFIGNLFVIVLMAMKKGNKRMVDNFVLNLAVADLVF<br>VCTLPLWALSAAHNNHWYFGESLCKVSSYIISVNRCSSILFLTGMGVERYLVVMKRVRSKSSVTKKCIFTTCGCIWALSL<br>LLGIPSLLYRKLNASDKFCQDEDSSAYRKYSLALLILTFLLPLGVILFCYCSICIKLLSHISMGKGTKNSLKIIFTIIST<br>FICSWLPFNAFKAFLILSSPSELNQESSLSCSAQTAIKWGLSLSIYLAFMNSCINPIIYAFMDHHFRRRVQLCCLHFFCA |

| | | | |
|---|---|---|---|
| | | | SQKPTQGLVFFSSAAESSLMFASRNKPSSSICHQPSM |
| | *Gekko japonicus* | Gene ID: 107110959<br>mRNA ID: XM_015411818<br>Protein ID: XP_015267304 | MGPAHLESGSAVTANWSTDFESFTEHYYGVGSDDSCPPQELPYAHIFIPTLNFITFFTGFFGNLYVILLMARKQGSKRIV<br>DTFVLNLAVADLVFVCTLPFWALSVALGYWCFGEVICKLSSYTISVNRCSSILFLMGMSVERFLVVTKHWDSRSIGTKKS<br>TGMSCGCIWVVSLLLGIPSLVYRKLIPMKDTSKPLCQDEVPPTTFSLVLLILTFLLPLGVILFCYCSIFFKLRGHVSRGN<br>RRNNALKIIFAIIGAFICFWLPFNTFNAFLIFAVSQNIYPSCRAQMALRWGLTISACLAFTNSCINPIIYVFMDQHFRQQ<br>ALKNCPMFCSKGSKMQNSALSFSSTESSLLFANKKKTTSFTSTHRLEDMKGMGRSYKPPQG |
| | *Gopherus evgoodei* (Goodes thornscrub tortoise) | Gene ID: 115651592<br>mRNA ID: XM_030562660<br>Protein ID: XP_030418520 | MNTEAWTGSYEASSAVYWDYNLLQDTCPPQDLPYACVYIPMLYFIAFFIGFIGNLLVIVLMAKKRGRKRLVDMFVLNLAV<br>ADLVFVLTLPFWAVSAAHGNKWYFGEGLCKFSSFIISVNRSSSILFLMGMSVERYLVVMKHVDSKSTGTKNCLIIVCGCI<br>WAVSLLLGIPSLMFRKLDTTGELCEDEDSVAFQVLSLALLILTFLLPLGVILFCYCSISFKLLNHISLGKGTKNSLQIIF<br>TIIGAFICCWLPFNAFKAFLFFSSMFQVADLSCRTLMALRRGLTISACLAFMNSCVNPIIYALMDRHFRRQLLPQRLWVC<br>GTFQKTRQTSGLSFSSTTESRLMFASRNKPSSPAFIRPEM |
| | *Gopherus flavomarginatus* (Mexican gopher tortoise) | Gene ID: 127052534<br>mRNA ID: XM_050956360<br>Protein ID: XP_050812317 | MNTEAWTGSYEASSAVYWDYNPLQDTCPPQDLPYAYVYIPMLYFIAFFIGFIGNLLVIVLMAKKRGRKRLVDMFVLNLAV<br>ADLVFVLTLPFWAVSAAHGNKWYFGEGLCKFSSFIISVNRSSSILFLMGMSVERYLVVMKHVDSKSTGTKNCLIIVCGCI<br>WAVSLLLGIPSLMFRKLDTTGELCEDEDSVAFQVLSLALLILTFLLPLGVILFCYCSISFKLLNHISLGKGTKNSLQIIF<br>TIIGAFICCWLPFNAFKAFLFFSSMFQVADLSCRTLMALRRGLTISACLAFMNSCVNPIIYALMDRHFRRQVLPQRLWVC<br>GTFQKTRQTSGLSFSSTTESSLMFASRNKPSSPAFIRPEM |
| | *Heloderma suspectum* (Gila monster) | Gene ID: 146348355<br>mRNA ID: XM_084848307<br>Protein ID: XP_084704433 | MEAAQLETGSAITQNWSSGFESSTIDYYGSSITNYDICLLEELPFAYIFIPIFYVITFLVGFFGNLLVILLMARKQGSKR<br>LVDTFVLNLAIADLVFVCTLPFWAISEALSYWCFGEGLCKLSSYAISVNRYSSMLFLTGMSVERFLVISKHWDSRSIGTK<br>KSTGMSCGCIWLVSLLLGIPSLVYRKLNLVEDDKSLCQGEEPSATFSLVILGLTFLLPLGIILFCYCSIFFKLRGHVSLG<br>KRRNNALKIIFAIIGAFICSWLPFNTVKAFFIFIIAQDIDLSCQVLTALKWALPISACLAFTNSCINPVIYVFMDQHFRQ<br>QVWKNFPAFCSTRENMQSSALSFSSTESSLLFASRKKLTPAMSTRMGDPNSRGVGL |
| | *Hemicordylus capensis* (graceful crag lizard) | Gene ID: 128323140<br>mRNA ID: XM_053245865<br>Protein ID: XP_053101840 | MEPARLGTESVTTENGNSSFEPFTVDYYGFMDYDKCYPQELPYAYIVIPTLYFMTFLIGFFGNLFVILLMARKQGSKRLV<br>DNFVLNLAVADLVFVCTLPFWAVSNALGYWSFGEGLCKFCSYTISVNRCSSILFLMGMSVERFLVVTKHWDSRSIGIKKN<br>IIMSLGCVWVTSLLLGVPSVVYRKLSPVDGTSKYLCEGEPPNAFSLVLLIVTFLLPLGVILFCYCSIFSKLRGHVSLGKR<br>RNNALKIIFAIVGFVICSWLPFHTLKAYITVIKAQNLDLSCQALVAFRWGVTISACLGFTNSCINPIIYVFMDQQFRQQA<br>LKSIPVFHNTKEDLQSSALSFSSTESSVLFGNRKKLPSATSMQVGGTKGKGRDV |
| | *Heteronotia binoei* (common prickly gecko) | Gene ID: 132567611<br>mRNA ID: XM_060233290<br>Protein ID: XP_060089273 | MGAADLVSGSAMTTNWSTDYSGVGSDDICPLQELPYAHIFIPTLNFITFFTGFFGNLYVILLMARKQGSKRLVDTFVLNL<br>AVADLVFVCTLPFWALSVALGYWCFGEVICKLSSYTISVNRCSSILFLMGMSVERFLVVMKHWDSRSTGTKKSIGISCGC<br>IWVVSLLLGIPSLVYRKLIPMKDTSQPLCQDEVPSTTFSLVLLILTFLLPLGVILFCYCSIFFKLRGHISRGNRRNNALK<br>IIFAIIGAFICFWLPFNTFNTFLIFAVSQDLYLSCPVQTALRWGLAITACLAFTNSCINPIIYVFMDQHFRQQALKNCPT<br>FCSKGSKMQNSALSFSSTESSLLFANKKITSATSTHCLEDVKGTRRGYKPPQDFIVDNM |
| | *Lacerta agilis* (Sand lizard) | Gene ID: 117048547<br>mRNA ID: XM_033152505<br>Protein ID: XP_033008396 | MEPAQMVTGSSMTETQSGDFESYTIGNYGDLVNYDICHLQELPFAYIFVPTIYYIAFLIGFFGNLFVILLMARKEGRKRL<br>VDTFVLNLAVADLVFVCTLPFWAVSKALGYWCFGEGLCKLSSYAITVNRCSSILFLMGMSVERFLVMTKQWDSRTFSTKK<br>SIAMSCGCIWVSSLLLGMPSLVYRKLNLVDDKYLCQDKPSPTFSLIMLILTFLLPLGVILFCYCSIFFKLQSHVHLGKRR<br>NNALKIIFAIVGAFICSWLPFNVLKAFFVFTMAQDIHLRCSAMMALTWGLTISACLAFANSCVNPIIYVFMDQQFRQQAL<br>KHFPAFCSKRENVQSSALSFSSTDTSLLFASRKKPTPATSARVGDPESKGVGL |
| | *Lepidochelys kempii* (Atlantic ridley) | Gene ID: 140901199<br>mRNA ID: XM_073319604<br>Protein ID: XP_073175705 | MTTEPWTVNYEASSAVYSDYNPLQNTCLPQDLPYAWCYIPILYFIAFFVGFTGNLLVIVLMAKKKGSKRLVDTFVLNLAV<br>ADLVFVCTLPLWAVSAAYDNKWYFGEGMCKFSSFIISVNRSSSILFLMGMSVERYLVVMKRVDSKSTGTKNCLIIACGCI<br>WAVSLLLGIPSMMYRKLSPSEDTTGELCEDEDSITFQVLSLALLILTFLLPLGVILFCYCSISFKLLNHISLGKGTKNSL<br>QIIFTIIGAFICCWLPFNAFKAFLFFSSTFQGADLSCWALMALRRGLTISACLAFMNSCVNPVIYALMDHHFRRQVLQQR<br>LRVCGAFQETRQTSVLSVSSATESSLMFASRNKPPSPASIRLEM |
| | *Lepidochelys olivacea* (Pacific ridley) | Gene ID: 311510870<br>mRNA ID: XM_084610923<br>Protein ID: XP_084467049 | MTTEPWTVNYEASSAVYSDYNPLQNTCLPQDLPYAWCYIPILYFIAFFVGFTGNLLVIVLMAKKKGSKRLVDTFVLNLAV<br>ADLVFVCTLPLWAVSAAYDNKWYFGEGMCKFSSFIISVNRSSSILFLMGMSVERYLVVMKRVDSKSTGTKNCLIIACGCI<br>WAVSLLLGIPSMMYRKLSPSEDTTGELCEDEDSITFQVLSLALLILTFLLPLGVILFCYCSISFKLLNHISLGKGTKNSL<br>QIIFTIIGAFICCWLPFNAFKAFLFFSSTFQGADLSCWALMALRRGLTISACLAFMNSCVNPIIYALMDHHFRRQVLQQR<br>LRVCGAFQETRQTSVLSVSSATESSLMFASRNKPPSPASIRLEM |
| | *Malaclemys terrapin pileata* | Gene ID: 128835391<br>mRNA ID: XM_054024842<br>Protein ID: XP_053880817 | MTTEAWTVSYEASSAVYWDYNPLQDTCPPQDLPYACVYIPILYFITFFIGFIGNLLVIVLMAKKRGSKRLVDTFVLNLAI<br>ADLVFVLTLPLWAVSAAHGNKWYFGESLCKFSSFIISVNRSSSILFLMGMSVERYLVVMKRVDSKSTGTKNCLIIACGCI<br>WAVSLLLGIPSLMYRKLSLSEDTTGELCEDEDSVTFQVLSLALLILTFLLPLGVILFCYCSISFKLLNHIRLGKGTKNSL<br>QIIFTIIGAFICCWLPFNAFKAFLFFSSMFQGADLSCWALMALRRGLTISACLAFMNSCVNPIIYALMDHHFRRQVLPQR<br>LRVCGAFQETRQTSGLSFSSTTESSLMFASRNKPSSPASIRPEM |
| | *Mauremys mutica* (yellowpond turtle) | Gene ID: 123368675<br>mRNA ID: XM_045013675<br>Protein ID: XP_044869610 | MTTEAWTGSYEASSDVYWDYNPLQDTCPPQDLPYACVYIPMLYFIVFFIGFIGNLLVIVLMAKKRGRKRLVDTFVLNLAV<br>ADLVFVLTLPFWAVSAAHGNKWYFGEGLCKLSSFIISVNRSSSILFLMGMSVERYLVVMKRVDSKSTGTKNCLIIACGCI<br>WAVSLLLGIPSLMYRKLSPSEDSTGELCEDEDSVAFQVLSLALLILTFLLPLGVILFCYCSISFKLLNHISLGKGTKNSL<br>QIIFTIICTFICCWLPFNAFKAFLFFSSMFQGADLSCWALMALRRGLTISACLAFMNSCVNPIIYALMDRHFRRQVLPQG<br>LQVCGTFQETRQTSALSFSSTTESSLMFASRNKPSSPASIRPEM |
| | *Mauremys reevesii* (Reeves's turtle) | Gene ID: 120405280<br>mRNA ID: XM_039538780<br>Protein ID: XP_039394714 | MTTEAWTGSYEASSDVYWDYNPLQDTCPPQDLPYACVYIPMLYFIAFFIGFIGNLLVIVLMAKKRGRKRLVDTFVLNLAV<br>ADLVFVLTLPFWAVSAAHGNKWYFGEGLCKLSSFIISVNRSSSILFLMGMSVERYLVVMKRVDSKSTGTKNCLIIACGCI<br>WAVSLLLGIPSLMYRKLSPSEDTTGELCEDEDSVTFQVLSLALLILTFLLPLVVILFCYCSISFKLLNHISLGKGTKNSL<br>QIIFTIICAFICCWLPFNAFKAFLFFSSMFQGADLSCWALMALRRGLTISACLAFMNSCVNPIIYALMDRHFRRQVLPQG<br>LQVCGTFQETRQTSALSFSSTTESSLMFASRNKPSSPASIRPEM |
| | *Natator depressus* (flatback sea turtle) | Gene ID: 141975719<br>mRNA ID: XM_074936311<br>Protein ID: XP_074792412 | MTTEAWTVNYEASSAVYSDYNPLQDTCPPQDLPYAWVYIPILYFIAFFVGFIGNLLVIVLMAKKKGSKRLVDTFVLNLAV<br>ADLVFVCTLPLWAVSAAYDNKWYFGEGMCKFSSFIISVNRSSSILFLMGMSVERYLVVMKRVDSKSTGTKNCLIIACGCI<br>WAVSLLLGIPSMMYRKLRPSEDTTGELCEDEDSITFQVLSLALLILTFLLPLGVILFCYCSISFKLLNHISLGKGTKSSL<br>QIIFTIIGAFICCWLPFNAFKAFLFFSSTFQGADLSCWALMALRRGLTISACLAFMNSCVNPIIYALMDHHFRCQVLQQR<br>LRVCGAFQETRQTSVLSVSSATESSLMFASRNKPPSPASIRPEM |
| | *Natrix helvetica* | Gene ID: 309389969<br>mRNA ID: XM_081948937<br>Protein ID: XP_081805063 | MKISELVTDLAMTKNWSRDLESATVDFFDLGFTNDDFCFMEKLPFASVFVSVIYFMIFFIGFFGNLFVILVMVLKRGNQR<br>LVDNFILNLAIADLVFVCTLPFWAMAEAFDSWQFGEVFCKFSSYAIFVNRCSSILFLMGMSVERFLVMNKHWDSRFLGTK<br>RNTSITCSCIWVFSFLLGIPSLLYRKVIPVGDRFACQDEGSSIIYTFTVLCLTFLLPLGVILFCYCSIFTMLQSHVNVGR<br>KTNHALKIIFVIIGAFICCWLPFNIFTAVITYGVVQNIEFSCEALKVLRRGVIISACLAFTNSCINPIIYAFMDQQFRQK<br>VLKSLLGCLCPPKENAQSSPLFFSSTESSVLFGSQKRFIPAITVPVSSRADLYASKREI |
| | *Notechis* | Gene ID: 113419021 | MKIAELVTDLAMTKNWSRDLESSTEDFFDLGLTDDDFCLIEKLPFASVFISVIYFMIFFIGFFGNLFVVLVMVQKRGNQR |

| | | | |
|---|---|---|---|
| | *scutatus* | mRNA ID: XM_026678216<br>Protein ID: XP_026534001 | LVDNFVLNLAIADLVFVCTLPFWALTEAFDSWWFGEELCKFSSYAISVNRCSSILFLMGMSVERFLVMNKHWDSRSLGTK<br>RNTNITCGCIWVFSFLLGIPSLLYRKLISVGDRFACQDVDPSVIYTFTVLCLTFLLPLGVILFCYCSIFSMLQSHVNVGR<br>KTNHALKIIFVIIGAFICCWLPFNTFTAVITYGMVQNIDFPCEAVKALRRGVIISACLAFTNSCINPIIYAFMDQQFRQK<br>NQAFCLAAKRGSYQPQLFQLAAGLIYMPLKDIYSCYC |
| | *Pantherophis guttatus* | Gene ID: 117664398<br>mRNA ID: XM_034415296<br>Protein ID: XP_034271187 | MKFAELVTDLAMTKNWSRDLESSTVDVFGLEFTNDDFCFMEKLPFASIFISVIYFMIFFIGFFGNLFVILVMVQKRGNQR<br>LVDKFVLNLAIADLVFVCTLPFWALTEASDSWWFGEELCKFSSYAISVNRCSSILFLMGMSVERFLVMNKHWDSRSLGTK<br>RNTNITCSCIWVFSFLLGIPSLLYRKLIPVGDRFACQDEDPSVIYIFTVLCLTFLLPLGVILFCYCSIFSMLKSHVNPGR<br>KTNHALKIIFVIIGAFICCWLPFNTFTAVITYYGVIQNIEFSCDALKALRRGVKISACLAFTNSCINPIIYAFMDQQFRQ<br>KVLKNFLGLCRPKENAQSSALFFSSTESSLLFGSKKRFIPATTVPINSGTDLYASKRQI |
| | *Paroedura picta* | Gene ID: 143835867<br>mRNA ID: XM_077334241<br>Protein ID: XP_077190356 | MTANGSTGLESFTEDYSGFDSDDNCLLHELPYAHIFIPTLYFITFFTGFFGNLYVILLMARKQGSKRLVDTFVLNLAVAD<br>LVFVCTLPFWALSVALGYWCFGEVICKLSSYTISVNRCSSILFLMGMSVERFLVVTKHWDSRSLGTKRSAGMSCGCIWVV<br>SLLLGIPSLLYRKLIPMIDTSKPLCQDEEPPTTFSLVLLILTFLLPLGVILFCYCSIFFKLRGHISRGNRRNNALKIIFA<br>IIGTFICFWLPFNVFNTFLIFVETQHVYLSCPAQMALRWGLTISACLAFTNSCINPIIYIFMDQHFQQQALKNCPMFCSK<br>GSKMQNSALSFSSTESNLLFANKKISSATSPRHLEDMKGMGRGYKPPQXLIVDNM |
| | *Pituophis catenifer annectens* | Gene ID: 139314145<br>mRNA ID: XM_070939684<br>Protein ID: XP_070795785 | MKFVELVTDLAMTKNWSRDLESSTVDVFGLEFTNDDFCFMEKLPFASIFISVIYFMIFFIGFFGNLFVILVMVQKRGNQR<br>LVDKFVLNLAIADLVFVCTLPFWALTEASDSWWFGEELCKFSSYAISVNRCSSILFLMGMSVERFLVMNKHWDSRSLGTK<br>RNTNITCSCIWVFSFLLGIPSLLYRKLIPVGDRFACQDEDPSVIYIFTVLCLTFLLPLGVILFCYCSIFSMLKSHVNPGR<br>KTNHALKIIFVIIGAFICCWLPFNTFTAVITYYGVIQNIEFSCDALKALRRGVIISACLAFTNSCINPIIYAFMDQQFRQ<br>KVLKNFLGLCRPKENAQSSALFFSSTESSLLFGSKKRFIPATTVPVNSGTDLYASKRQI |
| | *Podarcis bocagei* | Gene ID: 308586978<br>mRNA ID: XM_081194215<br>Protein ID: XP_081050341 | MEPAQMVTGSSMTETWSGDYESSAIDNYGDLVNYDICHLQELPFAYIFIPTVYYITFLIGFFGNLFVILLMARKEGRKRL<br>VDTFVLNLAVADLVFVCTLPFWAVSKALGYWCFGEGLCKLSSYAITVNRCSSILFLMGMSVERFLVMTKQWDSRTISTKK<br>SIAVSCGCIWVFSLLLGMPSLFYRKLNPVDDKYLCHDEPSPTFSLIMLILTFLLPLGVILFCYCSIFSKLQSHVHLGKRR<br>NNALKIIFAIVGAFICSWLPFNVLKVFFVFTMAQDIHPRCSAMMALTWGLTISACLAFANSCVNPIIYVFMDQQFRQQAL<br>KHFPAFCSKRGNVQSSALSFSSTDTSLLFASRKKPTPATSARVGDPESKGVGL |
| | *Podarcis cretensis* | Gene ID: 307218280<br>mRNA ID: XM_079980495<br>Protein ID: XP_079836621 | MEPAQMVTGSSMTETWSGDFESSAIDNYGDLVNYGICHLQELPFAYIFIPTVYYITFLIGFFGNLFVILLMARKEGRKRL<br>VDTFVLNLAVADLVFVCTLPFWAVSKALGYWCFGEGLCKLSSYAITVNRCSSILFLMGMSVERFLVMTKQWDSRTISTKK<br>SMAMSCGCIWVFSLLLGMPSLFYRKLNPVDDKYLCHDEPSPTFSLIMLILTFLLPLGVILFCYCSIFFKLQSHVHLGKRR<br>NNALKIIFAIVGAFICSWLPFNVLKVFFVFTMAQDIHPRCSAMMALTWGLTISACLAFANSCVNPIIYVFMDQQFRQQAL<br>KHFPAFCSKRGNVQSSALSFSSTDTSLLFASRKKPTPATSARAGDPESKGVGL |
| | *Podarcis erhardii* | Gene ID: 309344341<br>mRNA ID: XM_081866643<br>Protein ID: XP_081722769 | MEPAQMVTGSSMTETWSGDFESSAIDNYGDLVNYDICHLQELPFAYIFIPTIYYITFLVGFFGNLFVILLMARKEGRKRL<br>VDTFVLNLAVADLVFVCTLPFWAVSKALGYWCFGEGLCKLSSYAITVNRCSSILFLMGMSVERFLVMTKQWDSRTISTKK<br>SIAMSCGCIWVFSLLLGMPSLFYRKLNPVDDKYLCHDEPSPTFSLIMLILTFLLPLGVILFCYCSIFSKLQSHVHLGKRR<br>NNALKIIFAIVGAFICSWLPFNVLKVFFVFTMAQDIHPRCSAMMALTWGLTISACLAFANSCVNPIIYVFMDQQFRQQAL<br>KHFPALCSKRGNVQSSALSFSSTDTSLLFASRKKPTPATSARAGDPESKGVGL |
| | *Podarcis filfolensis* (Filfola wall lizard) | Gene ID: 310764277<br>mRNA ID: XM_083617857<br>Protein ID: XP_083473983 | MEPAQMVTGSSMTETWSGDFESSTIDTYGDLVNYDICHLQELPFAYIFIPTVYYITFLIGFFGNLFVILLMARKEGRKRL<br>VDTFVLNLAVADLVFVCTLPFWAVSKTLGYWCFGEGLCKLSSYAITVNRCSSILFLMGMSVERFLVMTKQWDSRTISTRK<br>SIAMSCGCIWVFSLLLGMPSLFYRKLNPVDDKYLCHDMPSPTFSLIMLILTFLLPLGVILFCYCSIFSKLRSHVHLGKRR<br>NNALKIIFAIVGAFICSWLPFNVLKAFFVFTMAQDIHLRCSAMMALTWGLTISACLAFANSCVNPIIYVFMDQQFRQQAL<br>KHFPAFCSKRGNVQSSALSFSSTDTSLLFASRKKPTPATSAQAGDPDSKGVGL |
| | *Podarcis gaigeae* (Skyros wall lizard) | Gene ID: 310847417<br>mRNA ID: XM_083745227<br>Protein ID: XP_083601353 | MEPAQMVTGSSMTETWSGDFESSAIDNYGDLVNYDICHLQELPFAYIFIPTVYYITFLIGFFGNLFVILLMARKEGRKRL<br>VDTFVLNLAVADLVFVCTLPFWAVSNALGYWCFGEGLCKLSSYAITVNRCSSILFLMGMSVERFLVMTKQWDSRTISTKK<br>SIAMSCGCIWVFSLLLGMPSLFYRKLNPVDDKYLCHDEPSPTFSLIMLILTFLLPLGVILFCYCSIFSKLRSHVHLGKRR<br>NNALKIIFAIVGAFICSWLPFNVLKAFFVFTMAQDIHLRCSAMMALTWGLTISACLAFANSCVNPIIYVFMDQQFRQQAL<br>KHFPAFCSKRGNVQSSALSFSSTDTSLLFASRKKPTPATSARAGDPQSKGVGL |
| | *Podarcis liolepis* (Columbretes wall lizard) | Gene ID: 310709760<br>mRNA ID: XM_083509795<br>Protein ID: XP_083365921 | MEPAQMVTGSSMTETWSGDFESSAIDNYGDRVNYDICHLQELPFAYIFIPTVYYITFLIGFFGNLFVILLMARKEGRKRL<br>VDTFVLNLAVADLVFVCTLPFWAVSKALGYWCFGEGLCKLSSYAITVNRCSSILFLMGMSVERFLVMTKQWDSRTISTKK<br>SIAVSCGCIWVFSLLLGMPSLFYRKLNLVDDKYLCHDEPSPTFSLIMLILTFLLPLGVILFCYCSIFSKLQSHVHLGKRR<br>NNALKIIFAIVGAFICSWLPFNVLKVFFVFTMAQDIHPRCSAMMALTWGLTISACLAFANSCVNPIIYVFMDQQFRQQAL<br>KHFPAFCSKRGNVQSSALSFSSTDTSLLFASRKKPTPATSARVGDPESKGVGL |
| | *Podarcis melisellensis* | Gene ID: 309372123<br>mRNA ID: XM_081915408<br>Protein ID: XP_081771534 | MEPAQMVTGSSMTETWSGDFESSTIDNYGDLVNYDICHLQELPFAYIFIPTIYYITFLIGFFGNLFVILLMARKEGRKRL<br>VDTFVLNLAVADLVFVCTLPFWAVSKALGYWCFGEGLCKLSSYAITVNRCSSILFLMGMSVERFLVMTKQWDSRTISTKK<br>SIAMSCGCIWVFSLLLGMPSLFYRKLNPVDDKYLCHDEPSPTFSLIMLILTFLLPLGVILFCYCSIFSKLRSHVHLGKRR<br>NNALKIIFAIVGAFICSWLPFNVLKAFFVFTMAQDIHLRCSAMMALTWGLTISACLAFANSCVNPIIYVFMDQQFRQQAL<br>KHFPAFCSKRGNVQSSALSFSSTDTSLLFASRKKPAPATSARAGDPETKGVGL |
| | *Podarcis muralis* (Common wall lizard) | Gene ID: 114599189<br>mRNA ID: XM_028733912<br>Protein ID: XP_028589745 | MEPAQMVTGSSMTETWSGDFESSAIDNYGDLVNYDICPLQELPFAYIFIPTVYYITFLIGFFGNLFVILLMARKEGRKRL<br>VDTFVLNLAVADLVFVCTLPFWAVSKALGYWCFGEGLCKLSSYAITVNRCSSILFLMGMSVERFLVMTKQWDSRTISTKK<br>SIAVSCGCIWVFSLLLGMPSLFYRKLNPVDDKYLCHDMPSPTFSLIMLFLTFLLPLGVILFCYCSIFSKLRSHVHLGKRR<br>NNALKIIFAIVGAFICSWLPFNVLKPFFVFTMAQDIHLSCSKMMALTWGLTISACLAFANSCVNPIIYVFMDQQFRQQAL<br>KHFPAFCSKRGNVQSSALSFSSTDTSLLFASRKKPTPATSARVGDPESKGVGL |
| | *Podarcis pityusensis* | Gene ID: 308479727<br>mRNA ID: XM_081010553<br>Protein ID: XP_080866679 | MEPAQMVTGSSMTETWSGDFESSAIDNYGDLVNYDICHLQELPFAYIFIPTVYYITFLIGFFGNLFVILLMARKEGRKRL<br>VDTFVLNLAVADLVFVCTLPFWAVSKALGYWCFGEGLCKLSSYAITVNRCSSILFLMGMSVERFLVMTKQWDSRTISTKK<br>SIAMSCGCIWVFSLLLGMPSLFYRKLNPVDDKYLRHDEPSPTFSLIMLILTFLLPLGVILFCYCSIFSKLQSHVHLGKRR<br>NNALKIIFAIVGAFIFSWLPFNVLKVFFVFTMAQDIHPRCSAMMALTWGLTISACLAFANSCVNPIIYVFMDQQFRQQAL<br>KHFPAFCSKRGNVQSSALSFSSTDTSLLFASRKKPTPATSARAGDPESKGVGL |
| | *Podarcis raffonei* (Aeolian wall lizard) | Gene ID: 128415969<br>mRNA ID: XM_053392885<br>Protein ID: XP_053248860 | MEPAQMVTGSSMTETWSGDFESSAIDNYGDLVNYGICHLQELPFAYIFIPTVYYITFLIGFFGNLFVILLMARKEGRKRL<br>VDTFVLNLAVADLVFVCTLPFWAVSKALGYWCFGEGLCKLSSYAITVNRCSSILFLMGMSVERFLVMTKQWDSRTISTRK<br>SIAVSCGCIWVFSLLLGMPSLFYRKLNPVDDKYLCHDMPSPTFSLIMLILTFLLPLGVILFCYCSIFSKLRSHVHLGKRR<br>NNALKIIFAIVGAFICSWLPFNVLKAFFVFTMAQDIHLRCSAMMALTWGLTISACLAFANSCVNPIIYVFMDQQFRQQAL<br>KHFPAFCSKRGNVQSSALSFSSTDTSLLFASRKKPTPATSARAGDPESKGVGL |
| | *Podarcis tiliguerta* | Gene ID: 310679915<br>mRNA ID: XM_083445863<br>Protein ID: XP_083301989 | MEPAQMVTGSSMTETWSGDYESSAIDNYGDLVNYDTCHLQELPFAYIFIPTVYYITFLIGFFGNLFVILLMARKEGRKRL<br>VDTFVLNLAVADLVFVCTLPFWAVSKALGYWCFGEGLCKLSSYAITVNRCSSILFLMGMSVERFLVMTKQWDSRTISTKK<br>SIAMSCGCIWVFSLLLGMPSLFYRKLNPVDDKYLCHDEPSPTFSLIMLILTFLLPLGVILFCYCSIFSKLRSHVHLGKRR<br>NNALKIIFAIVGAFICSWLPFNVLKAFFVFTMAQDIHLRCSAMMALTWGLTISACLAFANSCVNPIIYVFMDQQFRQQAL<br>KHFPAFCSKRENVQSSALSFSSTDTSLLFASRKKPTPATSARVGDPESNGVGL |

| | | | |
|---|---|---|---|
| | *Podarcis vaucheri* | Gene ID: 309534754<br>mRNA ID: XM_082227091<br>Protein ID: XP_082083217 | MEPAQMVTGSSMTETWSGDFESSAIDNYGDLVDYGICPLQELPFAYIFIPTVYYITFLIGFFGNLFVILLMARKEGRKRL<br>VDTFVLNLAVADLVFVCTLPFWAVSKALGYWCFGEGLCKLSSYTITVNRCSSILFLMGMSVERFLVMTKQWDSRTISTKK<br>SIAVSCGCIWVFSLLLGMPSLFYRKLNPVDDKYLCHDEPSPTFSLIMLILTFLLPLGVILFCYCSIFSKLQSHVHLGKRR<br>NNALKIILAIVGSFIFSWLPFNVLKVFFVFTMAQDIHPHCSAMMALTWGLTISACLAFANSCVNPIIYVFMDQQFRQQAL<br>KHFPAFCSKRGNVQSSALSFSSTDTSLLFASRKKPTPATSARAGDPESKGVGL |
| | *Pogona vitticeps* (central bearded dragon) | Gene ID: 110085269<br>mRNA ID: XM_020805267<br>Protein ID: XP_020660926 | MESVQLAVGSAMNGNWSSRFETYSMDYYGSGSMNDDTCLLEELPFAYIFIPSLYFTIFLIGFLGNLFVILLMAQKRGSTR<br>LLDTFVLNLAIADLVFVCTLPFWAVSEVLSYWCFGEVLCKLSSYAISVNRCSSILFLVGMSVERFLVMVKHWDSRSIGTK<br>KNIAMSCGCIWVISLLLGIPPLTYRKVILAEDNKSLCLDEKPSVIFILVMFFLTFLLPLGVILFCYCSIFSKLRSHISLG<br>RRRNNAVKIIFATIAAFVCSWLPYNTFKIFVNFVVYWSIDLSCHMLMALSWGLTISVCLAFTNSCINPIIYIFMDQHFRQ<br>QLLKDFLGFCRTRENMQSSAPSFSSTESSLLFVNRKKLTPATSTLVGDPNGRGVGL |
| | *Protobothrops mucrosquamatus* (brown spotted pit viper) | Gene ID: 114922686<br>mRNA ID: XM_029284468<br>Protein ID: XP_029140301 | MKISELVTDLAMTKNWTGDLELDYYGSGVTSDAFCLREKLPFASIFISVIYFMIFFIGFFGNLFVVLVMVQKRGNQRLVD<br>NFVLNLAIADLVFVCTLPFWALTEAFDSWWFGEELCKFSSYAISVNRCSSILFLMGMSVERFLVMNKHWDSRSLGTKRNS<br>NITCGCIWVFSFLLGIPSLLYRKLIPVGDRFSCQDEEPSVVYIFTVLFLTFLLPLGVILFCYCSIFSMLQSHVKLGRKTN<br>HALKIIFVIIGAFICCWLPFNTFTAFITYGVVQNIDFSCEVLKALRRGLIISACLAFTNSCINPIIYAFMDQQFRQKVLK<br>NFLGLFHPKENVQSSSLFFSSTDSSLMFGSKKRFIPATTVPVSSGAGLCASKRQI |
| | *Pseudonaja textilis* | Gene ID: 113446985<br>mRNA ID: XM_026716919<br>Protein ID: XP_026572704 | MKIAELVTDLAMTKNWSGDLESSTVDFFGLGLTDDDFCLIEKVPFASVFISVIYFMIFFIGFFGNLFVVLVMVQKRGNQR<br>LVDNFVLNLAIADLVFVCTLPFWALTEAFDSWWFGEELCKFSSYAISVNRCSSILFLMGMSVERFLVMNKHWDSRSLGTK<br>RNTSITCGCIWVFSFLLGIPSLLHRKLISVGDKFACQDVDPSIIYTFTVLCLTFLLPLGVILFCYCSIFSMLQSHVNVGR<br>RTNHALKIIFVIIGAFICCWLPFNTFTAVITYGMVQNIDFPCEAVKALRRGLIISACLAFTNSCINPIIYAFMDQQFRQK<br>NQAFCLEAKRGSYQPQLFQLAAGLIYMPLKDIYSCCC |
| | *Python bivittatus* (Burmese python) | Gene ID: 112541279<br>mRNA ID: XM_025170036<br>Protein ID: XP_025025804 | MKIAELVTDSAMTKNWSRDFVDYYNSGFNSDDFCLMEKLPFASIFISVIYFMIFFIGFFGNFFVVLVMVQKRGNQRLVDN<br>FILNLAIADLVFVCTLPFWALSEAFGTWWFGEELCKFSSYAISVNRCSSILFLTGMSVERFLVMNKHWDSRSFGTKRNTN<br>MTCSCIWVLSLLLGMPSLVYRKLILVGDKFSCQDEEPSVIYTFAVPFLTFLLSLGVILFCYCSIFSMLRGHVNLGRRRNH<br>ALKIIFVIIGAFICCWLPFNIFTAFITYSMVQNVNLSCEALKSLRRGLTISACLAFTNSCINPIIYVFMDQQFRQKVLKN<br>FLGLCHPRENAQSSAMSFSSTESSLLFASRKKCIPATTVSSGAGFCASKREI |
| | *Rhineura floridana* | Gene ID: 133386853<br>mRNA ID: XM_061630622<br>Protein ID: XP_061486606 | MEPIQMVTENWSRNFESSTTDYYDLSSVNDVCLPQELPFAYIFIPILYFITFLTGFFGNLFVILLMARKKGTKRLVDTFV<br>LNLAVADLVFVCTLPFWALSEALSYWCFGEGLCKFSSYVISVNRCSSILFLMGMSVERFLVMTKHWDSRTIRTKKNIGMS<br>CGCIWVVSLLLGIPSLVYRKLIPVDDTKSLCQDEEPSTTFSMIMLVFTFLFPLGVILFCYCSIFAKLQGHVSRSKRRNNA<br>LKIIFAIIGAFIFSWLPFNMFKAFFILTRAQDISLSCPALMALIWGLTISASLAFTNSCVNPIIYVFMDQQFRQQALKSF<br>TTFCSTRENVQSSATSFSSTESSLLFASRKKLTPATSVQVGDPKCMGVGL |
| | *Sceloporus undulatus* (fence lizard) | Gene ID: 121927517<br>mRNA ID: XM_042461183<br>XP_042317117 | MEAAQLATKSAITEKRTNSFEAYTIDYYGSDSMNYDICLPEELPFAYIFIPILYFLIFLIGFFGNLFVILLMARKQGSKR<br>LVDTFILNLAIADLVFVFTLPFWALSVALGYWCFGEGLCKLSSYTISVNRCSSILFLAGMSVERFLVMTKRWDARSIGTR<br>KNTIMSCGCIWVFSLLLGIPSLVYRKLDLVGDNNSSCQGEPSAAFSSVMLILTFLLPLGVILFCYCSIFFKLWNHASLGK<br>RRNNAFKIIFAIIGAFVCSWLPFNIFKIFLVFTLAQSIDLSCQALRALIWGLTITACLAFINSCINPIIYAFMDQHFQQQ<br>VLMFCKARGNAQTSTPSFSFTESSALFGSRKKLPPAMSAQI |
| | *Sphaerodactylus townsendi* | Gene ID: 125433045<br>mRNA ID: XM_048497342<br>Protein ID: XP_048353299 | MGPAHLVSESAMTANWSTGFEPFTEDYSGAGSDDICQLQELPYARIFIPILNFMTFFIGFVGNLYVILLMARKQGRKRLV<br>DTFILNLAVADLVFVCTLPFWALSVALGYWCFGEVICKLSSYTISVNRCSSILFLMGMSVERFLVVTKHWDSRFMGTKVS<br>IGMSCGCIWLFSLLLGIPSLVYRKLIPMEDTSKSLCQDEVPFTTYSIVLLIITFLLPLGVILFSYCSIFTKLRGHASRGK<br>RRNNALKIIFAVIGAFICFWLPFNTFNAFLIFAVSENMDLSCPAQMVLRWGLTISACLAFTNSCINPIIYVFMDQHFQQQ<br>ALKNCPIFCSKGGKMQTSALSSSTESSLLFANKKKLTSATSTRLEDMKGTGSGL |
| | *Terrapene triunguis* (Three-toed box turtle) | Gene ID: 112121048<br>mRNA ID: XM_024220160<br>Protein ID: XP_024075928 | MTTEAWTVSYEASSAVYWDYNPLQDTCPPQDLPYACVYIPILYFITFFIGFIGNLLVIVLMAKKRGSKRLVDTFVLNLAV<br>ADLVFVLTLPLWAVSAAHGNKWYFGEGLCKFSSFIISVNRSSSILFLMGMSVERYLVVMKRVDSKSTGTKNCLIIACGCI<br>WAVSLLLGIPSLMYRKLSLSEDTTGELCEDEDSVAFQVLSLALLILTFLLPMGVILFCYCSISFKLLNHIRLGKGTKNSL<br>QIIFTIIGAFICCWLPFNAFKAFLFFSSMFQGADLSCWALMALRRGLTISACLAFMNSCVNPIIYALMDRHFRRQVLPQR<br>LRVCGAFQETRQTSGLSFSSTTESSLMFASRNKPSSPASIRPEM |
| | *Thamnophis elegans* | Gene ID: 116509695<br>mRNA ID: XM_032218956<br>Protein ID: XP_032074847 | MKISELVTDLAMTQNWSRDLESSTVDFFDLGFTNDDFCFMELPFASVFISVIYFMISSLASLGTCCVLVIVLKRGNQRLV<br>DNFILNLAIADLVFVCTLPFWAVTEAFXFLQFGEEFLQNXQLCHFCERCSSILFLMGMSVERFLVMNKHWDSRFLGTRXN<br>TSITCSCIWVFSFLLGIPSLLYRKADPSGXDRFACQDEGSSIIYTFTVLCLTFLLPLGVILFCYCSIFTMLQSHVNVGRK<br>XSHALKIIFVIIGAFICCWLPFNTFTAVITYGVVQNIDFSCEALKVLRWGVIISACLAFTNSCINPIIYXFMDQQFRQKV<br>LKSVLGRLCPAKENAQSSTLFFSSTESSVLFGSKKRFIPATIVPVSSGADLHASKDRYSCYC |
| | *Thamnophis sirtalis* | Gene ID: 106549237<br>mRNA ID: XM_014066829<br>Protein ID: XP_013922304 | MKISELVTDLAMTKNWSRDLESSTVDFFDLGFTNDDFCFMEKLPFASVFISVIYFMIFFIGFFGNLFVILVIVLKRGNQR<br>LVDNFILNLAIADLIFVCTLPFWAVTEAFDSWQFGEEFCKISSYAIFVNRCSSILFLMGMSVERFLVMNKHWDSRFLGTK<br>RNTSITCSCIWVFSFLLGIPSLLYRKLIPVGDRFACQDEGSSIIYTFTVLCLTFLLPLGVILFCYCSIFTMLQSHVNVGR<br>KTNHALKIIFVIIGAFICCWLPFNTFTAVITYGVVQNIDFSCEAWKVLRWGVIISACLAFTNSCINPIIYAFMDQQFRQK<br>VLKSVLGRLCPAKENVQSSTLFFSSTESSVLFGSKKRFIPATIVPVSSGADLHASKRQI |
| | *Tiliqua scincoides* | Gene ID: 136648524<br>mRNA ID: XM_066624641<br>Protein ID: XP_066480738 | MEAGSVTTENWITLSPDYYGSGYVNYDSCPLQELPFAYIFIPILYFMNFLIGFFGNLFVILLMARKQGSKRLVDTFVLNL<br>AVADLVFVCTLPFWALSKALGYWCFGEGLCKVCSYTVSVNRCSSILFLMGMSVERFLVVTKQKDSRSIGTKKITGVSCGC<br>IWMVSFLLGMPSLFYRKLIPTEDTSKSLCEDEEPAIAYSLVFLILTFLLPLGVILFCYCSIFSKLQGHISLSKRRNNALK<br>IIFAIIGAFLCSWLPFNALNAFLIFIMIQKVTLSCQASMTFRWGMTISACLAFTNSCINPIIYVFMDQHFRQQVTKNIPD<br>LCSTRENAQSSTLSFTSTESSLLFANRKKLTPASSAPVEDTYNKGTGF |
| | *Trachemys scripta elegans* | Gene ID: 117876835<br>mRNA ID: XM_034769282<br>Protein ID: XP_034625173 | MTTEAWTVSYEASSAVYWDYNPLQDTCPPQDLPYACVYIPILYFITFFIGFIGNLLVIVLMAKKRGSKRLVDTFVLNLAV<br>ADLVFVLTLPLWAVSAAHGNKWYFGESLCKFSSFIISVNRSSSILFLMGMSVERYLVVMKRVDSKSTGTKNCLIIACGCI<br>WAVSLLLGIPSLMYRKLSLSEDTTGELCEDEDSVAFQVLSLALLILTFLLPLGVILFCYCSISFKLLNHIRLGKGTKNSL<br>QIIFTIIGAFICCWLPFNAFKAFLFFSSMFQGADLSCWALMALRRGLTISACLAFMNSCVNPIIYALMDRHFRRQVLPQR<br>LRVCGAFQETRQTSGLSFSSTTESSLMFASRNKPSSPASIWPEM |
| | *Varanus komodoensis* (Komodo dragon) | Gene ID: 123034042<br>mRNA ID: XM_044451090<br>Protein ID: XP_044307025 | METTELKTGPTATQSWSTGLESLTIDYTDDDICLQEELPFAFVFIPTLYMVIFFTGFFGNLFVILLMARKQSRKRLVDIF<br>VLNLAIADLVFVCTLPFWALSEALSYWCLGEGLCKFSSYALSVNRYSSMLFLTGMSVERFLVMSKHWDSRAIGTKKNICL<br>SCGVIWLISFLLGIPSLIYRKLVFVGNDEYLCQGEEPSATFNLFIFALTFLLPLGIILFCYCSIFSKLRGRASLGKRRSN<br>ALKIIFAIIGAFICSWLPFNTFKAAFIMVQKTDLSCKALTALKWGVHISICLAFTNSCINPVIYVFMDQQFRQQVWKNFP<br>AFCSMRGNIQSSALSLSSTESSLLFAGRKKPAQMGNPSSRVVGL |
| | *Vipera berus* (adder) | Gene ID: 308666168<br>mRNA ID: XM_081324760<br>Protein ID: XP_081180886 | MKISELVTDLAMTKNWSGDLELSTVDYYGSGVTSDDFCLREKLPFASIFISVIYFMIFFIGFFGNLFVVLVMVQKRGNQR<br>LVDNFVLNLAIADLVFVCTLPFWALTEAFDSWWFGEKLCKFSSYAISVNRCSSILFLMGMSVERFLVMNKHWDSRSLGTK<br>RNSNITCSCIWVFSFLLGIPSLLYRKLIPVGDRFSCQDEEPSVTYIFAILFLTFLLPLGVILFCYCSIFSMLRSHVKLGK<br>KTNHALKIIFVIIGAFICCWLPFNTFTAFITYGVVKNIDFSCEVLKVLRRGLIISACLAFTNSCINPIIYAFMDQQFRQK |

| | | | |
|---|---|---|---|
| | | | VLKNFLGLCHPKENVQSSSLFFSSTESSLMFGSKKRFIPATKVPVTSGAGLYASKRQI |
| | *Vipera ursinii* | Gene ID: 310638390<br>mRNA ID: XM_083359439<br>Protein ID: XP_083215565 | MKISELVTDLAMTKNWSGDLELSTVDYYGSGVTSDDFCLREKLPFASIFISVIYFMIFFIGFFGNLFVVLVMVQKRGNQR<br>LVDNFVLNLAIADLVFVCTLPFWALTEAFDSWWFGEKLCKFSSYAISVNRCSSILFLMGMSVERFLVMNKHWDSRSLGTK<br>RNSNITCSCIWVFSFLLGIPSLLYRKLIPVGDRFSCQDEEPSVIYIFAILFLTFLLPLGVILFCYCSIFSMLRSHVKLGR<br>KTNHALKIIFVIIGAFICCWLPFNTFTAFITYGVVKNIDFSCEVLKVLRRGLIISACLAFTNSCINPIIYAFMDQQFRQK<br>VLKNFLGLCHPKENVQSSSLFFSSTESSLMFGGKKRFIPATKVPVTSGAGLYASKRQI |
| | *Zootoca vivipara* (common lizard) | Gene ID: 118088240<br>mRNA ID: XM_035121611<br>Protein ID: XP_034977502 | MEPAQMVTGSSMTETWSGDFESSTIDNYGDLANYDICHLQELPFAYIFIPTVYYITFLIGFFGNLFVILLMARKEGRKRL<br>VDTFVLNLAVADLVFVCTLPFWAVSKALGYWCFGEGLCKFSSYAITVNRCSSILFLMGMSVERFLVMTKQWDSRTVSTKK<br>SIAMSCGCIWVFSLLLGMPSLFYRKLNLVDDRFLCHDKPSPTFSLIMLILTFLLPLGVILFCYCSIFSKLRSHVHLGKRR<br>NNALKIIFAIVGAFICSWLPFNVLKVFFVFTMAQDIHPPCSAMMALTWGLTISACLAFVNSCVNPIIYVFMDQQFRQQAL<br>KHFPAFCSKRGNVQSSALSFSSTDTSLLFASRKKPTPATSARVGDPESKGVGL |

**Table S4.** Possible *CXCL17* gene relics nearby *LIPE* gene in genome of some bird species. Information about these avian CXCL17 gene relics was retrieved from the NCBI databases.

| Species | NCBI information | Genomic DNA sequence | N-terminal signal peptide and C-terminal fragment |
|---|---|---|---|
| *Anser cygnoides*<br>(Swan goose) | Genomic Sequence: NC_089909.1;<br>Chromosome 37;<br>Reference Taihu_goose_T2T_genome Primary Assembly<br>Position: 1,010,785-1,011,138 | ATGGGTGCCCCCCCGGCGCTGCTGTTGCTGCTGCTGCTGCTCCTC<br>GGGGGGGGGGGCCTGGGGGCCCCCCCCGCCTCCCCGAGCTCGGG<br>ACCCCCCAGCAATTTGGGGCCCGTGACGTCACCGGCGCCCGTGAC<br>GTCACCGCAGCCCCCCCGCCAGGCCGAGGGGGTGCAAGAAATGG<br>GGGGGGGGGGGGGAAGGATTTGGGTGGGGGGGGGGGGTTATG<br>GGGGGGGGAGGGGGGGATTTTTTGGGGGGGGGTGGGGTGAAGCC<br>CCCCCCCTCATTTGTCCCCCCCCCCTCCCCAGGACGTCGCAGGA<br>GGGGGCGCTGCCCCCCCACCAAGGGGCTGGCGCTGCCCCTTTAA | MGAPPALLLLLLLLL<br>GGGGLG.........<br>SPPPHFVPPPSPGRRR<br>RGRCPPTKGLALPL* |
| *Caloenas nicobarica*<br>(Nicobar pigeon) | Genomic Sequence: NW_027017036;<br>Unplaced Scaffold Reference bCalNic1.hap1<br>Position: 35,548-35,864 | ATGGCCCCGTCGCTGCTGCTGCTGCTGCTGCTGCTGCTGGGGGGG<br>GTCCCCGCCGCCCCCCCGCCCCCCCAGACCCTCCCGGGGGGCCG<br>GGGGGGTCCCCGCCGCCCCCCCCCCCGTGCTGGGGGTCTCGCCCC<br>CCCCCGCTGGACACACGGTGAGGGGAGCGCCGGGGGGGGGCCC<br>GGGGGGGGGTTTGGGGCCCCCCCTGACCCCTGTGCTCCCCCCGG<br>ATTCCAGGGTCCTCTCGCCCCCCCCGGAGCCGCCGCCGGTGCCCC<br>CCCCGCTGCCGCCCCCGCGCGCCCACCAGGGGGCTCTCGCTGCCC<br>CTGTAG | MAPSLLLLLLLLLGG<br>VPAAPPPPQT.........<br>PLCSPRIPGSSRPPRS<br>RRRCPPRCRPRAPTR<br>GLSLPL* |
| *Ciconia boyciana*<br>(Oriental stork) | Genomic Sequence: NC_132963.1;<br>Chromosome 30;<br>Reference ASM3463844v1 Primary Assembly<br>Position: 1,728,539-1,728,902<br>Complementary strand | ATGGCTCCGTCGCTGCCGCTGCTGCTGCTGCTGCTGCTGCTGGGG<br>GTCCCCGCCGCCCCCCGGCCCCCCCGGAGCCTCCCGGCGCTGCTG<br>GGGGGGCGGGGGGGCCGCCCCGCCCGCATCGCGCCCCCGGCGCT<br>GCCCCCCGGCCCCGGGGCCGCTCACCGCGCCGGCGGCCCGGCGG<br>CCCCCCGGACGGCCGTGTGCCTGCGGGAGAAAGGGTGAGGCAGC<br>GCCGGGGGGGGGGGGTCGCCGGGGACCCCCTGACCCCTTCTGCC<br>GCCCAGGGTCCTCTCGCCCCCCAGGAGCCGCCGGCGGGGACGGT<br>GCCCCCGCGCTGCCGCCCGCGCGCCCCACCAAGGGGCTCTCCCTG<br>CCGCTGTAG | MAPSLPLLLLLLLLG<br>VPAA.........<br>GPPDPFCRPGSSRPPG<br>AAGGDGAPALPPARP<br>TKGLSLPL* |
| *Dromaius novaehollandiae*<br>(emu) | Genomic Sequence: NC_088137.1;<br>Chromosome 38;<br>Reference bDroNov1.hap1<br>Position: 90,344-90,723 | ATGTGCCGCCTGCCCGCCCTGGCTGCGCCGCCGCTGCTGCTGCTG<br>CTGCTGCTGCTGCTGCTGCTGCCGCCGCCGGGGACCCCGGGGCCC<br>CCCCGGGCTGCGCCGGCCCCCCCCGGCGTCGCCGGTGTCCGGCCC<br>GTGGCCGTCGCCAGGAGCCGCCAGCCGCCGTGCGCCTGCGGGAG<br>AGACGCCGCGGCGCCGCGCGGGGAGGGTGAGGCAGCGCCGGGG<br>GGCCGGGAGGGGCCGGGGGGGCTGGCGACCCCCTGACCCCCCCT<br>CTGTGCCCCCCCCCAGGGCTCCCCCCTGCCGGCGGCCGGGGCCG<br>GTGCCCGCCGTCCGCCCCCCGGCGCTGCCGCCCCCGCACCCCCAC<br>GAGGGGGCTCTCGCTGCCCCTCTAG | MCRLPALAAPPLLLL<br>LLLLLLLPPPG.........<br>PPLCPPPGLPPAGGR<br>GRCPPSAPRRCRPRT<br>PTRGLSLPL* |
| *Patagioenas fasciata*<br>(band-tailed pigeon) | Genomic Sequence: NC_092550.1;<br>Chromosome 31;<br>Reference bPatFas1.hap1<br>Position: 3,902,885-3,903,250 | ATGGCCCCGTGGCTCCTTCTGCTGCTCCTGCTGCTCCTGGGGGGG<br>GTCCCGGCCCCCCCCGCCCCCCAAAACGCCCCCCAACCCCCCCCT<br>CGGGGGGGGGCCCGTCGACCCCCCCCAAATTCGGGAGGTCCCGC<br>GTCCCCCTCTTGGACTCAAGGTGAGAATGGGGGGGGGGGGGGGG<br>GGGGAATTTTGGGGGGGTTTGGGGGTTATGGGGGTTATGGGGGGT<br>GGGGGGGGTATTTTGGGGACCCCCCCCTCCCCTGACCTTTGCCCT<br>CCCCTCCCCCCCATCCAGGCTCGTGCCGCCCGCCCCCCCGGAGCC<br>GCCGCCGCTGCCGCCCCCGCGGAGCCACCAGGGGGCTCTCGCTG<br>CCCATATAG | MAPWLLLLLLLLLG<br>GVPAPPA.........<br>PLPSPPPHPGSCRPPP<br>RSRRRCRPRGATRGL<br>SLPI* |
| *Struthio camelus*<br>(African ostrich) | Genomic Sequence: NC_090980.1;<br>Chromosome 39;<br>Reference bStrCam1.hap1<br>Position: 139,576-139,909 | ATGCGCCGCCCGCTCGCCCTGGCCCTGCTGCTGCTGCTGCTGCTG<br>CTGCTGGGGGGGGGGGACGGGACCCCCCGGCCCCCCCGGCCCCC<br>CCGGGGGGCACCGGCCCCCGTCGGCGCCGGGAGCCGCCGGCCGC<br>TCTGCACCTGCGGGAGAGGAGCCGCGAGTGAGGCAGCGCCCTTT<br>TTTGCGGGGGGGGGGATTTTTTGGGGTACCCCCCCTTTATTTCTTT<br>CTCTCCCCCCCCCCCGCAGGGGTCCCCGCCGCCCCCCGGGGCCGG<br>TGCCCGCCGCCCGCCCGGCGCTGCCGCCCCCGCGCCCCCACCAA | MRRPLALALLLLLLL<br>LLGGGDG.........<br>SPPPAGVPAAPRGRC<br>PPPARRCRPRAPTKG<br>LSLPL* |

| | | | |
|---|---|---|---|
| | | GGGGCTCTCGCTGCCCCTCTAG | |
| *Falco rusticolus* (gyrfalcon) | Genomic Sequence: NC_051189;<br>Chromosome 3;<br>Reference bFalRus1.pri Primary Assembly<br>Position: 103,615,328-103,615,767<br>Complementary strand | ATGGCTCCGCTGTTGCTGCTGGTGCTGCTGCTGCTGCTGGAGGGG<br>GGGCTCCCCTCCGCCCCCCTGCCCTTGCCACACTCGCCCCGGCAG<br>GGGGGGGGGCGGCCCAGCTGCCATCGGCCTTCCCCGGCGCTGCC<br>ACCGCCCCCCCCCCAGGTTGGGGGGGGGGGCGCCCCGGACCCCC<br>TTCCGCGGGGCTGCAGGAGAGACGGTGAGGGGGGGGGAGGGGC<br>GGGGGGCTGGGGGATCCCTCTACCCCCCCCTCGGCTCCTCTCACC<br>TCCACCCCCCAGGGCTGGGGGGGGAATGGGGGGTTTAGGAGGAG<br>TCCGTGTGCCCCCCCTGACCCCTGTGCCCCCCCAGGCAGCCCCCA<br>CTTGCCCCGGAGCCGCCGCCGGGGCCACCGCCCCCACCACTCCC<br>GCCCGCGCCCCCCCACCCGGGGGCTCCTGCTGCCGCTCTAG | MAPLLLLVLLLLLEG<br>GLPSAPLPLPHS……<br>…<br>APPDPCAPPGSPHLP<br>RSRRRGHRPHHSRPR<br>PPTRGLLLPL* |
| *Astur gentilis* (Northern goshawk) | Genomic Sequence: NC_064916;<br>Chromosome 37;<br>Reference bAccGen1.2<br>Position: 36,040-36,433<br>Complementary strand | ATGGGTCCGTTGCTGCTGCTGCTGCTGCTTCTGCTGCTGCCGCCGC<br>CACTCCCGGGGGTCCTCGCCGCTCCCCGGCCCCCCCAGAGCCTCC<br>CAGGGGTGGGGGCTGCCTGCCGGCATCGCCCCTCCCTGCCGATGC<br>CCCCCGCGAAATCTGGGGGTGCTTGCGGCGTTGGTGGCCTGGCGG<br>CCCACCCTGGACGGTCGTCGTGTGCCTGCAGGAGAAAGGGTGAG<br>GGAGTGAGGTGGGGGGGGGGGGTCGGTGGTTTTGGGGGACCCCT<br>CTGACCCCCTCTACCCTCCCCCCCCTCCAGGGTCCTCTCGCCTCCC<br>CAGGAGCCGCCAGGAGCGACGGCGAGCGCCACGCTGCTGCCTGC<br>GAGCCCCCACCAAGGGGCTCTCCCTGCCTCTCTAG | MGPLLLLLLLLLLPP<br>PLPGVL………<br>DPLYPPPPPGSSRLPR<br>SRQERRRAPRCCLRA<br>PTKGLSLPL* |
| *Cuculus canorus* (common cuckoo) | Genomic Sequence: NC_071435;<br>Chromosome 35;<br>Reference bCucCan1.pri<br>Position: 41,107-41,707 | ATGTCTCCGTGGCCGCTTCTGTTGCTGCTGCTGCTGCTGCTGCTGG<br>GGGAAGGGGAGGGCACTGCCCCCCCACGACCCCCCCGCCGCCAT<br>TGGGGGGTTCCTCGGAGACCCCCCACCCACCAGGACCCCCCCCA<br>GATCGGGGGGAAAAGAGGGGTGCATGGAGCCTGCGGGACGGAG<br>GGTGAGGAGGGGGGCAGGGGGTGTGGGGGGCCCTTTGACCCCCC<br>CCTCTAACCCCCCCCATGGGCCCCCAGAGGGTGGGTGGGGGGGG<br>CAGGAGTTTGGGAAACCCCCCTGACCCCCCCCCCCGACCCTACCG<br>GGACCCTCAGAGGGTGGAGGGGTTTTAGAGTTCGGGGGAGCCCC<br>CTGGCCCCCAAGGGACTCACAAAAGGGGGGACAGGGGGTTGG<br>GGTGCCCCTTTGATCCCAACTCCCTTTCCCCCCCCCAGGAACCCC<br>AAGAAGGCGGGGGAGGGGGGCAGGAGTTTGGAGAACCCCCTGA<br>CCCCCCCCCTCTGCCCCCCCAGGGACCCTCAGGAGCCACCGGCG<br>GGGATGGGACACCCCTCGCTGCCGCACCCCCACCAGGGGCCTCG<br>CGCTGCCCCTCTAGCGCTGCCCCTCTAG | MSPWPLLLLLLLLLL<br>GEGEGT………<br>PLTPPLCPPRDPQEPP<br>AGMGHPSLPHPHQG<br>PRAAPLALPL* |
| *Falco cherrug* (Saker falcon) | Genomic Sequence: NC_073699;<br>Chromosome 3;<br>Reference bFalChe1.pri Primary Assembly<br>Position: 104,998,636-104,999,077<br>Complementary strand | ATGGCTCCGCTGTTGCTGCTGGTGCTGCTGCTGCTGCTGGAGGGG<br>GGGCTCCCCTCCGCCCCCCTGCCCTTGCCACACTCGCCCCGGCAG<br>GGGGGGGGGCGGCCCAGCTGCCATCGGCCTTCCCCGGCGCTGCC<br>ACCGCCCCCCCCCCAGGTTGGGGGGGGGGGCGCCCCGGACCCCC<br>CTTCCGCGGGGCTGCAGGAGAGACGGTGAGGGGGGGGGAGGGG<br>CGGGGGGCTGGGGGATCCCTCTACCCCCCCCTCGGCTCCTCTCAC<br>CTCCACCCCCCCAGGGCTGGGGGGGGAATGGGGGGTTTAGGAGG<br>AGTCCGTGTGCCCCCCCTGACCCCTGTGCCCCCCCAGGCAGCCCC<br>CACTTGCCCCGGAGCCGCCGCCGGGGCCACCGCCCCCACCACTC<br>CCGCCCGCGCCCCCCCACCCGGGGGCTCCTGCTGCCGCTCTAG | MAPLLLLVLLLLLEG<br>GLPSAPL………<br>CAPPDPCAPPGSPHL<br>PRSRRRGHRPHHSRP<br>RPPTRGLLLPL* |
| *Falco peregrinus* (peregrine falcon) | Genomic Sequence: NC_073723;<br>Chromosome 3;<br>Reference bFalPer1.pri Primary Assembly<br>Position: 104,917,027-104,917,468<br>Complementary strand | ATGGCTCCGCTGTTGCTGCTGGTGCTACTGCTGCTGCTGGAGGGG<br>GGGCTCCCCTCCGCCCCCCTGCCCTTGCCACACTCGCCCCGGCAG<br>GGGGGGGGGCGGCCCAGCTGCCATCGGCCTTCCCCGGCGCTGCC<br>ACCGCCCCCCCCCCAGGTTGGGGGGGGGGGCGCCCCGGACCCCC<br>TTCCGCGGGGCTGCAGGAGAGACGGTGAGGGGGGGGGAGGGGC<br>GGGGGGCTGGGGGATCCCTCTACCCCCCCCCTCGGCTCCTCTCAC<br>CTCCACCCCCCAGGGCTGGGGGCGAATGGGGGGTTTAGGAGGAG<br>TCCGTGTGCCCCCCCTGACCCCTGTGCCCCCCCAGGCAGCCCCCA<br>CTTGCCCAGGAGCCCGCCGCCGGGGCCACCGCCCCCACCACTCC<br>CGCCCGCGCCCCCCCCACCCGGGGGCTCCTGCTGCCGCTCTAG | MAPLLLLVLLLLLEG<br>GLPSAP………<br>TPLCPPRQPPLAQEP<br>AAGATAPTTPARAPP<br>TRGLLLPL* |
| *Rhea pennata* | Genomic Sequence: NC_084698;<br>Chromosome 36; | ATGTGCCGCCTGCCTGCCCCAGCTGCGCCGCTGCCGCCGCTGCTG<br>CTGCTACTGCTGCTGCTGCTGCCGCCACTGCCAGGACCCCCCCGG | MCRLPAPAAPLPPLL<br>LLLLLLLPPLPGPPR |

| | | | |
|---|---|---|---|
| | Reference bPtePen1.pri Primary Assembly<br>Position: 60,362-60,806 | GCCGTGCCAGCCCCCCTTGGCATTGCCGGCCTCCGCGCCATGGCC<br>CGCCCCGGCCGCCGGCTGCCGTGTGCCTGCGGGAGAGCGGCCGC<br>CACGCCACGCGTGCCGGGTGAGGGGGTCAGGCATTCAGGGGGCA<br>GGAGGGGACAGGGGGGCCCTGGGGGTCAGGAGGGATTAGGGGG<br>ATGGGGGAGCTGGGGGGGTATGGGGGAGCTGGAGCGTTTGGGGG<br>TTCGGGGGTGCCGGGGGGGGTCAGAGTCCCCTGACACCCCTGCA<br>CCCCCCAGGACCCCCCAGGCGGTGCCTGCCACCCCTCCGCCGGC<br>GCTGCCGCCCCCGTGCCCCCACCAAGGGGCTCTCGCTGCCCCTCT<br>AG | ………<br>GGGQSPLTPLHPPGP<br>PRRCLPPLRRRCRPR<br>APTKGLSLPL* |
| *Calonectris borealis* (Cory's shearwater) | Genomic Sequence: NC_134346;<br>Chromosome 35;<br>Reference bCalBor7.hap1.2<br>Position: 528,795-529,160<br>Complementary strand | ATGGCTCCGTTGCTGCTGCTGCTGCTGCTGCTGCTGCTGGGGGTC<br>CCTGCCGCCCCCCGGCCCCCCCAGAGCCTCCCAGGGGTGGGGGG<br>TACCCGCCGGCATCACCCCCCCCCGGCAATGCCCCCCCCGAAGTC<br>TGGGGGTGCTCACAGCGTCAGCGGCCCGGCAGCCCCCCCGCGAC<br>GATCGTGTGCCTGCGGGAGACAGGGTGAGGGAGCGCCGGGGGTG<br>GCGGGGGGGGTGGTTTGGGGGACCCCTCTGACCCCCTCTGCCCCC<br>CCAGGGTCCTCTCACCCCTCCAGGAGCCGCCGGCAGGGACGGTG<br>CCCCCCGCGCTGCCGCCCGCGCGCCCCCACCAGGGGGCTGTCCCT<br>GCCGCTCTAG | MAPLLLLLLLLLLGV<br>PAAPRPPQSLPGVGG<br>TRRHHPPPAMPPPKS<br>GGAHSVSGPAAPPRR<br>SCACGRQGEGAPGV<br>AGGVVWGTPLTPSA<br>PPGSSHPSRSRRQGR<br>CPPRCRPRAPTRGLS<br>LPL* |
| *Buteo buteo* (common buzzard) | Genomic Sequence: NC_134201;<br>Chromosome 31;<br>Reference bButBut1.hap1.1<br>Position: 2,629,699-2,630,689<br>Complementary strand | ATGGGTCCGTTGCTGCTGCTGCTGCTGCTGCTGCTGGTGCCGGGG<br>GTCCTCGCCGCTCCCCGGCCCCCCAGAGCCTGCCGGGGGTGGG<br>GGCCGCCTGCCGGCATCGCCCCTCCCTGGCGATGCGCCCCGGGAA<br>ATCCGGGGGGGCGTGCGGCGTCGGGGGTCCGGCGGCCCCCCCCG<br>GACGGTCGTCGTGTGCCTGCAGGAGAAAGGGTGAGGGAGTGAGG<br>GGGGGGGGTGGAGGGGGGGGGGTGAGGTGGTTTTGGGGGACCC<br>CTCTGACCCCCTCTACCCCCCCCCCCCCCAGGGTCCTCTCGCCTCC<br>CCAGGAGCCGCCAGGAGCGACGGCGACCCCCGCGCTGCTGCCTG<br>CGCCCCCCCACCAAGGGGCTCTCCCTGCCGCTCTAA | MGPLLLLLLLLLVPG<br>VLAAPRPP………<br>*PPLPPPPPGSSRLPRS<br>RQERRRPPRCCLRPP<br>TKGLSLPL* |
| *Mycteria americana* (wood stork) | Genomic Sequence: NW_027445471;<br>Unplaced Scaffold Reference USCA_MyAme_1.0<br>Position: 50,440-50,819 | ATGGCTCCGTCGCTGCCGCTGCTGCTGCTGCTGCTGCTGCTGGGG<br>GTCCCCGCCGCCCCCCGGCCCCCCGGAGCCTCCCGGCGCTGCTG<br>GGGGGGGGGGGGGGCCGCCCCCGCCCGCATCGCGCCCCCCCGG<br>CGCTGCCCCCCCCCGGCCCCGGGGCCGCTCACCGCGGCGGCGGC<br>CCGGCGGCCCCCCCCGGACGGCCGCGTGCCTGCGGGAGAAAGGG<br>TGAGGCGGCGCCGGGGGCGGCGGGGGGGGGGGGGGTGGCGCC<br>GGGGGACCCCCCTGACCCCCTCTGCCGCCCAGGGTCCTCTCGCCC<br>CCCCAGGAGCCGCCGGCGGGGACGGTGCCACCCGCGCGCCCCCA<br>CCAAGGGGCTCTCCCTGCCGCTGTAG | MAPSLPLLLLLLLLG<br>VPAAPRPP………<br>VAPGDPPDPLCRPGS<br>SRPPRSRRRGRCHPR<br>APTKGLSLPL* |
| *Strix uralensis* (Ural owl) | Genomic Sequence: NW_027436856;<br>Unplaced Scaffold Reference bStrUra1<br>Position: 78,549-78,170<br>Complementary strand | ATGGCTCCGGCGCTGCTGCTGCTGCTGCTGCTGCTGCCGCTGCTG<br>GGGGGGGTCCCCGCCGCCCCCCGGCCCCCCCAGAGCTTCCCAGG<br>GGTGGGAGATGCCCGCCGGCATCGCCCCCCCTTGGCGATGCTCCC<br>ACCGAAGTCTGGGGGTGCTGACGGCATCGGCGGCCTGGCAGCCC<br>CCCCGGGACGGCCGTGTGCCTGCGAGAGAGAGGGTGAGGGAGC<br>GCGGGGGGCGGGGTGGGGCGTTCGGGGGACCCCTCTGACCCCCC<br>GGCCCCCTCTGCCTCCCCAGGGTCCTCTCGCGCCCCCCGGAGCCG<br>CTGGCGGGGACGGTGCTCCCGGCGCTGCCGCCCGCGCGCCCCCA<br>CCAAGGGGCTCTCCCTGCCCCTCTAG | MAPALLLLLLLLPLL<br>GGVPAAPRPP………<br>DPPAPSASPGSSRAPR<br>SRWRGRCSRRCRPR<br>APTKGLSLPL* |
| *Strix aluco* (Tawny owl) | Genomic Sequence: NW_027436525;<br>Unplaced Scaffold Reference bStrAlu1.hap1<br>Position: 14,908-15,287<br>Complementary strand | ATGGCTCCGGCGCTGCTGCTGCTGCTGCTGCTGCTGCCGCTGCTG<br>GGGGGGGTCCCCGCCGCCCCCCGGCCCCCCCAGAGCTTCCCAGG<br>GGTGGGAGATGCCCGCCGGCATCGCCCCCCCTTGGCGATGCTCCC<br>ACCGAAGTCTGGGGGTGCTGACGGCATCGGCGGCCTGGCAGCCC<br>CCCCGGGACGGCCGTGTGCCTGCGAGAGAGAGGGTGAGGGAGC<br>GCGGGGGGCGGGGTGGGGGGTTCGGGGGACCCCTCTGACCCCCC<br>GGCCCCCTCTGCCTCCCCAGGGTCCTCTCGCGCCCCCCGGAGCCG<br>CTGGCGGGGACGGTGCTCCCGGCGCTGCCGCCCGCGCGCCCCCA<br>CCAAGGGGCTCTCCCTGCCCCTCTAG | MAPALLLLLLLLPLL<br>GGVPAAPRPP………<br>DPPAPSASPGSSRAPR<br>SRWRGRCSRRCRPR<br>APTKGLSLPL* |
| *Grus americana* (Whooping crane) | Genomic Sequence: NW_026561436;<br>Unplaced Scaffold Reference | ATGGCTCCGGCGCTGCTGCTGCTGCTGCTGCTGCTGCTGCTGGGG<br>GTCCCCGCCGCCCCCCGGCCCCCCCAAAACCTCCCCGGGGCGGT | MAPALLLLLLLLLLG<br>VPAAPRPP……… |

| | | | |
|---|---|---|---|
| | bGruAme1.mat Primary Assembly<br>Position: 26,531-26,921<br>Complementary strand | GGGGGCCCCCCAGCATCCCCCCCCCCCCCGGCACGGCCTCCCCGA<br>GGCGCAGGGGGGCTCACGGCGGCCCCCCAGCCCCCCCCAGCCGG<br>TCGTGCGCCTGCAGGAGAGAGGGTGAGGGAGCGCTCGGGGGGG<br>GGGGGGGGCGTGGGGGGGGTTTGGGGGACCCCCAGAGCCCTCT<br>GACCCCCCCCTCTGCCCCCCTCCCCCCCCAGGGTCTTCTCGCCCC<br>CCCAGGAGCCGCCAGCGGGGCCGGTGCCCCCCCCGCTGCCACCT<br>GCACCCCCTCACCAAGGGGCTCTCCCTGCCCCTCTAG | PPLCPPPPPGSSRPPR<br>SRQRGRCPPRCHLHP<br>LTKGLSLPL* |
| *Harpia harpyja* (harpy eagle) | Genomic Sequence: NW_026293188; Unplaced Scaffold Reference bHarHar1 primary haplotype<br>Position: 65,284-65,673<br>Complementary strand | ATGGGTCCATTGCTGCTGCTGCTGCTGCTGCTGCTGCCGCGCTGCC<br>CGGGGGGTCCTCGCCGCCCCCCGGCCCCTCCAGAGACTCCCAGG<br>GGTGGGGGCTGCCTGCCGGCATCGCCCCTCCCTGGCGATGCCCCC<br>CCGCGAAGTCTGGGGGTGCTTGCGGCATCGGTGGCCCAGCAGCC<br>CGCCCTGGACGGTCGTCGTGTGCCTGCAGGAGAAAGGGTGAGGG<br>AGTGAGGTGGGGGCGGGCGGGGGGGGCGGTGGTTTTGGGGGAC<br>CCCTCTGACCCCCTCTGCCCCCCCCCCAGGGTCCTCTCGCTTCCCC<br>AGGAGCCGCCAGGAGCGACGGCGAACCCCACGCTGCTGCCTGCG<br>CACCCCCACCAAGGGGCTCTCCCTGCCTCTCTAG | MGPLLLLLLLLLPRC<br>PGGPRRPP………<br>PLTPSAPPPGSSRFPR<br>SRQERRRTPRCCLRT<br>PTKGLSLPL* |
| *Apteryx rowi* (Okarito brown kiwi) | Genomic Sequence: NW_020447324; Unplaced Scaffold Reference aptRow1<br>Position: 36,091-36,398 | ATGCGCCTCCTGCCCGCCCCGGCGCTGCTGCTGCTGCTGCCGCTG<br>CTGCTGCCGCTGCTGCTGGGGGTCCCGGGACCCCCCCGGGCTGCC<br>CCAGGCCCTCCCGCGGCAAGGAGCCGCCGGCCACCCTGTGCCTG<br>CGGGAGAGGTGAGGCAGCACTGGGGGGGTTGGGGGGGTGTTAGC<br>GCCCCCCTGACCCCCCCCCTTTTTGTGTTCCCCCCCAGGGCTCCCC<br>GCTGCCAGCAGCCGCTGCCCGCCACCCCCTGCCAGGCGCTGCCG<br>CCCCCGCGCCCCCACCCAGGGGCTCTCGCTGCCCCTCTAG | MRLLPAPALLLLLPL<br>LLPLLLGV………<br>*PPPFLCSPPGLPAAS<br>SRCPPPPARRCRPRA<br>PTQGLSLPL* |
| *Apteryx mantelli* (North Island brown kiwi) | Genomic Sequence: NW_027118507; Unplaced Scaffold Reference bAptMan1.hap1<br>Position: 162,154-162,461<br>Complementary strand | ATGCGCCTCCTGCCCGCCCCGGCGCTGCTGCTGCTGCTGCCGCTG<br>CTGCTGCCGCTGCTGCTGGGGGTCCCGGGACCCCCCCGGGCCGCC<br>CCAGGCCCTCCCGCGGCAAGGAGCCGCCGGCCACCCTGTGCCTG<br>CGGGAGAGGTGAGGCAGCACTGGGGGGGTTGGGGGGGTGTTAGC<br>GCCCCCCTGACCCCCCCCCTTTTTGTGTTCCCCCCCAGGGCTCCCC<br>GCTGCCAGCAGCCGCTGCCCGCCACCCCCTGCCAGGCGCTGCCG<br>CCCCCGCGCCCCCACCCAGGGGCTCTCGCTGCCCCTCTAG | MRLLPAPALLLLLPL<br>LLPLLLGVP………<br>PPPFLCSPPGLPAASS<br>RCPPPPARRCRPRAP<br>TQGLSLPL* |
| *Phaethon aethereus* | Genomic Sequence: NC_138606;<br>Chromosome 30;<br>Position: 1,704,307-1,704,680<br>Complementary strand | ATGGCTCCGTGGCTGCTGCTGCTGCTGCTGTTGCTGCTGGGGGTC<br>CCCGCTGCCCCCCAGCCTCCCCAGAGCTTCCAGGGGTGGGAGGT<br>GCCCCCCGGCATCACCCCGCCTCAGCGATACCCCCCCCGAAGTCT<br>GGGGGTGCTCACAGCATCGGCAGCCTGTCAGCCACCCATGGACG<br>GTCGTGTGCCTGCGACAGAAAGGGTGAGGGAGTGCTGGCGGGGG<br>GGCGCGGGGGTTTTGGGGGACCCCTCTGCCTGCCTCTGACCCTCT<br>CTGCCCCCCCAGGGTCCTCTCGCCTCCCCAGGAGCCGCCGGCGGA<br>GACGGTGCCCCCCCCGCTGCCGCCCACGTGCTCCCACCAAGGGG<br>CTCTCCCTGCCCCTCTAG | MAPWLLLLLLLLLG<br>VPAAPQPPQ………<br>LPLTLSAPPGSSRLPR<br>SRRRRRCPPRCRPRA<br>PTKGLSLPL* |

**Table S5.** Other CXCL17 orthologs used in this study. The information is downloaded from the NCBI gene database (https://www.ncbi.nlm.nih.gov/gene). [a] zebrafish CXCL17; [b] zebrafish CXCL17-like

| Class | Species | Gene ID | mRNA ID | Protein ID | Amino acid sequence |
|---|---|---|---|---|---|
| Mammals | *Homo sapiens* (human) | 284340 | NM_198477 | NP_940879 | MKVLISSLLLLLPLMLMSMVSSSLNPGVARGHRDRGQASR RWLQEGGQECECKDWFLRAPRRKFMTVSGLPKKQCPCDH FKGNVKKTRHQRHHRKPNKHSRACQQFLKQCQLRSFALPL |
| | *Mus musculus* (house mouse) | 232983 | NM_153576 | NP_705804 | MKLLASPFLLLLPVMLMSMVFSSPNPGVARSHGDQHLAPR RWLLEGGQECECKDWFLQAPKRKATAVLGPPRKQCPCDHV KGREKKNRHQKHHRKSQRPSRACQQFLKRCHLASFALPL |
| | *Vombatus ursinus* (common wombat) | 114040160 | XM_027858170 | XP_027713971 | MRAPVFFLLLLLLLPPLSVASSSPNPEDAEGQRNHKLPAKRH ARGGRQVCRCQDVFQNIHGGKRVQRAKPPARQCPCDRLK YKQKKPGLGHRTHWRRSCLGFLKQCQLRDITLPL |
| | *Sarcophilus harrisii* (Tasmanian devil) | 116422895 | XM_031963247 | XP_031819107 | MRTPVLSFLLLFLLLLLSITSFSSDPEAAGGQRHHKLHAWR HPQGRRQLCRCEDVFQKIHERRKLQVAKSPASHCPCVRPKH KWKKPGHQKHWRQSCFQFLKQCQLQSILLPL |
| | *Ornithorhynchus anatinus* (platypus) | 103166888 | XM_029066640 | XP_028922473 | MQLSTWSLLLLLLLLLTFATVTSVPHNPGGSRSHGERRQEA RRRLGQITHKCRCPGLSQELQRESRARRLQSPRGECPCDNL KVNLKKRPVWHHKGKEGRVHYRHKEACRLFWKQCELEG LSLPI |
| | *Tachyglossus aculeatus* (Australian echidna) | 119946135 | XM_038767563 | XP_038623491 | MRLTTCSLLLLLLLLLTFATVTSVPQNPGDRRSHGERRLAAR RRLGELTHKCRCPGSSQELQRESRARRLQSPRGECPCDDLK VNLKKRPMWHHKGKEGRVHYRRKEACRLFWKQCELEGL SLPI |
| Amphibian | *Xenopus tropicalis* (tropical clawed frog) | 116412198 | XM_031905867 | XP_031761727 | MRTVAVFMFLAAITLCYCWTPQRQDSSGRTNENRGLAKRN GHKDALGSPLRPERHHLRCCERLSPQHPQRKRRHMRAPRA HPIHQRRRCTTNCRGKKPSKFMVPLPLS |
| Fishes | *Danio rerio* [a] (zebrafish) | 100151367 | NM_001144821 | NP_001138293 | MKTMNFQILVLAFAVMIVTNIQCEARPQEGKSDKSAEVKG HAMPRKCNCQVRGTALDRNCVCEMPHKSRPTLNPEQKNM CLKKKIKTFRKCLQFMGANKKIAKGASLPI |
| | *Danio rerio* [b] (zebrafish) | 100536854 | NM_001386806 XM_073906074 | NP_001373735 XP_073762175 | MTKPICLVFALLILTIILCNNSVCSQRRSMKQSAVCGCKLYPD KGLKCTKRPNPKSRDEYYEILKCICRDTQIFSKSSRKEYLKR CNKFYPSLPL |
| | *Latimeria chalumnae* (coelacanth) | 106704074 | XM_014490303 | XP_014345789 | MKVSDVLVLLCAFTLSSCLQNSGSSDQEEGRVPEEASTVGA VKPQAGSCSCGDPVGSLQNRLVSLKAGPSKHCDCRRSLKE AVNIKRPQAWAWKSKPSRKRCKRKGDNGKIQIWGCRRKTI KPIIIPIPA |

**6×His-Cc-CXCL17**

```
1   CTT TAA GAA GGA GAT ATA ATG CAT CAT CAC CAC CAT CAC AGC CCG CAC GTG CCA GGT GGC GGT CAG GGC GCG GAA
    GAA ATT CTT CCT CTA TAT TAC GTA GTA GTG GTG GTA GTG TCG GGC GTG CAC GGT CCA CCG CCA GTC CCG CGC CTT
                             M   H   H   H   H   H   H   S   P   H   V   P   G   G   G   Q   G   A   E

76  GGT GAG CTG AAC CCT ACC CGT CGC CGT GCA ACG GGC GCC GGC GTG TGT GAA TGC CCG GGC TGG AGT GGT TCT CGT
    CCA CTC GAC TTG GGA TGG GCA GCG GCA CGT TGC CCG CGG CCG CAC ACA CTT ACG GGC CCG ACC TCA CCA AGA GCA
     G   E   L   N   P   T   R   R   R   A   T   G   A   G   V   C   E   C   P   G   W   S   G   S   R

151 CGC CAA CTG GCT GGC GAT CTC CGT GTT GGT GCA CGT CAC AAT GGC CTG TGT CCA TGC GAC TCC ATT GTA CCT CAC
    GCG GTT GAC CGA CCG CTA GAG GCA CAA CCA CGT GCA GTG TTA CCG GAC ACA GGT ACG CTG AGG TAA CAT GGA GTG
     R   Q   L   A   G   D   L   R   V   G   A   R   H   N   G   L   C   P   C   D   S   I   V   P   H

226 GGC CAG GGT CGT GGC CGC CAT CCG CAG GGT CGT GGC CGT CGC GGC CAA CAT CAC GGC CGT AAA CAG CAA CGC TGT
    CCG GTC CCA GCA CCG GCG GTA GGC GTC CCA GCA CCG GCA GCG CCG GTT GTA GTG CCG GCA TTT GTC GTT GCG ACA
     G   Q   G   R   G   R   H   P   Q   G   R   G   R   R   G   Q   H   H   G   R   K   Q   Q   R   C

301 CAT GGT ATC CCA TCA CGC TGC CGT CAG TTA GGC TTG GCG AAG GGT CTG TTG TTA CCG CTG TAA GCG GCC GCA CTC
    GTA CCA TAG GGT AGT GCG ACG GCA GTC AAT CCG AAC CGC TTC CCA GAC AAC AAT GGC GAC ATT CGC CGG CGT GAG
     H   G   I   P   S   R   C   R   Q   L   G   L   A   K   G   L   L   L   P   L   *
```

**6×His-SmBiT-Cc-CXCL17**

```
1   CTT TAA GAA GGA GAT ATA ATG CAT CAT CAC CAC CAT CAC GGT GTG ACC GGC TAC CGT CTG TTT GAA GAA ATT CTG
    GAA ATT CTT CCT CTA TAT TAC GTA GTA GTG GTG GTA GTG CCA CAC TGG CCG ATG GCA GAC AAA CTT CTT TAA GAC
                             M   H   H   H   H   H   H   G   V   T   G   Y   R   L   F   E   E   I   L

76  GGC GGC AGC CCG CAC GTG CCA GGT GGC GGT CAG GGC GCG GAA GGT GAG CTG AAC CCT ACC CGT CGC CGT GCA ACG
    CCG CCG TCG GGC GTG CAC GGT CCA CCG CCA GTC CCG CGC CTT CCA CTC GAC TTG GGA TGG GCA GCG GCA CGT TGC
     G   G   S   P   H   V   P   G   G   G   Q   G   A   E   G   E   L   N   P   T   R   R   R   A   T

151 GGC GCC GGC GTG TGT GAA TGC CCG GGC TGG AGT GGT TCT CGT CGC CAA CTG GCT GGC GAT CTC CGT GTT GGT GCA
    CCG CGG CCG CAC ACA CTT ACG GGC CCG ACC TCA CCA AGA GCA GCG GTT GAC CGA CCG CTA GAG GCA CAA CCA CGT
     G   A   G   V   C   E   C   P   G   W   S   G   S   R   R   Q   L   A   G   D   L   R   V   G   A

226 CGT CAC AAT GGC CTG TGT CCA TGC GAC TCC ATT GTA CCT CAC GGC CAG GGT CGT GGC CGC CAT CCG CAG GGT CGT
    GCA GTG TTA CCG GAC ACA GGT ACG CTG AGG TAA CAT GGA GTG CCG GTC CCA GCA CCG GCG GTA GGC GTC CCA GCA
     R   H   N   G   L   C   P   C   D   S   I   V   P   H   G   Q   G   R   G   R   H   P   Q   G   R

301 GGC CGT CGC GGC CAA CAT CAC GGC CGT AAA CAG CAA CGC TGT CAT GGT ATC CCA TCA CGC TGC CGT CAG TTA GGC
    CCG GCA GCG CCG GTT GTA GTG CCG GCA TTT GTC GTT GCG ACA GTA CCA TAG GGT AGT GCG ACG GCA GTC AAT CCG
     G   R   R   G   Q   H   H   G   R   K   Q   Q   R   C   H   G   I   P   S   R   C   R   Q   L   G

376 TTG GCG AAG GGT CTG TTG TTA CCG CTG TAA GCG GCC GCA CTC GAG CAC
    AAC CGC TTC CCA GAC AAC AAT GGC GAC ATT CGC CGG CGT GAG CTC GTG
     L   A   K   G   L   L   L   P   L   *
```

**Fig. S1.** The nucleotide and amino acid sequence of Cc-CXCL17 proteins in pET expression vector. The amino acid sequence of mature Cc-CXCL17 is shown in red, and that of SmBiT in blue.

##### Cc-GPR25 in pcDNA3.1 or PB-TRE vector

```
1    GCT AGC ATG ACA ACT GAA CCC TGG ACT GTA AAT TAT GAA GCA TCT TCT GCT GTC TAC TCG GAT TAT AAT CCC CTC
     CGA TCG TAC TGT TGA CTT GGG ACC TGA CAT TTA ATA CTT CGT AGA AGA CGA CAG ATG AGC CTA ATA TTA GGG GAG
              M   T   T   E   P   W   T   V   N   Y   E   A   S   S   A   V   Y   S   D   Y   N   P   L

76   CAA GAT ACC TGC CTA CCA CAG GAC CTA CCC TAC GCT TGG TGC TAC ATT CCC ATC CTC TAC TTC ATC GCT TTC TTC
     GTT CTA TGG ACG GAT GGT GTC CTG GAT GGG ATG CGA ACC ACG ATG TAA GGG TAG GAG ATG AAG TAG CGA AAG AAG
      Q   D   T   C   L   P   Q   D   L   P   Y   A   W   C   Y   I   P   I   L   Y   F   I   A   F   F

151  GTT GGC TTC ATT GGC AAC CTC TTG GTG ATT GTA CTG ATG GCC AAG AAG AAA GGC AGC AAG AGG CTG GTG GAT ACC
     CAA CCG AAG TAA CCG TTG GAG AAC CAC TAA CAT GAC TAC CGG TTC TTC TTT CCG TCG TTC TCC GAC CAC CTA TGG
      V   G   F   I   G   N   L   L   V   I   V   L   M   A   K   K   K   G   S   K   R   L   V   D   T

226  TTT GTC CTG AAT CTG GCT GTT GCC GAC CTG GTC TTT GTC TGC ACC TTG CCT CTC TGG GCA GTG TCA GCA GCT TAT
     AAA CAG GAC TTA GAC CGA CAA CGG CTG GAC CAG AAA CAG ACG TGG AAC GGA GAG ACC CGT CAC AGT CGT CGA ATA
      F   V   L   N   L   A   V   A   D   L   V   F   V   C   T   L   P   L   W   A   V   S   A   A   Y

301  GAT AAC AAG TGG TAC TTT GGT GAA GGC ATG TGT AAA TTC AGC AGC TTT ATC ATC TCT GTG AAC CGC AGC TCC AGC
     CTA TTG TTC ACC ATG AAA CCA CTT CCG TAC ACA TTT AAG TCG TCG AAA TAG TAG AGA CAC TTG GCG TCG AGG TCG
      D   N   K   W   Y   F   G   E   G   M   C   K   F   S   S   F   I   I   S   V   N   R   S   S   S

376  ATC CTG TTC CTA ATG GGC ATG AGC GTG GAG CGT TAC CTG GTG GTC ATG AAG CGT GTG GAC TCC AAG TCT ACT GGG
     TAG GAC AAG GAT TAC CCG TAC TCG CAC CTC GCA ATG GAC CAC CAG TAC TTC GCA CAC CTG AGG TTC AGA TGA CCC
      I   L   F   L   M   G   M   S   V   E   R   Y   L   V   V   M   K   R   V   D   S   K   S   T   G

451  ACC AAG AAC TGT CTG ATC ATT GCC TGT GGT TGC ATC TGG GCT GTC TCT CTC CTC CTA GGC ATT CCC TCT ATG ATG
     TGG TTC TTG ACA GAC TAG TAA CGG ACA CCA ACG TAG ACC CGA CAG AGA GAG GAG GAT CCG TAA GGG AGA TAC TAC
      T   K   N   C   L   I   I   A   C   G   C   I   W   A   V   S   L   L   L   G   I   P   S   M   M

526  TAT AGG AAG CTC AGC CCC TCA GAG GAT ACC ACT GGT GAA CTG TGC GAG GAT GAG GAC TCT ATC ACC TTC CAG GTG
     ATA TCC TTC GAG TCG GGG AGT CTC CTA TGG TGA CCA CTT GAC ACG CTC CTA CTC CTG AGA TAG TGG AAG GTC CAC
      Y   R   K   L   S   P   S   E   D   T   T   G   E   L   C   E   D   E   D   S   I   T   F   Q   V

601  CTC AGC CTG GCT TTG CTT ATC TTG ACC TTT CTC CTG CCC TTG GGA GTG ATC TTA TTC TGC TAT TGC TCC ATC TCC
     GAG TCG GAC CGA AAC GAA TAG AAC TGG AAA GAG GAC GGG AAC CCT CAC TAG AAT AAG ACG ATA ACG AGG TAG AGG
      L   S   L   A   L   L   I   L   T   F   L   L   P   L   G   V   I   L   F   C   Y   C   S   I   S

676  TTC AAG CTG CTG AAC CAC ATC AGC TTG GGA AAG GGG ACA AAG AAC TCC CTC CAG ATC ATC TTC ACG ATC ATC GGT
     AAG TTC GAC GAC TTG GTG TAG TCG AAC CCT TTC CCC TGT TTC TTG AGG GAG GTC TAG TAG AAG TGC TAG TAG CCA
      F   K   L   L   N   H   I   S   L   G   K   G   T   K   N   S   L   Q   I   I   F   T   I   I   G

751  GCC TTC ATC TGC TGC TGG CTG CCC TTC AAT GCC TTT AAA GCG TTC CTC TTC TTC TCC TCT ACG TTC CAG GGA GCA
     CGG AAG TAG ACG ACG ACC GAC GGG AAG TTA CGG AAA TTT CGC AAG GAG AAG AAG AGG AGA TGC AAG GTC CCT CGT
      A   F   I   C   C   W   L   P   F   N   A   F   K   A   F   L   F   F   S   S   T   F   Q   G   A

826  GAT CTG TCT TGC TGG GCA TTA ATG GCT CTC AGA CGG GGC CTG ACC ATC AGT GCT TGC TTG GCC TTC ATG AAC AGC
     CTA GAC AGA ACG ACC CGT AAT TAC CGA GAG TCT GCC CCG GAC TGG TAG TCA CGA ACG AAC CGG AAG TAC TTG TCG
      D   L   S   C   W   A   L   M   A   L   R   R   G   L   T   I   S   A   C   L   A   F   M   N   S

901  TGC GTA AAT CCC ATC ATC TAT GCC TTG ATG GAT CAC CAT TTC CGA CGC CAG GTG CTG CAA CAA CGG CTC CGG GTC
     ACG CAT TTA GGG TAG TAG ATA CGG AAC TAC CTA GTG GTA AAG GCT GCG GTC CAC GAC GTT GTT GCC GAG GCC CAG
      C   V   N   P   I   I   Y   A   L   M   D   H   H   F   R   R   Q   V   L   Q   Q   R   L   R   V

976  TGT GGT GCT TTC CAG GAG ACA CGG CAA ACC TCG GTC TTG TCC GTC TCA TCC GCT ACT GAG TCC AGC CTC ATG TTC
     ACA CCA CGA AAG GTC CTC TGT GCC GTT TGG AGC CAG AAC AGG CAG AGT AGG CGA TGA CTC AGG TCG GAG TAC AAG
      C   G   A   F   Q   E   T   R   Q   T   S   V   L   S   V   S   S   A   T   E   S   S   L   M   F

1051 GCC AGC CGG AAC AAG CCA CCA TCA CCT GCC TCC ATT CGG CCG GAG ATG TAA GCG GCC GC
     CGG TCG GCC TTG TTC GGT GGT AGT GGA CGG AGG TAA GCC GGC CTC TAC ATT CGC CGG CG
      A   S   R   N   K   P   P   S   P   A   S   I   R   P   E   M   *
```

##### Cc-GPR25-LgBiT in pTRE3G-BI vector coexpressing with SmBiT-ARRB2

```
1    GCT AGC ATG ACA ACT GAA CCC TGG ACT GTA AAT TAT GAA GCA TCT TCT GCT GTC TAC TCG GAT TAT AAT CCC CTC
     CGA TCG TAC TGT TGA CTT GGG ACC TGA CAT TTA ATA CTT CGT AGA AGA CGA CAG ATG AGC CTA ATA TTA GGG GAG
              M   T   T   E   P   W   T   V   N   Y   E   A   S   S   A   V   Y   S   D   Y   N   P   L

76   CAA GAT ACC TGC CTA CCA CAG GAC CTA CCC TAC GCT TGG TGC TAC ATT CCC ATC CTC TAC TTC ATC GCT TTC TTC
     GTT CTA TGG ACG GAT GGT GTC CTG GAT GGG ATG CGA ACC ACG ATG TAA GGG TAG GAG ATG AAG TAG CGA AAG AAG
      Q   D   T   C   L   P   Q   D   L   P   Y   A   W   C   Y   I   P   I   L   Y   F   I   A   F   F

151  GTT GGC TTC ATT GGC AAC CTC TTG GTG ATT GTA CTG ATG GCC AAG AAG AAA GGC AGC AAG AGG CTG GTG GAT ACC
     CAA CCG AAG TAA CCG TTG GAG AAC CAC TAA CAT GAC TAC CGG TTC TTC TTT CCG TCG TTC TCC GAC CAC CTA TGG
      V   G   F   I   G   N   L   L   V   I   V   L   M   A   K   K   K   G   S   K   R   L   V   D   T

226  TTT GTC CTG AAT CTG GCT GTT GCC GAC CTG GTC TTT GTC TGC ACC TTG CCT CTC TGG GCA GTG TCA GCA GCT TAT
     AAA CAG GAC TTA GAC CGA CAA CGG CTG GAC CAG AAA CAG ACG TGG AAC GGA GAG ACC CGT CAC AGT CGT CGA ATA
      F   V   L   N   L   A   V   A   D   L   V   F   V   C   T   L   P   L   W   A   V   S   A   A   Y
```

```
301  GAT AAC AAG TGG TAC TTT GGT GAA GGC ATG TGT AAA TTC AGC AGC TTT ATC ATC TCT GTG AAC CGC AGC TCC AGC
     CTA TTG TTC ACC ATG AAA CCA CTT CCG TAC ACA TTT AAG TCG TCG AAA TAG TAG AGA CAC TTG GCG TCG AGG TCG
      D   N   K   W   Y   F   G   E   G   M   C   K   F   S   S   F   I   I   S   V   N   R   S   S   S

376  ATC CTG TTC CTA ATG GGC ATG AGC GTG GAG CGT TAC CTG GTG GTC ATG AAG CGT GTG GAC TCC AAG TCT ACT GGG
     TAG GAC AAG GAT TAC CCG TAC TCG CAC CTC GCA ATG GAC CAC CAG TAC TTC GCA CAC CTG AGG TTC AGA TGA CCC
      I   L   F   L   M   G   M   S   V   E   R   Y   L   V   V   M   K   R   V   D   S   K   S   T   G

451  ACC AAG AAC TGT CTG ATC ATT GCC TGT GGT TGC ATC TGG GCT GTC TCT CTC CTC CTA GGC ATT CCC TCT ATG ATG
     TGG TTC TTG ACA GAC TAG TAA CGG ACA CCA ACG TAG ACC CGA CAG AGA GAG GAG GAT CCG TAA GGG AGA TAC TAC
      T   K   N   C   L   I   I   A   C   G   C   I   W   A   V   S   L   L   L   G   I   P   S   M   M

526  TAT AGG AAG CTC AGC CCC TCA GAG GAT ACC ACT GGT GAA CTG TGC GAG GAT GAG GAC TCT ATC ACC TTC CAG GTG
     ATA TCC TTC GAG TCG GGG AGT CTC CTA TGG TGA CCA CTT GAC ACG CTC CTA CTC CTG AGA TAG TGG AAG GTC CAC
      Y   R   K   L   S   P   S   E   D   T   T   G   E   L   C   E   D   E   D   S   I   T   F   Q   V

601  CTC AGC CTG GCT TTG CTT ATC TTG ACC TTT CTC CTG CCC TTG GGA GTG ATC TTA TTC TGC TAT TGC TCC ATC TCC
     GAG TCG GAC CGA AAC GAA TAG AAC TGG AAA GAG GAC GGG AAC CCT CAC TAG AAT AAG ACG ATA ACG AGG TAG AGG
      L   S   L   A   L   L   I   L   T   F   L   L   P   L   G   V   I   L   F   C   Y   C   S   I   S

676  TTC AAG CTG CTG AAC CAC ATC AGC TTG GGA AAG GGG ACA AAG AAC TCC CTC CAG ATC ATC TTC ACG ATC ATC GGT
     AAG TTC GAC GAC TTG GTG TAG TCG AAC CCT TTC CCC TGT TTC TTG AGG GAG GTC TAG TAG AAG TGC TAG TAG CCA
      F   K   L   L   N   H   I   S   L   G   K   G   T   K   N   S   L   Q   I   I   F   T   I   I   G

751  GCC TTC ATC TGC TGC TGG CTG CCC TTC AAT GCC TTT AAA GCG TTC CTC TTC TTC TCC TCT ACG TTC CAG GGA GCA
     CGG AAG TAG ACG ACG ACC GAC GGG AAG TTA CGG AAA TTT CGC AAG GAG AAG AAG AGG AGA TGC AAG GTC CCT CGT
      A   F   I   C   C   W   L   P   F   N   A   F   K   A   F   L   F   F   S   S   T   F   Q   G   A

826  GAT CTG TCT TGC TGG GCA TTA ATG GCT CTC AGA CGG GGC CTG ACC ATC AGT GCT TGC TTG GCC TTC ATG AAC AGC
     CTA GAC AGA ACG ACC CGT AAT TAC CGA GAG TCT GCC CCG GAC TGG TAG TCA CGA ACG AAC CGG AAG TAC TTG TCG
      D   L   S   C   W   A   L   M   A   L   R   R   G   L   T   I   S   A   C   L   A   F   M   N   S

901  TGC GTA AAT CCC ATC ATC TAT GCC TTG ATG GAT CAC CAT TTC CGA CGC CAG GTG CTG CAA CAA CGG CTC CGG GTC
     ACG CAT TTA GGG TAG TAG ATA CGG AAC TAC CTA GTG GTA AAG GCT GCG GTC CAC GAC GTT GTT GCC GAG GCC CAG
      C   V   N   P   I   I   Y   A   L   M   D   H   H   F   R   R   Q   V   L   Q   Q   R   L   R   V

976  TGT GGT GCT TTC CAG GAG ACA CGG CAA ACC TCG GTC TTG TCC GTC TCA TCC GCT ACT GAG TCC AGC CTC ATG TTC
     ACA CCA CGA AAG GTC CTC TGT GCC GTT TGG AGC CAG AAC AGG CAG AGT AGG CGA TGA CTC AGG TCG GAG TAC AAG
      C   G   A   F   Q   E   T   R   Q   T   S   V   L   S   V   S   S   A   T   E   S   S   L   M   F

1051 GCC AGC CGG AAC AAG CCA CCA TCA CCT GCC TCC ATT CGG CCG GAG ATG CCA CCG GTC GGT ACC GGC GGA GGG TCT
     CGG TCG GCC TTG TTC GGT GGT AGT GGA CGG AGG TAA GCC GGC CTC TAC GGT GGC CAG CCA TGG CCG CCT CCC AGA
      A   S   R   N   K   P   P   S   P   A   S   I   R   P   E   M   P   P   V   G   T   G   G   G   S

1126 AGC AGT GGC GGT GGG ATG GTC TTC ACA CTC GAA GAT TTC GTT GGG GAC TGG GAA CAG ACA GCC GCC TAC AAC CTG
     TCG TCA CCG CCA CCC TAC CAG AAG TGT GAG CTT CTA AAG CAA CCC CTG ACC CTT GTC TGT CGG CGG ATG TTG GAC
      S   S   G   G   G   M   V   F   T   L   E   D   F   V   G   D   W   E   Q   T   A   A   Y   N   L

1201 GAC CAA GTC CTT GAA CAG GGA GGT GTG TCC AGT TTG CTG CAG AAT CTC GCC GTG TCC GTA ACT CCG ATC CAA AGG
     CTG GTT CAG GAA CTT GTC CCT CCA CAC AGG TCA AAC GAC GTC TTA GAG CGG CAC AGG CAT TGA GGC TAG GTT TCC
      D   Q   V   L   E   Q   G   G   V   S   S   L   L   Q   N   L   A   V   S   V   T   P   I   Q   R

1276 ATT GTC CGG AGC GGT GAA AAT GCC CTG AAG ATC GAC ATC CAT GTC ATC ATC CCG TAT GAA GGT CTG AGC GCC GAC
     TAA CAG GCC TCG CCA CTT TTA CGG GAC TTC TAG CTG TAG GTA CAG TAG TAG GGC ATA CTT CCA GAC TCG CGG CTG
      I   V   R   S   G   E   N   A   L   K   I   D   I   H   V   I   I   P   Y   E   G   L   S   A   D

1351 CAA ATG GCC CAG ATC GAA GAG GTG TTT AAG GTG GTG TAC CCT GTG GAT GAT CAT CAC TTT AAG GTG ATC CTG CCC
     GTT TAC CGG GTC TAG CTT CTC CAC AAA TTC CAC CAC ATG GGA CAC CTA CTA GTA GTG AAA TTC CAC TAG GAC GGG
      Q   M   A   Q   I   E   E   V   F   K   V   V   Y   P   V   D   D   H   H   F   K   V   I   L   P

1426 TAT GGC ACA CTG GTA ATC GAC GGG GTT ACG CCG AAC ATG CTG AAC TAT TTC GGA CGG CCG TAT GAA GGC ATC GCC
     ATA CCG TGT GAC CAT TAG CTG CCC CAA TGC GGC TTG TAC GAC TTG ATA AAG CCT GCC GGC ATA CTT CCG TAG CGG
      Y   G   T   L   V   I   D   G   V   T   P   N   M   L   N   Y   F   G   R   P   Y   E   G   I   A

1501 GTG TTC GAC GGC AAA AAG ATC ACT GTA ACA GGG ACC CTG TGG AAC GGC AAC AAA ATT ATC GAC GAG CGC CTG ATC
     CAC AAG CTG CCG TTT TTC TAG TGA CAT TGT CCC TGG GAC ACC TTG CCG TTG TTT TAA TAG CTG CTC GCG GAC TAG
      V   F   D   G   K   K   I   T   V   T   G   T   L   W   N   G   N   K   I   I   D   E   R   L   I

1576 ACC CCC GAC GGC TCC ATG CTG TTC CGA GTA ACC ATC AAC AGT TAA CAT ATG
     TGG GGG CTG CCG AGG TAC GAC AAG GCT CAT TGG TAG TTG TCA ATT GTA TAC
      T   P   D   G   S   M   L   F   R   V   T   I   N   S   *
```

**sLgBiT-Cc-GPR25 in PB-TRE vector**

```
1    GCT AGC ATG AAC TCC TTC TCC ACA AGC GCC TTC GGT CCA GTT GCC TTC TCC CTG GGC CTG CTC CTG GTG TTG CCT
     CGA TCG TAC TTG AGG AAG AGG TGT TCG CGG AAG CCA GGT CAA CGG AAG AGG GAC CCG GAC GAG GAC CAC AAC GGA
              M   N   S   F   S   T   S   A   F   G   P   V   A   F   S   L   G   L   L   L   V   L   P

76   GCT GCC TTC CCT GCC CCA GTC TTC ACA CTC GAA GAT TTC GTT GGG GAC TGG GAA CAG ACA GCC GCC TAC AAC CTG
```

CGA CGG AAG GGA CGG GGT CAG AAG TGT GAG CTT CTA AAG CAA CCC CTG ACC CTT GTC TGT CGG CGG ATG TTG GAC
A A F P A P V F T L E D F V G D W E Q T A A Y N L

151 GAC CAA GTC CTT GAA CAG GGA GGT GTG TCC AGT TTG CTG CAG AAT CTC GCC GTG TCC GTA ACT CCG ATC CAA AGG
CTG GTT CAG GAA CTT GTC CCT CCA CAC AGG TCA AAC GAC GTC TTA GAG CGG CAC AGG CAT TGA GGC TAG GTT TCC
D Q V L E Q G G V S S L L Q N L A V S V T P I Q R

226 ATT GTC CGG AGC GGT GAA AAT GCC CTG AAG ATC GAC ATC CAT GTC ATC ATC CCG TAT GAA GGT CTG AGC GCC GAC
TAA CAG GCC TCG CCA CTT TTA CGG GAC TTC TAG CTG TAG GTA CAG TAG TAG GGC ATA CTT CCA GAC TCG CGG CTG
I V R S G E N A L K I D I H V I I P Y E G L S A D

301 CAA ATG GCC CAG ATC GAA GAG GTG TTT AAG GTG GTG TAC CCT GTG GAT GAT CAT CAC TTT AAG GTG ATC CTG CCC
GTT TAC CGG GTC TAG CTT CTC CAC AAA TTC CAC CAC ATG GGA CAC CTA CTA GTA GTG AAA TTC CAC TAG GAC GGG
Q M A Q I E E V F K V V Y P V D D H H F K V I L P

376 TAT GGC ACA CTG GTA ATC GAC GGG GTT ACG CCG AAC ATG CTG AAC TAT TTC GGA CGG CCG TAT GAA GGC ATC GCC
ATA CCG TGT GAC CAT TAG CTG CCC CAA TGC GGC TTG TAC GAC TTG ATA AAG CCT GCC GGC ATA CTT CCG TAG CGG
Y G T L V I D G V T P N M L N Y F G R P Y E G I A

451 GTG TTC GAC GGC AAA AAG ATC ACT GTA ACA GGG ACC CTG TGG AAC GGC AAC AAA ATT ATC GAC GAG CGC CTG ATC
CAC AAG CTG CCG TTT TTC TAG TGA CAT TGT CCC TGG GAC ACC TTG CCG TTG TTT TAA TAG CTG CTC GCG GAC TAG
V F D G K K I T V T G T L W N G N K I I D E R L I

526 ACC CCC GAC GGC TCC ATG CTG TTC CGA GTA ACC ATC AAC AGT GGT GGC GGC TCT GGT GGT GGC AGC GGC GGT GGT
TGG GGG CTG CCG AGG TAC GAC AAG GCT CAT TGG TAG TTG TCA CCA CCG CCG AGA CCA CCA CCG TCG CCG CCA CCA
T P D G S M L F R V T I N S G G G S G G G S G G G

601 ACC ACA ACT GAA CCC TGG ACT GTA AAT TAT GAA GCA TCT TCT GCT GTC TAC TCG GAT TAT AAT CCC CTC CAA GAT
TGG TGT TGA CTT GGG ACC TGA CAT TTA ATA CTT CGT AGA AGA CGA CAG ATG AGC CTA ATA TTA GGG GAG GTT CTA
T T T E P W T V N Y E A S S A V Y S D Y N P L Q D

676 ACC TGC CTA CCA CAG GAC CTA CCC TAC GCT TGG TGC TAC ATT CCC ATC CTC TAC TTC ATC GCT TTC TTC GTT GGC
TGG ACG GAT GGT GTC CTG GAT GGG ATG CGA ACC ACG ATG TAA GGG TAG GAG ATG AAG TAG CGA AAG AAG CAA CCG
T C L P Q D L P Y A W C Y I P I L Y F I A F F V G

751 TTC ATT GGC AAC CTC TTG GTG ATT GTA CTG ATG GCC AAG AAG AAA GGC AGC AAG AGG CTG GTG GAT ACC TTT GTC
AAG TAA CCG TTG GAG AAC CAC TAA CAT GAC TAC CGG TTC TTC TTT CCG TCG TTC TCC GAC CAC CTA TGG AAA CAG
F I G N L L V I V L M A K K K G S K R L V D T F V

826 CTG AAT CTG GCT GTT GCC GAC CTG GTC TTT GTC TGC ACC TTG CCT CTC TGG GCA GTG TCA GCA GCT TAT GAT AAC
GAC TTA GAC CGA CAA CGG CTG GAC CAG AAA CAG ACG TGG AAC GGA GAG ACC CGT CAC AGT CGT CGA ATA CTA TTG
L N L A V A D L V F V C T L P L W A V S A A Y D N

901 AAG TGG TAC TTT GGT GAA GGC ATG TGT AAA TTC AGC AGC TTT ATC ATC TCT GTG AAC CGC AGC TCC AGC ATC CTG
TTC ACC ATG AAA CCA CTT CCG TAC ACA TTT AAG TCG TCG AAA TAG TAG AGA CAC TTG GCG TCG AGG TCG TAG GAC
K W Y F G E G M C K F S S F I I S V N R S S S I L

976 TTC CTA ATG GGC ATG AGC GTG GAG CGT TAC CTG GTG GTC ATG AAG CGT GTG GAC TCC AAG TCT ACT GGG ACC AAG
AAG GAT TAC CCG TAC TCG CAC CTC GCA ATG GAC CAC CAG TAC TTC GCA CAC CTG AGG TTC AGA TGA CCC TGG TTC
F L M G M S V E R Y L V V M K R V D S K S T G T K

1051 AAC TGT CTG ATC ATT GCC TGT GGT TGC ATC TGG GCT GTC TCT CTC CTC CTA GGC ATT CCC TCT ATG ATG TAT AGG
TTG ACA GAC TAG TAA CGG ACA CCA ACG TAG ACC CGA CAG AGA GAG GAG GAT CCG TAA GGG AGA TAC TAC ATA TCC
N C L I I A C G C I W A V S L L L G I P S M M Y R

1126 AAG CTC AGC CCC TCA GAG GAT ACC ACT GGT GAA CTG TGC GAG GAT GAG GAC TCT ATC ACC TTC CAG GTG CTC AGC
TTC GAG TCG GGG AGT CTC CTA TGG TGA CCA CTT GAC ACG CTC CTA CTC CTG AGA TAG TGG AAG GTC CAC GAG TCG
K L S P S E D T T G E L C E D E D S I T F Q V L S

1201 CTG GCT TTG CTT ATC TTG ACC TTT CTC CTG CCC TTG GGA GTG ATC TTA TTC TGC TAT TGC TCC ATC TCC TTC AAG
GAC CGA AAC GAA TAG AAC TGG AAA GAG GAC GGG AAC CCT CAC TAG AAT AAG ACG ATA ACG AGG TAG AGG AAG TTC
L A L L I L T F L L P L G V I L F C Y C S I S F K

1276 CTG CTG AAC CAC ATC AGC TTG GGA AAG GGG ACA AAG AAC TCC CTC CAG ATC ATC TTC ACG ATC ATC GGT GCC TTC
GAC GAC TTG GTG TAG TCG AAC CCT TTC CCC TGT TTC TTG AGG GAG GTC TAG TAG AAG TGC TAG TAG CCA CGG AAG
L L N H I S L G K G T K N S L Q I I F T I I G A F

1351 ATC TGC TGC TGG CTG CCC TTC AAT GCC TTT AAA GCG TTC CTC TTC TTC TCC TCT ACG TTC CAG GGA GCA GAT CTG
TAG ACG ACG ACC GAC GGG AAG TTA CGG AAA TTT CGC AAG GAG AAG AAG AGG AGA TGC AAG GTC CCT CGT CTA GAC
I C C W L P F N A F K A F L F F S S T F Q G A D L

1426 TCT TGC TGG GCA TTA ATG GCT CTC AGA CGG GGC CTG ACC ATC AGT GCT TGC TTG GCC TTC ATG AAC AGC TGC GTA
AGA ACG ACC CGT AAT TAC CGA GAG TCT GCC CCG GAC TGG TAG TCA CGA ACG AAC CGG AAG TAC TTG TCG ACG CAT
S C W A L M A L R R G L T I S A C L A F M N S C V

1501 AAT CCC ATC ATC TAT GCC TTG ATG GAT CAC CAT TTC CGA CGC CAG GTG CTG CAA CAA CGG CTC CGG GTC TGT GGT
TTA GGG TAG TAG ATA CGG AAC TAC CTA GTG GTA AAG GCT GCG GTC CAC GAC GTT GTT GCC GAG GCC CAG ACA CCA
N P I I Y A L M D H H F R R Q V L Q Q R L R V C G

1576 GCT TTC CAG GAG ACA CGG CAA ACC TCG GTC TTG TCC GTC TCA TCC GCT ACT GAG TCC AGC CTC ATG TTC GCC AGC
CGA AAG GTC CTC TGT GCC GTT TGG AGC CAG AAC AGG CAG AGT AGG CGA TGA CTC AGG TCG GAG TAC AAG CGG TCG

```
     A   F   Q   E   T   R   Q   T   S   V   L   S   V   S   S   A   T   E   S   S   L   M   F   A   S

1651 CGG AAC AAG CCA CCA TCA CCT GCC TCC ATT CGG CCG GAG ATG TAA
     GCC TTG TTC GGT GGT AGT GGA CGG AGG TAA GCC GGC CTC TAC ATT
     R   N   K   P   P   S   P   A   S   I   R   P   E   M   *
```

##### Ap-GPR25-LgBiT in pTRE3G-BI vector coexpressing with SmBiT-ARRB2

```
1    GCT AGC ATG GCC CCT GAA GCC ACC GAG GAA CCT GAG GCT AGC GCC GAT TAC GAG TAC CCT GCT GGC GGC ACC GCC
     CGA TCG TAC CGG GGA CTT CGG TGG CTC CTT GGA CTC CGA TCG CGG CTA ATG CTC ATG GGA CGA CCG CCG TGG CGG
             M   A   P   E   A   T   E   E   P   E   A   S   A   D   Y   E   Y   P   A   G   G   T   A

76   CCA GGA TCT CCT GAG CAG CTG TTC ACC ACA GTG GTC ATC CCT GTG CTG TAC AGC TTC ATC TTC CTG CTA GGC TTC
     GGT CCT AGA GGA CTC GTC GAC AAG TGG TGT CAC CAG TAG GGA CAC GAC ATG TCG AAG TAG AAG GAC GAT CCG AAG
     P   G   S   P   E   Q   L   F   T   T   V   V   I   P   V   L   Y   S   F   I   F   L   L   G   F

151  GCC GGA AAC CTG TTC GTG ATC GTC CTG CTC GCT AAG AGA TCC GGC GGA AGA CGC GCC GTG GAC ACC TTC GTG CTG
     CGG CCT TTG GAC AAG CAC TAG CAG GAC GAG CGA TTC TCT AGG CCG CCT TCT GCG CGG CAC CTG TGG AAG CAC GAC
     A   G   N   L   F   V   I   V   L   L   A   K   R   S   G   G   R   R   A   V   D   T   F   V   L

226  AAT CTG GCC GTG GCC GAC GTG GTG TTC GTG TGC ACC CTG CCA TTC TGG GTT GCA GCT GGC GCC AGA GGC AAC CGG
     TTA GAC CGG CAC CGG CTG CAC CAC AAG CAC ACG TGG GAC GGT AAG ACC CAA CGT CGA CCG CGG TCT CCG TTG GCC
     N   L   A   V   A   D   V   V   F   V   C   T   L   P   F   W   V   A   A   G   A   R   G   N   R

301  TGG CTG CTC GGA GAA GGC CTG TGT AAA GCC TCT AGC TAC GCC ATC AGC GTG AAT AGA TGC TCT AGC ATC CTG TTC
     ACC GAC GAG CCT CTT CCG GAC ACA TTT CGG AGA TCG ATG CGG TAG TCG CAC TTA TCT ACG AGA TCG TAG GAC AAG
     W   L   L   G   E   G   L   C   K   A   S   S   Y   A   I   S   V   N   R   C   S   S   I   L   F

376  CTC ACC GCC CTG AGC GTG GAA CGG TAC CTG GTG ATT TGG AAG GTG CTT GAT ACC AGA AGG GGC GGC TCT AGA CGG
     GAG TGG CGG GAC TCG CAC CTT GCC ATG GAC CAC TAA ACC TTC CAC GAA CTA TGG TCT TCC CCG CCG AGA TCT GCC
     L   T   A   L   S   V   E   R   Y   L   V   I   W   K   V   L   D   T   R   R   G   G   S   R   R

451  CAC GTG CCT CTG GCC TGT GGC CTG ATC TGG GCC GTG TCC CTG CTG CTG GGA GCC CCT GCC CTG TGG TAC CGG AGA
     GTG CAC GGA GAC CGG ACA CCG GAC TAG ACC CGG CAC AGG GAC GAC GAC CCT CGG GGA CGG GAC ACC ATG GCC TCT
     H   V   P   L   A   C   G   L   I   W   A   V   S   L   L   L   G   A   P   A   L   W   Y   R   R

526  CTG GAC GGC GAT GAC TGC TGG GAC GAG GAT GGC GAG GAC TTC AGC CTG GCC ATG GTG TTC CTG ACC TTT CTG CTG
     GAC CTG CCG CTA CTG ACG ACC CTG CTC CTA CCG CTC CTG AAG TCG GAC CGG TAC CAC AAG GAC TGG AAA GAC GAC
     L   D   G   D   D   C   W   D   E   D   G   E   D   F   S   L   A   M   V   F   L   T   F   L   L

601  CCA CTG GGC GTG ATC TCT TTC TGC TAC TGT TCC ATC TAC TGC AGA CTG CAG AGA CAC GTG CGG CTG GGC AGA GGC
     GGT GAC CCG CAC TAG AGA AAG ACG ATG ACA AGG TAG ATG ACG TCT GAC GTC TCT GTG CAC GCC GAC CCG TCT CCG
     P   L   G   V   I   S   F   C   Y   C   S   I   Y   C   R   L   Q   R   H   V   R   L   G   R   G

676  GTG CGG CGG AGC CAC AGA GTG ATC GTG GCT ATC GTT GCC GCC TTT ATC TGC AGC TGG CTG CCC CTG AAC GCC TGC
     CAC GCC GCC TCG GTG TCT CAC TAG CAC CGA TAG CAA CGG CGG AAA TAG ACG TCG ACC GAC GGG GAC TTG CGG ACG
     V   R   R   S   H   R   V   I   V   A   I   V   A   A   F   I   C   S   W   L   P   L   N   A   C

751  AAG GTG CTG CTG TTT CTG CTG GCC AAG GGC ACA CTG GCC CTG CCT GCC GGC CAG GAG GGC ACA CTG AGA TGG GTC
     TTC CAC GAC GAC AAA GAC GAC CGG TTC CCG TGT GAC CGG GAC GGA CGG CCG GTC CTC CCG TGT GAC TCT ACC CAG
     K   V   L   L   F   L   L   A   K   G   T   L   A   L   P   A   G   Q   E   G   T   L   R   W   V

826  GTG GCC ACA AGC ACC TGT CTG GCT TTT GCC AAC AGC TGC GTG AAC CCC GTG ATC TAT GCC CTG ATG GAC CCC AGA
     CAC CGG TGT TCG TGG ACA GAC CGA AAA CGG TTG TCG ACG CAC TTG GGG CAC TAG ATA CGG GAC TAC CTG GGG TCT
     V   A   T   S   T   C   L   A   F   A   N   S   C   V   N   P   V   I   Y   A   L   M   D   P   R

901  CTG CGG TGC CCC CCC GCC CTG GGC GCT AGA ACC CCT CCT AGC ACA ACC GCG AGC AGC CCC AGC CCT GGC TCC GGC
     GAC GCC ACG GGG GGG CGG GAC CCG CGA TCT TGG GGA GGA TCG TGT TGG CGC TCG TCG GGG TCG GGA CCG AGG CCG
     L   R   C   P   P   A   L   G   A   R   T   P   P   S   T   T   A   S   S   P   S   P   G   S   G

976  ACC AAG CTG CCT GCT GTG CCC AAC CCT AGA CCT GGA ACA CCA CCG GTC GGT ACC GGC GGA GGG TCT AGC AGT GGC
     TGG TTC GAC GGA CGA CAC GGG TTG GGA TCT GGA CCT TGT GGT GGC CAG CCA TGG CCG CCT CCC AGA TCG TCA CCG
     T   K   L   P   A   V   P   N   P   R   P   G   T   P   P   V   G   T   G   G   G   S   S   S   G

1051 GGT GGG ATG GTC TTC ACA CTC GAA GAT TTC GTT GGG GAC TGG GAA CAG ACA GCC GCC TAC AAC CTG GAC CAA GTC
     CCA CCC TAC CAG AAG TGT GAG CTT CTA AAG CAA CCC CTG ACC CTT GTC TGT CGG CGG ATG TTG GAC CTG GTT CAG
     G   G   M   V   F   T   L   E   D   F   V   G   D   W   E   Q   T   A   A   Y   N   L   D   Q   V

1126 CTT GAA CAG GGA GGT GTG TCC AGT TTG CTG CAG AAT CTC GCC GTG TCC GTA ACT CCG ATC CAA AGG ATT GTC CGG
     GAA CTT GTC CCT CCA CAC AGG TCA AAC GAC GTC TTA GAG CGG CAC AGG CAT TGA GGC TAG GTT TCC TAA CAG GCC
```

```
     L   E   Q   G   G   V   S   S   L   L   Q   N   L   A   V   S   V   T   P   I   Q   R   I   V   R

1201 AGC GGT GAA AAT GCC CTG AAG ATC GAC ATC CAT GTC ATC ATC CCG TAT GAA GGT CTG AGC GCC GAC CAA ATG GCC
     TCG CCA CTT TTA CGG GAC TTC TAG CTG TAG GTA CAG TAG TAG GGC ATA CTT CCA GAC TCG CGG CTG GTT TAC CGG
      S   G   E   N   A   L   K   I   D   I   H   V   I   I   P   Y   E   G   L   S   A   D   Q   M   A

1276 CAG ATC GAA GAG GTG TTT AAG GTG GTG TAC CCT GTG GAT GAT CAT CAC TTT AAG GTG ATC CTG CCC TAT GGC ACA
     GTC TAG CTT CTC CAC AAA TTC CAC CAC ATG GGA CAC CTA CTA GTA GTG AAA TTC CAC TAG GAC GGG ATA CCG TGT
      Q   I   E   E   V   F   K   V   V   Y   P   V   D   D   H   H   F   K   V   I   L   P   Y   G   T

1351 CTG GTA ATC GAC GGG GTT ACG CCG AAC ATG CTG AAC TAT TTC GGA CGG CCG TAT GAA GGC ATC GCC GTG TTC GAC
     GAC CAT TAG CTG CCC CAA TGC GGC TTG TAC GAC TTG ATA AAG CCT GCC GGC ATA CTT CCG TAG CGG CAC AAG CTG
      L   V   I   D   G   V   T   P   N   M   L   N   Y   F   G   R   P   Y   E   G   I   A   V   F   D

1426 GGC AAA AAG ATC ACT GTA ACA GGG ACC CTG TGG AAC GGC AAC AAA ATT ATC GAC GAG CGC CTG ATC ACC CCC GAC
     CCG TTT TTC TAG TGA CAT TGT CCC TGG GAC ACC TTG CCG TTG TTT TAA TAG CTG CTC GCG GAC TAG TGG GGG CTG
      G   K   K   I   T   V   T   G   T   L   W   N   G   N   K   I   I   D   E   R   L   I   T   P   D

1501 GGC TCC ATG CTG TTC CGA GTA ACC ATC AAC AGT TAA
     CCG AGG TAC GAC AAG GCT CAT TGG TAG TTG TCA ATT
      G   S   M   L   F   R   V   T   I   N   S   *
```

##### Gg-GPR25-LgBiT in pTRE3G-BI vector coexpressing with SmBiT-ARRB2

```
1    GCT AGC ATG GAG TAC GCG GAG GCG GAT TAC TAC TAC GGC GAA GCC GAA GAG GAG GAA GAG GGC AAC GGA ACA GCC
     CGA TCG TAC CTC ATG CGC CTC CGC CTA ATG ATG ATG CCG CTT CGG CTT CTC CTC CTT CTC CCG TTG CCT TGT CGG
              M   E   Y   A   E   A   D   Y   Y   Y   G   E   A   E   E   E   E   E   G   N   G   T   A

76   TGT GCT TGG CAG GCT GAT TGG GAG GCC TCT TTC GCT CTG TTG CCT GCC TTG TAC GCT CTG GTC TTT GTC TTG GGA
     ACA CGA ACC GTC CGA CTA ACC CTC CGG AGA AAG CGA GAC AAC GGA CGG AAC ATG CGA GAC CAG AAA CAG AAC CCT
      C   A   W   Q   A   D   W   E   A   S   F   A   L   L   P   A   L   Y   A   L   V   F   V   L   G

151  CTG GCT GGC AAC GCT TTG GTG CTG TTG ACC GTC TGG AGA GGT CCA AGA GCC AAG AGG CGG AGT GCT GAT GCC TAC
     GAC CGA CCG TTG CGA AAC CAC GAC AAC TGG CAG ACC TCT CCA GGT TCT CGG TTC TCC GCC TCA CGA CTA CGG ATG
      L   A   G   N   A   L   V   L   L   T   V   W   R   G   P   R   A   K   R   R   S   A   D   A   Y

226  ATC GGC AAC TTG GCT CTG GCT GAC CTG GCT TTC GTG GCT ACA CTG CCT CTG TGG GCT GCC TAC ACA GCT CTG AGG
     TAG CCG TTG AAC CGA GAC CGA CTG GAC CGA AAG CAC CGA TGT GAC GGA GAC ACC CGA CGG ATG TGT CGA GAC TCC
      I   G   N   L   A   L   A   D   L   A   F   V   A   T   L   P   L   W   A   A   Y   T   A   L   R

301  TTC CAC TGG CCT TTC GGA GCA GCC TTG TGC AAG CTG AGC AGC TAC CTG GTG CTG CTG AAC ATG TTC GCC AGC GCT
     AAG GTG ACC GGA AAG CCT CGT CGG AAC ACG TTC GAC TCG TCG ATG GAC CAC GAC GAC TTG TAC AAG CGG TCG CGA
      F   H   W   P   F   G   A   A   L   C   K   L   S   S   Y   L   V   L   L   N   M   F   A   S   A

376  TTC TGT CTG GGA GGC TTG AGC GCT GAG AGG TAC AGA GCC GTG CTG AGA GCT GCT CCA CCA CCT CCA AGA GCT GCC
     AAG ACA GAC CCT CCG AAC TCG CGA CTC TCC ATG TCT CGG CAC GAC TCT CGA CGA GGT GGT GGA GGT TCT CGA CGG
      F   C   L   G   G   L   S   A   E   R   Y   R   A   V   L   R   A   A   P   P   P   P   R   A   A

451  TCT GCT AGG CTG AGA AGG CCT GCT GCT CTG GGT CCA TTG GCT GCA CTG TGG GCC GCC GCA GCA GCA GCT GCA TTG
     AGA CGA TCC GAC TCT TCC GGA CGA CGA GAC CCA GGT AAC CGA CGT GAC ACC CGG CGG CGT CGT CGT CGA CGT AAC
      S   A   R   L   R   R   P   A   A   L   G   P   L   A   A   L   W   A   A   A   A   A   A   A   L

526  CCT GCT CTT CTG CTG AGA GAG GCC AGA AGA GAC GCC AGA AAC AGA ACA CTG TGT GAC CTG AGA CTG GGT GGT GAA
     GGA CGA GAA GAC GAC TCT CTC CGG TCT TCT CTG CGG TCT TTG TCT TGT GAC ACA CTG GAC TCT GAC CCA CCA CTT
      P   A   L   L   L   R   E   A   R   R   D   A   R   N   R   T   L   C   D   L   R   L   G   G   E

601  GGT GGC GGA GCT GAA AGG GCC GCA GCT GCC TTG TCT CTG GGC ACC ACC GCT TTG GGT TTC GCA GCT CCA CTG CTG
     CCA CCG CCT CGA CTT TCC CGG CGT CGA CGG AAC AGA GAC CCG TGG TGG CGA AAC CCA AAG CGT CGA GGT GAC GAC
      G   G   G   A   E   R   A   A   A   A   L   S   L   G   T   T   A   L   G   F   A   A   P   L   L

676  CTG ATG GCT GTC TGC TAC TGC TGC GTC GGA TCT GCT GTG AGA CGA CAT CTG AGG CCT AGA AGG GCT GAG GCT GCT
     GAC TAC CGA CAG ACG ATG ACG ACG CAG CCT AGA CGA CAC TCT GCT GTA GAC TCC GGA TCT TCC CGA CTC CGA CGA
      L   M   A   V   C   Y   C   C   V   G   S   A   V   R   R   H   L   R   P   R   R   A   E   A   A

751  GCC AGA AGA AGG CTG CTG AGA CTG ATC GCT GCA CTT GTG GGA GTG TTC GCA GGC TGT TGG CTG CCT TTC CAC CTG
     CGG TCT TCT TCC GAC GAC TCT GAC TAG CGA CGT GAA CAC CCT CAC AAG CGT CCG ACA ACC GAC GGA AAG GTG GAC
      A   R   R   R   L   L   R   L   I   A   A   L   V   G   V   F   A   G   C   W   L   P   F   H   L

826  CTG AAG AGC CTG TTC GTG CTG GCT GCC GCT GGT CTG TTG GAG CTG CCA TGT GCT CTG CTG GGT CTG ATC AGC AGG
     GAC TTC TCG GAC AAG CAC GAC CGA CGG CGA CCA GAC AAC CTC GAC GGT ACA CGA GAC GAC CCA GAC TAG TCG TCC
```

```
      L   K   S   L   F   V   L   A   A   A   G   L   L   E   L   P   C   A   L   L   G   L   I   S   R

901  CTG CAC CCT TAC GCC ACA TGT CTG GCC TAC CTG AAC AGC TGC CTG AAC CCT CTG CTG TAC GCC TTT CTG GAC GGC
     GAC GTG GGA ATG CGG TGT ACA GAC CGG ATG GAC TTG TCG ACG GAC TTG GGA GAC GAC ATG CGG AAA GAC CTG CCG
      L   H   P   Y   A   T   C   L   A   Y   L   N   S   C   L   N   P   L   L   Y   A   F   L   D   G

976  AGA TTC AGG GCT CAG TGC AGA GCC TTG CTG GGA CTG AGA GGT GCT AGA CCT CCA CCT GCC GCT TCC AGC ACA CCT
     TCT AAG TCC CGA GTC ACG TCT CGG AAC GAC CCT GAC TCT CCA CGA TCT GGA GGT GGA CGG CGA AGG TCG TGT GGA
      R   F   R   A   Q   C   R   A   L   L   G   L   R   G   A   R   P   P   P   A   A   S   S   T   P

1051 AGC GCT CCA ACA CAG AGA AGC GAG TTG CCT TCC TCG GGG GGC ACC AAG GTG CCA CCG GTC GGT ACC GGC GGA GGG
     TCG CGA GGT TGT GTC TCT TCG CTC AAC GGA AGG AGC CCC CCG TGG TTC CAC GGT GGC CAG CCA TGG CCG CCT CCC
      S   A   P   T   Q   R   S   E   L   P   S   S   G   G   T   K   V   P   P   V   G   T   G   G   G

1126 TCT AGC AGT GGC GGT GGG ATG GTC TTC ACA CTC GAA GAT TTC GTT GGG GAC TGG GAA CAG ACA GCC GCC TAC AAC
     AGA TCG TCA CCG CCA CCC TAC CAG AAG TGT GAG CTT CTA AAG CAA CCC CTG ACC CTT GTC TGT CGG CGG ATG TTG
      S   S   S   G   G   G   M   V   F   T   L   E   D   F   V   G   D   W   E   Q   T   A   A   Y   N

1201 CTG GAC CAA GTC CTT GAA CAG GGA GGT GTG TCC AGT TTG CTG CAG AAT CTC GCC GTG TCC GTA ACT CCG ATC CAA
     GAC CTG GTT CAG GAA CTT GTC CCT CCA CAC AGG TCA AAC GAC GTC TTA GAG CGG CAC AGG CAT TGA GGC TAG GTT
      L   D   Q   V   L   E   Q   G   G   V   S   S   L   L   Q   N   L   A   V   S   V   T   P   I   Q

1276 AGG ATT GTC CGG AGC GGT GAA AAT GCC CTG AAG ATC GAC ATC CAT GTC ATC ATC CCG TAT GAA GGT CTG AGC GCC
     TCC TAA CAG GCC TCG CCA CTT TTA CGG GAC TTC TAG CTG TAG GTA CAG TAG TAG GGC ATA CTT CCA GAC TCG CGG
      R   I   V   R   S   G   E   N   A   L   K   I   D   I   H   V   I   I   P   Y   E   G   L   S   A

1351 GAC CAA ATG GCC CAG ATC GAA GAG GTG TTT AAG GTG GTG TAC CCT GTG GAT GAT CAT CAC TTT AAG GTG ATC CTG
     CTG GTT TAC CGG GTC TAG CTT CTC CAC AAA TTC CAC CAC ATG GGA CAC CTA CTA GTA GTG AAA TTC CAC TAG GAC
      D   Q   M   A   Q   I   E   E   V   F   K   V   V   Y   P   V   D   D   H   H   F   K   V   I   L

1426 CCC TAT GGC ACA CTG GTA ATC GAC GGG GTT ACG CCG AAC ATG CTG AAC TAT TTC GGA CGG CCG TAT GAA GGC ATC
     GGG ATA CCG TGT GAC CAT TAG CTG CCC CAA TGC GGC TTG TAC GAC TTG ATA AAG CCT GCC GGC ATA CTT CCG TAG
      P   Y   G   T   L   V   I   D   G   V   T   P   N   M   L   N   Y   F   G   R   P   Y   E   G   I

1501 GCC GTG TTC GAC GGC AAA AAG ATC ACT GTA ACA GGG ACC CTG TGG AAC GGC AAC AAA ATT ATC GAC GAG CGC CTG
     CGG CAC AAG CTG CCG TTT TTC TAG TGA CAT TGT CCC TGG GAC ACC TTG CCG TTG TTT TAA TAG CTG CTC GCG GAC
      A   V   F   D   G   K   K   I   T   V   T   G   T   L   W   N   G   N   K   I   I   D   E   R   L

1576 ATC ACC CCC GAC GGC TCC ATG CTG TTC CGA GTA ACC ATC AAC AGT TAA
     TAG TGG GGG CTG CCG AGG TAC GAC AAG GCT CAT TGG TAG TTG TCA ATT
      I   T   P   D   G   S   M   L   F   R   V   T   I   N   S   *
```

**Fig. S2.** The nucleotide and amino acid sequence of Cc-GPR25, Ap-GPR25, and Gg-GPR25 in different expression constructs. The amino acid sequence of GPR25 is shown in red, that of LgBiT is shown in blue, and the signal peptide of sLgBiT is shaded.

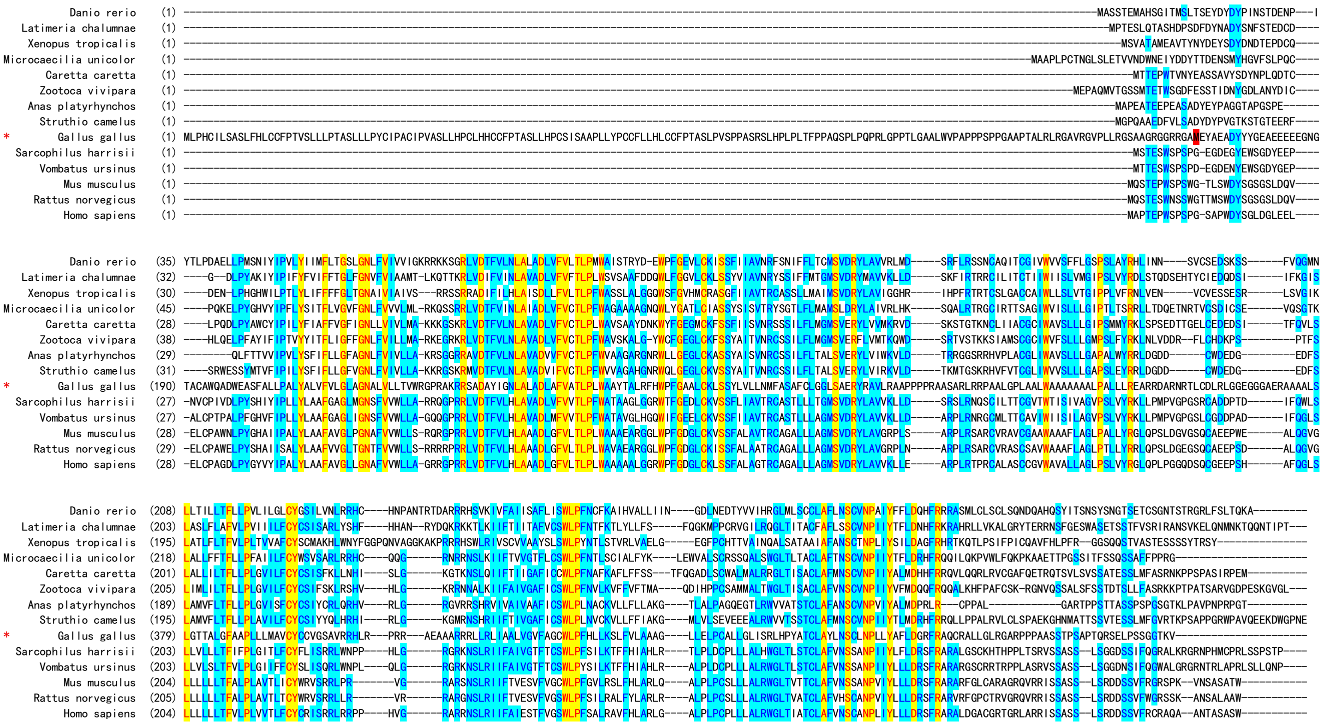

**Fig. S3.** Amino acid sequence alignment of chicken GPR25 with other GPR25 orthologs from mammals, birds, reptiles, amphibians, and fishes. The chicken GPR25 (Gg-GPR25) is indicated by a red asterisk, and the second methionine residue is highlighted in red.

**A:** Gene synteny and architecture

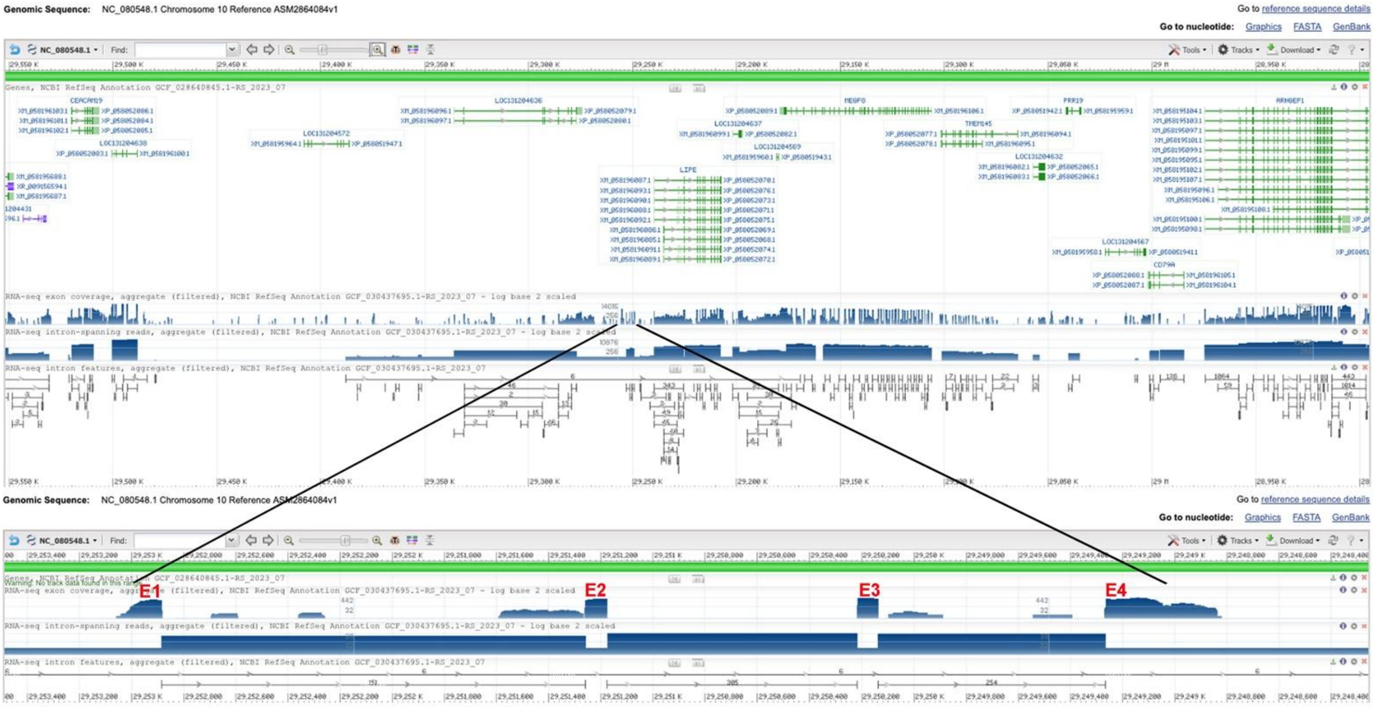


**B:** Sequence of genomic DNA (exons in red and highlighted in yellow, coding region underlined)

gagagaaaaaatggaaaaaaaaaaagaaagaaagaaagaaacaccacacatgggttggttcaaatacgaaaagttattagcggaagtagcggccattgaccgagagagccgaggaccttgctagcaaaatggaccgattcttcatctatctgatgttagtgctct
tggttggcttggttggctgctcagaaagaggtaagaaggttcggaaagggttaaatggaagatggttgcatgagaagaaaggagaggaaggggaggaagatccctggatccatgccagacaagcatttattccctcaacaccattcagctggtcattaaggtg
gaagctttcaagaagagattgggctgccctatggcagaaatggtgtaaggcagaggtcttcaaacttggcagctttaaggcttgtggggttggactggatgacctaacagggtcccttccaactcttgttaatctgtttaaaaataaactatgctggctggagaattct
gggagttgaagtccacaaggcttaaagctgccaagtttgaagacttctggtgtagggtctcctgcttgggcagaggaaggggttggactggatgacctaacagggtcccttccaactcttgttaatctgtttaaaaataaactatgctggctggagaattctgggagt
tgaagtccacaaggcttaaagctgccaagtttgaagacttctggtgtagggtctcctgcttgggcaggggaaggggttggactggatgacctaacagggtcccttccaactcttgttaatctgtttaaaaataaactatgctggctggaaaattctgggagttgaagt
ccacaaggcttaaagctgccaagtttgaagacttctggtgtagggtctcctgcttgggcaggggaaggggttggactggatgacctaacaggatcccttccaactcttattttaatctgtttaaaaataaactatgctggctggagaattctgggagttgaagtccaca
aggcttaaagctgccaagtttgaagacctctggtgtagggtctcctgcttgggcaggggaaggggttggactggatgacctacagggtcccttccaactgtaagggagggagggagggaaggggg agggagaggaggaagggagggagggaggagga
gaaggaaaggaaggaaaagaggggaagaaaggagaagaggggaaggaaagaagaaaagaggaaggaaggaaaagaggggaagggagggagaaagagaagaaaggaaggagaagaggggaaggaaaaagggaaggaaggaagaaggagg
gaaggaaggaaaagaggggaaggaaggaaggaaagaaaaagggaaggaaggagaagaggggaaaggaagaagggaaggaaggaaggaaaaggggaaggaaggaaggaaaagaggggagggagagggagggaaggaaggaagaaaggg
agggagggagggagaaagattggaacagtgaatggatccatgtacctcagctagaatggaaactctttttaatttgatagttggaggtaaaagatttggggagcacactcctccctagttctctcaagtcctccatttctcgggtcctcttttttcctcacctgttcccagt
gggatggtgtgaaagaaaggaggacgaactgggaggatcacccccaaagaagggcacccacctcccctacctcggttggaaggaagcagggagagttggtcatcctttctccaggatcgccaagtccacgacccctcgtagccccttccaacctgctcccg
tctttctccttttgttgtgggggggggggatgcaggtggaagctacggcgaaggaaaagagcgttcgctgagcaggtcgaaacgaagcaagttgtggcatccgtcctgcacttgtccaggtgagtcccgtctcttgacttccatgggggggggggagttgggag
aaagtttgcctgagggtttactttagctccgtgatggtgaacctacggcacgcgtgctacaggtggcacacagagccctttcaacgggcacgccagcccagctgcactgcacatgcgcacacctgcctcccaccagccagctggtctttgggcctatgccgtgc
atgcacaggggtgggcgcatacatacatggggggggggggcaggggataggcatgggggcgctcacattgcatttgggggggttcttgcgcgccccgttttgggactggaaagcctcttgcagcaacctggaagccaaaatggggaacgggagtggg
gggaaatgaggaaatgctgggttctgacgatcccagctgagttgcctgatcgccagaaccttttaaaagcatttttacaacctcttcaaccaaagagtttgtagaaaaaatgcttttaaaaggctctgacgagaaccgctgagctgcacgatcatcagaggcttttttt
tttttacttttaaaagcatttttcagccgaagaaaaaatgcttttaaaagtaaaacctctgataatcatgtggatcagctgggcattagggggggggcaggtatttttcctaccagttctccgaaccacctgccgtcatcgctaccggatcaggcgatccggtccgatct
gggagcatttcaccccctgcttgtggggtctgggcaactgaatggagctagcagagcgttttactgaccggatgcccttcctgttgccaatgcagagttttgttcagttatattctcattatgtccagagagagaaatatctgcctctacttaggctcgaactcacggcct
cttgattgtgaggcgagagctctacctctaggccaccgcaccatccctgacgtccttctaatgatggttcccaattgcttttagatttaaatgaactccaacatccaaagtattataaacatgcctgtccctgtgatggattcagcttacatagtttgtcaggtaagaaaat
gctcctgggtgtaactcgtgtgggaggtaaaggaagatgggtccgtgggtgtagcatcttcttaggcgtggacaaaaggaagaaatcagaaaccacccatcggaacttttctagaatactggctcgttttaattttgtccgcccattaaaatcacttttcgttgtgcag
gatactacagatagtagtatcagcaattgaacccagaattataaaacccaaaatataggactcaattgtggtcgtaagtagaggattactgttgtgcctcggccgccctcctctccacagccgggcccctcccatctcctgctatctcagccagagactgatagtga
agaagaatggcctggcatgcctccagcccccagccctggcaccatgcacagacagactgagcaaaaaacctccctcctatagcatgtgagcatgaagccagccacgagctagagttgccggcagctgagcaggaggatgaaaagacacagtggacagat
ccccgcttccggagaatggagaggcgacgtcagcaaaaggaagggaggggcaggcctggataagtgctgagtcatggagccacaccccatggcctatataaaggatctgctttctggcattccttgagtcaggcaaagtctcatttggttgctgaagtcacac
cttggattcctacctgccctgagaactcttgacaggactttggcaaagctgcagaggcttcgtggccacgcttgatacagacttcccagacccggccgtcagaggaggagtggggcacgacaattaccttctacctttcagtcttcaactgaccgattaaaagctg
cattattttattttattttttttctctgttccctttcagagcaccgcagtcctcgccgccatcatcatcatcatcgcccgaacaagcgacctgtgctccaaaagacaaaatgcaggaggtttttgagacactgcgcttttaaggggatccacgttcccttatagtttatcccg
gttcttggcaagacccgaagagatttattacccatgacgcaatattgattctttggttatgcacctttctcgtttgttttttttaaatatcattgtttgctctaggaaaacgttggttg

**C:** Sequence of cDNA (coding region in red and highlighted in yellow)

Gagagaaaaaatggaaaaaaaaaaagaaagaaagaaagaaacaccacacatgggttggttcaaatacgaaaagttattagcggaagtagcggccattgaccgagagagccgaggaccttgctagcaaaatggaccgattcttcatctatctgatgttagtgctc
ttggttggcttggttggctgctcagaaagaggtggaagctacggcgaaggaaaagagcgttcgctgagcaggtcgaaacgaagcaagttgtggcatccgtcctgcacttgtccagatttaaatgaactccaacatccaaagtattataaacatgcctgtccctgtg
atggattcagcttacatagtttgtcagagcaccgcagtcctcgccgccatcatcatcatcatcgcccgaacaagcgacctgtgctccaaaagacaaaatgcaggaggtttttgagacactgcgcttttaaggggatccacgttcccttatagtttatcccggttcttgg
caagacccgaagagatttattacccatgacgcaatattgattctttggttatgcacctttctcgtttgttttttttaaatatcattgtttgctctaggaaaacgttggttg

**D:** Amino acid sequence (signal peptide shaded)

MDRFFIYLMLVLLVGLVGCSERGGSYGEGKERSLSRSKRSKLWHPSCTCPDLNELQHPKYYKHACPCDGFSLHSLSEHRSPRRHHHHHRPNKRPVLQKTKCRRFLRHCAFKGIHVP
L

**Fig. S4.** Gene synteny and architecture (**A**), genomic DNA sequence (**B**), cDNA sequence (**C**), and encoded protein sequence (**D**) of the unannotated *cxcl17* gene in the NCBI reference genome of *Ahaetulla prasina*. The information was downloaded from the NCBI gene database (https://www.ncbi.nlm.nih.gov/gene).

**A:** Gene synteny and architecture

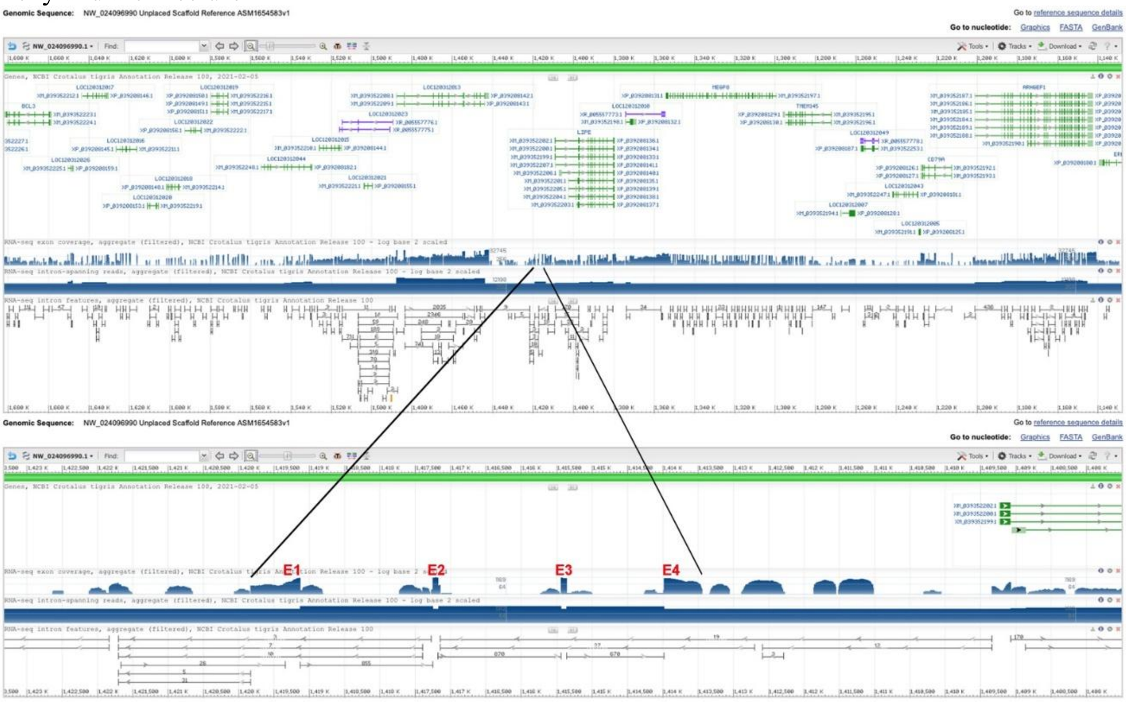


**B:** Sequence of genomic DNA (exons in red and highlighted in yellow, coding region underlined)

ctgatatgattttcttatttttccccaagcacttcccagaggtattagccgggaagaaaaaaaaaaatttttttaaataaaataaatacgtcggttggttaagtacaaaaggttattagccgaagcagtggccattgacggagagagagccaaggacgtcgctggcg
aaatggacagattctttttattttgatgtcagcgctcctcgttggcctggtctgcagctcagaaaaaggtaggagttggagagagttattgggaattgttgcacgggaagaaaggagaggaaagtgagggaggtctctggattggcaccagacaagaaatgtccc
ctcagcaacattaacctggtcattaaattaaccctttttcctctagatttacacaataccctgagccataaaatggctaggaagaaataacaactccctaagtttagtatattgtagaaagccatcatggtgggagtggagctggtccaacctgtgcttgaacaccaccc
aggtcttctcactgatctttgaccaaggatctgtagactgtccagaggtctgtagtgaagaggaaggaaggaaggaaggaaggaaggaaggaaggaaggaaggaaggaaggaaggaaggaaggaaggaaggaaggaaggaaggaaggaaggaaggaaga
agggagggagggagggaggggagggaagggagaggaaggagggaaagaaggaaaggaaagggagggagggagggagggaggaaggaaagagggaagggagggagggaggggagggagggagggaaagaaagaagggagggagaga
aagaaggaagggaggggaggggagggagggaaagaaaaaagggaggaggagggagggaggaaggaaggaaggagggagggaggaaaagaagggagggagggagaaagggagagaaaggaggagggagggaaagaagaaaagggagg
gaggaaggaaagagggagggaggggagggagagaaagaaggaagggaggagggaggaaagaaggaaaggaaagaaggaagggaaagaaggggagggagagggagagagggaaagaaggagagggagggaaagaaggaaagggagg
gagagagggagggaaggaaggaaggagggaggggagggaggagagaaagaaaagggaggagggagggaaagggaaaggaaagaagggagggagggaaagaaggaaggaggagaaagggagagagggaaagaaaggaggagggaggg
gaaggaagggaggggagaagggaggatgaaggaagggagagggagggaggggagggagaagaaggaagggaggagggagggaaagaaggaaaggaaagaagggagggagggaaagaagggaaggagggagaaagggagagagggaaa
gaaaggaggagggagggaaagaaggaaagggagggagagagggaggaatgaaggaaggaaagagggagggaggggaggggagggagagaatgaaggaagggaggagggagggaaagaagaaggaaggaaggaaggaaggaaggaagg
aaggatggatggaagaaggaaggaaggaaggaaggatggaagaaggaaggaaggaaggaaggaaggaaggaaggaaggaaggaaggaaggaaggaaggaaggaaggaaggaaggaaggaaggaaggaaggaaggaaggaagggacagattggaaca
gtggggtccttggggggttccctgaccttgcttgctttcttgcagacgtttcattacccaaactagttaacatcaccagtgctagattgaaacagtgaatgaatctacgtacctcacctagaatggaaactggttttaatttgatagctggaggtaaaagatttaggctgca
ctcctccgtggttctctcaggtcccccttttctcacctgctccccatgggatgatgtgaaaggaagagggacgaactgggaaaaccacccccaaagaaaggcacccccccctcccctacctcggttgaaggcagtggagaccattggcgatccttcctccaggat
catcaagtccacaagagcttcttgaacctcatcggtagaatccttcatccctcgtagccccttctgagctgctctcacctttcgccttttgggcttaggtggaaggtccgaccaagcgagtttgctgagcaggtcgaaacgaagcaggctggagcctcggtcctgcc
cttgtccaggtgagtcccatctcttgatctccatcggggagctggaagaaaatttgcctgcaagtttactttgggttcttctggcatccaggatcacgtggtacaagtgcaaagaagagcaactaagatgattaaaggcctggagactaaaacataggaagaacgg
ttgcaggaaccggatatggccagtctagagaaaagaaggacaggaggggggggacatgatagcagtgatccaatatttgaggggctgccacaagacggagggggggggggtgaacttattttccaaatcacctgaaggcaggataagaagcaatggttggaa
agtaatcaaggagagaagcagcctggaattaaggagaaacttcctaacagggaggacaattaaccagtggaacagctggccaccagaaagcgtgggcgctccatcattggagatttttaagaagagactagacagtcacttgtctgaaatggcataggttctct
tgcttgagcaggggggtggactagaagacttccaaggtcccattctattctattctattctattctattctattctattctattctattctattctattctattctattcttctacttcattccattattctatcctatcctatcctattctacttcattccattattctattctattctattctatc
ctagcctagcctagcctagcctattcttctacttcattcctattctattctattctattctattctattctattctattctattctattctattctattctattattctattctattctattctattctattctttattctattctattctattctattctttattctattctatactattctattctattc
tttattctattctattctattctattctattcttctacttcattccattattctatcctatcctatcctattctacttcattccattattctattctattctatcatagcctagcctagcctagcctattcttctacttcattccattattctattctattttattctattctattttattctattctattctat
tctattctattctttattctattctattctattctattctttattctattctattctattctattctattctattctttattctattctattctattccattctatattctattctattctattctattctattgtaaaggactggggtcacatgatagcagtgttctgatatttgaggggctgcc
acaaagaagaaggggtcaacttattttcgaaagtaccagaatgcaggacaagaagcaatggatgaaaactaatcaaagagagaagcaacctaggaactaaggagaaatttcttgacaggaagaactattaatcagtggaacaacctgcctccagaatatcact
agaggtttttaagaagagactggacagccacttgtctgaaatggcataggttctcctgcttgactagaagacctccaaggtcccttccagctctattttgattgattgattgattgattatgggttgtctctttgggtcttttctctctgttccaggtgtctgacttaaaaacag
acaagccttgttgttatccttgttactcttttgaggattgcatttcttcaaggatggcttatgggtcaggaggaaaaaccttgcgatctttttctgatgccccatgttttcctggcgtccttctaatgctggttcctaatcatttttagatttaagtgaactgcaatacccccacaa
agacaaacatccttgcccctgtgatggattcaccttaagtagtttgtcaggtaagaaaacactcccgggtgtaactcctgtggaaggtaaatgaagaagggtgtgtgggtgtgggtggctgttgtgtagcctaagcatcttattaggcctgggtgagaaggaaaact
cagaaatcacccattgggatttttctaccatccccatttattttcattttgtccacccattaaaatcacttcttactttgcaggacactacaggtaatcctcgacttacgatcacagtggatcccataattgctgttgctaagtgaaacattgggtgagtcttgccccccatttta
cgacacagttaaatgaagttggtaacaaagctgttaagtgaaattggctaccccattgacagaaggttgcacaaagggtcccaggatgctgcaaccgtcataaatatgagtcaaaaattggaaaaaaaaatggacttattaccgtttttcatactcacaacccttaggg
caggggtaggcaaccttggctcttttatgactcgtggacttcaactcccagaattcctgagccagcatggctgagcttggaaacctctccctttcccctcagcgcggctggctcaggaattctgggagttgaagtccacaagtcataaaagagcccaggttgccta
ccctaggattagactaacatcatctacatcatcatctaacatcatcagtaaaataataaaataaactagacatatatgaatgtatgtatgtatgtatctacctacctactatctatctatctatctatctatctatctatctatctatctatctatctatctatctatctatctatct
atcacagggctaaacagaaaaactctgacactgacatcccctaaacccaggggtaggcaaccttgggtcttatatgactggtggacttcaactcccagaattcctgagccagcatggctcaacatggaaacctctccctttcccctcagcgcggctggctcagga
attctgggagttgaagtccacaagtcataaaagagccaaggttgcctacccctgccttagggcatccctgtggtcacgtgatcaaaattcagaggctcggcaactggcttgtgcttatgacagttgcagtgccctggggtccatgggatcactttttatgatcttctga
ccaatgggggagccagattcacttaacaaccatgttactaacttaaaagcaatgattcacttaacaactgtggcaagaaaggtcgtaaaatggggcaaagctcacttaaccactctctcacttagcaacggaaatgttaggctcaattgtggtttgtatgtcgaggac
tacctctatacctttggtgttcaattgaccgattaaaagattctttttttttctgtttctttttagaacactccagtcatcatcatcatcctcctcgcaagaacaagcaacttgtgcttcataagaaaaaatgcaggagatttttgagacactgcgcttttaaagggatccatatt
cccttatagcttctcccaagggaattatccgtgacgccatattgattctttgcttatgtacctttctctttttttaaatatcattgtttgctctagcaaactgcttgctcggttggacataaagctccaggctcgaggcctttgtggaagtgaacggcctgtaaacaactgtatta
aataataatatgttactatatgtggggcaagctgcccacgccgcatattttgcaatcccttgtaaaaccgggagaggtgcagaaaataaggggagtttcaatgggattgcagtctttagcctacctatctcagggattgcc

**C:** Sequence of cDNA (coding region in red and highlighted in yellow)

Agtacaaaaggttattagccgaagcagtggccattgacggagagagagccaaggacgtcgctggcgaaatggacagattctttttattttgatgtcagcgctcctcgttggcctggtctgcagctcagaaaaaggtggaaggtccgaccaagcgagtttgctg
agcaggtcgaaacgaagcaggctggagcctcggtcctgcccttgtccagatttaagtgaactgcaatacccccacaaagacaaacatccttgcccctgtgatggattcaccttaagtagtttgtcagaacactccagtcatcatcatcatcctcctcgcaagaaca
agcaacttgtgcttcataagaaaaaatgcaggagatttttgagacactgcgcttttaaagggatccatattcccttatagcttctcccaagggaattatccgtgacgccatattgattctttgcttatgtacctttctctttttttaaatatcattgtttgctc

**D:** Amino acid sequence (signal peptide shaded)

MDRFFLFLMSALLVGLVCSSEKGGRSDQASLLSRSKRSRLEPRSCPCPDLSELQYPHKDKHPCPCDGFTLSSLSEHSSHHHHPPRKNKQLVLHKKKCRRFLRHCAFKGIHIPL

**Fig. S5.** Gene synteny and architecture (**A**), genomic DNA sequence (**B**), cDNA sequence (**C**), and encoded protein sequence (**D**) of the unannotated *cxcl17* gene in the NCBI reference genome of *Crotalus tigris* (Tiger rattlesnake). The information was downloaded from the NCBI gene database (https://www.ncbi.nlm.nih.gov/gene).

**A:** Gene synteny and architecture

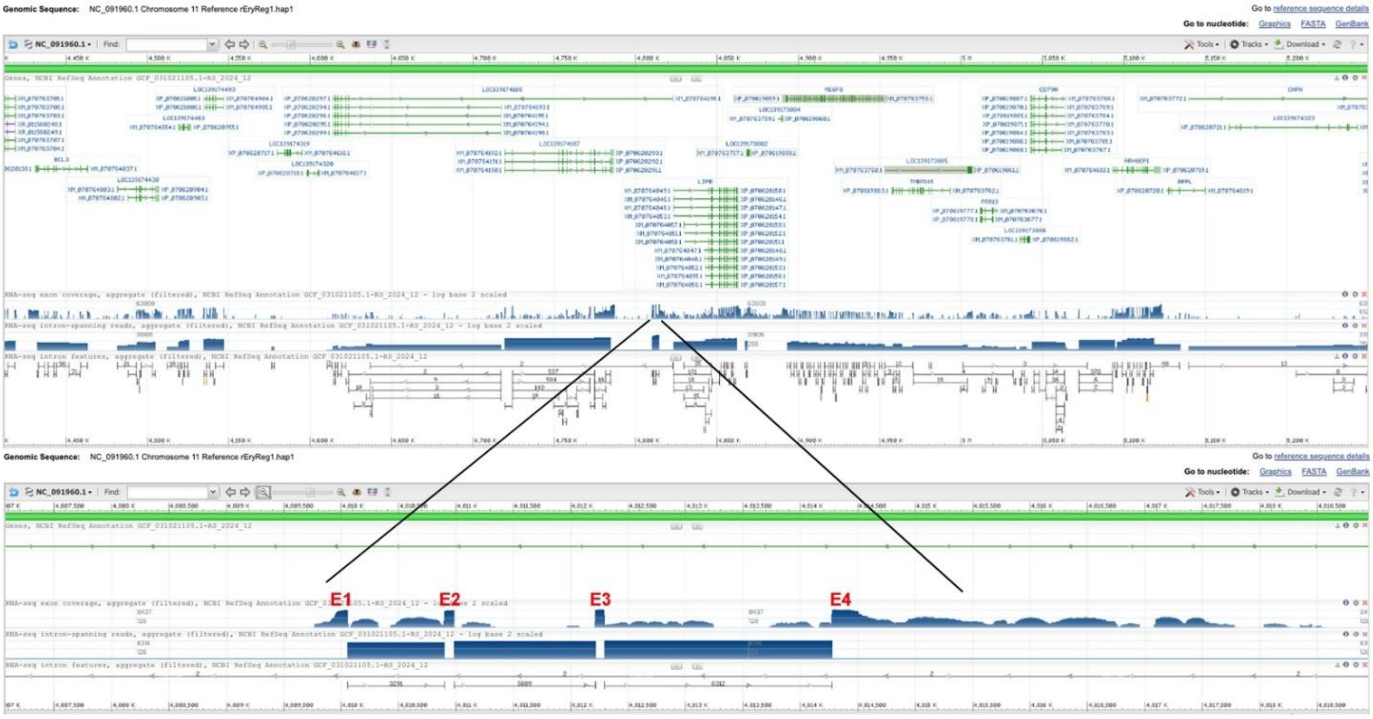


**B:** Sequence of genomic DNA (exons in red and highlighted in yellow, coding region underlined)

ttgcacagtggattggggactgccctgatttcatattgttataagagcctttcctggaaatattaactaactgggaggagagagagagaaagaaaatgcaaaagaaaatttaaataaataaatacaagttggtgaagtacaagtgcgaaaaagttattagcggaagc
ctgtggccattgacagagggcatcaaggacattgctatcaaaatgggccgattcgtcttctttctgatgttagcggtcctcggtggcttggtttgctgctcggaaggaggtaagagcgttattggagattatttcgttttgttttgttttgttttattttattttgtccagtacata
ttggtagtatacataggtagaacactgtttatatacataagatgggtactaatgagagggaaatattagggcaggaattgagcaaggtcaatagtggagggtctaagggtaaagttttcggggtttggtgacgaaactatagaggcaggtagtgagttccaggtatt
aaccactgattagggaagggaagaagagagggaagggagggagggagggaggaaggaaggaagatgggaaggaaggaagatgggaaggaagaagaaaggaaggaagaaggaaggaaggagggagggggaggagggagaaaaggaagggag
gaaagagaagaggggaaggaaagaaggaaagggaggaagagagggaaatgaaggaggaaggaaggaggaatggaagaagggagggaaggaaggaaggaagactggaacaatgaattgatccatatatctcacctaggacggaaactgattttgata
gctagaggtaaaagattttgggggcactgcccccccccccatagctctctcagctcctccttttcccaactgttcctaatgggatgatgtgaaaaggacaggaattgggaagatcacccccaaagaagggcacccacctccccaaccttggttgaagagaacggg
ggggggggggagttggcggtcctttctccaggatcgccaagtcccccatagcttcttgaatcacgccgttggcttctttcatccctcgtagccccttccaacctgcactcatctttctccttttgggggggggggcagatggcagtcacagtgaaggaaacgaagc
tttgctgagccggtcgaaacgaagcaagcagtggcctgagtcgtgcccttgtccaggtgagtctcacctcttgaccctcatcagggagttggaagaaaatgcacctgagggtttacttcaggccagcggttgtcaacctttctaatgtcgtgaccccttaacacagt
tccttgggttgtggtgacccccaaccataagtcgaacgccaatccttccaacagagcttaaagctgattggccggaagatcagagggacgcccccactgccaacacctgattggtcggattgtaaaaatacgttccaagcagtgaagggccaccgtcgccggc
acctaccagtacagtgatacctcgtcttacaaacgcctcgtcatacaaacttttcgagatacaaacccggggtttaagattttttgcctcttcttacaaactattttcaccttacaaacccaccgccgctgctgggatgccccgcctccggacttccgttgccagcaaa
gcacccgtttttgcactgctgggattcccctgaggctcccctccatgggaaaccccacctccagacttccgtgtttttgtgttgctgcaggggaatcccaacaggggaatccgagcagcgcaaaaatgggcgcttcgctggcaacggaagtccggagggaatg
cttcccagtgaggggagcctcaatgaaatcgcagcattgcaaaaacaaagaggtccagagatggggtttcgaggactttggtgtttttgtgatgctgcgatttcactgatgctcccgtcgctgggaaaccccacctccggacttccgttgcctgtgaagcgcccatt
tttgcgatgctgggattcccctgcagcatcacaaaaacacagaagtccagaggtggggtttcccatagaggggagcctcaggggattcccagcagtgcaaaaacgggcgcttcgactggcaaaaggggtgaatttggggcttgcacgcattaatcgcttttcca
ttgattcctatgggaaacattgtttcgtcttacaaacttttcaccttaagaacctcgtcccggaaccaattaagtttgtaagacaaggtatcactgtatttctacttggtgcttcccagagccggagtgggcgcacgcgtgcggctgctgtgcaattttgcctctgtgtatg
cgcccggagcaaaacctcacatgatttcagcgattttcacctatatatttgcttctttgcctgtgcaattaactcctaggacccctgacatccttctaatgatggttcccaattgcttttagatttaaatgcacttggacgtctaaagaaacacgaacatgcctgtccctgcg
atggattcagagtgcgtagtttgtcaggtaaataagaaaaatgctcctgggtgtaactcatgtgaaagttaaaggcggggttgggtctgtttgtttagcatcttcttaggcgtggaggaaaaggagaaatgagaaagcacccagcagggttttttctaatattctgcttc
atttttaattttgcccgcccattaaaataacttttattgtgcaggatactacaagtgtaactcatgtgaaagttaaagggagttgggtctgtgtgtttagcatcttcttgggtgtggaggaaaaggagaaatcagaaagcactgaacagagtttttctaatattctgtttcat
ttttaattttgcccgcccattaaaataacttttattgtgcaggatactacaagtgtaactcatgtgaaagttaaagggagttgggtctgtgtgcttagcatcttcttaggtgtggaggaaaaggaggaatcagaaagcacacagccggattctaatacaggggtacctc
tacctacgaacgcctctacttacaaacttttctagataagaaccgggtgttcaagatttttcccccctctcctcaagaaccattttccacttacaacttactttaaaacttaaagataaagagatgtaagatctgtggaaagttttatgaacggttggtaactaaaaataattg
agaagtattttcttttaattactttggaagaaatacatgatgattccattttaatttattaaagacccaggtgacgaaataaggattggtggcagtattggataaggagttaacacagagacgaaaggattaatattgagttaagatgttaagatgtttatgttttattacaaat
agaaaaagggaatataagggactagatttcataatattgttatctgtagatgaaatatatctcatgtgtagttaattttaattaacttagcaaagaagaggataaaaagtacatattgaaagtattattagcatttttttctctgtatttatttatttatttatctatctatctatctatct
atctatctatcatctatctatctatctatctatctatctatctatttatatgccgcccctctcagcagactcggggcggctaacaacagcaataaaacagcatataataataatccaatagtaaaaacagtttaaaacccattattataaaaaccagacatacatacatacgatat
accatgcataaaattgtagaggcctagaggaaagagtatctcaattcccccaagcctggcggcagaggtgggttttaagcagcttacgaaaggcaaggagggtgggggcaattctaatctctgggaggagttggttccagagggctggggccgccacagag
aaggctcttcccctgggtcccgccaagcgacattgtttagttgacgggacccggagaagacccactctgtgggacctaactggtcgctgggattcgtgcagcagaaggcggtccctgagataatgtttaatgtatcaaagaaatatcattaattattattttaagcatt
gattctgtgtttttattgctcatgttgttttattgtatgttggaagaaaaataaagacaaattttatacacacacaccctgccccccaaaaaagttaagtgagctttgccccattttatgaactttcttgccacggttgttaagtaaatcactgcaggtgaaaagttcgtaaca
cggttgtttaaacgaatctggctcccccactaaagcaaaaatatcgctgaaaatcggcgagatcttgccggcgtgcgcattatttattcataaagacatatttacttcatttatttacattgtactttttttcttaaattttttttcgtcctccttttcattgtaaaaaaaattggtcgcct
tacttcaatgtagggctcgattttggtcgttaagtggagggttgttttccaccaatcaattgaccgattaaaagattcgtaattttttttttccctgttctctttcaggacatcgcccacatgaccatcattttcacaagcgatacctgtcccgtagggcactatgcaagaagtt
tttgcgacactgcacttttcaggggatccacattcccttgtagctgatcctggttcttgacaagatccaaggggatttattatccgtgacgcaatattgattcttcagttatgcacctttctcggggggttttaaacattgtttattcgaggaaactgctgggttgtttggaaat
aaagctccgagtttgatgtctttgtgggagtgaatggtctgtattgaattgtgtggacaaatacagtggtacctctctttacgaacttaattcgttctgtgaccaggttcttaagtagaaaggtttgtaagaagaagcaatttttcccatatgaatcaatgtaaaagcaaata
atgtgtgcg

**C:** Sequence of cDNA (coding region in red and highlighted in yellow)

Agtgcgaaaaagttattagcggaagcctgtggccattgacagagggcatcaaggacattgctatcaaaatgggccgattcgtcttctttctgatgttagcggtcctcggtggcttggtttgctgctcggaaggagatggcagtcacagtgaaggaaacgaagcttt
gctgagccggtcgaaacgaagcaagcagtggcctgagtcgtgcccttgtccagatttaaatgcacttggacgtctaaagaaacacgaacatgcctgtccctgcgatggattcagagtgcgtagtttgtcaggacatcgcccacatgaccatcattttcacaagcg
atacctgtcccgtagggcactatgcaagaagtttttgcgacactgcacttttcaggggatccacattcccttgtagctgatcctggttcttgacaagatccaaggggatttattatccgtgacgcaatattgattcttcagttatg

**D:** Amino acid sequence (signal peptide shaded)

MGRFVFFLMLAVLGGLVCCSEGDGSHSEGNEALLSRSKRSKQWPESCPCPDLNALGRLKKHEHACPCDGFRVRSLSGHRPHDHHFHKRYLSRRALCKKFLRHCTFQGIHIPL

**Fig. S6.** Gene synteny and architecture (**A**), genomic DNA sequence (**B**), cDNA sequence (**C**), and encoded protein sequence (**D**) of the unannotated *cxcl17* gene in the NCBI reference genome of *Erythrolamprus reginae* (royal ground snake). The information was downloaded from the NCBI gene database (https://www.ncbi.nlm.nih.gov/gene).

#### A: Gene synteny and architecture

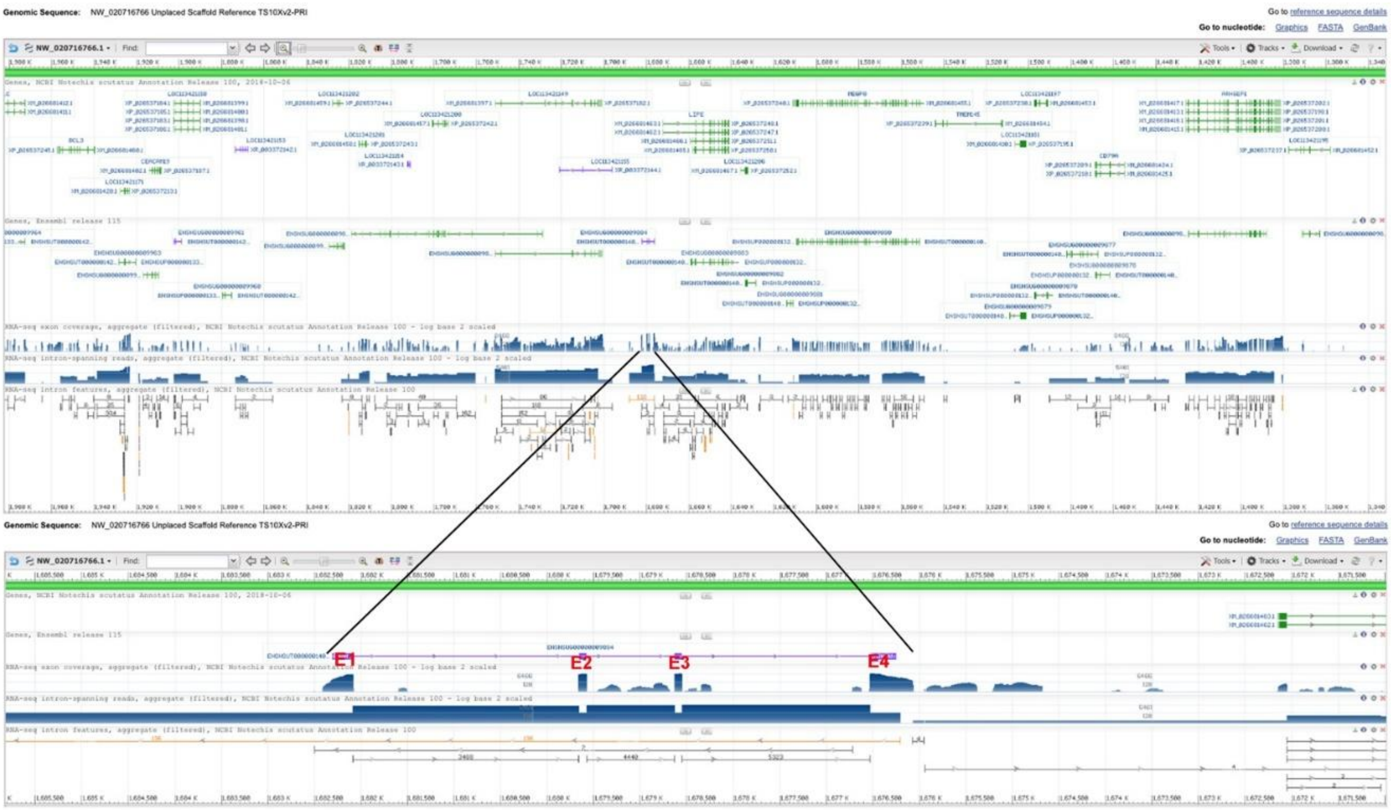


#### B: Sequence of genomic DNA (exons in red and highlighted in yellow, coding region underlined)

ggaaaaaaaaggtttgcacagtggcttggggactgccctgatatgatttcttatttctacacaaagcctttcccagaaatattacccagctgggaggaggggagaaaaaacaagttggtttgttcaagtaggaaaaatcagtagaagccgtggccgttgacggag
agacccaaggaacgccgctgtcgaaatggaccggttcttcttctttctggtgttaggggtccttgtcagcttggtttgctgctcagaaaaaggtaagaggattgggagagggggagggagggagggagggagggagggagggagggagggnnnnnnnn
nnnnnnnnnnnnnnnnnnnnnnnnnnnnnnnnnnnnnnnnnnnnnnnnnnnnnnnnnnnnnnnnnnnnnnnnnnnnnnnnnaaggaaggagaagaagggaagggaaggaggaaaagaagaaagagggagga
aaagagagaaggaaggaaggggaaggaaagaaaaaaggaaaagaggtgaaggaagaaaggagaagaggggaagaaaagaaggaaagggagggagggaggaaaagaggggaaggaaggagaagaggggaaggaaggagaagaggggaag
gaaagaaggaaagggagggaggaaaagaaggaagggagggagggagggagggagggagggagggagggagggagggaggaaggaaggaaggaaggaaggaaggaaggaaggaaggaaggaaggaaggaaggagagattgacacagtga
atggatccatgtgcctcacctagaatggaaattgtttttgatagctggaggtaaaagatttggaatgcactcctctatggttctctcaggtcccccttttctcacttgttcccaaatgggatgatgtgaaaaagagaaggaaaaactaggaaaatcacccccaaagaag
agcacccacttcccctaccttggttgaagggagtggggacaatcggtcatctcttctccaggatctccaaatctatgatagcttcttcttctttttttaaacaagttttattaattttcaaagcacacaaatcatcattatccattacatgtttaaaccttttttccattataaatttaa
caagtcagataaactataattctgttttaaaccatttccatataaacatacaaaaacctcttgtccaatttctctttacttctctttactcttattcaatttcattattattatttcatctatttctttctgacttagttaccattattctcattctatatttctgcaaggttttattaattctcccc
ccaaaaaaaacatatacacacaacacataaaacataaaaaaaaatctttttggaaagtaacttggacccaggtatctctctcttcatggacttcaattatattcatgctcatcttacatatcattacttattttatttaccattctaagctttctatattaacttctccacatatttcac
tcttcttaattctatcttaattaatcatcttaatttgagtcattatatttgtatcttttgtcccttctttattgtccctttgtctcgttccttataaaaaaaaaaagaagtttctcctgaaatcactaatcagagtatgatgcctttctatctccccccctaaattaatcaaattaacctttaaaa
taacgcccaatacccaatccgattttgcttggtacattctatctaaatagtacatccagagtaattaaatacatagttaaacaataaatctaaatattaaaaacaaatactataatagaaattttataataacatcaataccataatcttattaaactctttaattaaaactcaata
ttctaaacctcccatcatattattaaattttaaaaccaatcattattaaacatttcaacattttaatctcttatcatctcttaaaccatttaagttttctatttctaataccatactacattatcatcttatgtaccttgcaaaatctttttgaagaaattatactttatcctttatccctaaaa
aagattccttttttcccatttaatttcaaaactttgtagcttttctccaccttttccctttcttcttctcccaacaaacctttctacccccacttcttctagcctctgccttactggcttcccttctctgtcgtgttcctttttcccttttgtttactatcattatattgattttgcaccttattttt
ccctttcttaagtgttttttaaatcttaagtcttttatttctcttcctttcactcttccttctttcttggtcaagttctgaaatttattaaaaatggcttcctctccttcttcttcttctgctgtcttcatctgttgtaatgtttgttgatagttaataagcgccatttctaaaaaaaggaaggc
ttctttcaaaccagatttaccattttttttgcttgttcataatcttttctcagttcctcaatagaaagagccattataaatcttttcttttttcccttctttcttcttacaacaaccagaatattccaaagaaaaaataaaacagtcccaaagttatcaatcagtccaaaccaaggcag
agttctcactttctatgacagcttcttgatccacctcggtggcttccttcatccctcatagccccttccaactgttctcgtctttttcctctttgtggggttcaggtacaaggtacggcgaagaaaaagagagtttgctgagaaggtcgaaacgaagcaagctgtggcatc
cgtcctgcacttgtccaggtgagtctcagctcttcatctccattggggagctggaagaaaatttgcctggtgggtgtcggagttccagaaaatctccactccaaccaagaactctgagacaggcatcttctcaaaagttctttttatttgggataataaactggcacatc
agggaaaacccgaatctgaaagttccaggcttccttacaataagaaaaaaactacaaatccacccccctgcgcccatcattgtatcacatgccccaatcgccatccgttccagcttgagatgtcctctcacccaggggcagagataacatgaccttgacttagaaaaa
gaatgttattatgtctaaccacatacgctccaagaaattccccccctcccattttcccaggctcaaaatgtggcagtgtggaagggcctagtttcaatatggcttccaaagctgacaggcagtttacttcagattcttccggcgccaggattgtgtggttgttttgttcatgg
gttgtcttttgggttttttttctctctgtctctgttctaggtgcctgactcaaaagcaaacaagctttgttgttcatcgtggtttcccttctgagggttgcatttcttcaaggggtgacctgtgggtcaggagaaagaacctctgtttccttctctgagacaccatgttttctcttcc
agacattctttcaatcgtcttcttttaatgaccccataaatcccttgcggaggaggagcctggccaactgaatggagctagcagaatgtttattggccagatgcccttcctgtcaccaatgctgagtttttgttcagcagagatattctccagagagagaaatatctgcct
ctacctaggatcgaactcacagtctcttgattgtgaggcgagagctctacctctaggccactgcatcacccctgacaaccttctaatgatggttcctaattgcttttagatttaaatgaattccaacagctaaagaagcacaaccatgcctgcccctgtaatggactcag
tttacatagtttgtcaggtaagaaaataatcctgggtgtaactcgtgtggaaggtcaaggaagatgggtgtgggtgttgtatagcctaaggatcttctcaggcgtgggagaaaggaagaaatcagaaaccacccatcgggacattttgagaatactccctcgttttaa
ttcagtccatttattaaaatcactttttgattttgcaagatactacagatagtagtatcaacttacaaccacaattgaaacctaaattgctgtggctaagcaaaacattgtttaagtgagttttgccccattttacgacctttctggccacagttcttaagtgaatcactgtgtcag
gattctgacacttaaccccatttaaatcatgagtctcgagccaaacttccaaaacttcaaaaggatttattgggaattaaactgtccggcattgtttgttgtcttctttaaaatacatatgtgcttattaaacagtgtattgtctggatcaatttgcttatacaagataatagtaa
ggtactagtagtaataataataatgatgataatagccataataatagtactgatgtttgttgatatataataacattcttacatcatttgcttatacatatcagtaatctttcatcttctaatttctactccttttgtctaacccagggatccccaaacttggcaagtttaaaactggt
ggactttaactcccagaattcctgagctaacatgactggaggaggaattctgggaattgaagtccacaagtcttaaagctgtcgaatttgaagacccctgatctaaccatcgagaaaacatatatcccatgttgaaaaatagcttgtttcttccttatcttgcctagtgcc
caaattttgttcacgtgaccattggggactgtgaggactggtcctaaatcatgttgctcggtgttgttataactttgagtggtcatgaaatgtggtaagttgaggactttcctgtacaaggaagtgaaactggcaggatgaacttgagactttagcacctgagggataga
tcaacgttgccaggggtggtattctacaggtttggactggttcaggtgaaccagtacttctggtgatcgtttggcccctgccctagtctgttctttatgttttgcagctcaactggttagcgtactcaccgaaccagttaaatcccaccactgaatgttttgttttatgaaact
cttccgaaggacccacaatacaacaaaccaccctatcagcaagccaactggcttttcaaatattcaaacatacagcccacccacccccaaaaaacttaaaacacccaacctcaaggtgaataccagcccaaacataatctacttctaggtagtctcaatttgcagc
cattcgtttgctgactgttcgaagataggggtgccctgaaaaaagggacttatgaccgtttttcagacagtgaggaggttggacagaaacctgggccagtcctggagtctggggaatgctctgacgagggctctgcgtcggaggcagagatggggccagggc
cgtctgacagtgatcagctgccttcggagtcaggaatgagtggggaagaagaacagctggagcctgttcccagtgtgcacatgcacagagctgccaggagaagggaacagttaaggaacaagggctgacttgggagtaaaggctcaggtggacggtgaat
gggtcacgggatccacatttggaagcttaacaactggttcgtatttatgatggtcacagtatccccggggtcaacgtggtcaccttttgcaaaacaagccaagtcaatgtggagatcagattcccttaataaactgtgttactatcttaacactgcagtggttcgcttaa
caacagtggtgggaaaggtcgtaagatgggacaaagctcccttaacaatggtctcgcttaacaatggaaaagtcaattgtggtcgtaagtcaaggactttctgcatacctttcagtcgattaaaagattcattattttattttattttttccctgttccctttcagaacatcg
caatcctcatccccatcgcaagagcaagaaacatgcactccataagaaaaaatgcaagatgtttttgagacactgcgcttttaacgggatgcacattcccttataggttatcttggttcttggcaagacccctgaggggattattacccacgacgcaatgttgattcttc
tgtatgcgcctttctctttttaaaaaaaaatattattgtttgctctcggaaaccacttgggttggttggaaataaagctcggagcttgaggcctttgtgggagtgaatggtccgtaaagagttgtgtggacaaataaaagtaataatagaacttgctatatgttgggcacatt
gcctgctccgcacattttg

#### C: Sequence of cDNA (coding region in red and highlighted in yellow)

caagtaggaaaaatcagtagaagccgtggccgttgacggagagacccaaggaacgccgctgtcgaaatggaccggttcttcttctttctggtgttaggggtccttgtcagcttggtttgctgctcagaaaaaggtacaaggtacggcgaagaaaaagagagttt
gctgagaaggtcgaaacgaagcaagctgtggcatccgtcctgcacttgtccagatttaaatgaattccaacagctaaagaagcacaaccatgcctgcccctgtaatggactcagtttacatagtttgtcagaacatcgcaatcctcatccccatcgcaagagcaag
aaacatgcactccataagaaaaaatgcaagatgtttttgagacactgcgcttttaacgggatgcacattcccttataggttatcttggttcttggcaagacccctgaggggattattacccacgacgcaatgttgattcttctgtatgcgc

#### D: Amino acid sequence (signal peptide shaded)

MDRFFFFLVLGVLVSLVCCSEKGTRYGEEKESLLRRSKRSKLWHPSCTCPDLNEFQQLKKHNHACPCNGLSLHSLSEHRNPHPHRKSKKHALHKKKCKMFLRHCAFNGMHIPL

**Fig. S7.** Gene synteny and architecture (**A**), genomic DNA sequence (**B**), cDNA sequence (**C**), and encoded protein sequence (**D**) of the unannotated *cxcl17* gene in the NCBI reference genome of *Notechis scutatus* (mainland tiger snake). The information was downloaded from the NCBI gene database (https://www.ncbi.nlm.nih.gov/gene).

**A:** Gene synteny and architecture

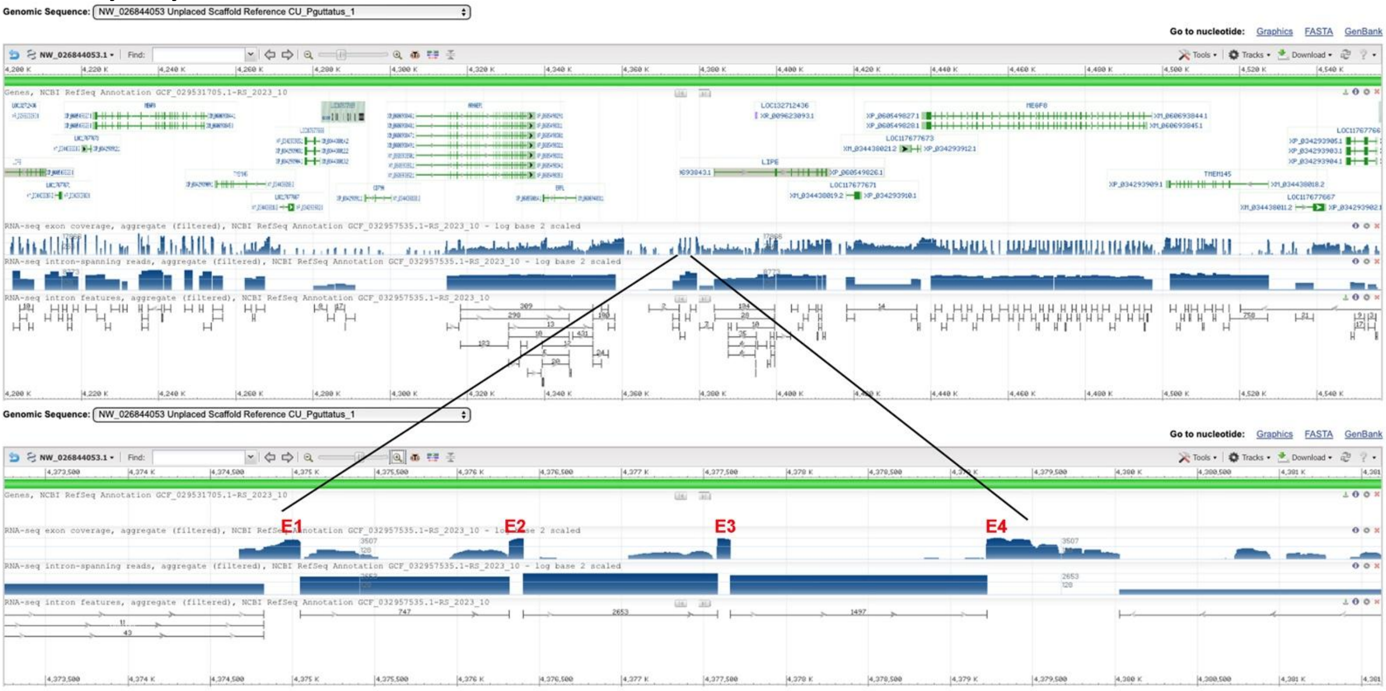


**B:** Sequence of genomic DNA (exons in red and highlighted in yellow, coding region underlined)

agctctctaaatcagtaaatattaaggtagaattacaggaagggaaaaacaaagtttgcacaatgtcttggggattgccccgatatgatttcttacttttacagaaacctttcccagaaatattacttagctgggaggagagagaaaaaatgcaaacaaacaaaaaaa
atataatacaagttgattggttcaagtacgaaaagttattagcggaagcagcggccattgatcgagaacgccgaggacgttgctctcgaaatggaccgattcttcatctttctgatgttagcgctctctgttggcttgctttgtcgctcagaaacaggtaagagggtttg
gagagggttattagattgttgcatgagaagaaaggagaggaaggggaggaagatccctggatctgtgctggacgagcaatgttccctcaacaccgttaagctggtcgttaaggtggaagctttcaagaagagactgggctgccctctggcagaaacggtgtag
ggcagaggtcttcacacttggcagctttaagacttgtggacttcaacccccagaattctccagccagctatgctggatgacgtccaggcgggtaaacagcctgaactggggcgaaccggtagaaactcacccctgcactaaatgttattaagttatggtactcaga
atcatacaacgcccaggaaaaaaataaccactccttaagtttaatgtatcgtacaaagtcattgagtttgtagtggagctggttcaacctatgtgagaagattacataagtcttctcacggatcttcaaccaaggatcaggaagggaggaagggaggaagggagga
aggaaggaaggaaggaaggaaggagagaaagaaagagaaggagggaggaaaggagggagggagggagggggagggggagaaagaaagagaaggcgggagggagggagggaggaaagggggaagggagggaggaaggaaggaaggga
ggggagaaagaaagagaaggtgggagggaaggaacgaaggagagaaagaaagggaggaaggaaggaaggagagaaagaaagagaaggagggagagagggaaggagggagggaggaaggaaggaaggaatggagggagggagggggag
aaagaaagagaaggcaggagggagggagggagggaaagtgggaagggaggaaggaaggggagggggagaaagaaagagaaggagggagggaggaagaaaggaaggaaggggggaacccccctccccaaaaatcacactttttttagtggattataatt
ttttaattaggtagctggagataaaagattttggggtgcactcctccatagttctcctcctttttctttttctcacctgttcccagtgggatggtgtgaaagaaagagggacgaactgggaagatcacccccgcacacacaaagaagggcacccacctcccctacctcg
gttgaagggagcggggagagttggtcatcctttctccaagatcgccaagtccacggtagcttcttgaaccacatccggtggctgctttcatccctcgtagccccttccaacctgatcccgtctttctccttttggggggggggtgtccaggtggaagctacggcgaag
gacaagagagtttgctgagcaggtcgaaacgaagcaagttgtggcatccgtcctgcccttgtccaggtgagtctcatctctggatctccatcggggagttgggagaaaattctcctgagggtttactttaggtctgtgatagcgaacctatgacacgcgtgctacag
gtggcatgcagacccctctctgtgggcacgccagctgttgccccagcccagctgcaccgcacatgcgcgccaggagtggattctatttacctttacaaccagttcgcaacgggatcgcccgtgcgcaactcacttcactcgcgcatgtgccttgcttctgcgcat
gtgcagaaacttctgacaatgtcagggtcagtgggcggatccttccaccggttttattaccggttctatagaaccggtccgaaccgggggcaacccaccactgaagcacatgcgcttctcgccagccagcagctcttcgggcctgtgccgtgcatgcgcgtggg
gcagggcacatgtgggggtgcacgcgtacatacatgggggcagggtgcatgcccagggggtgcatgcatgggggcgcttgcattgcattttgggggtgcacaccctgttttgggcctggaaagcctcttgcaccaatctggaagccaaaacaggggacagc
ggggatggggaacattcgtggggaggccagggagcatgcatggcgacaggggttcgggcacgcacatgctttgggcactcggcaccaaaaaggttaatcatcactgctttaggttcttccggcgccaagtttgtgtggttgtttgttcatgggttgtctctgggtt
ttttctctctctgttcttaggtgcctgttgctcaaaagcaaacaagctatgttgttaatcatggtttctcctttgagggttgcatttcttcaagggagggacctgtgggtcaggatgaggaacctcaatgttttctcttcctgacatctttctaatcatcttcttttaacgactccat
aattccttgaggggtaagagtctgggcaactgaatggagctagcagagcgttttactggccggatgcccttcttgttgccgatgcggagtttttattcagcagatatattctcattgtgcccagagagagagaaagatctgcctctacctaggatcgaactcacagcctc
ctgattgtgaggtgagacctctaccgccaccgcaccacccccgacgtccttctaatgatggttcctaattgcttttagatttaaatgaacgccaacatccaaagaaacatgaacatgcctgcccctgtgatggattcagcttacgtagtttgtcaggtaagaaaatgct
cctgggtgtaactcatgtggaactcataaaggaagatgggtccatgggtgtagcatcttcttaggcgtggatgaaaggaagaaatcagaaaccacccatcgggacttttctagaatactggctcgttttaattttgtccgccgattaaaatcacttttcattgtgcagga
tactgcttctctgaaaataagacatcccctgataataagctcaatcagacttttgaacgtaagcgctaaaataacccccccccaaaaaataagccctccctgaaaatatttaaacacatgtgcagccgctccccgccatttcctctggttagggttagggagacagag
ctggaaatcaggtaagacggcaagaggagccccgtctcgttccatgtgccccaaaaataataagacctccctgaaaataaggccaagcgcttatttcagggttcaaaaaaatataagacagggtcttattttcggggaaacatcaggggtgaaatgttcccggtt
cggaccggatcacccgatccggtagtgatggaggcgggtggttcggagaaccggtagcaaaaatccctgccccaccccggccatgcccagctgaaccgcgcgatcctcagaggctttttttgcttttaaatgcctctgacgatccaagctgagttgcctgatcgt
cagaggcatttaaaagaaaaaaagcctctgccgatcaggcaactcagctgggatcgtcagagccttttgaaagcatttttttacaacctcgtcgccgaaaaggttgtctaaaaaatggttttaaaagttaaaaaaaaaaaattggccacgaccacccagtcacattaa
accccccaccaccaccaccaagccacgtccacagaacaggtagtaacaaattttagatttcaccactgggaaacatggtacagatagtaacatcaacaattgaacccaaaattgctgtggctaagcgagacatcggttaagtgagttttgcctcgttttacgacctt
tcttgccacgtttgttaagtgaatcactgcagttgataaattagtaacacggttgttaagtgaatttggctttcccgttgactttcctggtgagaaggtcgcaaaaagggatcatgtgacccccaggacactgcgaccgtcgtagaccgactggctctatctctgcgtcg
gacggagagcatccatcggagcctcccccagactccagcactggtccaagttcctccgcaacctcctcatcgtccaagtctaccgccagttctgctgacagctggcggagccacaacaacaagcaaagtcaatggggaagttagattcccttaacaaccgcat
tactaacttaacagttgcagtggttcgcttaacaacggtggcgagaaaggtcgtaaaatgggacaaagctcccttatcaatggtctcgcttagcaacagaaatatacgactcaattgtggtcgtaagtggaggattatcttctaccgttcagtcttcaattgaccgacc
aaaagatgcactaatttattttatttttctctgttccctttcagaacatcgcaatcatcatcgtcatcatcatcgcaagaacaagcgacatgtgctccataagacaaaatgcaggaggtttttgaggcactgcacttttaaggggatccatgttcccttatagttcttggcac
gacccaaggagatttattatccatgacgcaatattgattcttcggttatgcgcctttctcgtttttttttaaaaaaaatatcattgtttgccctaggaaactgcttgattgtttggaaataaagctccgagcttgaggcctttgtgggagtgaatggcctgtaa

**C:** Sequence of cDNA (coding region in red and highlighted in yellow)

agttattagcggaagcagcggccattgatcgagaacgccgaggacgttgctctcgaaatggaccgattcttcatctttctgatgttagcgctctctgttggcttgctttgtcgctcagaaacaggtggaagctacggcgaaggacaagagagtttgctgagcaggt
cgaaacgaagcaagttgtggcatccgtcctgcccttgtccagatttaaatgaacgccaacatccaaagaaacatgaacatgcctgcccctgtgatggattcagcttacgtagtttgtcagaacatcgcaatcatcatcgtcatcatcatcgcaagaacaagcgacat
gtgctccataagacaaaatgcaggaggtttttgaggcactgcacttttaaggggatccatgttcccttatagttcttggcacgacccaaggagatttattatccatgacgcaatattgattcttcggttatgcgcctttctcgtttttttttaaaaaaaatatcattgtttgcccta
gg

**D:** Amino acid sequence (signal peptide shaded)

MDRFFIFLMLALSVGLLCRSETGGSYGEGQESLLSRSKRSKLWHPSCPCPDLNERQHPKKHEHACPCDGFSLRSLSEHRNHHRHHHRKNKRHVLHKTKCRRFLRHCTFKGIHVPL

**Fig. S8.** Gene synteny and architecture (**A**), genomic DNA sequence (**B**), cDNA sequence (**C**), and encoded protein sequence (**D**) of the unannotated *cxcl17* gene in the NCBI reference genome of *Pantherophis guttatus*. The information was downloaded from the NCBI gene database (https://www.ncbi.nlm.nih.gov/gene).

**A:** Gene synteny and architecture

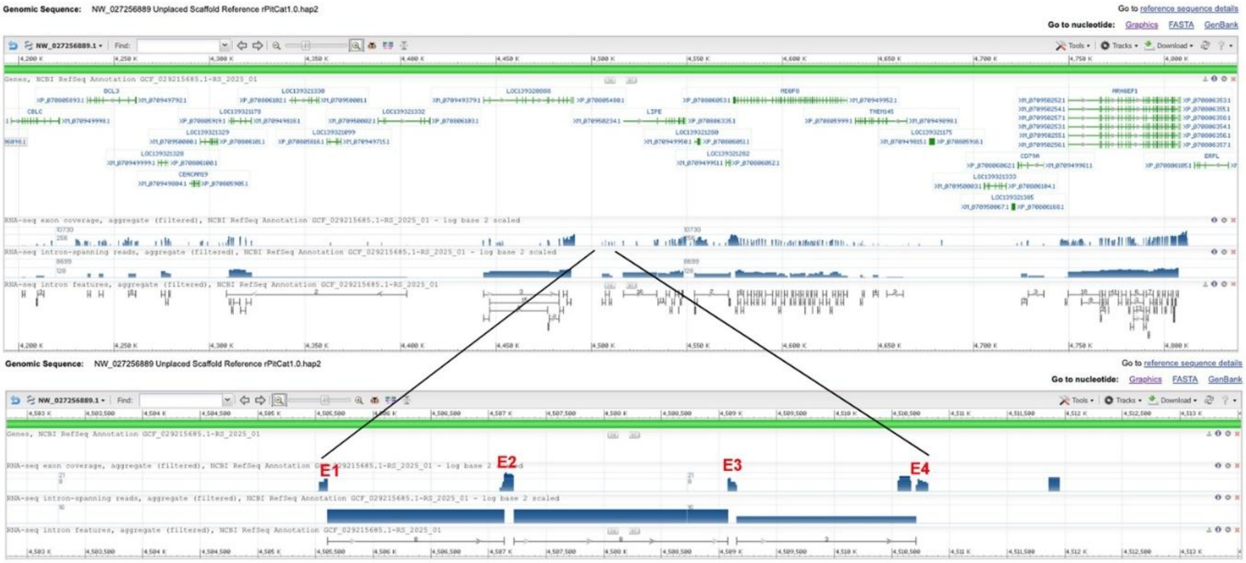

**B:** Sequence of genomic DNA (exons in red and highlighted in yellow, coding region underlined)

tctggcagctctctaaatcagtaaatattaaggtagaattacaggaagggaaaaacaaagtttgcacaatggcttggggattgcccacgatatgatttcttacttttacagaaacctttcccagaaatattacctagctgggaggagagagaaaaaatgcaaacaaaa
aaaaatatatataatacaagttgattggttcaagtacgaaaagttattagcggaagcagcggccattgatcgagaacgccaaggacgttgctctcaaaatggaccgattcttcatctttctgatgttagcgctctctgttggcttgctttgtcgctcagaaacaggtaagag
ggtttggagagggttattagattgttggatgagaagaaaggagaggaagggcagaaacggtgtagggcagaggtcttcacacttggcagctttaagacttgtggacttcaacccccagaattctccagccagctatgctggatgacgtccaggcgggtaaaca
gcctgaactggggcgaaccggtagaaactcacccctgcactaaatgttattaagtcatggtactcagaatcatacaacgcccaggaaaaaaataaccactccttaagtttaatgtatcgtacagtcattgagtttgtagtggagctggttcaacctatgtgagaaggc
tacataagtcttctcaaggatcttgaaccaaggatcaggaaggaaggaaggaaggagggaaggaaggaaggaaggaaggaaggaaggaaggaaggaaggaaggaaggaaggaaggaagaggggagggaggggaggggaggggaggggagg
ggagaaagaaagagaaggtgggagggagggagggaggaagggaggggaggggagaaagaaagagaaggtgggagggagggagggaaggaaggagagaaagaaagagaaggagggagggaggaaaggagggagggatcaagaaaggaac
ggaaggagggagggagggagggaggggaggggagaaagaaagaaggtgggagggagggagggagggaagggggaagggaggaagggagggagggagggaggaaggaagaggaggggaggggagaaagaaagagaaggcgggagggagg
gaggaaggaaggggaggggaggggagaaagaaagagaaggtgggagggagggaaggaaggagagaaagaaagggaggaaggaaggagagaaagaaagaaggagggagggagggaaggagggagggtggaagggagtgaggaaggaag
gaaggaaggaaggaacggagggagggaggggaggggagaaagaaagagaaggcaggagggagggagggagggagggaaggtggaaagggaggaaggaaggggaggggagaaagaaagagaaggagggagggagggagggagggagg
gagggaggaaggaaggggaggggaggggagaaagaaagagaaggtgggagggagggagggagggagggagggaggaaggaaggaaggggggaaccccctcccccaaaaatcacactttttggtggatgataatttttttaattaggtagctggaaata
aaagattttggggtgcactcctccatagttctcctccttttctttttctcacctgttcccagtgggatggtgtgaaagaaagagggacgaactgggaagatcaccccccccacacacacaaagaagggcacccacctcccctacctcggttgaagggagcggggag
agttggtcatcctttctccaagatcgccaagtccacgatagcttcttgaaccacatttggtggctgctttcatccctcatagccccttccaacctgatcccgtctttctccttttggggggggggggcgtccaggtggaagctacggcgaaggaaaagagagtttgc
tgagcaggtcgaaacgaagcaagttgtggcatccgtcctgtccttgtccaggtgagtctcatctctggatctccatgggagagttgggagaaaattcgcctgagggattactttaggtctgtgatggcgaacctatgacacgcgtgctacaggtggcatgcagacc
cctctctgtgggcacgccagccgttgccccagcccagctgcaccgcacatgcacgccaggattcgggttctatttacctttactactggttcgcaacaggattgcccatgtgcaactcatttcactcgcgcatgtgtcttgcttctgcgcatgcgcagaaacttctga
caatgtcagggtcagggggcggatccttccaccggttttattaccggttctatagaaccggtccgaaccgggggcaacccaccactgaagcacgcgcgcttctcgccagccagctgctcttcgggcctctgccgtgcatgcgcgtgggacggggcatatgcg
ggggtgcacgcgtacatacatgggggtaggacgcatgcaccgggtgtgaattcatgggggcgcttgcattgcattttgggggtgcacgccctgttttgggcctggatgttgtggctccgacccccgaggccgacttatcagagggagaaggctccgaacatga
aggagaggagttggccaggccttcctctccaggaccctcctctggcaatgcctcaggtgtctcccgtaggagaagagattgaccaggagccaagtatctcctcacagtcggacagtgaaggggataagtcttggctggatccacgccagcgtcggagaga
caagcgcactcaacagaagaagaggcatgccagagattaagctgagtcactggaccacaccccacaggggatataagggagtgaggtggccattcagactctgtgacggacaaaactaactactatcgagtcacattgcatctttggaaacaaggttatctgc
cttggatttaattatggactggacttgacaattcttgaactgtgtgtatcttgtttatgagtaaagttcttggcaagcagctactgggttgctgccaatccttttatcgaaagaatttgttgtctgccagtttatttaacccctggctgtctgtttgtgtgtgtgtgtgagacggg
gacagaacactggaaagcctcttgcaccaacctggaagccaaaacgggggatggggggacagggaacatacgtggggaggccagggagcatgtatggtgacaggggtttgggcacgcgcacatgcatttgggcacgcgcatgctttgggcactcggc
accaaaaaggttaatcatcactgctttaggttcttccggcgccaagattgtgtggttgtttgttcatgggttgtctctgggtttttctctctctgttcttaggtgcctgttgctcaaaagcaaacaagctatgttgttaatcatggtttctcgtttgagggttgcatttcttcaagg
gggggacctgtgggtcaggatgaggaacctcaatgttttctcttcctgacatctttctaatcatcttcttttaacgattccataatcccttgggaggtaagagtctgggcaactgaatggagctagcagagcgttttactggccaaattgctgagcacattgtaagccgc
cctgagtcttcggagaagggcggcatataaatcaaattaaaaaaaaaaaaataaaaaaaaaatgcccttcttattgccgatgcagagtttttattcagcagatatattctcattgtgcccagagagagaaagatctgcctctacctaggatcgaactcacagcctcctgat
tgtgaggtgagagctctaccgccaccgcaccacccccgacgtccttctaatgatggttcctaattgcttttagatttaaatgaacgccaacatccaaagaaacatgaacatgcctgtccctgtgatggattcagcttacgtagtttgtcaggtaagaaaatgctcctgg
gtgtaactcgtgtggaaggtaaaggaagatgggtccatgggtgtagcatcttcttaggcgtggatgaaaggaagaaatcagaaaccacccattgggacttttctagaatactggctcgttttaattttgtccgccgattaaaatcacttttcattgtgcaggatactgctt
ccctgaaaataagacatcccctgataatacgcccaatcaggcttttgagcataagcgctaaaataaaccgcccccctgaaaataagccctccctgaaaatatttaaccacaggcacagccagtccccgccatttcctctgtttagggttagggagacagagctgg
aaatcaggtaagacggcaagaggagccccgtctcgctccatgtgccccaaaaataataagacctccctgaaaataaggccaagcgcttatttcagggttcaaaaaaatataagacagggtcttattttcggggaaacatcagggatgaaatgctcccagttcgga
ccggatcgcccgatccggtagcgatggaggcgggtggttcggagaaccggtagcaaaaatccctgcccccccccatgcccagctgaaccacgcgatcatcagaagcttttttcttttaaatgcctcagacgatccaagctgagttgcctgatcgtcagaggcatt
taaaagaaaaaaattctctgccgatcaggcaactcagctgggatcatcagagccttttgaaagcactttttacaacctcttcgccgaagaggttgtcgaaaaaatggttttaaaactttaaaaaaaaaattggccacgcccacccaatcacatttccccaccaccacc
accaccaccaccaagccacgcccacagaaccggtagtaacaaattttagatttcaccactgggaaacatggtacagatagtaatatcaacaattgaacccaaaattgctgtggctaagcgagacatcggttaagtgagttttgcctcgttttacgacctttcttgcca
cgtttgttaagtgaatcactgcagctgataaattaataacacggttgttaagtgaatttggctttcccgttgactttcctggtgagaaggtcacaaaaagggatcatgtgacccccaggacactgcgaccgtcgtagaccgactggctctatctctgcgtcagatgga
gagcatccatcggagcctcctccagactccagcactggcccaagttcctccgcaacttcctcatcatccaagtctaccgccagctctgctgacagctggcggtgccacaacaagcaaagtcaatggggaaatcagattcccttaacaaccgcgttactaacttaa
cagttgcagtggttcgcttaacaacggtggcgagaaaggtcgtaaaatgggacaaagctcccttatcaacggtctcgcttagcaacggaaatatacgactcaattgcggtcctaagtggaggattatcttctaccgttcagtcttcaattgaccgactaaaagatgc
actaatttattttattttttctctgttccctttcagaacaccgcaatcatcatcatcatcgtcatcatcatcgcaagaacaagcgacatgtgctccataagacaaaatgcaggaggtttttgagacactgcacttttaaggggatccacgttcccttatagttcttggcacgac
ccaaggagatttcttatccatgacgcaatattgattctttggttatgcgcctttctcgtttttttaaaaaaaaatcattgtttgccctaggaaactgcttgattggttggaaataaagctccgagcttgaggcctttgtgggagtgaatcgcctgtaaagaattgtgtggacaa
ataatagtagcaatggaacttgctatacgccgggcacgcttcctgctgcaaacattttgcaatcccttgtaaaaacccagagaggtgcagaaaataaaggcaacggggatatcatttccaaaagggatggaagtctttagcctaccctttgcaaaatgctaaccgg
aagatggaggtagtcttctacctacaatcagaatagagt

**C:** Sequence of cDNA (coding region in red and highlighted in yellow)

Agttattagcggaagcagcggccattgatcgagaacgccaaggacgttgctctcaaaatggaccgattcttcatctttctgatgttagcgctctctgttggcttgctttgtcgctcagaaacaggtggaagctacggcgaaggaaaagagagtttgctgagcaggt
cgaaacgaagcaagttgtggcatccgtcctgtccttgtccagatttaaatgaacgccaacatccaaagaaacatgaacatgcctgtccctgtgatggattcagcttacgtagtttgtcagaacaccgcaatcatcatcatcatcgtcatcatcatcgcaagaacaagc
gacatgtgctccataagacaaaatgcaggaggtttttgagacactgcacttttaaggggatccacgttcccttatagttcttggcacgacccaaggagatttcttatccatgacgcaatattgattctttggttatgcgcctttctcgtttttttaaaaaaaaatcattgtttgc
cctaggaaactgcttgattggttggaaataaagctccgagcttgaggcctttgtgggagtgaatcgcctgtaaagaattgtgtggacaaataatagtagcaatggaacttgctatacgccgggcacgcttcctgctgcaaacattttgcaatcccttgtaaaaaccca
gagaggtgcagaaaataaaggcaacggggatatcatttcca

**D:** Amino acid sequence (signal peptide shaded)

MDRFFIFLMLALSVGLLCRSETGGSYGEGKESLLSRSKRSKLWHPSCPCPDLNERQHPKKHEHACPCDGFSLRSLSEHRNHHHHRHHHRKNKRHVLHKTKCRRFLRHCTFKGIHVP
L

**Fig. S9.** Gene synteny and architecture (**A**), genomic DNA sequence (**B**), cDNA sequence (**C**), and encoded protein sequence (**D**) of the unannotated *cxcl17* gene in the NCBI reference genome of *Pituophis catenifer annectens*. The information was downloaded from the NCBI gene database (https://www.ncbi.nlm.nih.gov/gene).

**A:** Gene synteny and architecture

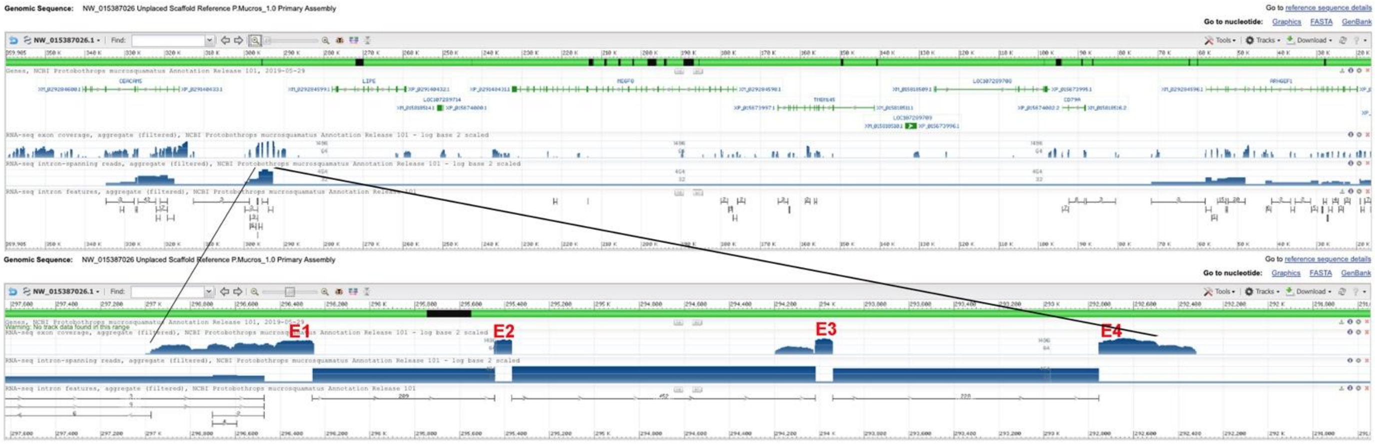


**B:** Sequence of genomic DNA (exons in red and highlighted in yellow, coding region underlined)

ccaagcatttcccagaagtattagccgggagaagaagaaattttttttaaaaaaaaataataataaatacgtcggttggtttaagtacaaaaggttattagccgaagcggtggccattgacggagagagagagccaaggacgtggctggagaaatggacagatt
cttcttatttttgatgtcagcgctcctcgttggcctggtctgcagctcagaaaaaggtaggaggagttggagagacttattgggaattgttgcatgggaagaaaggagaggaaagtgaaggagatccctggattggcaccagacaagcaacgttccctcagcaac
attaacctggtcattaaattaacccttttctctagatttgcacaataccctgagccataaactggctaggaagcaataacaactccctaagtttagtatatcgtacaaagccatcgtggtgggagtggggctggtccaacctgtgcttgaacaccacccaggtcttctc
actgatctttgaccaaggatctgtagactgtctagaggtctgtagtgaagaggaaggaaggaaggaaggaaggaaggaggagagggaagggagggagggagagaaagaagggagggagggagggacagaggaagggagagaaaggaggaggggag
ggaaagaagaaggaaaggaaagggagagagggagggaggaaggaaggaaggaaggaaggaaggaaggaaggaaggannnnnnnnnnnnnnnnnnnnnnnnnnnnnnnnnnnnnnnnnnnnnnnnnnnnnnnnnnnnnnnnnnnnnnnnnnnn
nnnnnnnnnnnnnnnnnnnnnnnnnnnnnnnnnnnnnnnnnnnnnnnnnnnnnnnnnnnnnnnnnnnnnnnnnnnnnnnnnnnnnnnnnnnnnnnnnnnnnnnnnnnnnnnnnnnnnnnnnnnnnnnnnnnnnnnnnnnnnnnnnnnccaggatcac
caagtccacgagagcttcttgaaccacatcagtggattccttcatccctcgtagcctcttccaacctgctctcacctttctccttttgggcttaggtggaaggtacgaccaagcgagtttgctgagcaggtcgaaacgaagcaggcaggggcctcggtcctgcccctt
gtccaggtgagtcccgtctcttgatttccatcggggagctggaagaaaatttgcctgcaagtttactttgggttcttctggcatccaggatcacatggcgcaaagtgcaaagaagagcaactaagatgattaaaggcctggagactaaaacataggaagaacggtt
gcaggatttgggtatggccagtctagagaaaagaaggactgggggagacatgatagcagtgatccaatatttgaggggctgccacaaggcagaggggggggggtcaacttcttttccaaatcacctgaaggtaggataagaagcaatggttggaaagtaatc
aaggagagaagcaacctggaattaaggagaaagttcctaatagggaggacaattaaccagtggaacggcttgccttcagatgttgtgggtgcttcatcactggaggtttttaagaagagactggacagccacttgtctgaaatgacataggttctcctgcttgagc
agggggggtggactagaagacctccaaggtccctttccattccattccattctattctattctattctattctgttctattctattctattcttctacttcattccattattctattctattctattctattctgttctgttctgttctgttctgttctgttctgttctgtttttttcctattcttc
tacttcattccattcttctattctattctattctattctattctattctattctattctattctgttctgttctgttctattctattctattctattctattctaaaggactggggtcacatgatagcagtgttctgatatttgaggggctgccgcaaagaagaggggggtcaacttattttcg
aaagtaccagaatgcaggacaagaagcaacgatgaaaactaatcaaagagagaagcaacctagaactaaggagacatttcctgacaggaagaactattaatcagtggaacaacctgcctccagaaatcatgggcactccatcactggaggttttcaagaaga
gactggacagccacttgtctgaaatggcataggttctcctgcttgactagaagaccaccaaggtctcttccagctctattctgattgattgattgattgattgattgattgattgattatgggttgtttctttgggtcttttctctctgttccaggtgtctgacttaaaaacagac
aagctctgttgttatcctcattactcttttgaggattgcatttcctcaaggatggcttatgggtcaggaggaagaacctcgctgtctttctctgatgccccatgttttctcttcctggcgtccttctaatgctggttcctaatcatttttagatttaagtgaactgcaataccccaa
aaaggacaaacatccttgtccctgtgatggactcaccttaagtagtttgtcaggtaagaaaacactcccgggtgtaactcctgtggaaggtaaatgaagaagggtgtgtgggtgtgggtggctgttgtgtagcctaagcatcttattaggcctggatgagaaggaa
aaatcagaaatcgcccattgggatttttctaccatccccatttattttcattttgcccacccattaaaatcacttcttactttgcaggatactacaggtaatcctcgacttacgaccacagtggatcctaaaattgctgttgctaagtgaaacattggttaagtgagtcttgccc
cattttacgacacagttaaatgaagttggtaacatggctgttcagtgaaattggcttccccactgacagaaggtcgcaaaaaggaggccgcatggttccaggatgctgcaaccgtcataaatatgagtcaaaaattgaaaaaaaaatggacttattaccgtttttcac
actcacaacccttagggcaggggtaggccactttggctcttttatgactagtggacttcaattcccagaattcctgagccagccatgctgaggggaaagggagaggtttctaagctgagccatgctggctcaggaattctgggaattgaagtccacccatctccga
gtggccaaggtggagaaatacaggcctaggccaggagtaggcaaccttggctcttttatgactcgtggacttcaactcccagaattcctgagccagccgtgctgaggggaaagggagaggtttccaagctgagccctgctggctcaggaattctgggagttga
agtccacgagttataaaagagccaaggttgcctaccccctgccttagggcatccccatggtcacgtgatcaaacttcggaggctcagcaactggcttgtacttatgacagttgcagtgtcctgggtccatgggatccctttttacgaccttctgaccaatggagcca
gattcacttaacaaccatgttactaacttaacaacagcaatgactcacttaacaactgtggcaagaaaggtcgtaaaatggggcaaagctcacttaaccactgtctcacttagcaatggaaatgttaggctcaattgtggtttgtaggtcgagaactacctatatacctt
tcggtcttcaattgaccgattaaaagattcattattaatttttttttctgttcctttttagaacaccccagtcatcatcatcatcatcctcctcgcaagaacaagcaacttgtgcttcgtaagaaaaaatgcaggagatttttgagacactgcgcttttaaagggatccatattc
ccttatagcttctcccaaggggattatccgtgacgccatattgattctttgcttatgtacctttctcttcttttaaatatcattgtttgctctaggaaactgcttgcttggttggaaataaagctccaggcttgagggctttgtggaagtgaacggcctgtaaataattgtgtaga
caagttaaataagaatatgttactatatgtgt

**C:** Sequence of cDNA (coding region in red and highlighted in yellow)

Gtacaaaaggttattagccgaagcggtggccattgacggagagagagagccaaggacgtggctggagaaatggacagattcttcttatttttgatgtcagcgctcctcgttggcctggtctgcagctcagaaaaaggtggaaggtacgaccaagcgagtttgct
gagcaggtcgaaacgaagcaggcaggggcctcggtcctgcccttgtccagatttaagtgaactgcaataccccaaaaaggacaaacatccttgtccctgtgatggactcaccttaagtagtttgtcagaacaccccagtcatcatcatcatcatcctcctcgcaag
aacaagcaacttgtgcttcgtaagaaaaaatgcaggagatttttgagacactgcgcttttaaagggatccatattcccttatagcttctcccaaggggattatccgtgacgccatattgattctttgcttatgtacctttctcttcttttaaatatca

**D:** Amino acid sequence (signal peptide shaded)

MDRFFLFLMSALLVGLVCSSEKGGRYDQASLLSRSKRSRQGPRSCPCPDLSELQYPKKDKHPCPCDGLTLSSLSEHPSHHHHHPPRKNKQLVLRKKKCRRFLRHCAFKGIHIPL

**Fig. S10.** Gene synteny and architecture (**A**), genomic DNA sequence (**B**), cDNA sequence (**C**), and encoded protein sequence (**D**) of the unannotated *cxcl17* gene in the NCBI reference genome of *Protobothrops mucrosquamatus* (brown spotted pit viper). The information was downloaded from the NCBI gene database (https://www.ncbi.nlm.nih.gov/gene).

**A:** Gene synteny and architecture

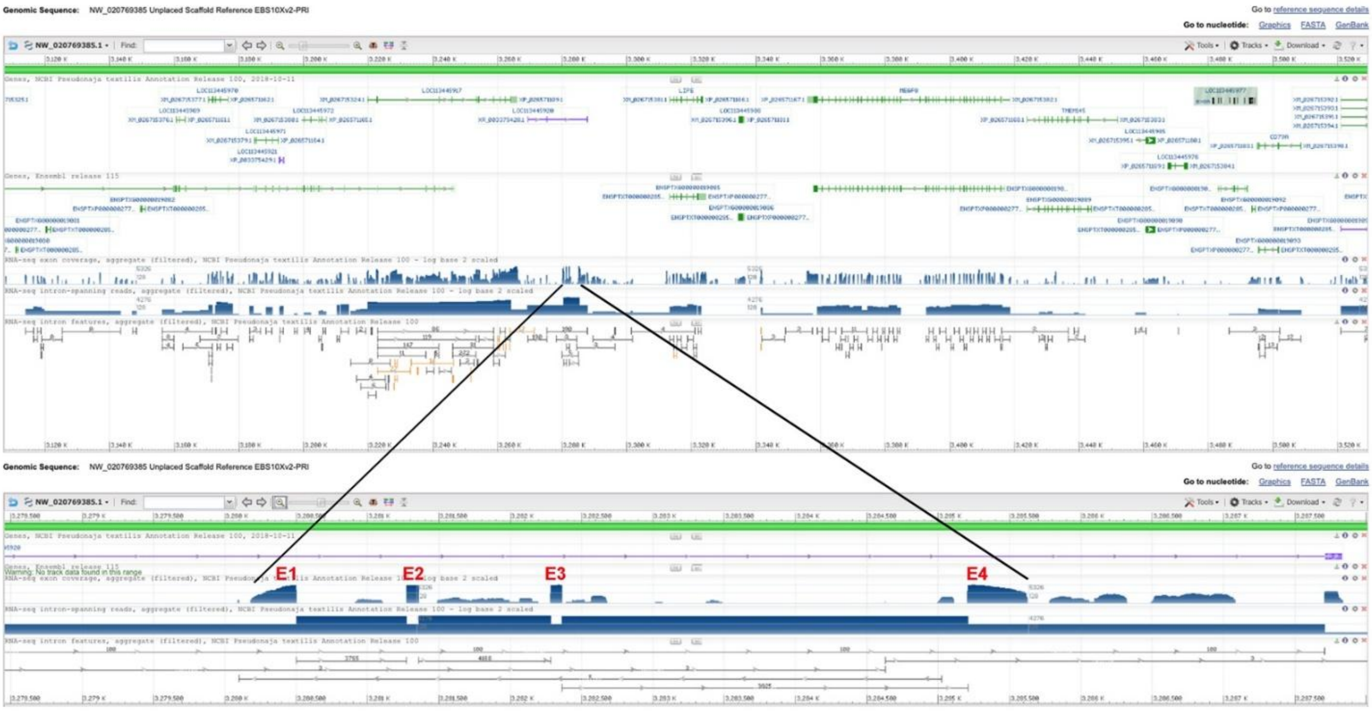


**B:** Sequence of genomic DNA (exons in red and highlighted in yellow, coding region underlined)

gactgccctgatatgatttcttattcctacacaaagcctttcccagaaatattacccagctgggaggaggggagaagttggttggttcaagtaggaaaaataagtggaagccgtggccgttgacggagagacccaaagaacgctgctgccgaaatggaccggtt
ctccttctttctggtgttgggggtccttgtcagcttggtttgctgctcagaaaaaggtaagacgatttggagacggggagggagggagggaggaaggaaggaaaagaggagaaggaaggaaggggaagtggggaaggaaagaagggagagaggggag
ggagggaggaaaaggggaaaggaaggagaagaagggaaggaaaggaggaaaagaagaaagggagggaggaaaagagagaaggaaggnnnnnnnnnngaaggaaggaaggggaaggaaagaaaaaggaaaagaggtgaaggaaggaaa
gagaagaggggaaggaaagaaggaaagggagggagggaggaaaagagggaagggaggaaggaaaagaggggaaggaaggagaagagggaaaagaaagaaggaaagggagggaggaaaagaaggaaggaaggaaggaaggaaggaagg
aaggaaggaaggaaggaaggaaggaaggagagagattgacacagtgaatggatccatgtgcctcacctagaatggcaactgtttttgatagctggaggtaaaagatttggaatgcactcctctatgattctctcaggtcccccttttctcacctgttcccaatggg
atgatgtgaaaaagagagggacaaactaggaaaatcacccccaaagaagagcacccacctcccctagcttggttgaagggagtggggacaatcagtcatctcttctccaggatctccaaatctatgatagcttcttgatctacctcggtggcttctttcatccctcat
agccccttccaactgttctcgtctttttccttttgtggggttcaggtagaaggtacggcgaagaaaaagagagtttgctgagaaggtcgaaacgaagcaagctgtggcatccgtcctgcacttgtccaggtgagtctcagctcttcgtctccattggggagctggaa
gaaaatttgcctggcgggtgtcagcgttccagaaaatctccactccaaccaagaactccgagacaggcatcttctcaaaagttctttttattttggataataaattggcacatcagggaaaacccgaatctgaaagttccaggcttccttacaataagaaaaaactaca
aatccacccctgcacccatcattgtatcacatgccctaatcgccatccgttccagcttgagatgtcctctcacccagaggcagagataacatgaccttgacttagaaaaagaatgttattatgtctaactgcatacgctccaagaaattcccccccctccctttttcccag
gctcaaaatgcggcagtgtggaagggcctagtctgaatatggcttccaaagctgacaggcagtttacttcaggttcttccggtgccaggattgtgtggttgttttgttcatgggttgtcttttggggtttttttctctctgtctctgttctaggtgcctgactcaaaagcaaa
caagctttgttgttcatcatggttttgtcttctgagggttgcatttcttcaaggggtgaactgtgggtcaggagaaggaacctgtttccttctctgaaacaccatgtttttttcttccagacatcctttcaatcgtcttcttttaatgaccccataaagccaactgaatggagcta
gcagaatgtttactggccagatgcccttcctgtaaccaatgctgagtttttgtttagcagagatattctccagagagagagaaatatctgcctctacctaggatcgaactcacaatctcctgattgtgaggcgagagctctacctctaggccactgcatcacccctgacaa
tcttctaatgatggttcctaattgcttttagatttaaatgaattccaacacctaaagaagcacaaccatgcctgcccctgtaatggactcagtttacatagtttgtcaggtaagaaagtaatcctgggtgtaactcatgtggaaggtcaaggaagatgggtgtgggtgttg
tatagcctaaggatcttcttaggcgtggaagaaaggaagaaaccagaaactacccatcgggacttttcgagaatactcgctcgttttaattctgtccatttattaaaatcgcttttgattttgcaggatactacagatagtagtatcgactgacaaccacaattgaaacct
aaattgctgtggctaagcaaaacattgtttaagtaagttttgccccattttatgacttttcgggccacagttcttaagtgaatcactgtgtcaggattctggcacttaaccccatttaaatcatgagtctcgggccaaacttccaaaacttcaaaaggatttattgggaattaa
actgtccggcattgcctgggtgtagccgactctgagacacatcattcattgtcacccttgaccctagtttttccccaccccctgggtgtattgtaattcatattctccaatgggacacatccatagccaagaggcatcaatcatgctctccagttagaattttgcgcggca
gaaacttcagtcatctagttactatctttgaaaaacatttatggaatgcagtttcggtttgcaagctgctacattgctccccccctctctttctgttggcaaccctaaaaggaccagagaaatatggatttatgagttgacacactgcaattgataagttagtaacatggttgtt
aagtgaatctggctcccccattaactttgcttgtcagaaggttgcgaaacaaggatctcgtgaccccaggacactgcagctgacataaatacaagtcagttgccaagcacctgaattttgatcatgtgaccatgggaaatctgcaagtgtagaagtgtgaaaaatgg
tcagaagtcactttttttcagtgccgctgtaacttcaagcagtcattaaatgaactgtcacaagtcaaggactacctgtatccagcccattactcttctacacaagagaatctttagcattgccccatggttgattttgcaaatttccctgaatctaagttgcagtttcaatac
atacaggtaacccttaacttacaactagcggggccagaattttcttcttaagcagagcaattttataactgttttgctgaagctgttaagcaaatccctgaaatttggtaagtgaatcatgtaaattcatgctggtggtctagtgcccacatttttcatttcatttcatttatttat
acatttgttatcttctttaaaaatacatatgtgcttattaaacagtgtattgtctggatcaatttgcttatacaagataatagtaaggtacaagtagcaataataataatgatgaaaatagccataataatagtactgatgtttgctgatatataataacattcttacatcatttgctta
tacatatcagtaatctttcatcttctaatttctactccttttgtctaacccagggatccccaaacttggcaagtttaaaactggtggactaactcccagaattcctgagctaacaggactggaggaggaattctgggaattgaagtccacaagtcttaaagctgtcgaattt
gaagacccctgatctaaccatcgagaaaacatatatcccatgttgaaaaatagcttgtttcttccttatcttgcctagtgcccaaattttgttcacgtgaccactgggactgtgaggactggtcctaaatcatgttgctcggtgttgttataactttgagtggtcatgaaatg
ttgtaagttgaggactttcctgtacaaggaagtgaaactggcaggatgaacttgagagtttagcacctgagggatagatcaacgttgccaggggttggactggttcgggtgaaccagtagttctggtgatcatttggcccccgccctactctgtcctttatgtccatgt
tttgcaactcagctagttagcgtactcaccgaaccagtggaatcccaccatggaatgttttgtttgatgaactcttctgaaggacccacaatacaacaaaccacccttccagcaagccaactggcttttcaaatattcaagcataaagcccacccacccccaaaaaac
ttaaaacacccaacctcaaggtgaataccagcccaaacataatctgcctctaggtagtctcaatttgcagccgttcgtttgctgactgttcgaagataggggtgccctgaaaaaagggacatgaccatttttcagacagtgaggaggttggacagaaacctgggtc
agtcctggagtctggggaaggctctgatgagggctctgcgtcggaggcagagatggggccagggccgtctgacagttatcagctgccttcagagtcaggaatgagtggggaagaagaacagctggagcctgttcccagtatgcacatgcacagagctgcca
ggagaagggaacagttaaggaacaagggctgacttgggagtaaaggctcaggtggacgatgaatgggtcacgggatccacattcggaagctcgtatttatgatggtcacagtatccccgggatcacgtggtcaccttttgcaaaacaagccaagtcaatgtgg
agatcagattcccttaataaactgtgttactatcttaacactgcagtggttcgcttaacaacagtggtgggaaaggtcgtaagatgggacaaagctcccttaacaatggtctcgcttaacaacggaaaagtcaaatgtggtcgtaagtcaaggactctctgcatacctt
tcagtcgattaaaagattcattattttattctattttttcccctgttccctttcagaacatcacaatcctcaccctcatcgtaagagcaagaaacatgcactccgtaagaaaaaaatgcaagatgtttttgagacactgcgcttttaatgggatgcacattcccttatagtttatct
tggttcttggcaagacccctgaggggattattatccacgatgcaatgttgattcttttgcatgcgcctttctcttttttaaaagaaatattattgtttgctctcggaaaccatt

**C:** Sequence of cDNA (coding region in red and highlighted in yellow)

Agtggaagccgtggccgttgacggagagacccaaagaacgctgctgccgaaatggaccggttctccttctttctggtgttgggggtccttgtcagcttggtttgctgctcagaaaaaggtagaaggtacggcgaagaaaaagagagtttgctgagaaggtcga
aacgaagcaagctgtggcatccgtcctgcacttgtccagatttaaatgaattccaacacctaaagaagcacaaccatgcctgcccctgtaatggactcagtttacatagtttgtcagaacatcacaatcctcaccctcatcgtaagagcaagaaacatgcactccgt
aagaaaaaaatgcaagatgtttttgagacactgcgcttttaatgggatgcacattcccttatagtttatcttggttcttggcaagacccctgaggggattattatccacgatgc

**D:** Amino acid sequence (signal peptide shaded)

MDRFSFFLVLGVLVSLVCCSEKGRRYGEEKESLLRRSKRSKLWHPSCTCPDLNEFQHLKKHNHACPCNGLSLHSLSEHHNPHPHRKSKKHALRKKKCKMFLRHCAFNGMHIPL

**Fig. S11.** Gene synteny and architecture (**A**), genomic DNA sequence (**B**), cDNA sequence (**C**), and encoded protein sequence (**D**) of the unannotated *cxcl17* gene in the NCBI reference genome of *Pseudonaja textilis*. The information was downloaded from the NCBI gene database (https://www.ncbi.nlm.nih.gov/gene).

**A:** Gene synteny and architecture

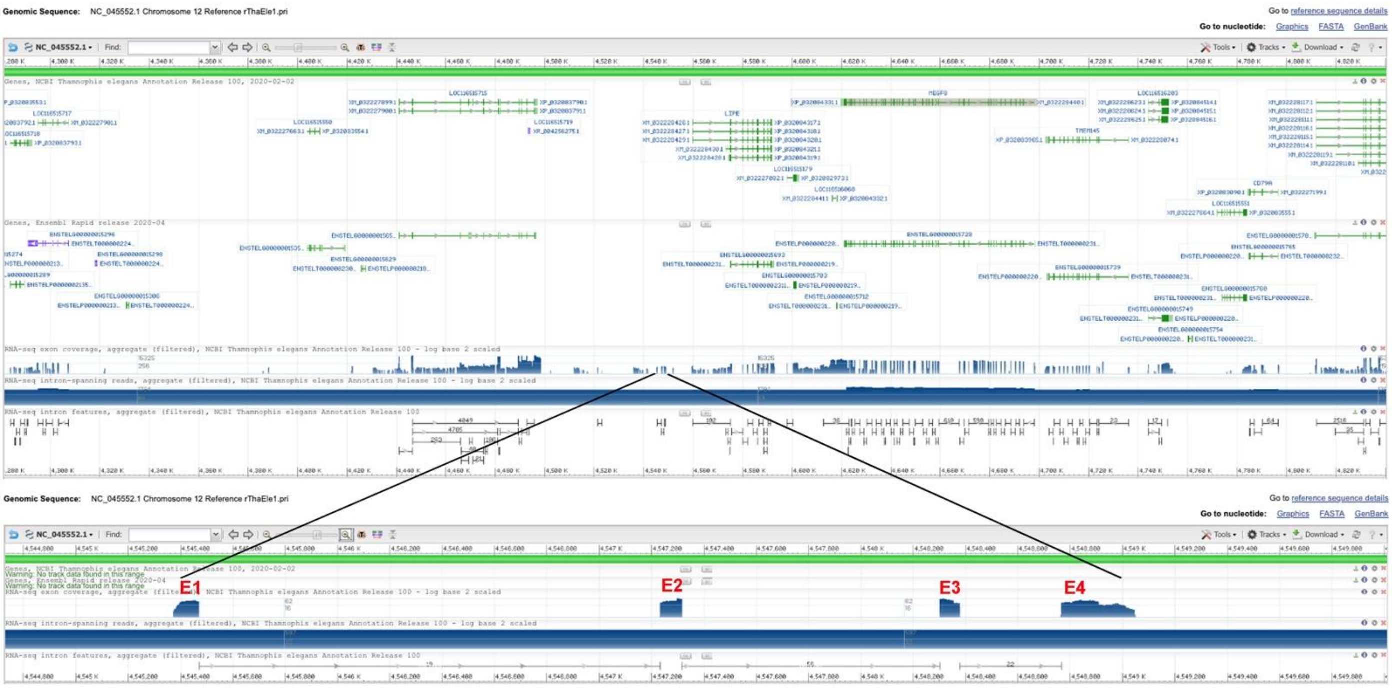


**B:** Sequence of genomic DNA (exons in red and highlighted in yellow, coding region underlined)

ggagtttgcacaatggctcggggattgccctgatccgattttcttatttttacggaagccttttctagaaatattaccggttgggaggagagattaaaatgcaaacaaaaaaacaaaacgaaaaaaaccagtttggttggttcaagtagtagtggccactgacggaca
cagcgaaggacgtttactatcgaaatggtccgattcttcatctttctggtgttagtggtcctccttggcttggttggttgctcagaagaaggtaagaggatttggaaagggttcttggagatggttgcgtgagaaggaaggagaggaagatccctggacaagcagtc
tcccctcaacaccattcgcttggtcattaagctggaagctttcaagagactgaaatgcccactatcaaaaatggtgcagttgaatctatggcatacgtgccacaggtggcacacacggagccatatttggtggcacgccagcctgctgatttttcacccttctggag
ggccaggagtggttgttttcaccctcaccaggcttcagggaagcctctggaggctggggagggcgaaaaagggccgaaagagaagttcgggagggagggagggaaaggaaggagggagaagaggggaaggagggagggaggaagagaggaagg
aagagagggaaaggaaggaaggagaagaggggggagggaggaagagagggaaaggaaggagaagaggggaaggaaggaggaagaaagagaggaaggaagagaggaaaaggaaggagaagaggggaaggaaggaaggagagaagaaggg
aggaagaaagggaaagggaggaaggagaagaggggaaggagggagggagtaagagagggaaaggaaggagaagaggggaaggaaggaaggaggaagaaagagaggaagggaggaaaggaaggaaggagaagaggggaaggagggagg
gaggaagagaggaaggaagagagagaaaggaaggaaggagaagaggggaaggaaggagggaggaagagagggaaaggaaggagggagaagaggggaaggagggaggaaggaagagagggaaaggaaggagaagaggggaaggaaggag
gaagaaagaggagaagagggaaggaaggagggatgaagagagggaaaagaaagaagagaagagaggaaggaaggaggaagaaagagaggaaggaagagagggaaggg aaggagggagaagagggg aaggaaggagggaggaagagag
ggaaaggaaggagggagaagaggggaaggagggaggaaggaagagagggaaaggaaggagaagaggggaaggaaggaggaagaaagaggagaagagggaaggaaggagggatgaagagagggaaaagaaagaagagaagaggggaag
gaaggaggaagaaagagaggaaggaagaagggaagggaaggagggagaagaggggaaggaaggagggaggaagagagggaaaggaaggagggagaagaggggaaggaaggaggaagaaagagaaggagaagagggaaggaaggaggg
atgaagagagggaaaagaaagaagagaagagggggaaggaaggaggaagaaagagaggaaggaagagagggaaaggaaggagggagaagaaggaaaggagggagggaaaaagattgacacagtgaatggatccatgtacctgacctagcatgga
aactgtttttaatttggtagctggaggtaaaagatttggtctgcactcctccgtagttctcccaggtcccccttttctcagtcccccccccctttctcacctgttcccaatgggatggtgtgaaagaaaaagtgacaaactggaagatcgcccccaaagggggcatcca
cctcctccacctgggttgaagggagcggggacagtggctcatcctttctccaggagcgccaagtccacaaaaatcttcctgaaccacgtcgacggtggcttctgtcgtccctcgtagccccttccaacctcctcttgtctttctccttttgggggggccagatggaa
gggacggcgaagggaaagagagtttgctgcgcaggtcgaaacgaagcaggttggggcatccgaactgcccttgtccaggtgagcctcgtcttttgatttccatcggggagcgggaagaaaatttgcctgcaagattactttcggtttcttctggcgcaaggattg
tgtggttgtgtgttcataggttgtctccttggggtttttttttctccctctgttcttagggccctggctcaaaagcaaacaagctttgttgttaatcagggttcctcttttgagggttgcgtttcttcaaggggtgacctcaggggtgggaccggtttgggcaaactggatgct
agttttacatctgattcaccgaactggtagttggaaggactggctggccccgcccgcccctctccttccctctaggagtctcaacgcggcccgttcatcatgccaggtaactgcagggcctgtgtggaggctctgggagggccaaaaaatgggcctccagaaga
ccctccgggcccgtttctggatcccggagggcctccagagcctgggggaggctatttccaccccctccccggaggcttaaggacagcctccagagcctggggagggtgtaaacgctgcccccccccatggtgcaggaggcccgagtaggctatgcccacc
atagccaagcccacccagcaaccgggccgagaaccggttgctgaaatttttcaatcccaccgctgagtgatctgtggatcaggaggaggtacctctttttcttcctctgatgtgccatgttttctcttcctgacttccttccaatcatcttcctttagtaatcccttgcagg
gtggagtctgggcaactgaatggagcgagcagagtgtttcctggccggatgccctccactgtcgccagtatggagttttgttcagcagatatattctcagtgtgcccaaagagagaaatatctgcctcgacctaggattgaactcacggcctcctgatggtgaggc
gggaggttgacttctaggccaccgcaccacccttctaatgacggttcctaattgcttttagatctagatggactccgacgcccaaagagacgcagccctgagtgtccctgtgatggattcagcttacgtagtttgtcaggtaagacaatactcctgggtgtaactcgt
gtgaaagtaaaggaagatggatccgtgggtgtagcatttccgtaggtgtggaagaaagggagaaatcagaaactacccatcgggatgtttctagaaccctggcttgttttaattgtgtccgcccattaaaaccactttttatggggcaggatactacacagccgccg
tggttcacttaacaactgtggcaagacagggcgtacaacaggacaaagctcccttatcaacggtctcgcttagccgcagaaatacaagttctggtcgtaagtggaggattatcctctacctttctgcctccaactgacccattaaaagatgcattgttttattttattttttt
tcccacgttccctttcagaacatcacaatcaacgtcattctcgcaagagcaagcatcacgtgtcccataagaacaaatgtaggaggtttttgaaacactgcgcttttaaagggatccatgtccctttatagtttatcctggttttctcggcaacacccaaggggattttttt
atccaaggtgcaatgttgattcttgcgcctttctgatttttttaaaatatccttgtttgctctaggttggttggaaataaagctctgaatttggggcctttgtgggtgtgtgaatggcctgtgtaaagaattgtgtggacaaataaaaacaataatgaaacttgctagacacctg
cttcgcacattttgcaaccccttgtaaaacccggaga

**C:** Sequence of cDNA (coding region in red and highlighted in yellow)

gttcaagtagtagtggccactgacggacacagcgaaggacgtttactatcgaaatggtccgattcttcatctttctggtgttagtggtcctccttggcttggttggttgctcagaagaagatggaagggacggcgaagggaaagagagtttgctgcgcaggtcgaa
acgaagcaggttggggcatccgaactgcccttgtccagatctagatggactccgacgcccaaagagacgcagccctgagtgtccctgtgatggattcagcttacgtagtttgtcagaacatcacaatcaacgtcattctcgcaagagcaagcatcacgtgtccca
taagaacaaatgtaggaggtttttgaaacactgcgcttttaaagggatccatgtccctttatagtttatcctggttttctcggcaacacccaaggggatttttttatccaaggtgcaatgttgattcttgcgcctttctgattttttaaaa

**D:** Amino acid sequence (signal peptide shaded)

MVRFFIFLVLVVLLGLVGCSEEDGRDGEGKESLLRRSKRSRLGHPNCPCPDLDGLRRPKRRSPECPCDGFSLRSLSEHHNQRHSRKSKHHVSHKNKCRRFLKHCAFKGIHVPL

**Fig. S12.** Gene synteny and architecture (**A**), genomic DNA sequence (**B**), cDNA sequence (**C**), and encoded protein sequence (**D**) of the unannotated *cxcl17* gene in the NCBI reference genome of *Thamnophis elegans* (Western terrestrial garter snake). The information was downloaded from the NCBI gene database (https://www.ncbi.nlm.nih.gov/gene).

**A:** Gene synteny and architecture

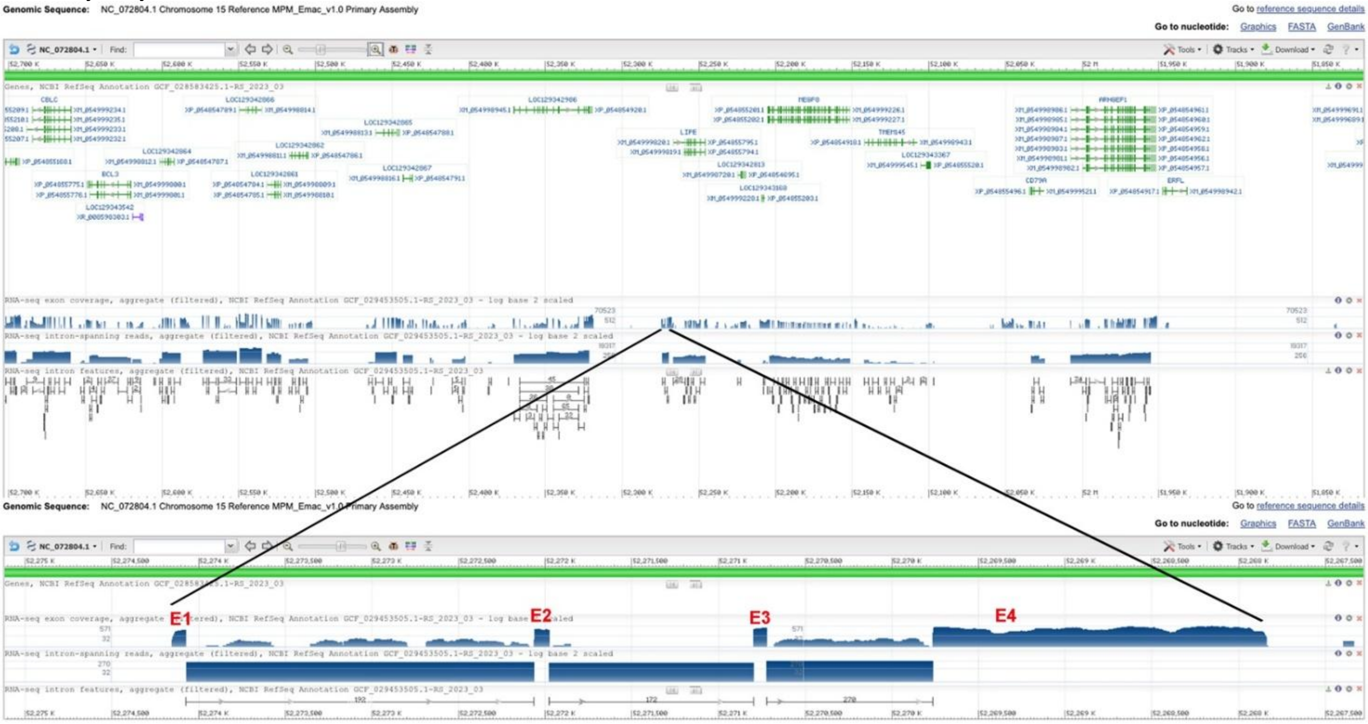


**B:** Sequence of genomic DNA (exons in red and highlighted in yellow, coding region underlined)

agatcctcttacaggaaacttttccgcaccgctgctcctggggattgcccccttttttactctgagccttcccaaaaatgacatcaggagaaggccctaatcataaaaggctagaagtgaaagctgcgctggacattgccactaaaatgggagggttgctagtctgg
acgctgctgctgctcttgctatctgtcagcatctttggctcctcaaccagggggaaaggtgagtgaaaaggagaatctgggtcccggagcattttcaaggaggaggagggactctcgtccctttaactcattccacattatggtccgccccgcctgcacatatttag
caggggagagaacagttccttttattaccacaacagagctggactcagggttgccaactccagcttgggaaattcttgagatttgggtgtggagcctggaggagggcagggtttggggagggtagagtccaccctccaaagcagccgttttctccagaggaact
gatctctgtagtctagagacgagttgtgatttcagatctccaggcctcacctggaggttggcaatgctacctggaatggcggaagttgactgtggggttgagacggaccaaggaagtgctttccccacacctaacatgcactacttcctgtatagtacagacctgtc
cagtcttgcaggtcaccatccaaggccttctgcagcatgaatggaaaagagatactgtctgctacaggcggggcctttgctgtggtggcacctcccttgggaaattcctcctctgcagagattcatccagcacctactctttggaatattaggtgtccaagaaagga
agttcccatctaggcctttgaagggtgcttgcatttttgtggatttgttttgatgacttcaaatctcgttttaatttggctgttttagtcatttgtactttcgcctggattgtttaaagccttctttgctgttttaatttgctgcttttcctctccctgcattctgaactgtcttgaactttgg
tggagaggctgtccagaaacgaagaagcatgaaatgaacttgtggaactcactgccacaagatgcagtgctggctacagctgatttattgcttactgaatcaaaacttctacatgttatctttcgtctgcttggcagcaatatgtaaaattacttttccatagctggtatta
ctactatttaaaaaaaagagagaaaaccctctgctggcgctcttcccaccaaggccaggcagcattaggtgttggtggtaggcttctccttcctggcagatacagtttgtagctcctgatttttcctttttaattgtgactggtgaaagcacaggagcaggggagataat
gtaaaaatgccccccttctcccaattattcttagagatcagacaaaaaaattcatggaagcagtttgtatctatgtctgttaaccacaatccttaaaaggaaccttcatgttcagaggcagtatactctgaatgttaaatgctaggggaaaggcatgtgactggccgctgct
ggaacacagtctcccaatgcatgttatcatattgaggaggggttgggtgggaaagtcatgggaaaatgctgtggaagggtagcggttttaatggtagagtatctgccttccatgcagaaggtcccaggttcagtaacacgatccagtaacaggagatgatgggaa
acctctgtaggtgacccgacagaactactgatagtctgcaaaggaaggctgaccaatttcttggctccggagccctacagtgaatgtatatgcttgtacctgcatgtgtgcatctgtttgcaagaaacagtcaatgtatattcattttacaaagcaaaccaggacca
gtccctagatcagtgtggggtatacatcacacactgagcagctgtgcgtgcattgaacatgattcaactcgtgtatagagaaaaatcaggggtacactggccacagattgtatatgtgtttgctgtaactcgtggacagcactattataaggctgtttcgggtgtttatg
agaatggaaaacccgctcctttcaaattctgttatcccaaggaacatggggcctttctctcctccctacagtgtatctgttaaattttgtcacctgtactttgcctgttgggtcaggccagttctagagaacccttcgtgtgcatgtggttgtgccacctttcggagctgctc
ggctgctaatttatccctgtttctgggtcagatcaaggattgcacgtcggccagacagagcgggagctgagtggatctgaccaaagccagaatgggagtggattctgccgctgcccaggtgagtgccgccgcacctcccgcagccggataaaaggaggataa
ggcccaaattcttgtttttaatttctctgtggaaactggcctggtttgtttgctatccgggtgtgtcttttgtaacttcctgactcttctacagatacctgtggcatcatgccctctgctttgacaattccccagagaggttaaccgtttttccagctagctagctagcgtatttt
ttaccccagtttccttcaaaagccccagggcagggtacatggtttcttccttccttccacagccctggaaggtaggtcaaatagcttcatgccagagcaggatttgaacccaagactctcagatcctaccctgacactctagttaagtttagtttagtttattcggtgttta
gcccaccctccccgcaagcaggctcaggacggggtacagcagtcttaaaataccattccacaacgggataaaatacaatacaataaaatagaaaaaatcaccaattaaaatcatatacacagaatcatagatggctgcttccacatttctgctccccaacatacaa
agagggagggaaacggacaggtaagaggtaatcagatacatacgcaggacagctcataacagtcttaaatagtctcaagctttgcaggggggccggtgagaggctcaatgatggtcctagtaccggcctcaaccatatgcctggcagaacatctctgtcttgca
ggcccgccgaaatgatacaagatcttggcgggcccgggtgtcatcagacaaagagttccaccgggttggggccaggaccgaaaaggccctggccctggtcgaagccaggtgaacattgcactgtcttgatccagcaggggggttctgtttgcacacttcgcctt
gggctggttaagggtggtggaggagagtgccctcaagtcatagccaacttatggcaaccctgggtggggttttcatggcaagagactaacagaggtggtttgccattgcctgcctctgcaaccctgcaattactaatcaaggccggtcctgcttagcttctgagat
ctgtcaagatcaggctcacctgggctatccaggtcaaggcgctggttaaggaggcctgttgatcatcgcattcccccgaattctgacattcttcctgattaactcataccttctgatgtcattttaggctcaccgaggcctcagaagcagaagaaacacaggcatgtg
tgtccctgtgatgacgttgccgtatctcagccaggtaagaaaaatgttctgaagcacccctgatcctggggagggaatgaatagggtcacatcatggactgctctgtcaccagctctgaaaagactgaaagtacttaaagtgatcttccatcagagctccgtccca
gacttttggaatgtcagattcctaatctaagacttacaggaaggaccaccagccctgcagagaactgggaacgtgtcctggctcagccagctttgccaaaggactcacttggtgtgtgtttctggcagacaaaggacaccataaaaaggtccctttgatttctggttt
caaccaggcaggatttttttttcagtgtatgcggcgaagacgtggaatggatgagtgcgggcagcaacaaccccacatcaatagagagagtcatgaagtattttgttgaggtgggggggggagccatgggagaaaagcacaggagcggaagggactctag
ctcttctgggtcaggcagaagaagaatccgcaggaagcatattttgtctgctttccttccagagggtatttttgcccacatctggggtgacagaggctgctggcaaggtgctagaaaaacaagttacctggtgccagaattaagacccacagctctcatgttctaaat
agatgcaagagggaccgaaaggagataggatctactgttcgacgggaagagaaataataacaacaatatttgatttatataccacccttcacgacaacttgaggcccactcagggtggtttccaaagtgggttattgttatgctcacgacaatcaccctgtgaggt
gggtggggctgagagagctctgagagagctgtgactggcccaaggtcacccagctggcttcaagcagaggagtggggaatcaaacctggctctccagattagagtcctgctgctcttaaccactacatcaacatgtgctcggtttcttctgtccccctttcccag
ggcgtcatcgtcgccatcctcacagcaagaacaagcattttcctcgccacaggaagcagaggtgccggacatcccacagcaaatgcaaacaccggcatctaatgaacgggcttcagattccactgtagctctctggggctcctcagtaaagcacgtccaccca
cttcctatagaaagggccttctttctggggtgcatgtcacttctggccttgaaagtgaacagatagagaatggacaactcagaaccgatgctgaaactccagttcaaccataggcttaccaggcctattgctagaacctcccagcctcagtttgtcatctgtaaaatg
ggaatacacaatgtgtgtgtgtatatgtatatatatgtatatgt

**C:** Sequence of cDNA (coding region in red and highlighted in yellow)

Aggctagaagtgaaagctgcgctggacattgccactaaaatgggagggttgctagtctggacgctgctgctgctcttgctatctgtcagcatctttggctcctcaaccagggggaaagatcaaggattgcacgtcggccagacagagcgggagctgagtggat
ctgaccaaagccagaatgggagtggattctgccgctgcccaggctcaccgaggcctcagaagcagaagaaacacaggcatgtgtgtccctgtgatgacgttgccgtatctcagccagggcgtcatcgtcgccatcctcacagcaagaacaagcattttcctcg
ccacaggaagcagaggtgccggacatcccacagcaaatgcaaacaccggcatctaatgaacgggcttcagattccactgtagctctctggggctcctcagtaaagcacgtccacccacttcctatagaaagggccttctttctgggg

**D:** Amino acid sequence (signal peptide shaded)

MGGLLVWTLLLLLLSVSIFGSSTRGKDQGLHVGQTERELSGSDQSQNGSGFCRCPGSPRPQKQKKHRHVCPCDDVAVSQPGRHRRHPHSKNKHFPRHRKQRCRTSHSKCKHRHLMN
GLQIPL

**Fig. S13.** Gene synteny and architecture (**A**), genomic DNA sequence (**B**), cDNA sequence (**C**), and encoded protein sequence (**D**) of the unannotated *cxcl17* gene in the NCBI reference genome of *Eublepharis macularius*. The information was downloaded from the NCBI gene database (https://www.ncbi.nlm.nih.gov/gene).

**A:** Gene synteny and architecture

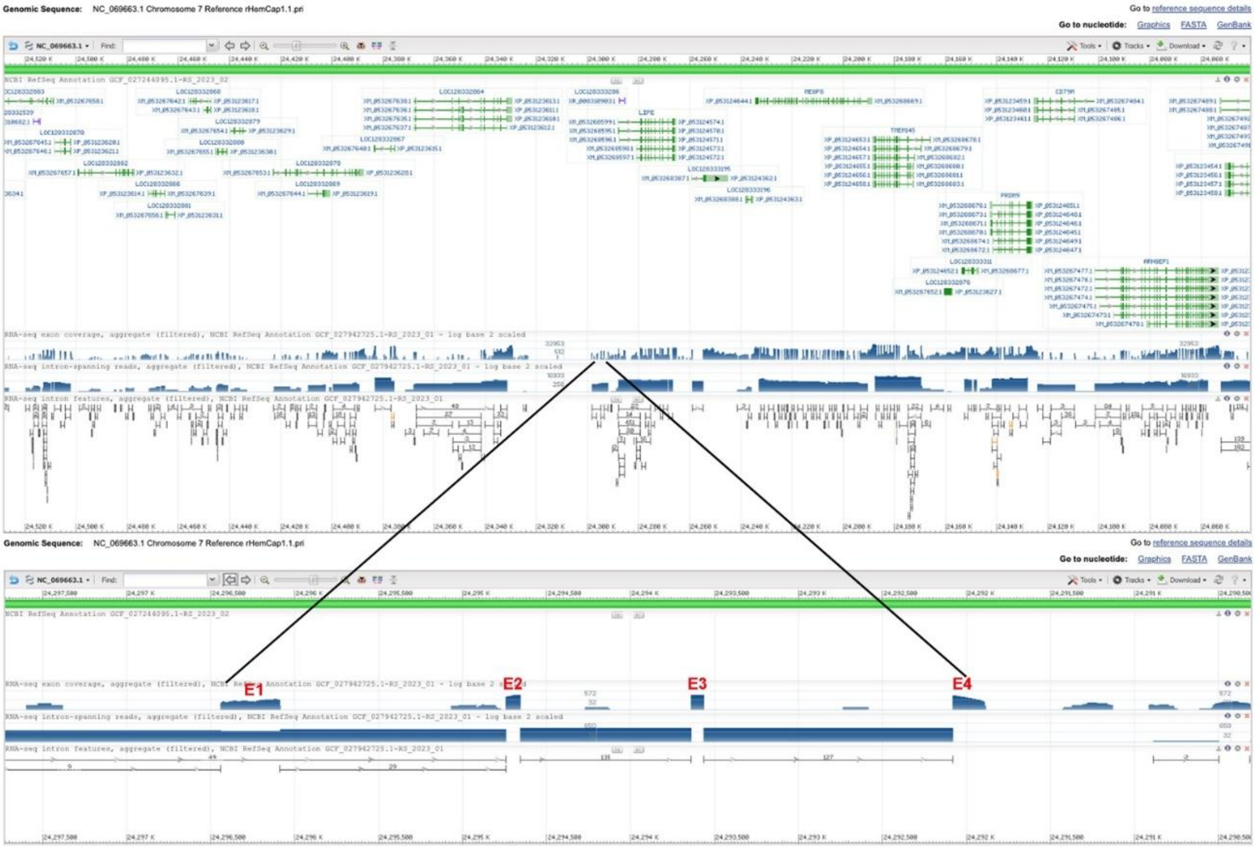


**B:** Sequence of genomic DNA (exons in red and highlighted in yellow, coding region underlined)

gtgccagggaccctggggattgccccgaagagtgcttttctccttaatccttcccataaatgacatcagaaggagcattaagggaggagcccatctactaatataagcggggaaaccctgaatgatagaggtgcactccagagacccttgcagagacacagct
gcccagatggacaggttcttcgccttgacactactgttgctgctgctgcagtctgtcagcgtcacttgctcagccagaaagaaaggtgagaggaaggatggggtgggggagatcggagggacccttgtaggtataatgcccccccccccgcccctggctgtg
ctactttggttacccgcttcctctgcctctctctgttcctgtgctgttagcagcaccctgacatgcagaaatttaacaagggaacccccttcctctccctcagtaattgagcttgtgtggggggagtgataaagaaaagggttgccagttctaacggaagctattcctgga
gatattgccccctccctgcaataatttatccaatatgtttcataacaacaagctattaaaatctctcagaatgcactgcattcaatagacatttgggaagtgatgctcattcctgaagacagagggttgggaaaaatcggaaagaaccctgttaactggacaaagaggc
agctttttaaagtgggggctagcttatctttagcagggagagagcaactatccctgttcaggccagcatagcgtccttccccatcaccttctatccatttggccattgtggctgggagtgatgggaattgtagtccaacaatttctcgcagaccccctagcctgcagtcc
tgattctcctgtggtcttttcccactctgtctggagtggctgctgatggtaatgtgtgtgttagattgtgagccccttttggaatagggaatgatctcctcatccctttagctatgtaaactgctttgagaactgttgtttgttgaaaagtggtatacaaatattctaaataaaagt
aaagatgccaccttgccgtccttgatcgtgatcactttagtttagacaacggacttgcagggttgccaaattaagcaaggattagagttcctctgcctttaataggtacagggaagaaagatttgggaggcacacagctatctccccctgccggggataaacaattgt
gggagaagctgcacttgttaaaattctcgcttaacttgcatgtcagggctccagggaaagtgaggggaaactcagtgcatgggagacatacacaggcttacggttgtgaataaaagccagcacgtggccagggggcaactcttctgcttagtatgcagaaggc
ctgctaattccgtccctggcatgagagcccatgtaaagtgtttctggaagcccattccaaagggtgcctttactgaacattacaaggattttttttacaaggcagaatgtgaatatcgcccagagggagagggaaattctcctctttgggaatctattgtcttgagccac
caccttcttgcgatacacggaggctgtgtgtctgtgatattacaactgatccgctggtaatatttctcctttctgcatcagatggagggctacaggttgatcaggcagtggcaaagctgagccgatctagacggagctacgatgggaacaaactctgcccctgtcca
ggtgtgtctcttccccatgcctacagctgaggaaatacatgctgaaagaactctgccaaagctgcctttgctttcccttttagcaactgatgtatgctggctgttcttttcaaaaccttttgcctctctccagatgctttctttcttgcgtgatgttccatgctgagccctttct
gggacaggttagcccttgtttgaaggatgctactcaatggggccctcctcttggggttattccagtgtgaaatgtaggtctgtgtcttgcacttggacacactacgctccacactctggtcacaccccctacagccgctctcagtcatgatagcaactaatttcccccag
ggagttgagaagacagtttggggggggaaactttgtttgtgctcaaaattagggtgaatatattttgatatacacactctccattcctctgtaggttaactgggatttctcatctctattagtgattgtatggatacagggtggtgttctggattaattcagagtataaatgtgt
ggtcttgcataggtagcccaagagcctgagccattttggaagggcggtataaaaatcaaataaataaaataaaataataaataaataaaagagattgtgtagagggatggggatgctgatggaaatagcggtgctatggaaaaagggggaagcagttctttagttat
atggcagagcaggggttcccagctatgagaccccagatgttgatggactacaacgtccatcatcccctgccacaatggccttcagtggggatgatgggagctgtagtcccaacaaaatctggggagccaaagttgggagcccctgctgtaatgaactgaaagg
tgatattttaacactaagcaaacagaatattgtttttgatttgcatgagttattctctgaggtgctatatctgtctgcattgctagttaagggggggcccacggatggtctcttgcctgggggccactctctctctctccctctctctcttcaacgattcatatcattttaggcttg
catggactccataggcagaatcatcaccagcataggtgtccctgtgatggcctcaagctgtatctgccaggtaagaaaacattcccttaggtgtagccaatgatgtggagaggatgtttggggatgacatccctcaatttgatagaggaggaacaattatcttgtatc
ctaggcagctagaggagtgttgatggaaggcagacagcacccatcccagttgccttcatcctcttggttccctgccaggattgctctcagttttggaaggtgtgccccaggtcctccatcatctacagaagggtggaaggactgagagagagagcttgtcgggac
tcccatcagtagttctggcatctccggagctacctgttgtgttgaaattgccacagctgcagctctgcggggactgaagagtggatgaagagacgttctgtcattgctcatcccagacacttcacttgtcctgaaatggctgccctggtctcctgctgggcttgccatc
agctctgcagaactgggggtcttcacagcagaggatactgagcatatacccgcttctctgaactgagcttctcagagatacaggccaaagacctgagtcaagtaggggtgacaagagatggtgatctaaactgccagagccaggtggcatccacccaagagtt
ctaaaggacacaaagtgaaattgctaatctcttagcataaatatgtaacgtgtccctactagcaggctctgttctagaggactggaaagtagccagtgcaacaccggttttcaaaaagggatcgggggtggatctggggaattacaggccagttagcttaaaatctg
tcccaggtaagttgatggaaagcatacttaaggataaaatgggatctactgttggaggagaagagataaactgtaggggtgtgcaaagttgggtcgaggaggttggacctggtctgaaatgggccaggctggagggtgtcagagagacttggaccctgaccta
agaaagcctagttcagatcagagttttacaaccctgttaggagaaaactagagccctgctccaaatggtggctggaggtgtttttgtatgtgattgcagttggggtgagtgggagacggagacctcttctgtcctctctacctgccagcctgcccaaacatttctttct
gtcttttaaaaactttcttaggaagtttttaacttgcagtttaaaagcggccgatttccaggtttcccttgataccctccccctgcaattcctccccctgcaattcccggctcttaagtcacttcccctgcaattctctggtgttaggtaaattttcttccaggtttcacagcga
gtatcctggtactcttaagaattaatttagaggatggtcaggaaaatacccaatcctgaccccgaccctaacattcaaaagggaagttttgaaacatttgtgggatgttctgaaaacatttatgggattgggtcagatctggtgcacaatagtgcacactgctaaaaaa
accacccccaaacatgattgttctttgtttctccacctgcagaacactctcgccatcatcgataccgtcatggcagaagcagtcggcacctgaactccaggaggcaagagtgccagaagttcctgaaattctgccaagatcccatgcgggggctccagattcctct
ctagctggtcaccacccaaaaggggtttgaacttgagggctcagcaataaagaaaattcattttcttagagaacgagccttttttttcccttttttttttttttttttgctgtgtg

**C:** Sequence of cDNA (coding region in red and highlighted in yellow

gggaaaccctgaatgatagaggtgcactccagagacccttgcagagacacagctgcccagatggacaggttcttcgccttgacactactgttgctgctgctgcagtctgtcagcgtcacttgctcagccagaaagaaagatggagggctacaggttgatcagg
cagtggcaaagctgagccgatctagacggagctacgatgggaacaaactctgcccctgtccaggcttgcatggactccataggcagaatcatcaccagcataggtgtccctgtgatggcctcaagctgtatctgccagaacactctcgccatcatcgataccgt
catggcagaagcagtcggcacctgaactccaggaggcaagagtgccagaagttcctgaaattctgccaagatcccatgcgggggctccagattcctctctagctggtcaccacccaaaaggggtttgaacttgagggctcagcaataaag

**D:** Amino acid sequence (signal peptide shaded)

MDRFFALTLLLLLLQSVSVTCSARKKDGGLQVDQAVAKLSRSRRSYDGNKLCPCPGLHGLHRQNHHQHRCPCDGLKLYLPEHSRHHRYRHGRSSRHLNSRRQECQKFLKFCQDPMR
GLQIPL

**Fig. S14.** Gene synteny and architecture (**A**), genomic DNA sequence (**B**), cDNA sequence (**C**), and encoded protein sequence (**D**) of the unannotated *cxcl17* gene in the NCBI reference genome of *Hemicordylus capensis* (graceful crag lizard). The information was downloaded from the NCBI gene database (https://www.ncbi.nlm.nih.gov/gene).

**A:** Gene synteny and architecture

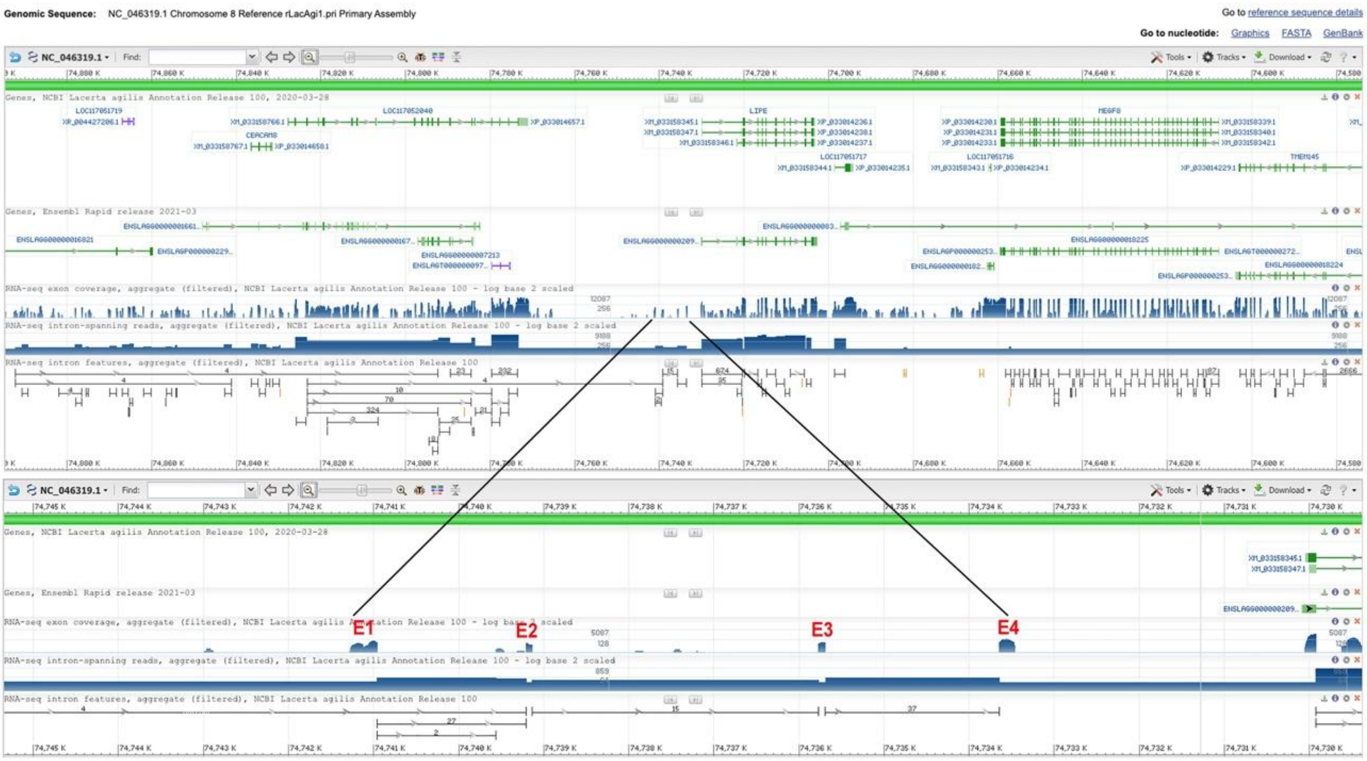


**B:** Sequence of genomic DNA (exons in red and highlighted in yellow, coding region underlined)

cccacctcttggccccaggtgagggtgagggttggccttggactggggggggggagggagggaccctcccctcttcttttgtatttttatcctctgatagctttgctttcattcaaaaaattccccaagctgaaactcgaagcccaatggtccgggaagtaatatgat
ccactctttcttgtttggcagacaccttttgtgtctgtcgctctgatctgaatttggcaagccacgctatttgccaactccttctgccaggtacaaaagctttatgtgtatacggatcgagaggagatttcccatatcacaaaatatgcatttcgaaacttctaggagcttttg
aaagcagctctctccaaataaacaaacatgttacaggagagcatttctgctccagtgatccagggcctgccccgaaataatccctttcttccccaactcttcccgtaaatgacatcaagcataaaggcagtcgactcaaacagcagcgcaagaggagagatccct
cgccgctacaaatttgtgccttgcagacctccgcggacgccgctctttgccatgggccgattcctagctttgactctcctgctcctgtcgctcagcatcctttgctgcacggccaaggataaaggtgagagggaaggatggaaactgggaggaggaggaatcgg
tacgtcaattcttccccctttggtgggcgcagttgaactgcggaacccgctgccgcaggacgcagcaacggccaccaatttggatggtttcggaaaaggattggacagatttgtggaggagacgtctgctggtggctactagccatgatggctgtgctctgcttcc
atggctggaggcagtaatgcaggggcggaccctggtaataggctgccctgtgtgaggccaaggctttgggaaggaggcatgaggttctggcaaggtcctagagccctcctacctctttccccaagcccagtaagggctttgcttgcccagctttgggaaggag
gcaggagccctcactgggctttgagaaggcaccttccttggctgcacgcccagtgtgatggcaccgtttgcgctgccgtaaatccacctctgtagtaatgctcctgaatgccagttgctgggaggcacagcaggggggagagtatttttgtgtttggatcctgcttgc
cggattcccacaggcatctggttggcccctgtgggaacaggatgctggactaggtgggctattggcctgatcttgcaggtcccagtggcatagcatgggttgccagtgccctgggccgcgtggagggagggaaatggatggagtacacacatgcattcagttg
cctaacagcctccgtgtcacaaaagagattgcagccgcagagataagaaaagaagagtcattccattgtctagagaggttgcttcttggttctcatggaaaggggtcagcatggcacagaacatcctgctttgccagccacaggtgccaggctcaatcccttgcc
cttcccagctcagggtggtcctgggttgggagggcttgaggtttgccgctgcatgagatgctggagaaacgctgctggaataagctagccttccctagtctaggactcttcatatagaatcacagagtttccacttcatgggtgagtggggattcgaacccagatc
ctagtctgacattccaactattacaccacactgactgaaacgcacatatcagctaagcaaaaacagtcaaggaaggcactgagcaggaccggtaaggggcatggtctggcgaaatgtaatgaccaaggagaatcctgggggccagacagagatgtctgggg
gccgcattaagctgccaaggccccccccccatgatataggaggtggtaagataagaaacaagtctcccgtccctttctcccccaggaccatccttggatggccatctgtggtggatgctttagctgagattcctgcattgcagagggttggactagatggccccc
aggcaccctttccgattctatggttcttctcccctctccagacacctgcgcccctcccaaaacctttccagagggtggtgggctccttctcaatgaggaatgcagcaggagcgggcagagacggctgcgagagggaagagtcacccagaataggtcttggggc
gtaaggaagttcatggcagcaaaaaatccaccatcttctttcctccagcattgttcagaagagtctcccaagtggctttgcagagggcacagtgtgaaggagaaggcggaaacttatccttctgtgtctcgaaatctaacttgctccgtttaaataacccgccctcgtt
ctcagagtagctgtcttgctgttcggagtttacccaattgaccatcgttcccttttctctgccaggtagaagattccgttcaggccacaaggagggagagccagcccaaggaggccaccgtccctgcagttgttccggtgagtctctttcttgcaccaccaaccccg
acagttgagaatatggaaactctcccccccccccagttccgctttccgctttcggtttcctctgcagcagctggcccggcctgtttgttgctgaggtgcggcattcaaaacaagatgtggccagtaacctcttgcacaaatcagtctggctcatcgcttggtttgagagt
gccacaaaccaaacccgagtccccccccactgcaacctgactctgaaatttcccagtgtgaaaaacggagatggatgatggtcatcgtccgttcatttagaacaggggtggccaactcccaagagactgcgatctactcacagagttaaaaactggcagtgatct
acccccttttgagctttttaaagaaggaggaaggcccatttggggggggggggggttcgggtcaaagttgttgagtttttcaggaaggaggtaaaaggttgagctttttaggggagccacagttgttcagcttctttgggggggagccaatgatctaccagtgat
ctaccacagacgtccagtgatctaccggtagatcaggatctacctgttggacatgcctgattagaacaattaatgttgaggattaataatgcagcaatacgccgcctaacgccaaattaggcttcggagtggtcttattaaaaaaaaacaccaaccaattacgcaag
catttaaatataaagcacccataattaaaatgcacagacaatcccattagcaattaaaatgtgagcatccgtagttaaaatatgcaacgaagcaaaccgatgcagtaattaaagtagggaaagccggtgtatttatttatttatttatttattattgtgatacattatataatt
ccatttactcataactcacacatacacatataacacacatccaattacattcatacaattacaattacaataatacaattataatagttacaataccttataccaatacatatacatatatccaaatacttttatatcttccaaactcattttagaccaattatatttttattacataga
ttatttttaaaaatataaacttgttctttctagccttccgcttaccacggtatgactaaattgttaaataatgtccatcttcgacttccctgtccatctcttcattaaaatatgtaagaaaattgttcaccataaataaaattataacaccgacctcttttactttatttcttatatttcttg
atggtttaatcttgtgtgctttttattcctatgtcgttttccaaataacaggggttattccaaagctagtactgtgttaatgctgcatattttttttaggtattgcaaaaagctgctccattcttcaatgaacttttctctcggattgtcttttatttttctgagaaatatgcaagctct
atatattctaaagttttgtataaccattccctcctggttggtattattgcacttttccacttagatccaattaagatttttgccgctgttgaggcatataagaaaaaaattcctcatttttcttggtatgtctggtatagttatacctatgaggtaagcctctggtgttttagggaatttg
gttccaaaaaatttcttccatttctgaatgtatttcattccagaaatttgctattttgccacactcccaccagacatggtaccagtttccaacttccttacacctccaacagatattactcacatttttaaacatttttgagagcttatagggagtgaaatgccatctataaaacatc
tttatcatattctctctaatcgaattgcatgttgtgaagctccaactagatctccatagattttcccatcttaccatcattatattgtggcctacatcctgagcccaatgcgtcatagtggacttaatttgctcgtccattactgtccagtccaataatagcttatatgcttttgaaa
gggttttagttttagaatttactacttctatttgaaatttagatggccctttttcaaaacctaatttactatccattttatacctttcaaaaagttggtaatatgtaaaccaatcatttactgtgccttttagactctcatatgattttagttctaatttattattcttttcctccaataagtcc
tgatatgtcagccattagtgagtgaggtgaggccgggactcggcatggtctactatggattaactcagaacataacaatggaacctcatctatggagaatggggaaagctctcctttgtgtgactaatctatggccctccagatgtcatttatggccctccagctccta
tcatccccggccaatggccatgctggctggagctgatgggagttgcagaacatctggagaaccacccctggttcactcccacagtggagaattggaagaggatggggcagcggaggaggatttgcatgtggtttggctcgctttcacgagacatctctgcagt
caacacccctaacataggaacccaagaagagccaatggcctatttagtccaacatccttcttctcacgcccagaggcctatgtgagaaaccctctatcgggacctgagcacaagagcaccctcccctctcgtggtttccggcaactggtattcagaagtgttgctg
gctccaaccgtggaagaaccagtagccattgtctgccctctccctgaatttgtccagtcctcctttaaagccatctaagttggtggctcagggacgcaggtggcactgtgggttaaaccactgtgccgcttgggcttgccaatcgaaaggtcggcggttcgaatcc
ccgcgacggggtgagctcccgttgttcggtcccagctcctgccaacctagcagttcgaaagcacacaatgcaattagataaataggtaccactctggtgggaaggtaaacagtgtttccgtgcgctgctctggttttgccagtagcggcttagtcatgctggccac
atgacccggaaaaactgtctgcggaagcggacaaacgccagctctccctcggcctgtaaagtgagatgagcgccgcaaccccagagacattcgcaactgaacttaattgtcagggttccctttacctttaccttttaagttggtggctatgattgcatcatgtggca
aggagttccgcagtttaactatgtgcatacagtttctttggatgtccacaagttctaaggttaggagcgctctgcaatattctgcttgcgtgccctaaccctttctggttatggggtgggaggtggggaggaccataaaagcactcctccatcccaaatcttgcacttgc
ctcttcccgctgccagttaccatacctaattcctcatcctttgttggcttatgttgtttcagacctgcatgggccccggaatccaaagaaacacggacggctgtgtccttgtgatggcatccttaccggtttgtctcttccaggtaacaaaatacctttagacagggctgct
ggtggagataagatgaatgggaaacatttatcctcccccccccccaaaagaccttaaaggggagagataatttattgtgtatcttaggcaactagaatcttattgatgggagggagagagttgtagtcaacacctgggtcccctaaccagatcagtctcaactctgtat
ggtttgtgcttgtacagtgtctcccatttgtggccctgtgagcttaaatcagctgtgcctaaccattgggaaagggttgtagccgggggcgtagcagggtaaattggtgcccaggggcaaaaaattgtttgtcacccccccccccagacatgggaaggtcaggtggt
acccagtgaggaaaatttcttgtcaaccccccccattgatcccccccccccccgcaaaaatggttttacgttatttcatactttcttacaaaggaataataacaagtaataataaaaaaaacttttatttatatcccaccctccccagctgaagtcgggttcagtgtggctaac
atcagataaataacattgctataaaatcaaacaataattaaattaacctcctaaaaacaactcaaaatcaaattaaagtctaattagatggctttccacagggttagggttgggaacattaagtgttctctgaactgaaatttcagccttcatgacataggaaaacagtca
agtgaacagctattttggtggaggagggtcaagatattaaccgatatgcatgaaatttcatatatagctatataatccctagtatagtaagataagaccttcggagcaggcattaaaaaaaatatatattccttgataatttgtcacccccttcattatggaacccggggc
gacccgcccacccccttgctacgcccctggttgtagctccaagttagagcatctgttttgaatgcagaaggtcccaggttcaatccagggctgggagggatcttgtgaacgagctttatgtgactcctgtaaaaagtgtttgttctacagatcaaagcgaatgaaagc
atacattgggcaggggtaaaactagctttatttcatctgggtcaaagacccagtttggcacactgaccccaagcgcatttgtgtaaacacacacacaaacacacacagaggctgggagcctcaaggggggcccctctggagacttcccggggggggggatgggac
cagctagccccctcccctagctacagcactgcatcaggtaaggtgatctcacaagtcaactgctccctaattgttaagggattaatatactaatagtaacaccttgatttttgcctagtagcaattcagcaattcagatctccgagcagaggtgttttatgtgtgggggt
ggggtggggggctcatacacacttccacttgttccacttgcctataagacaagtgtccgaatggttttccccttgactggctcctttcccagggaaaacctgctctttgcagctagatcggagcaaacgtcaatttggagaaaacccgaattgccatttcctcctgttg
agcactaaagagtggtttcctctggaggaaagcgcgggacaagaggaagtttggataaacccagggtctcgctcctgccaacagaattctgcgctagctgcagcatccaaatcaaatgcaagggaagccccacacagaggtcatagcagttgaattggggta
acctgtgtaaaaggggcaggaacataggaatataccgagtctgaccactggtctatctagctcagtgtttctcaaacttgggtttccagctgtttttgggggggggactacaactcccatcatccctagctagcaggacaagtagtcagggattatgggaattgtagt
ccagaaactacagctgtccaggtattccagagggggccattcccaaccctccttggagatgccaggggttgaaccaaacagctgaattatgcaaagcagatgcttgatcactaatcaagagcatcagaagggaagatataaactgtcctcccccccttttttttgc

aggacagctccatcaccacccccgtggcatggccaagcggcatttgcaccagagaaggcatcggtgccaggcatttgtgaataagtgccacatgatgcgggtcgacatacccttgtagctttctctgcctccaagcgcagggcttaattctttgtgttaaccataa
aacaaatccctcttcggtatctgctgtctttacttcgtttgtttccccacaacggccctgcgaggtaggtgaagctgagaggcagcgaccggcctatgtttacccagggagcttcatggccgagcgggggactgaaccttggtctcgcaggtcctaatccaacaca
ccaaccactaccataacctccaagtatcctaattccccagggacgggttcctgaattacagaagccatcccaattccagaatgccctggtttttccattttcctctcc

**C:** Sequence of cDNA (coding region in red and highlighted in yellow)

tcgactcaaacagcagcgcaagaggagagatccctcgccgctacaaatttgtgccttgcagacctccgcggacgccgctctttgccatgggccgattcctagctttgactctcctgctcctgtcgctcagcatcctttgctgcacggccaaggataaaggtagaa
gattccgttcaggccacaaggagggagagccagcccaaggaggccaccgtccctgcagttgttccgacctgcatgggccccggaatccaaagaaacacggacggctgtgtccttgtgatggcatccttaccggtttgtctcttccaggacagctccatcacca
cccccgtggcatggccaagcggcatttgcaccagagaaggcatcggtgccaggcatttgtgaataagtgccacatgatgcgggtcgacatacccttgtagctttctctgcctccaagcgcagggcttaattctttgtgttaacc

**D:** Amino acid sequence (signal peptide shaded)

MGRFLALTLLLLSLSILCCTAKDKGRRFRSGHKEGEPAQGGHRPCSCSDLHGPRNPKKHGRLCPCDGILTGLSLPGQLHHHPRGMAKRHLHQRRHRCQAFVNKCHMMRVDIPL

**Fig. S15.** Gene synteny and architecture (**A**), genomic DNA sequence (**B**), cDNA sequence (**C**), and encoded protein sequence (**D**) of the unannotated *cxcl17* gene in the NCBI reference genome of *Lacerta agilis* (Sand lizard). The information was downloaded from the NCBI gene database (https://www.ncbi.nlm.nih.gov/gene).

### **A:** Gene synteny and architecture

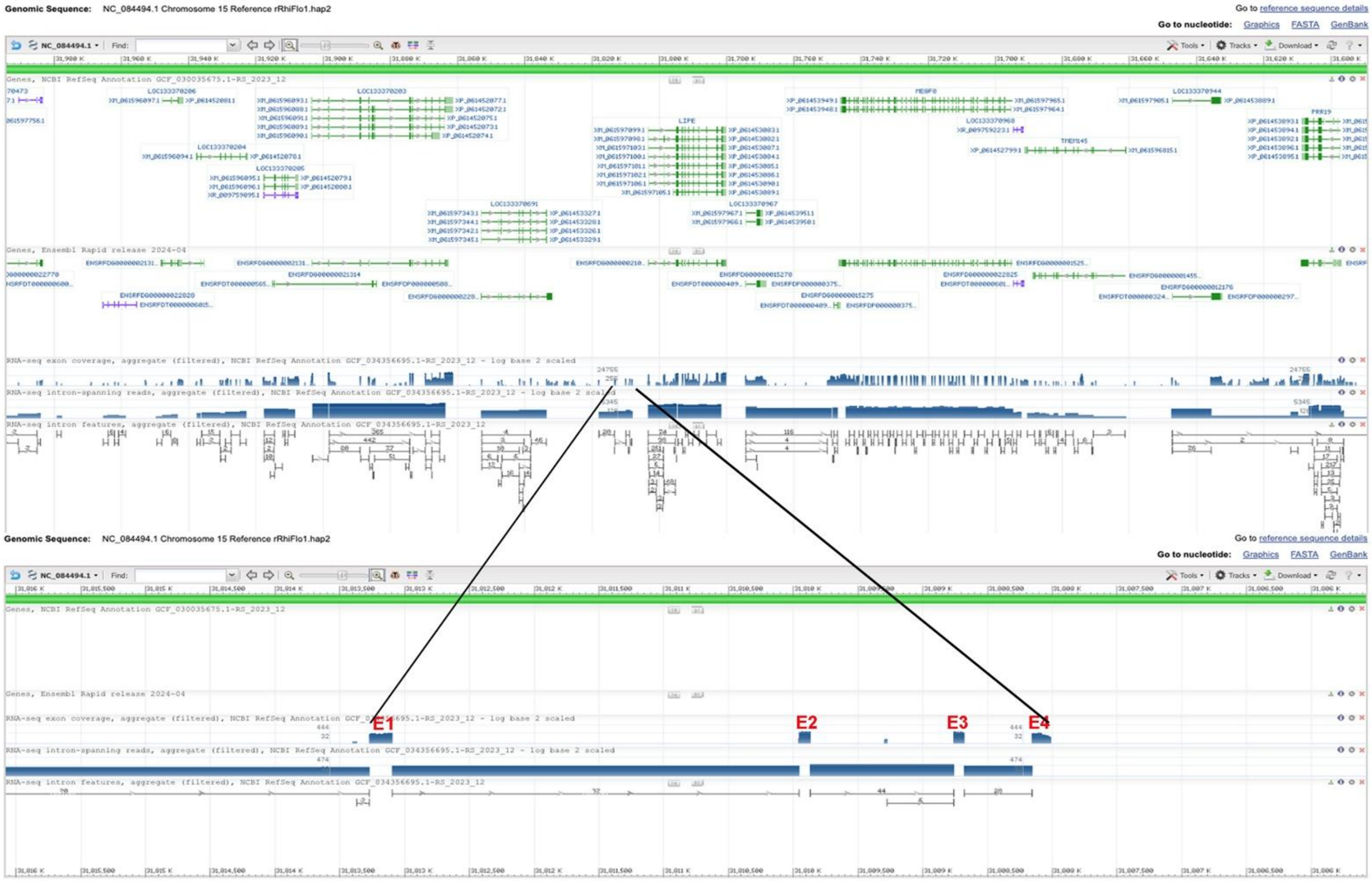


### **B:** Sequence of genomic DNA (exons in red and highlighted in yellow, coding region underlined)

caggaagtaatacaataccttttatttggtttgacaaacatctctttggactctcgctctgatttggctctgatgaagttggcaattcataccttttgcagaccgcaactgtcaagtacaaaaggcatgtgtgtgtggtgggggtgtatgtgcggggaagttcacctttct
cacaggatgtatctccttgaaacttttgaagagctctccaaatagctctccaaataaataaaggtaaggttacaggaaagtgtttctgcaccagctaccgtggtgattgctctgaattaatccttttcttctctaacccttcctgtaaatgacatcaggcgagagacttaag
gaagcagcccaagccgtaccacaagtggaggaatccctcattctctaaggctgggcatcacagacatttgtggatgaggctgctgctgccatgcacagattccgagccttgacccttctgctgttgtcactcagcatcatttactgcttggctaaggagaaaggtg
agcaggaagggtgggagtgggagaaggaggaatcgtggcaccggttcttcccatttcgtggaagacaggcacatccaggtctgatagccacacgcccatctctccaagcactgaactttcaggggttccagttagggctgatgggaggtgtagtccaaaaca
tttggagggcctcaggttggtgacgcctgtactaccatgatgggagaagctatactttgcatgccggggccctacgaagatgaaaaagagctgggtgagtacacaggagacatgaacagtccatcagttgagcggtggttagcagtttattggtctgcttgacgg
agaaaggagcatcatggcagagcatatgttttgcatgccaaaggttccaggctagtcccttgcatgcctgcctaagaaaggctcttgggtagcagggcttggggtatgcctctacctgaggccttggagaattgctgcaggtcactcagagtacaccagccttcc
caacctggggccctccagatgtaggactataattcccatcagccccagctgacaggctgaggctgatgggaattgtagccctaaacatctggagggtcccaagctagggaaggctgtggtagacactagccttccatctccttttgatatttgatgctctatatattg
cagaagtggataaactttatcagtcgatggtcacatgctctggcggagaaccttcatgagggctacatgttgacaatgggcagggccagagcaaaaatcagatattttttctatacactctatatgtgcccacttggctgcttagatctgctgaactggcactttcacg
agcaccacaggtttgtttgttccatgcctgcctgctgatatttggcaaacgccttcactccattgtttttggtgcctgctaaaaccttctttgttccaggaagcctgcttgagcgtgcaattttagcagtgttcagaaattacagattttgttttatattttcatttcccccccccg
tgtaccttgttttcaatgtttgtttcttattggtatttttaaataaatagtttagacagtctgttctagaccacatagagatcttttgttctagtaagtggcgtatacattttcttaagaaaaaaaaaatatcaccatcctccaccaaggcaaggaagaagcattaccatggttcaaa
gagacataccagccaggcaaaatcactcaaggagggcgttgagcaggatcagtgagaggcattgcctgtggaaaaggaatggcctaggaagagtcttgagggccacatctgtcccctgtgtctgaagtccccaccgatgatctattagaacctaagggccca
tccatacgactgtataacctgctatgtattagctttgtgtcctcgttaattcactgttgtccatatgatgttgcaagctagtacggattgtcaggttcacaaatctgcattagaaatagcaccaaaaaccagatatgctttcctaaactgataatcagttgcattaacatccat
gcacccggatgcaatgctgcagactttcctgcaataggcagtccatgggtgttccccccaatcatttcccaagccccttgtctccttcccatccaatagcacatccacaaaccatgcatgggaattaaaaaaaaaaagatatacgtttcggggctcttccaaaaaag
cgcgcaaaaagcaaacgactgttatggaccgatcactgaaaagggatggacttaggggagtagccagtgctaaagcaattcagaccaaaaaccccacaccccaacgtatggatgcttgatgattgggttttcacaataatccgaccacaaacaggacatgtat
ggatcttgagaccatcaaccgaatatggaaaatgcagattttgcggttaggtatgaatggagcctacctaagataagaaactttacagaaggaaaatgtcttgtccgcttctcccccccctctgcagtcacctataccccctctcaaaccctttccagagggtcatgtgg
gggaggctcctattcaatgagagatgaagcattagaagctgctgtgataaggtgaggaacttcagagaagttaaattttccatctcctcctccagcaggccgggagccccgacccgcacattgttcaggaggggaccttgaccctccagcaaccttttcagagg
gcacgggggggggaggaggcaggaaacactctgtaagtgaccctctttaagttcatgtctcaggatctaaatcattatgtagaaataatttgcctgaactcctgaaatacctgtgttgcctgtttcacatgccatgcatgcaccatacatttaaagcacctttaaagtcca
tagctgtctccaaagaatcttgggaactgtagtttacccctcagacagctacaattcccagcacccttaacaaacagcagttcccaggattctttgggggaaatcatgtgcttcgaatgtgcttcaaatgtatttatttaacaaaaagtgtataccacttgattgtaatgaa
acctctaatgcaggggttcctaaactgtggtctgtggaccagcagtagtccactagctttgttcaggtggtgtgtggtctgtctttgaagaatggtggagcagcagctgtgaacggtgggggcagtgactgctccgtcattcttagaacccagcagtccagtccagt
ctgctggctgcaagatgagcaggatagactggaagcgacgtggggtggtgatggtggagaaagaggcacttggaaaaggttctgtggaatttcaattgtaacacaataaagtacaatatacaagaaagaaaagaagaaacaaatacatttaaaaaccaaacag
catctatcacagtgcattacaactgctaacaacaggcagaaaaatcattagggctgcaatccaatacacactttcctgggagtaagtcccactgaacccagtggatcttacttctgaatgtactggattgcactgtaagtggtccgccaagaccatcagcaattttca
agtggtccctgggggaggggagggaaggataggaaggataggaaccactgctctaatgtatggcagggtcagagctgttcagctggtcattctttccccttttctctttcagatggtggtttccatttaggccgcaaggagagggagctgagcagacatcagcca
ggccggcatgggcatgaaccctgcagttgcgctggtgagactgtttccccacagttgcagttgagcaaagggaggtggaggaggaaactgccccatgttctgctttctgtttcctccacaacagctggcaggatttgttagtcattgaagtgtagcactcaactcct
cctacaggtacctcttgcgtaatatttggtgccaaccattatgtgaaagtcacaaatgaaactggaaccgcacccccaccccactgcaatctgattctaaaattttccagtgtaataaagtgaggccattattattattattattattattattattattattattattattatg
ttggtaattaatattgtcacaatgattaagaaatggctagtcactgattaatgtactatttaatgccgaaacaggcttctaagtggttcacaagcatagaaaccagtgacacaagcattgaaatgtacacattcataattcaaatacaacatgaaacaatcccattaaaatt
aaaacagaaagcctccataattaaaaattgcaacagagctattccatgaaagcaacacaggaattaaagcaggagacttgggtgcagagatcaaaggaatgcctgtataaacaggtaagttttcaaaaaccagtggattcaactaggggtttttctaattgccatta
gtgattgtgacaatagatccagcatggtgtgctctggattaattcagaacataaaggtatggccttgtcccaagccaagggaaggcgatactgttcctgctatggagaactatccttggcagcagctaacctctggcccttcaggtgttgctggactacagctccgat
tgtccctgaccagtagccatactggctggagctgatgggagcccaagtatatctggagagccaaaaggtacttcagtcacagcatggagaattaggaaatgattttgggggtgggattgcatgtggtctggctagcttgcatgacccatctctgcagcctgctacc
ctaacccttgctggttaagcctgggggagggcacataaaaggtcactccccaaatccagcacctgcttcttcccactaccacacatacatacgctcaccctcagggtggctcatgtcatttcagacttgcgtggaccccagaaacaaaagaaacatgggcacatg
tgtccctgtgacggcatcagtaacaatttatctctgccaggtaagaaaacgtcattaggcatggctgatgatgaagataagatgaatgggaaatatctttccccccccccttttagggaggaaggataatttatttttgtatcctagacaactggaaggtcattgatggg
agggagagagcagtggtcatcacttgagttccctgccaagatcaccctctcggctctgcagggcatgggcttgcatcctgtttgttgttctgtaggcttaaagcagccgttcctaaccttgggaaacagccatagctcattcgtagagcatctgatttgcatacagaa
gttgccgaatactaagctttatatggagtcacaccattggtccaactagcttgtctgcactgagtggtggctgttctccagggtttcagacaggagatgttccctgccttacttggagatccagggattgagcctgggaccttctaaatgcaaagcagacactctacca
ctaagctacagcatcagaaggggaaatataaactggcctgttttcttcttcttctgtagaacagccccatcctcatcgccatggcaagaccaagcggcaccataaatggcaccagtgccagatgttcctgagacactgctatctgaagcggctccacattccgctgt
agctatctctgcccccaagcgggggattaactctgtgggttatcattaaaacaaatccctcttcttggggtgaaagactttctgttgtttgtctgtatggggttcagaccagaggcattgctaggcacttaaaagatttggggcacacacccatgatatgcaaatttgcat
gaaatttgcacgtgcaaattgatatgcatagataccctttggcaaatttttattgagcattatttatgatttgtgctcacgatgttactggaaagattacagtataagggatcacgcattcagtgccatttgcacctgcaaaagttttggggcagtcgtaccgctgtgtttgta

### **C:** Sequence of cDNA (coding region in red and highlighted in yellow)

gcgagagacttaaggaagcagcccaagccgtaccacaagtggaggaatccctcattctctaaggctgggcatcacagacatttgtggatgaggctgctgctgccatgcacagattccgagccttgacccttctgctgttgtcactcagcatcatttactgcttggct
aaggagaaagatggtggtttccatttaggccgcaaggagagggagctgagcagacatcagccaggccggcatgggcatgaaccctgcagttgcgctgacttgcgtggaccccagaaacaaaagaaacatgggcacatgtgtccctgtgacggcatcagta
acaatttatctctgccagaacagccccatcctcatcgccatggcaagaccaagcggcaccataaatggcaccagtgccagatgttcctgagacactgctatctgaagcggctccacattccgctgtagctatctctgcccccaagcgggggattaactctgtggg
ttatcattaaaacaaatccctcttcttggggtgaaagactttctgttgtttgtctgtatggggttcagaccagaggca

**D:** Amino acid sequence (signal peptide shaded)

MHRFRALTLLLLSLSIIYCLAKEKDGGFHLGRKERELSRHQPGRHGHEPCSCADLRGPQKQKKHGHMCPCDGISNNLSLPEQPHPHRHGKTKRHHKWHQCQMFLRHCYLKRLHIPL

**Fig. S16.** Gene synteny and architecture (**A**), genomic DNA sequence (**B**), cDNA sequence (**C**), and encoded protein sequence (**D**) of the unannotated *cxcl17* gene in the NCBI reference genome of *Rhineura floridana*. The information was downloaded from the NCBI gene database (https://www.ncbi.nlm.nih.gov/gene).

**A:** Gene synteny and architecture

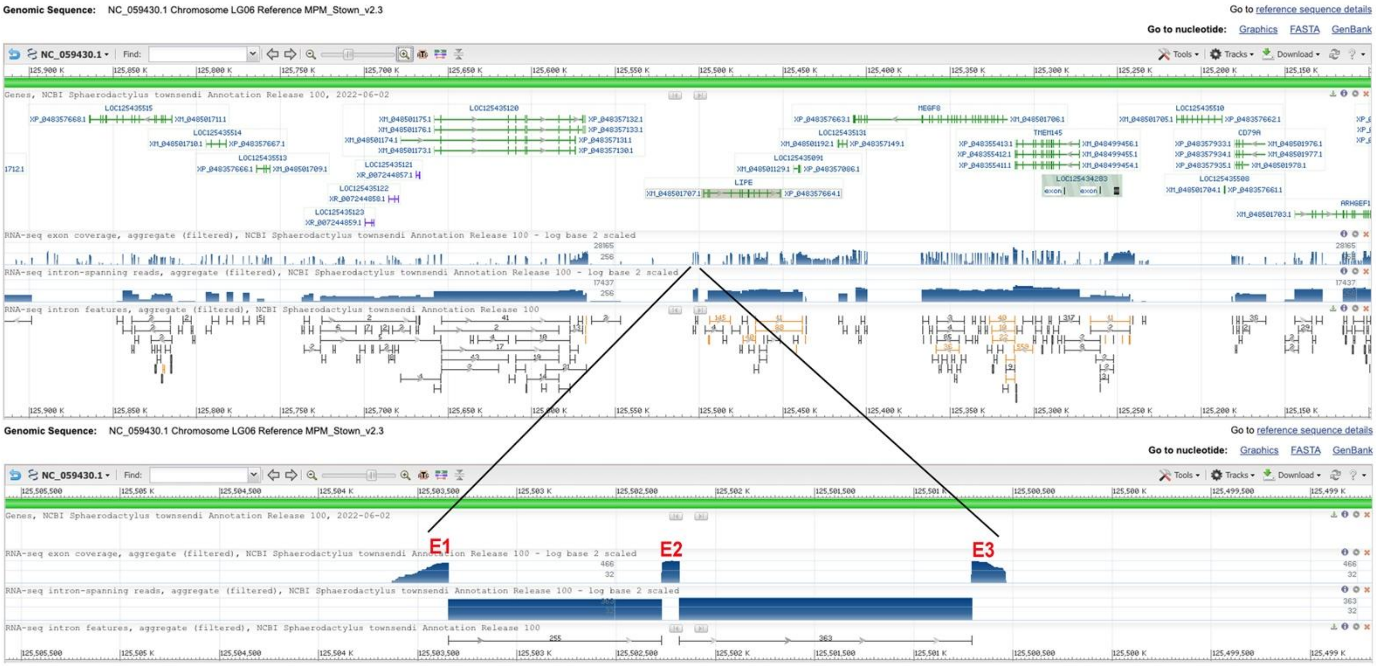


**B:** Sequence of genomic DNA (exons in red and highlighted in yellow, coding region underlined)

atccctttctctaaaccttcgcaaaaatgacatcagaagaacgaaggaggtggcacaggcgaaaagatccctaatcacaaaagtcagaggagagagccccaggggacactgtgattaaaatgggaagattcctagccttggtgatcttgctgctactgctatctg
tcagtaccttttgcttctcagccggcgagaaaggtgaggcaagaaagatggggtgggcccaggtgggatcagccccctttattcctttccactttatgatctgcccagtctgaagacaactagcaggagggagaagagttgctttccttatgagggcaggacagg
aataaaagaagttgtctgcaaaaccgagacacgccaacatagtaccttccttgtgccgaagaggcacctcttcctgtacagacctgcctagacattccggttgccatccaaggacttctgctgtgtgaacagatgcgagatattatctgttacctgtgtgaacagatc
tgtgtgaacagataatcgacagggccttctccatggtggcacctcccttgggaaatgcctctggtgctaactcttttgaattgtgagcactgggggaaagcacttctggtctaggcttctgcattcgtgtcgatatacgttgctgaccttgaaacttcatttctgtgttgct
gttttagttgtttatcgttcggcctggatttttttaaaaaaaattgttgtttttaaatggctggttttactattatggaatttctgacgtaggtttgctaggttatgtggtcatgatctggttgtttttgattctaatgttttcacccgcacacgtgggcgaaacaggagcagcagcaa
cagggccacagccatgcaggtcggaaaacccgcaacacccagttgattctggctgtgaaagccaagaatatatgatggggttttcattgcacattaagctatacataccttgaacatgaaaatgaagcaatgcatgtacaaagaaaaatcaagtgtacactgtaca
tggattgtacatgtgttgattgaaatctgtgaacaaggctattagaagactacttcggggtgttcagaaaatgggaacaatctccttacacaggggtgtggctgccctattctcctctccccggcttttctgactggcctctcttaaattttgtcatctcttctctgcctgcca
ggtcaggacaaatctgggaaatgtgtggggctgttttaccttttggcactgatgcgctgctaatttatccctttctttctgggtcagatcatggattccacattcgccacgtggagagaaagctgagcagatctgagctaggtcacggcaggagtggactttgcccttg
cccaggtgagggccgtctcatatcctacagccaggggaaaagaagttaaagggcaaactctgtttttgctttcccaagtagaaactgggctggcttgtttgccacctgggtgtgcctttcaaaacttttctgcctctcctacactcatctgtggcattaagtcctctgac
gtttccccagacaggtgacccattatttgtaactaggtggcatatttttagcgcgcttttcttgaaagtgctcaggtgtgatttcttccttcttccttttatcacaacctggcctgttccccactgcaaccagcttgatcccagcacggagggcggaatcatcctgtgcctct
tcgccgctccgttccgattggcttctgttctacggccatgttctgtctatccccacacacgttataaaaaaaaactgccacaagaatggagggacaaaggtggcgtttttttgattggccagctgtacgcatgcctgaaacactcagctgtgattggctgaatgggggac
tcctggcacccaagagattccagcactttactgcgaatcgagctgggttaagcgtggttcccctgaaaagtagtagttcccaactggagtcggaaatttgaagccgttacgcggggctgaagctggtacctgaaaccatgtcgatcccagtggttgtgcggagca
cttaggtcgaacgcagctcgaacttaggtcaatagcgcaagtgcggaatcgacccagatgtgtttccacactcctacattcctgctgggtgaccttgagctagtcacagttcttcggaactctctcagccccacctacctcacaaggtgtctattgtgggcagagga
agggaaagtcaccttaagtcacctaacaagagagaaagggggggatataaatccaacccctcctcctcctcctcctcctcctccttctcctcctcctcctcctcctcctcctcctcttcttcttcttcttcttcttcttcttcttcttcttcttcttcttcttcttcttcttcttct
tcttcttcttcttcttcttcttcttcttcttcttcttcttcaagcttcccattgatttctggttccagtaggggccatttttttagtgcacgtggtaaagatatttggaatgggtgggtgaggacagggagaaccccacaccaacaaagagggatacaaggtgaggttgggga
gggggagagacccatggaagaaacgaactcccactcttccaagcaaagccgaatccgcaggaagtacatttcgccttgctgtcgtctcagaggatctttttgcccgtgtcttggggggatcgaggccacttgcaagcttccagaagaaggagttagctggcca
cacaattaagactcacagctttgtgttgtgtacatgaatacgaaaaggagatgggggtccagcatcagaagggacgagaaaacctactttaagtcagtttcttctttccccctctcgcagggtaccaccgtagccattcccacggcaagaacaaacatttccctcac
cgcatgaagcagaagtgcaagtcgtaccagagacactgccaacaccggactctcatgggggggacttcagattcctctgtagctttctgaggctgttgaataaagcaagcccgcctgcttccggcagagagtcttcttttgggtcgtgtgt

**C:** Sequence of cDNA (coding region in red and highlighted in yellow)

gaaaagatccctaatcacaaaagtcagaggagagagccccaggggacactgtgattaaaatgggaagattcctagccttggtgatcttgctgctactgctatctgtcagtaccttttgcttctcagccggcgagaaagatcatggattccacattcgccacgtgga
gagaaagctgagcagatctgagctaggtcacggcaggagtggactttgcccttgcccagggtaccaccgtagccattcccacggcaagaacaaacatttccctcaccgcatgaagcagaagtgcaagtcgtaccagagacactgccaacaccggactctcat
ggggggacttcagattcctctgtagctttctgaggctgttgaataaagcaagcccgcct

**D:** Amino acid sequence (signal peptide shaded)

MGRFLALVILLLLLSVSTFCFSAGEKDHGFHIRHVERKLSRSELGHGRSGLCPCPGYHRSHSHGKNKHFPHRMKQKCKSYQRHCQHRTLMGGLQIPL

**Fig. S17.** Gene synteny and architecture (**A**), genomic DNA sequence (**B**), cDNA sequence (**C**), and encoded protein sequence (**D**) of the unannotated *cxcl17* gene in the NCBI reference genome of *Sphaerodactylus townsendi*. The information was downloaded from the NCBI gene database (https://www.ncbi.nlm.nih.gov/gene).

**A:** Gene synteny and architecture

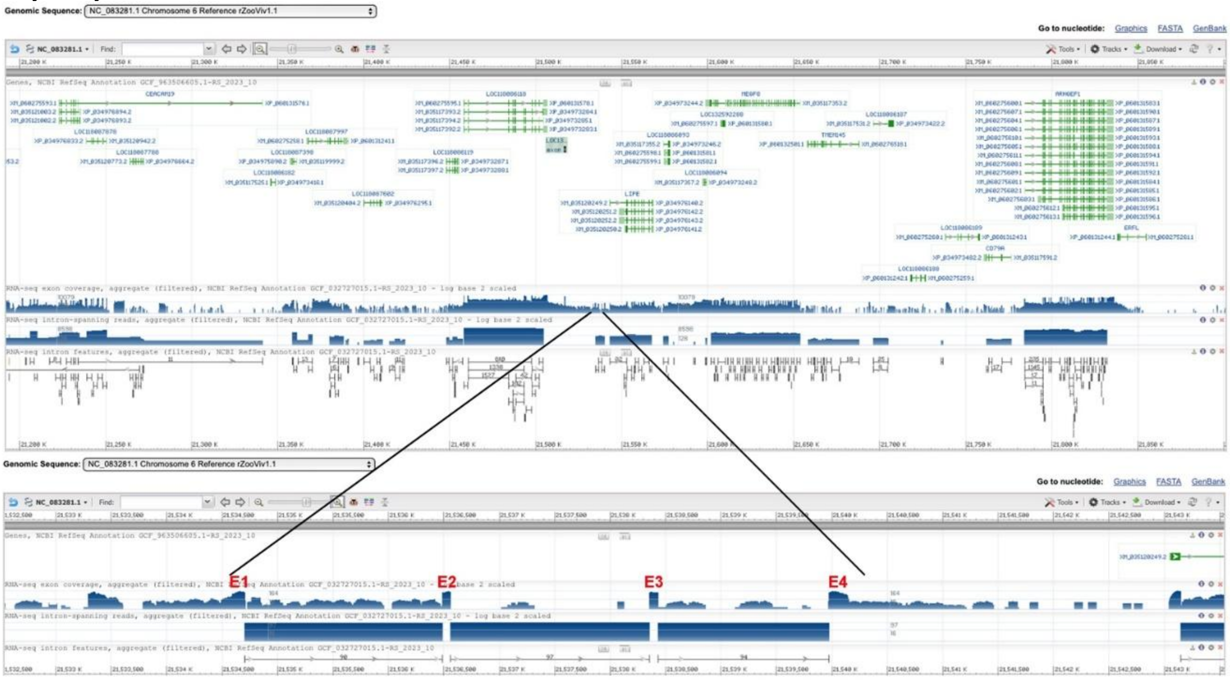


**B:** Sequence of genomic DNA (exons in red and highlighted in yellow, coding region underlined)

cgtatacgggtcgagaggagatttccctcttcacaaaatatgcatctcgaaacttccaggagcttttgaaagcagctctctccaaaaaaacaaacatgttacaggagaatatttctgctccagtgatccagggcctgccccaaaataatccctttcttccccaactcttc
ccgtaaatgacatcaagcataaaggggagttgactcaaacagcagcacaagcagatagatcccccgtcgctacaaatttgtgccttgcagacttctgcgggcgtcgctctttgccatgggccgattcctagctttgactttcctgctcctgtcgctcagcatcctttg
ctgcaccgccaaggataaaggtgagagggaaggattgaaactgggaagaggaggaattgctacgtcaattcttccccctttggtgggcgcagttgaacagtggaactcgctgccgcaggacgcagcgacggccaccaatttggatggctttggaaaaggatt
ggacagatttgtggaggagaagtctgctggtggctattagccatgatggctgtgctctgtctccacggctgaaagcagtaatgcaggggcggactctggtaaaaggccgccctgtgcagactaattgacaccctccctccagccctcatggcaccagtttaagc
caggctttgggaaggaggcatgaggttctggcaaggtcctagtgccctccttcctctttccccaagcccggtaagggctttacttgcctatctttgggaaggagacaggagcccttactgggctttgagaaggcaccttccttggctgcacgcctggtgtgacggc
accgtttgcgctgctgtaaatccacctctgtagtaatgctcctgaacgccagttgccgggaggcacagcagggggagagtatttttgggtttggatcttgcttgtcggattgccacaggcatctggttggcccctgtaggaacaggatgctggactagatgggcca
ttgacctgatcctgcaggtcccagtggtgtagcatgggttgccagtgccccgggctgcatggagggagggaaacagatggagtacacatgtgccttcagctgcctgacggcctccacatcacaaaggagattgcggccgcagatctaagaaaagaagagtc
gttccattgtctagagagatcgcttcttggttctcatggaaaggggtcagcatggcatagaacaccctgctttgccaaccaaaggtgccaggctcaatcccttgctttcgcagctcaggtgctcctgggttgggggggcgtgaggtttgccactgtgtaagacact
ggagaaacgctgttagaataaactagccttccccagtctaggactctctatatagaatcaaagagtttccacttcatgtgtgagttgggattcgaatccagatcctagtcggacattccaactattataccacactgactgaaatgcacatatcagctaagcaaaaaca
gtcaaggaaggcactgagcaggaccggtaagaggcatggtctggcgaaacataacgaccaaggagaatcctgggggccaggcagagatgtctggggggccgcattaggctgccaagccaccccacctgaggtggaaagataagaaacaagtttcccgtc
cccttctcccccaggaccatccacctgtcatggatgctttagttgagattcctgcattgcagagggttggactagatgacccacagggaccctttctgattctatgattcttctcccctctccagacacctgcgcccctcccaaaacctttccagagggtggtgggctc
cttctcaatgaggaatgcagcaggagcgggcagagactgctgcgagagggaagagtcacccagaataggtcttgggacgcaaggaagttcatggcagcaaaaaaaacttatttcctccagcatcgttcaggagagagccgtgactcccctgagtggctttgc
agagggcacagtggggaggagcaggcggtaacttatccttctgtgtctctaaatctaacttgctccatataaataacccgccctcattctcagagtagctgtcttgctgttcagagtttacccaattgaacattttctttcccattttctctgccaggtagaagattccgttc
aggccacaaggagcgacagccagcccggggaggccaccaaccctgcagttgttccggtgagtcaccttcttgcaccaccaaccccggcagttgagaatctggaaactctctcccccaagttccgcttttggtttcctctgtagcagctggcccggcctgtttatt
gctgaggtgtggcattcaaaaaagataggaccggtaacctcttgcacaaatcactcaggctcatcgcttggtttgagagtggcacaaaccaaacctgactgagtgcaatttgactctgaaattttccagtgtgaaaaatggaggtggatgatggtcatcgctcattta
tatagaataataaatgttgacggttaataatgcagcaatacaccgcttaacaccaaattaggcttctaagtggtcttattaaaaaaaaaaccaattacgcaagcatttaaatataaagcacccataattaaaatgcacagacaatcccattaaaagttagaatgtgagca
tccgtagttaaaatatgcaatacagtaaagccaaatgatgcattaattaaagcagggaaatcctgctggaggggtgaaggaaatgcctgtgtaaacaggcgcatctttaaaagccggtgaattagattgggtttttttgcatcattagtgagtgaggcaaggactca
gcatggtctgctatggattaactcagaacataaaggtggaacttcatctatggagaatggggaaaactctcctttgtgtgactaatctatggccctccagatctcatttatggccctccagctcctatcctccccaaccattggccatgctggttgcggctgatgggtgt
tgaagaacatctggagaaccacccctggctcactcccacaatggagaattggagggggggcagcggaggaggaattgcatgtgctttggctcgttttcacgagacatctctgcagtcaacacccctaacataggaacctgagaagagccaatggcctatttagt
ccaacatccttcttctcacgcccagaggcctatgtgaaaccctcagttgggaccctagcacaagagcaccctccccactcatggtttccagcaactggtattcagaagcgttgcttgctagcaaccatggaagcagaggcgtgccacaaggaccataagccact
gtctgccctctccctgaatttgtccagtcctcctttaaagccatctaagttggtgtctcagggatgcaggtggcgctgtgggttaaaccactgtgccgcttgggcacagtgagctcctgttgctcggtcccagctcctgccaacctagcagttcgaaagcacatcaaa
gtgcaagtagataaataggtaccactctgacaggaaggtaaacggcgtttccgtgcactgctctggcttcaccagaagcggcttagtcatgctggccacacgacccggaaaaactgtctccggaagcggacaaacgccggctcccttggcttataaagcaaga
tgagagccgcaaccccagagtcgtttgtgactggactttattgtcagtggtccctttacctttacctttaaatttggtggctatgattgcatcctgtggcaaggagttccgcagtttactatgtgcatacagtttctttggatgtccacgagttctacggtagagttaggagc
gctctcctatattctgcttgcatgccctaaccctttttggttatgggggaggagggaggaccataaaagcaccccccccacaccccaaatcttgcacgtgcctcttcccactgccagttaccatacctaattacggtactcatggcttatgttactttgttggcttatgttgtt
ttagacctgcgtgagccctggaatccaaagaaacacggacggccgtgtccttgtgacggcatccttactggtttgtctctgccaggtaacaaaatacctttagacagggctggtggtagagataagatgaatgggaaacatttatcctccacccccccacccccccca
agaaaaagacttcaaaggggagaggtaatttattgtgtatcctaggtaactagaatctcattgatgggagggagaaagttgtagtcaacacctgggtcccctaaccagatcagtctcaactctgtatggtttgtgcttgtgcttgtatctcctatttgtggccatgtaagc
gtaaatcagccatgcctaaccattgggaaagggttgtagctccaagttagagcatctgttttgaatgcagaaggtcccaggttcaatccaggggtgggagggaccttgtgaacgacctttatgtgaccccctataaaaagtgtttgttctacagatcaaagcgactga
aagcataccttgggcaggggtgaaactaggctttatttcacccagtttggcacactgaccccaagcgcatttgtgtaaacacacacacacacacacacacacagagagagagagagagagagagagggaggggggagggagggagggaggctgggagcct
caagggcgcctccctggaggcttcatgggggggcaggcaccagctagccccctcccccagctacagcactgcatcaggtaaggtgatctcacaagtcaactgctccctaatggttaagggattaatatagtaatagtaacaccttattttggcccagtagcaatt
cagcaattcagatctctgagcagaggtgttttatgtgtgggggggaggggggtggctcgtacacacttgcacttgtccacttgcctataagacaagtgtccaaatggtttccccccttgatccgctcctttcccagggaaaacctgctctttacagctagatcggagcag
acgtcaatttggagtaaaataaaaaaaattccttcagtagcaccttaaagaccaactaagtttttattttggtataagctttcgtgtgcatgcacacttcttcagatacgctgaaataggagaaaacccaaattgccatttgctccagttgagcactaaagagtggtttccc
ccggaggaaagcagcgggagaagaggaagtttggataaaaccagggtctcgctcctgccagcagcattctgcgctagctgcagcatccagatcaaatgcaagggaagccccacacagagtgcattgcagttgaagtggggtaacctgtgtaaaagggggca
agaacattggaatatgccttttactgagtctgactactggtctatctagcttggtgtttctcaaacttaggtccccagctgttttacagccccccatcatccctagccagcaggaccagtgggcagggatgatgggaagtgtagtctagaaacaacagctttccaggtttt
ccagagaggggccattcccaaccctccttggagatgccaggggttagggttagggttagggtcgatcactaagcaagagcatcagaagggaagatataaactgtcacccccctttttttttttgcagaacagctccatcaccaccaccgtggaatggccaagcg
acatttgcaccagagaaggcatcggtgccaggcgtttgtgaataagtgccacatgatgcgggttgacatacccttgtagctttctctgcccccaagcacagggcttaattctttgtgttaaccataaaacaaatccctcttcggtacctgctgtctttacttcgttcgtttc
cccacaacagccctgcgaagtaggtgaagctgagaggcagcgatcggcctacgtttacccag

**C:** Sequence of cDNA (coding region in red and highlighted in yellow)

Gcagcacaagcagatagatcccccgtcgctacaaatttgtgccttgcagacttctgcgggcgtcgctctttgccatgggccgattcctagctttgactttcctgctcctgtcgctcagcatcctttgctgcaccgccaaggataaaggtagaagattccgttcaggcc
acaaggagcgacagccagcccggggaggccaccaaccctgcagttgttccgacctgcgtgagccctggaatccaaagaaacacggacggccgtgtccttgtgacggcatccttactggtttgtctctgccagaacagctccatcaccaccaccgtggaatgg
ccaagcgacatttgcaccagagaaggcatcggtgccaggcgtttgtgaataagtgccacatgatgcgggttgacatacccttgtagctttctctgcccccaagcacagggcttaattctttgtgttaaccataaaacaaatccctcttcggtacctgctgtc

**D:** Amino acid sequence (signal peptide shaded)

MGRFLALTFLLLSLSILCCTAKDKGRRFRSGHKERQPARGGHQPCSCSDLREPWNPKKHGRPCPCDGILTGLSLPEQLHHHHRGMAKRHLHQRRHRCQAFVNKCHMMRVDIPL

**Fig. S18.** Gene synteny and architecture (**A**), genomic DNA sequence (**B**), cDNA sequence (**C**), and encoded protein sequence (**D**) of the unannotated *cxcl17* gene in the NCBI reference genome of *Zootoca vivipara* (common lizard). The information was downloaded from the NCBI gene database (https://www.ncbi.nlm.nih.gov/gene).

**A:** Gene synteny and architecture

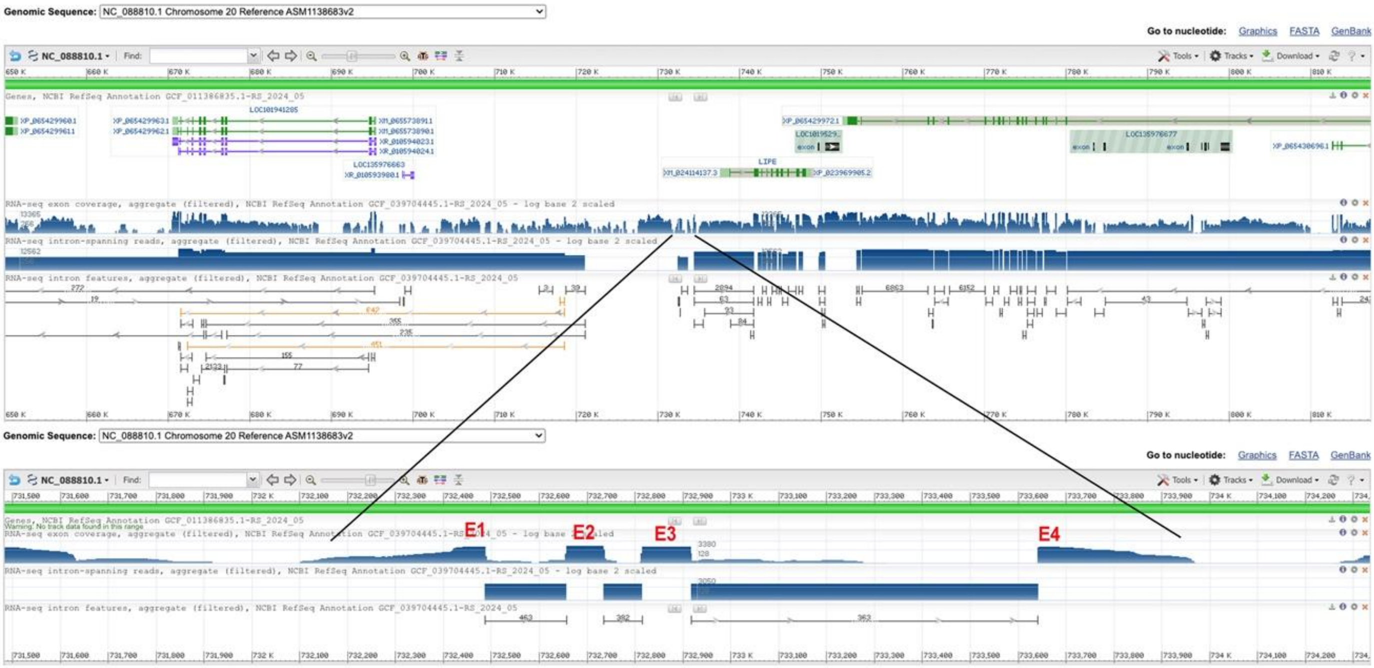


**B:** Sequence of genomic DNA (exons in red and highlighted in yellow, coding region underlined)

aatgggtccccccccgctgggccaaaccgcagccaatgggcaccgggctggcagccgctggggtataaaggctgcggcagccccaagacagcgctgcggaaatggaccaactcctgtgcctgttcgtgctgctgtctgtaacaatcacaacctgctcgcc
ccatgtgccaggtgagaggggatgggggtacctgtataacccgcccccacaccccaccccgagccagctgcacctcagcaccgggcaaggggtccctgtataaacagcccctgcgccccaccccagaggggccgcatctcagcaccgggcgtttagctt
cccttctcctgtctgacgcaggtggtggacacggagcggacggggagctgaacccagtcagacggcgcgcgaccggagcagggctctgtgagtgcccaggtcagtgtcccctgccatggggtgaagttgggtgggggaggagccacacccacacccac
gtgatgtctcctgtcatttcaggctggtcgggatccaagcgccagctcggagacctcagggatggggccagacacaatgggctgtgcccctgtgacggcatcgtcccccatggccaaggcagggggtgagagccgctcctagcatcgggggtgtgggagca
gggccagggaatgggatctgctaacggggcagggtcagggttaggggctgagtggcatcagggaacgagggaaggggatttgagtctcatcacaaggtgcaaggaatggggtttaggactgaggtggaggggggcaaggattgggggatgggattgag
gggaagaggggaggaattggggtctagtaatggggtgcagtgaatggggttggggatgaggtaggggttgagattggggtctgacactgaggggaagagaggatggattggggtctggtaacggggggcagagaatggggttggggatgaggtagggg
ttgagattggggtctgacactgaggggaagagaggatagattggggtctggtaatggggggcagagaggaggtttgagattgaggtgggatgcagggattggggtctgacactgaggggaagagaggatggattggggtctggtaatggggggcagaga
ggaggtttgagattgggatgggtgcagggattggggtctgacattgaggggaagagaggatggattggggtctggtaatggggggcagagaggaggtttgagattgggatgggtgcagggattggggtctggcattgagagggggtctggggtctagctgg
gggaagggcaggagacacccctgctaacccccccccccccgtctctcccgcagggcgtcacccccagggccggggcagacggacccagcaccacaggcgcagacggcagcggtgccgcgagatcccgagccgatgccgggagaagggcctggcca
aggggctgctgctgcccctctagcggcccgtccccggggccctgagcggagccagtgggggggcgtcgcctttcgccccgtcaataaaaacccagcgcgaggggacgcgccccgtctggtgtccgtgtgcggctccttcaccctgacctctgaccccacg
gccggggtcaaaatcgcccccctgacacctcccccctggggccctgcaccgacccctcccccgctcctggggcagcagcatgggggggggggcttcttctctccctgcggggcgttaac

**C:** Sequence of cDNA (coding region in red and highlighted in yellow)

ggggtataaaggctgcggcagccccaagacagcgctgcggaaatggaccaactcctgtgcctgttcgtgctgctgtctgtaacaatcacaacctgctcgccccatgtgccaggtggtggacacggagcggacggggagctgaacccagtcagacggcgcg
cgaccggagcagggctctgtgagtgcccaggctggtcgggatccaagcgccagctcggagacctcagggatggggccagacacaatgggctgtgcccctgtgacggcatcgtcccccatggccaaggcagggggcgtcacccccagggccggggca
gacggacccagcaccacaggcgcagacggcagcggtgccgcgagatcccgagccgatgccgggagaagggcctggccaaggggctgctgctgcccctctagcggcccgtccccggggccctgagcggagccagtgggggggcgtcgcctttcgcc
ccgtcaataaaaacccagcgcgaggggacgcgccc

**D:** Amino acid sequence (signal peptide shaded)

MDQLLCLFVLLSVTITTCSPHVPGGGHGADGELNPVRRRATGAGLCECPGWSGSKRQLGDLRDGARHNGLCPCDGIVPHGQGRGRHPQGRGRRTQHHRRRRQRCREIPSRCREKGL
AKGLLLPL

**Fig. S19.** Gene synteny and architecture (**A**), genomic DNA sequence (**B**), cDNA sequence (**C**), and encoded protein sequence (**D**) of the unannotated *cxcl17* gene in the NCBI reference genome of *Chrysemys picta bellii* (western painted turtle). The information was downloaded from the NCBI gene database (https://www.ncbi.nlm.nih.gov/gene).

**A:** Gene synteny and architecture

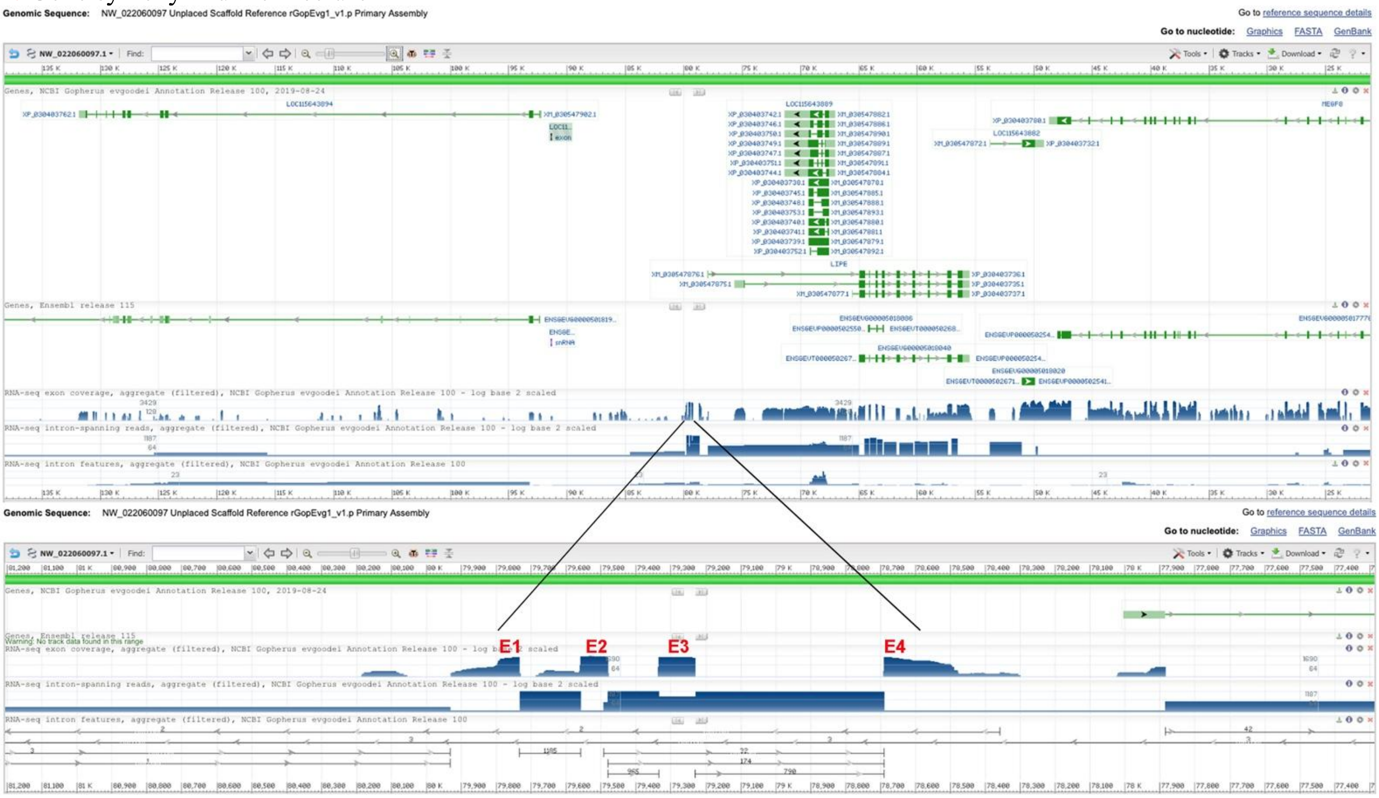


**B:** Sequence of genomic DNA (exons in red and highlighted in yellow, coding region underlined)

ttcctgctcggttcccgtcagagtcgtcagggtgacagaacaatgggtcccccccgctgggccaaaccgcagccaatgggcactggtctggcaccgggacggcagccgctggcatataaaggctgtggcagccccgagacagcgctgccaaaatggacc aactcctgtccctgttcctgctgtctgtaggcatggcagcctgctcggcccatgtgccaggtgagaggggatgggggtacctgtataaccagcccccacactccaccccagagccagctgcatctcggcactgggcgaggggtccctgtataaacagcccctg tgtcccaccccagagggggccgcatctcagaaccaggcaagcatttagcttcccttctcgtgtctgacgcaggtggtggacgcagagcggaaggggagctgaacccagtcagacagcgcgcaaccggagcagggctctgtgagtgtccaggtcagtgtccc ctgtcacagggtgaagttgggtgggggaggagccacacacacacacacacacacacacacacacacacacacacacacacacacacacacacacacacacacacacgtctcctgtcatttcaggctggtcgggatcacagcgccagc tcgcaggagatctcgggtttggggccagacacaatggggtgtgcccctgtgacagcatcatcccccacggccaaggcaggagtgagagctgctcccagcatcaggggtggggtgggtggggtgagtgagcagggccagggaatgggatctgctaacga ggtggggtcagggttaggggctagaaccgacagaggcagagaatggcatcagggaaagagggaagggtattcgagtctcctcacagggtgctaggaatggggttggggactgaggtggaggggggcaaggattgggggatgggactgaggggaaga ggagaggaattggggtctggtaacggggtgcagagaatggagtgtaggatggggtggggtgcagggagtgggtctggtaacagtgcagagattggggtttgagattggggtgggatgcagagattgaggtctggcattgaggggaagagaggatggact gaggtctgataacggggggcagagaacaggtttgagattggggtggggtgcagggattggggtctggcattgagagggggtctggggtctagctgggggaagggcaggagacacccctggtaactgccaccccccgtctctcccgcagggcatcacccc cagagccggggcaggcggagcgaacaccacaggcgtagacggcagcggtgccgcaagaccccgaggcgatgccggcggcagggcctgaccaaggggctgctgctgcccctctagctgccgtcccgggggctctgagtgcagccagtggggggcgt tgccttctgcccgtcaataaaaacccagctcgaggggatgctcccgtgtggtgtccgggtgtggctccttcacccctgacctctgacccccccagccggggtcaaaatcgccc

**C:** Sequence of cDNA (coding region in red and highlighted in yellow)

Gacggcagccgctggcatataaaggctgtggcagccccgagacagcgctgccaaaatggaccaactcctgtccctgttcctgctgtctgtaggcatggcagcctgctcggcccatgtgccaggtggtggacgcagagcggaaggggagctgaacccagtc agacagcgcgcaaccggagcagggctctgtgagtgtccaggctggtcgggatcacagcgccagctcgcaggagatctcgggtttggggccagacacaatggggtgtgcccctgtgacagcatcatcccccacggccaaggcaggaggcatcacccccca gagccggggcaggcggagcgaacaccacaggcgtagacggcagcggtgccgcaagaccccgaggcgatgccggcggcagggcctgaccaaggggctgctgctgcccctctagctgccgtcccgggggctctgagtgcagccagtgggggcgttg ccttctgcccgtcaataaaaacccagctcgagg

**D:** Amino acid sequence (signal peptide shaded)

MDQLLSLFLLSVGMAACSAHVPGGGRRAEGELNPVRQRATGAGLCECPGWSGSQRQLAGDLGFGARHNGVCPCDSIIPHGQGRRHHPQSRGRRSEHHRRRRQRCRKTPRRCRRQGL TKGLLLPL

**Fig. S20.** Gene synteny and architecture (**A**), genomic DNA sequence (**B**), cDNA sequence (**C**), and encoded protein sequence (**D**) of the unannotated *cxcl17* gene in the NCBI reference genome of *Gopherus evgoodei* (Goodes thornscrub tortoise). The information was downloaded from the NCBI gene database (https://www.ncbi.nlm.nih.gov/gene).

**A:** Gene synteny and architecture

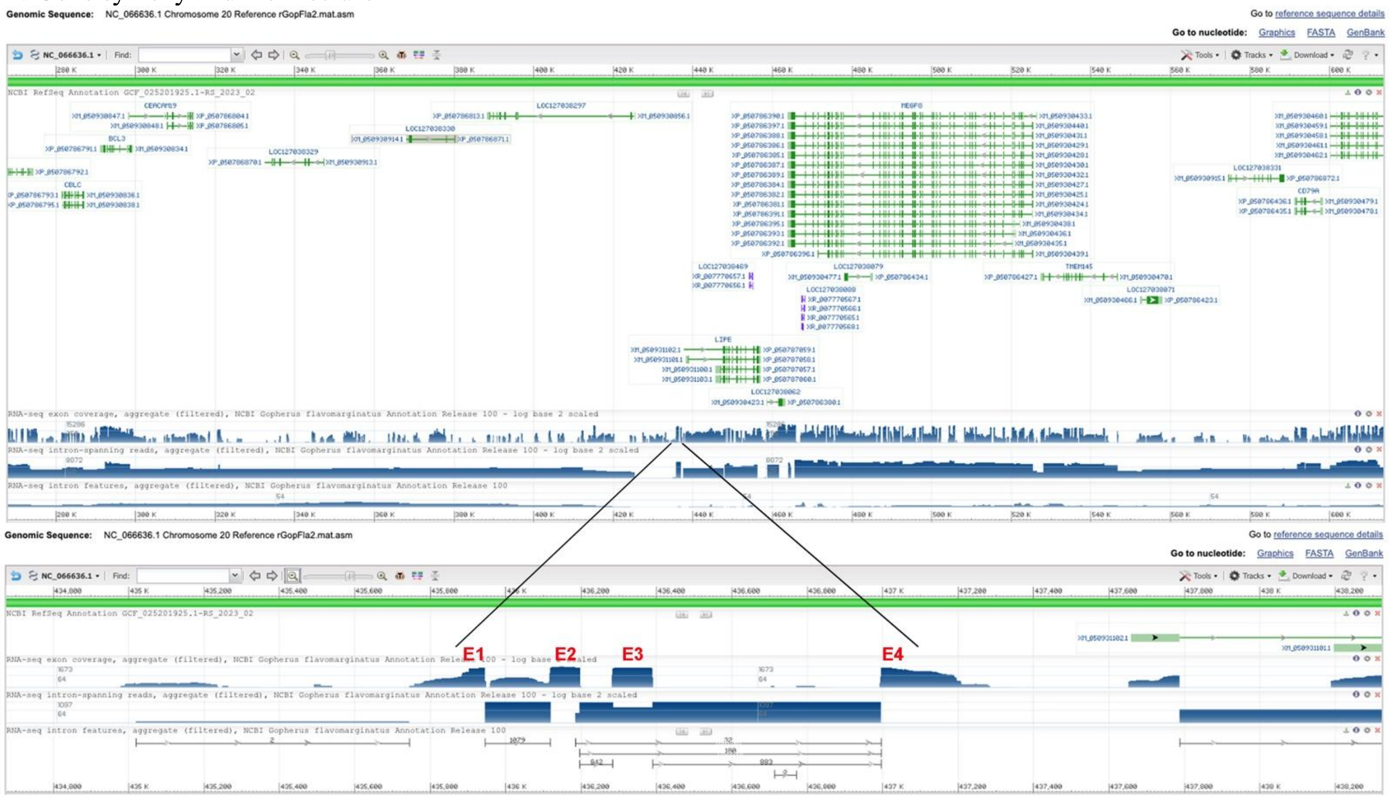


**B:** Sequence of genomic DNA (exons in red and highlighted in yellow, coding region underlined)

ggggcactgcacgttctgggaatcagcccaaaagtcaccgttctgcactaatccttcctgctcggttcccgtcagagtcgtcagggtgacagaacaatgggtcccccccgctgggccaaaccgcagccaatgggcaccggtctggcaccgggacggcagcc
gctggcgtataaaggctgtggcagccccgagacagcgctgccaaaatggaccgactcctgtccctgttcctgctgctgtctgtaggcatggcagcctgctcggcccatatgccaggtgagaggggatgcgggtacctgtataatcagcccccacactccaccc
cagagccagctgcatctcggcactgggcgagggatccctgtataaacagcccctgtgtcccaccccagagggccgcatctcagaaccaggcaagcatttagcttcccttctcgtgtctgacgcaggtggtggacgcagagcggaaggggagctgaaccca
gtcagacagcgcgcgaccggagcagggctctgtgagtgtccaggtcagtgtcccctgtcacagggtgaagttgggtgggggaggagccacacacacaaacacaccacacatacgtctcctgtcatttcaggctggtcgggatcacggcgccagctcgcag
gagatctcggggttggggccagacacaatggggtgtgcccctgtgacggcatcatcccccacggccaaggcaggggtgagagctgctcccagcatcaggagtgggaggggtgggggggggtggggtgagtgagcagggccagggaatgggatctgcta
acggggtggggtcagggttaggggttagaaccgacagaggcagagaatggtatcagggaacgagggaagggtattcgagtctcctcacagggtgctaggaatggggttggggactgaggtggagggggggcaaggattgggggatgggactgaggga
agaggggaggaattggggtctggtaatggggtgcagagaatggagtgtaggatggggtggggtgcagggattgggtctggtaacagtgcagagaatggggttgtggatgaggtaggggttgagattggggtctggtaacaaggtgcagagaatggggtttg
agattggggtgggatgcagagattgaggtctggcattgaggggaagagaggatggactgaggtctgataacggggggcagagaacagctttgagattggggtggggtgcagggattggggtctggcattgagagggggtctggggtctagctgggggaa
gggcaggagacacccctggtaaccgccacccccccgtctctcccgcagggcgtcacccccagagccggggcaggcggagcgagcaccacaggcgtagacggcagcggtgccgcgagaccccgaggcgatgccggcggcagggcctgaccaaggg
gctgctgctgcccctctagctgccgtccctggggctctgagtgcagccagtggggggcgttgccttccgctccgtcaataaaaacccagctcgaggggatgctcccgtgtggtgtccgggtgtggctccttcgccgctgacctctgacccccccagccggggtc
gaaatcgccccctgacacctgcccccagggctctgcaccaac

**C:** Sequence of cDNA (coding region in red and highlighted in yellow

Ggctgtggcagccccgagacagcgctgccaaaatggaccgactcctgtccctgttcctgctgctgtctgtaggcatggcagcctgctcggcccatatgccaggtggtggacgcagagcggaaggggagctgaacccagtcagacagcgcgcgaccggag
cagggctctgtgagtgtccaggctggtcgggatcacggcgccagctcgcaggagatctcggggttggggccagacacaatggggtgtgcccctgtgacggcatcatcccccacggccaaggcaggggggcgtcacccccagagccggggcaggcggag
cgagcaccacaggcgtagacggcagcggtgccgcgagaccccgaggcgatgccggcggcagggcctgaccaaggggctgctgctgcccctctagctgccgtccctggggctctgagtgcagccagtggggggcgttgccttccgctccgtcaata

**D:** Amino acid sequence (signal peptide shaded)

MDRLLSLFLLLSVGMAACSAHMPGGGRRAEGELNPVRQRATGAGLCECPGWSGSRRQLAGDLGVGARHNGVCPCDGIIPHGQGRGRHPQSRGRRSEHHRRRRQRCRETPRRCRRQG
LTKGLLLPL

**Fig. S21.** Gene synteny and architecture (**A**), genomic DNA sequence (**B**), cDNA sequence (**C**), and encoded protein sequence (**D**) of the unannotated *cxcl17* gene in the NCBI reference genome of *Gopherus flavomarginatus* (Mexican gopher tortoise). The information was downloaded from the NCBI gene database (https://www.ncbi.nlm.nih.gov/gene).

**A:** Gene synteny and architecture

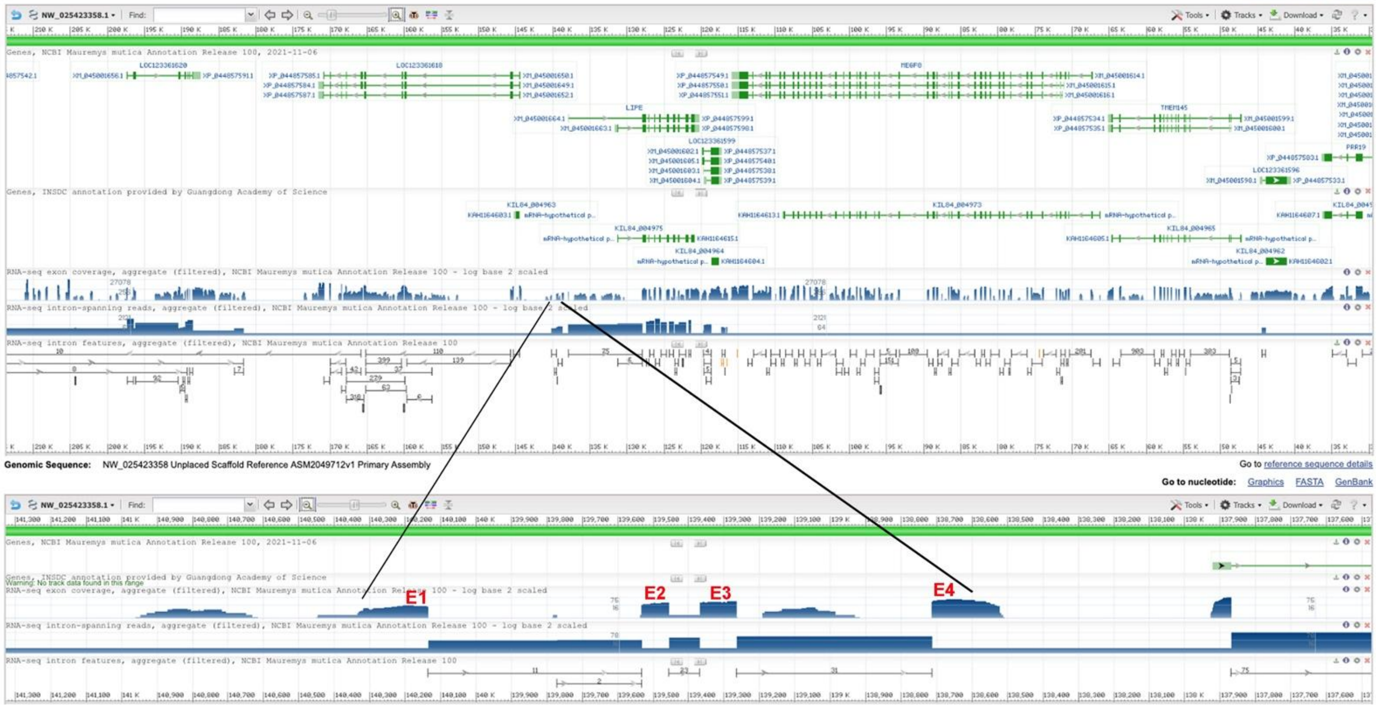


**B:** Sequence of genomic DNA (exons in red and highlighted in yellow, coding region underlined)

cagggtgacaggacaatgggtcccccccgctgggccaaaccgcagccaatgggcaccgggatggcaccgggatggcagccgctggggtataaaggctcggcagcccaaagacagcgctgccgaaatggaccgactcctgtgcctgttcctgctgctgtc
tgtaggcatggcagcctgctcggcccctgtgccaggtgagaggggatgggggtctctgtataaccagcccccacactccaccccagaggtggctgcatctcggcaccgggcgaggggtctctgtataaacagcccctgtgccccaccccagagccagctg
catctcggcacagggcgaggggtccctgtataaacagcccctgtgccccaccccagagccagctgcatctcggcaccgggcgaggggtctctgtataaacagccccccaccccagagccagctgcatctcggcacagggcgaggggtccctgtataaaca
gcccctgtgctccaccccagagccagctgcatctcggcaccaggcaaggggtccctgtataaacagcccctgtgccaggtgagagggggtggggtacctgtataaccagccctcacatcccaccccagagccagctgcatctcggcaccgggcgaagggt
ccctgtataaacagcccctgtgccccaccccagagccagctgcatctcggaaccgggcgaggggtccctgtataaacagcccctgtgccccaccccagaggggccgcatctcagcacccggcgagcgtttagcttcccttctcgtgtctgacgcaggtggtg
gacacggggcggaaggggagttgaacccagtcagacagcacacgaccggagcagggctctgtgactgcccaggtcagtgtcccctgtcacggggtgaagttgggtgggggaggagcccccacacacacatacacacatgatgtctcttgtcatttcaggc
tggtcgggatcccagcgccagctcgcaggagaccttggggttggggccagacacaatgggctgtgcccctgtgacagcatcgtcccttatggccaaggcagggggtgagagtcgctcccagcattgtgggtgggggagcagggccagggaacgggatctg
ctaacggggcggggttagggttaggggctagaaccgacggaggcagagaacggcatcagggaacgagggaaggggattcgagtctcctcacagggtgcaaggaatggggttggggactgaggtggaggggggcaaggactgggggatgggaatga
ggggaagagggaaggaattggggtctagtaacagggtgcagataacggagtgtaggatggggtggggtgcagggattgggtctggtaacagtgcagagaatgggggtggggatgaggtaggggtcgagattgggatctgacactgaggggagagagg
atggactagggtctgataacggggtgcagagaggaggtttgagatggggtgggatgcacggattgggggctggcattgaggggaagggggaaggcttggggtctggcattgagaggggtctggggtctagctgggggaagggcaggagacacccctgc
taacccccccccgtctctcccgcagggcatcacccccagggccggggcaggcggagcgagcaccacaggcgccgacggcagcggtgccgcgagatcccgagccgatgccggcggcagggcctggctaaggggctgctgctgcccctctagctcccgt
ccccggggccatgagtggagccagtggggggcgtcgccttccgccccgtcaataaaaacccagctcgagggg

**C:** Sequence of cDNA (coding region in red and highlighted in yellow

Tataaaggctcggcagcccaaagacagcgctgccgaaatggaccgactcctgtgcctgttcctgctgctgtctgtaggcatggcagcctgctcggcccctgtgccaggtggtggacacggggcggaaggggagttgaacccagtcagacagcacacgacc
ggagcagggctctgtgactgcccaggctggtcgggatcccagcgccagctcgcaggagaccttggggttggggccagacacaatgggctgtgcccctgtgacagcatcgtcccttatggccaaggcagggggcatcacccccagggccggggcaggcg
gagcgagcaccacaggcgccgacggcagcggtgccgcgagatcccgagccgatgccggcggcagggcctggctaaggggctgctgctgcccctctagctcccgtccccggggccatgagtggagcca

**D:** Amino acid sequence (signal peptide shaded)

MDRLLCLFLLLSVGMAACSAPVPGGGHGAEGELNPVRQHTTGAGLCDCPGWSGSQRQLAGDLGVGARHNGLCPCDSIVPYGQGRGHHPQGRGRRSEHHRRRRQRCREIPSRCRRQG
LAKGLLLPL

**Fig. S22.** Gene synteny and architecture (**A**), genomic DNA sequence (**B**), cDNA sequence (**C**), and encoded protein sequence (**D**) of the unannotated *cxcl17* gene in the NCBI reference genome of *Mauremys mutica* (yellowpond turtle). The information was downloaded from the NCBI gene database (https://www.ncbi.nlm.nih.gov/gene).

**A:** Gene synteny and architecture

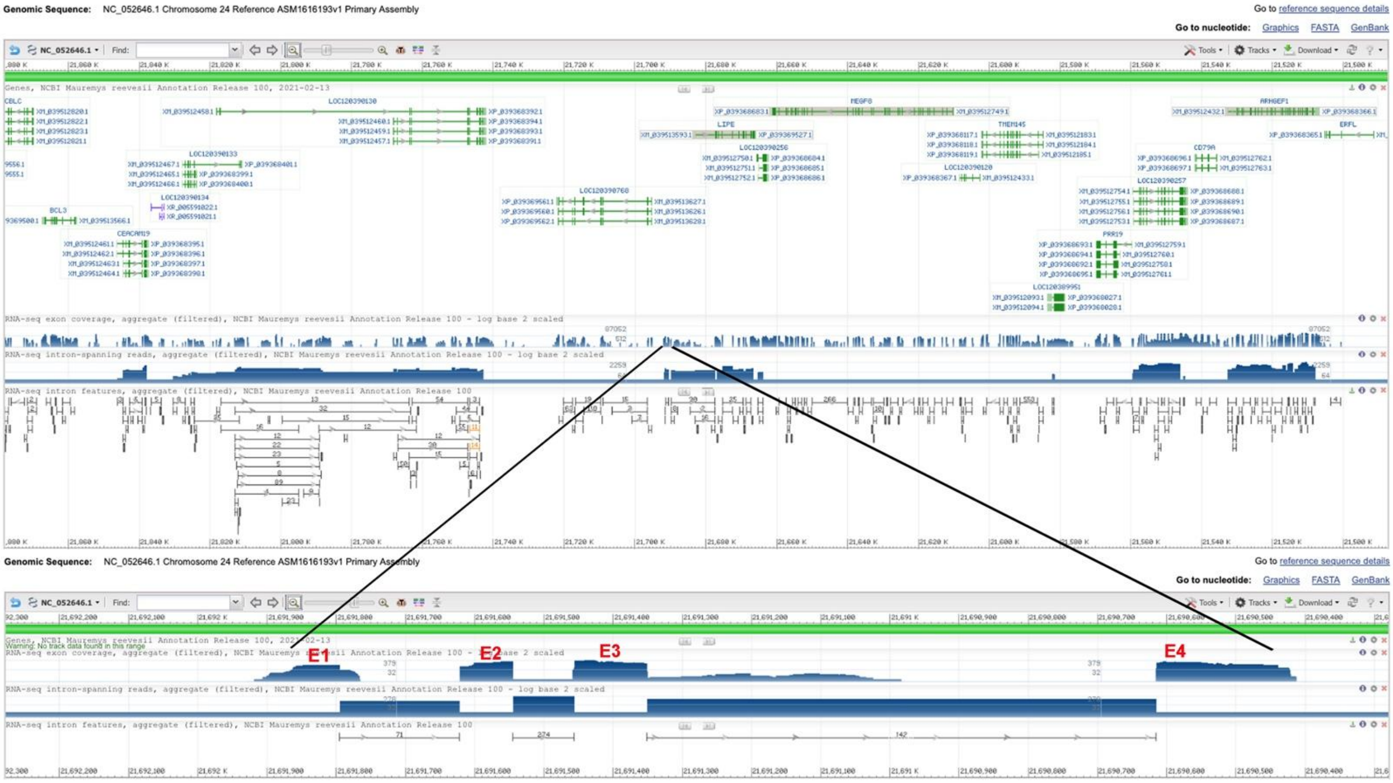


**B:** Sequence of genomic DNA (exons in red and highlighted in yellow, coding region underlined)

cactgcacgctctgggaatcagcccaaaagtcaccgctccgcactaatccttcctgctcggttcccgtcagagtcgtcagggtgacaggacaatgggtccccccccccgctgggccaagccgcagccaatgggcaccgggacggcagccgctggggtat
aaaggctgcggcagcccccagacagcgctgccgaaatggaccgactcctgtgcctgttcctgctgctgtctgtaggcatggcagcctgctcggcccctgtgccaggtgagaggggatgggggacctgtataaccagcccccacaccccaccccagagcca
gctgcagctcggcaccgggcgaggggtccctgtataaacagcccctgtgccgcaccccagaggggctgcatctcagcacctggcaagcgtttagcttcccttctcgtgtctgacgcaggtggtggacacagagcggaaggggagctgaacccagacagac
ggcacgcgaccggagcagggctctgtgagtgcccaggtcagtgtcccctgtcacggggtgaagttgagtgggggaggagcccccccccatacacatacacacatgatgtctcttgtcatttcaggctggtcgggatcccagcgccagctcgcaggagacct
cggggttggggccagacacaatgggctgtgcccctgtgacggcatcgtcccccatggccaaggcagggggtgagagctgctcccagcgttgagggtgggggagcagggccagggaatgggatctgctaacggggcggggtcagggttaggggctagaa
ccgacagaggcagagaacggcatcagggaacgagggaagggaattcgagtctcctcacagggttcaaggaatggggttggggactgaggtggagggggtcaaggattgggggatgggattgaggggaagcgggaaggaattggggtctggtaacgg
ggtgcagagaatggtgtgtaggatggggtggggtgcagggattgggtctggtaacagtgcagagaatggggttggggatgaggtaggggttgagattgggatctgacactgaggggaagagaggatggactgtggtctggtaacgaggcagagaatggg
gtttgagatggggtgggatgcatggattggggtccggcattgaggggaagagaggatggattggggtctggtaacgggggcagagaagaggtttgagatggggtggggtgcagggattggggtctggcattgaggggaagagaggatggattggggtct
ggtaacagggggcagagaagaggtttgagattggggtggggtgcagggattggggtctggcattgaggggaagggggaaggcttggggtctggcattgagaggggtctggggtctagctgggggaagggcaggagacacccttgctaaccccaccccc
ccgtctctcccacagggcatcacccccagagccggggcaggcggagcgagcaccacaggcgccgacggcagcggtgccgcgagatcccgagccgatgccggcggcagggcctggccaaggggctgctgctacccctctagctcccgtccccgggg
ccctgagtggagccagtgggggcgtcgcctttcgccgcgtcaataaaaacccagctcgaggggacgttcccgtctggtgtccgtgtgcggctccttcgcccctgacctctgaccccacggccggggtcaaaatcacccctgactcctccccctggggccctg
caccaaccccgcccctgctcctggggcaccagcacgggggggggggcgtcctctctccctgcggggtgttaaccctggagcctagtgatggcggctggggtcagccaagggcgcaaggagggagggggcccggggtgtcccgacggggggggggagg
ggagcccccagcacagggtcgggggggggggcggccctaggggaggt

**C:** Sequence of cDNA (coding region in red and highlighted in yellow)

aaaggctgcggcagcccccagacagcgctgccgaaatggaccgactcctgtgcctgttcctgctgctgtctgtaggcatggcagcctgctcggcccctgtgccaggtggtggacacagagcggaaggggagctgaacccagacagacggcacgcgacc
ggagcagggctctgtgagtgcccaggctggtcgggatcccagcgccagctcgcaggagacctcggggttggggccagacacaatgggctgtgcccctgtgacggcatcgtcccccatggccaaggcagggggcatcacccccagagccggggcaggc
ggagcgagcaccacaggcgccgacggcagcggtgccgcgagatcccgagccgatgccggcggcagggcctggccaaggggctgctgctacccctctagctcccgtccccggggccctgagtggagccagtgggggcgtcgcctttcgccgcgtcaat
aa

**D:** Amino acid sequence (signal peptide shaded)

MDRLLCLFLLLSVGMAACSAPVPGGGHRAEGELNPDRRHATGAGLCECPGWSGSQRQLAGDLGVGARHNGLCPCDGIVPHGQGRGHHPQSRGRRSEHHRRRRQRCREIPSRCRRQG
LAKGLLLPL

**Fig. S23.** Gene synteny and architecture (**A**), genomic DNA sequence (**B**), cDNA sequence (**C**), and encoded protein sequence (**D**) of the unannotated *cxcl17* gene in the NCBI reference genome of *Mauremys reevesii* (Reeves's turtle). The information was downloaded from the NCBI gene database (https://www.ncbi.nlm.nih.gov/gene).

**A:** Gene synteny and architecture

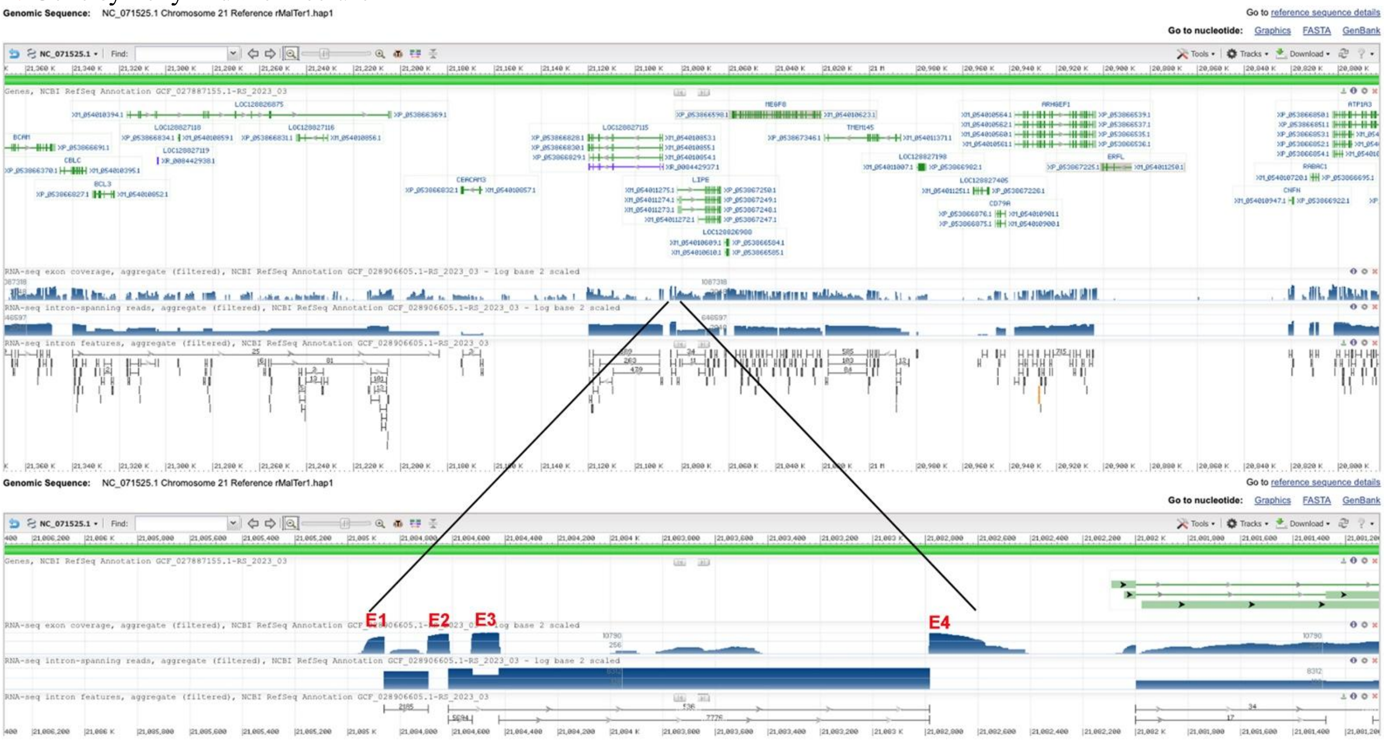


**B:** Sequence of genomic DNA (exons in red and highlighted in yellow, coding region underlined)

cactaatccttcctgctcggttcccgtcagagtcatcagggtgacaggacaatgggtcccccccccccccccgctgggccaaaccgcagccaatgggcaccgggctggcagccgctggggtataaaggctgcggcagccccaagacagcgctgcggaaa
tggaccaactcctgtgcctgttcctgctgctgtctgtaacaatcacaacctgctcgccccgtgtgccaggtgagaggggatgggggtacctgtataaccgcccccacaccccaccccgagccagctgcacctcagcaccgggcaaggggtccctatataaaca
gcccctgcgccccaccccagaggggccgcatctcagcaccgggcgtttagcttcccttctcctgtctgatgcaggtggtggacacggagcggacggggagctgaacccagtcagacggcgcgtgaccggagcagggctctgtgagtgcccaggtcagtgt
ccctgccatggggtgaagttgggtgagagaggagacacccccccacacacacacacacatgtgatgtctcctgtcatttcaggttggtcgggatccaagcgccagctcggagacctcggggatggggccagacacaatgggctgtgcccctgtgatggcat
cgtcccccatggccaaggcagggggtgagagcccagcatcgggggtaggggagcaaggccagagaatgggatctgctaacggggcggggtcagggttaggggctagtaatgacaggggcagagtggcatcagggaacgagggaaggggatttgagtc
tcatcacagggtgcaaggaatggggtttaggactgaggtggaggggggcaaggattgggggatgggattgaggggtagagaggaccaaatggggtctggtaaagggtgcagagaatgggattggggatgaggtaggggttgagattggggtctgacatc
gaggggaagaggggatggattggggtctggtaaaagtgcagagaatagggttggggatgaggtaggggttgagattggggtctgacactgaggggaagagaagatggattggggtctggtaatggggggcagagaggaggtttgagattgaggtgggtg
cagtgttgcggagtgtgggggagtcagggccctgcacccctcttcctgagattcactgcgactctcaaccagccagtaaagcagaaggtttatttaaggacaggaacacagtctcaagcagagcgtgtaggtacgaccagacccctcaattaggtccctctgg
gaggttcagggagcttagaccccagcttggggttccctgcgttgcaccacccagcccaaactgaaaccaaaccaaaactcctcctgcagctcctctcttccccctccccccagctcctcctctcctttgttcagtctcccgggcagaaggtgttaattctccccacc
cccattcctggctcaggttacagcacaggtagcttccttcaagggaagtcccacatccccactgcaacccccctgcaacattcccaggtcaaatctgccccgctccctgctccgtcacatgcagggattgggatctgacactgaggggaagagaggatggattg
gggtctggtaatggggggcagagaggaggtttgagaatgggatgggtgcagggattgggatctgacactgaggggaagagaggatggattggggtctggtaaggggatgcagagaggaggtttgagaatgggatgggtgcagggattgggatctgacac
tgaggggaagagaggatggattggggtctggtaatggggggcagagaggaggtttgagaatgggatgggtgcagggattggggtctgacactgaggggaagagaggatggattggggtctggtaaggggatgcagagaatggggttggggatgaggta
ggggttgagattgggatctgacactgaggggaagagaggatggattggggtctggtaatggggggcagagaggaggtttgagattgggatggatgcagggattggggtctgacactgaggggaagagaggatggattggggtctggtaatggggggcag
agaggaggtttgagattgaggtgggatgcagggattgggatctgacactgaggggaagagaggatggattggggtctggtaatggggggcagagaggaggtttgagattgggatgggtgcagggattggggtctggcattgagaggggggtctggggtcta
gctgggggaagggcaggagacacccctgctaagcccccccccccgtctctcccgcagggcgtcacccccggggccggggcagacggacccagcaccacaggcgcagacggcagcggtgccgcgagatcccgagccgatgccggcagacgggcct
ggccaaggggctgctgctgcccctctagctccccgtccccggggccctgagcggagccagtggggggcgtcgcctttcgccccgtcaataaaaacccagctcgaggggacgcgcccgtctggtgtccgtgtgcggctccttcaccccctgacctctgacccc
acggccggggtcaaaatcgccccctgaccccctcccccggggccctacaccgacccctcccccgctcctggggcagcagcacggggggggcgttaaccctggagcctagtgatggcggctggggtcacccgagggcaggaggcgggagaggcctgg
ggtgtcccgaccggggggctgggggagagggggaggggagccccccactcccagcacagggtcgggggggggcggccctaggggtggggccagaggaggtgacactgccacgtgg

**C:** Sequence of cDNA (coding region in red and highlighted in yellow)

Tgggccaaaccgcagccaatgggcaccgggctggcagccgctggggtataaaggctgcggcagccccaagacagcgctgcggaaatggaccaactcctgtgcctgttcctgctgctgtctgtaacaatcacaacctgctcgccccgtgtgccaggtggtg
gacacggagcggacggggagctgaacccagtcagacggcgcgtgaccggagcagggctctgtgagtgcccaggttggtcgggatccaagcgccagctcggagacctcggggatggggccagacacaatgggctgtgcccctgtgatggcatcgtccc
ccatggccaaggcagggggcgtcacccccggggccggggcagacggacccagcaccacaggcgcagacggcagcggtgccgcgagatcccgagccgatgccggcagacgggcctggccaaggggctgctgctgcccctctagctccccgtcccc
ggggccctgagcggagccagtggggggcgtcgcctttcgccccgtcaataaaaacccagctcgaggggacgcgcccgtctg

**D:** Amino acid sequence (signal peptide shaded)

MDQLLCLFLLLSVTITTCSPRVPGGGHGADGELNPVRRRVTGAGLCECPGWSGSKRQLGDLGDGARHNGLCPCDGIVPHGQGRGRHPRGRGRRTQHHRRRRQRCREIPSRCRQTGL
AKGLLLPL

**Fig. S24.** Gene synteny and architecture (**A**), genomic DNA sequence (**B**), cDNA sequence (**C**), and encoded protein sequence (**D**) of the unannotated *cxcl17* gene in the NCBI reference genome of *Malaclemys terrapin pileata*. The information was downloaded from the NCBI gene database (https://www.ncbi.nlm.nih.gov/gene).

**A:** Gene synteny and architecture

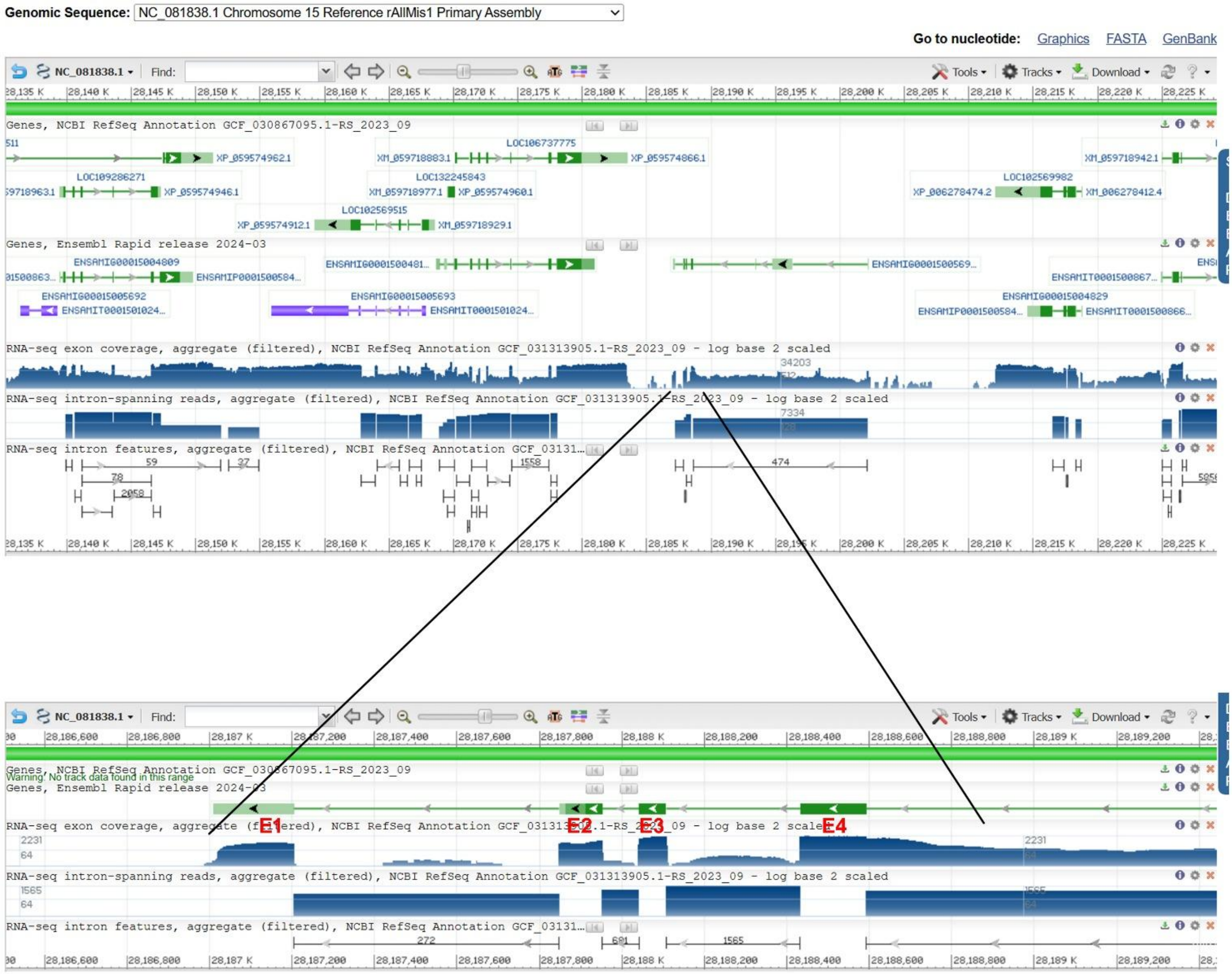

**B:** Sequence of genomic DNA (exons in red and highlighted in yellow, coding region underlined)

caaaagcctgggagggttaggagccagaatgcctgggttcatgtttcagtttggtttcacctggcccgggcacaccttctgggaaagcaaattctttgtcacggcaagtgctgggcttggctctgctttgaatcaactctgctttgtttctccttccttctggcacctctt
ctgggactttccatggggcagccaatgagacctggctgggcaaaaacgggcagggggaaggcccgcagtgataaagccagggctgcggatgcctctctcagtgccaccaacatggacctacgccttagcttcgtgctgctgctgctgctgctggggctggg
atcggctgtgaagaagccaggtgagaggggaaggggcccagggaccagctgtaatgtccccaccccagagcgagttgcattgtggcaccaggcaaggggtcctggtatggccaggctccaatatattccatcccagaggtgcccagctcagcctcaatgc
gctgcaccagagccagctgcatctcagggtcctgttatacccagccctgacgtcccatcccagaacaggctgcatcccagtgtgtcccaccccaaaaccagctgcatcccagcccagggggagggatccctctctagccccagccccatgccagtgcctacc
ttgccttctccccttgggcacaggcaggcgtccagcccagaaggaaaatggagactcagagacatccctcgacctctgctcatgcgaaggtgagtgtgtcccaccctgaaccatggcagggtgtgagggcagggggataacccagccccccctcgccacac
acaccgtctcgtttcaggctggttgggcgcctggggccagctcgagggcggcgtctgggggggggcaggggcacaacagcgcatgcccctgcaggggcatccacccccacgaggtgatgacaggtgagagctggccaggggtggggacaggggctgg
gacgggttaagtgggggcacagcatggggtcagaaggatggaggtagtagggggcactggggagcaggggcactggggacgggtaagtgggggcacagcatggggtcagaagggtggaggtagtagggggcactggggagcaggggcactggg
gacgggtaagtggggacacagcatggggtcagaagggtggaggtagtagggggcactggggagcaggggcactggggacgggtaagtgggggcacagcatggggtcagaagggtggaggtagtagggggcactggggagcaggggcactgggg
acgggtaagtgggggcacagcatggggtcagaagggtggaggtagtagggggcactggggagcaggggcactggggacgggtaagtgggggcacagcatggggtcagaagggtggaggtagtagggggcactggggagcaggggcgtgggcaa
ggaatgtgggctgggggagaggatgtgcggtaagagggttggggcaggtagggggaattgggggaagggggcaggggtagggagcacagcctggggggtgggacggggtcacaagggcaccactcaccccccttctccccttctccctgtagccccgca
ccgccccacacagggccgggcccagggcaggcccaagaggcaccagcgggtctggcgatgccggcagttcctgcgacggtgccagcagcgcagcctgatgcaggggctgctcctgcccctctaggggctgtggggagggtctccagctccaagctgg
tcattaaaccccctggaagcacctggcatctgcacgtgcttccttgaaccctgaccttggaccctgcctttggggcctgacctgcactagggaccccagggctggtggtgccagtgccctctgaccccggggcctggccctgccttctgatgccaggaacccccc
tgaccc

**C:** Sequence of cDNA (coding region in red and highlighted in yellow)

gtgataaagccagggctgcggatgcctctctcagtgccaccaacatggacctacgccttagcttcgtgctgctgctgctgctgctggggctgggatcggctgtgaagaagccaggcaggcgtccagcccagaaggaaaatggagactcagagacatccctc
gacctctgctcatgcgaaggctggttgggcgcctggggccagctcgagggcggcgtctgggggggggcaggggcacaacagcgcatgcccctgcaggggcatccacccccacgaggtgatgacagccccgcaccgccccacacagggccgggccca
gggcaggcccaagaggcaccagcgggtctggcgatgccggcagttcctgcgacggtgccagcagcgcagcctgatgcaggggctgctcctgcccctctaggggctgtggggagggtctccagctccaagctggtcattaaaccccctggaagc

**D:** Amino acid sequence (signal peptide shaded)

MDLRLSFVLLLLLLGLGSAVKKPGRRPAQKENGDSETSLDLCSCEGWLGAWGQLEGGVWGGQGHNSACPCRGIHPHEVMTAPHRPTQGRAQGRPKRHQRVWRCRQFLRRCQQRSLM
QGLLLPL

**Fig. S25.** Gene synteny and architecture (**A**), genomic DNA sequence (**B**), cDNA sequence (**C**), and encoded protein sequence (**D**) of the unannotated *cxcl17* gene in the NCBI reference genome of *Alligator mississippiensis* (American alligator). The information was downloaded from the NCBI gene database (https://www.ncbi.nlm.nih.gov/gene).

**A:** Gene synteny and architecture

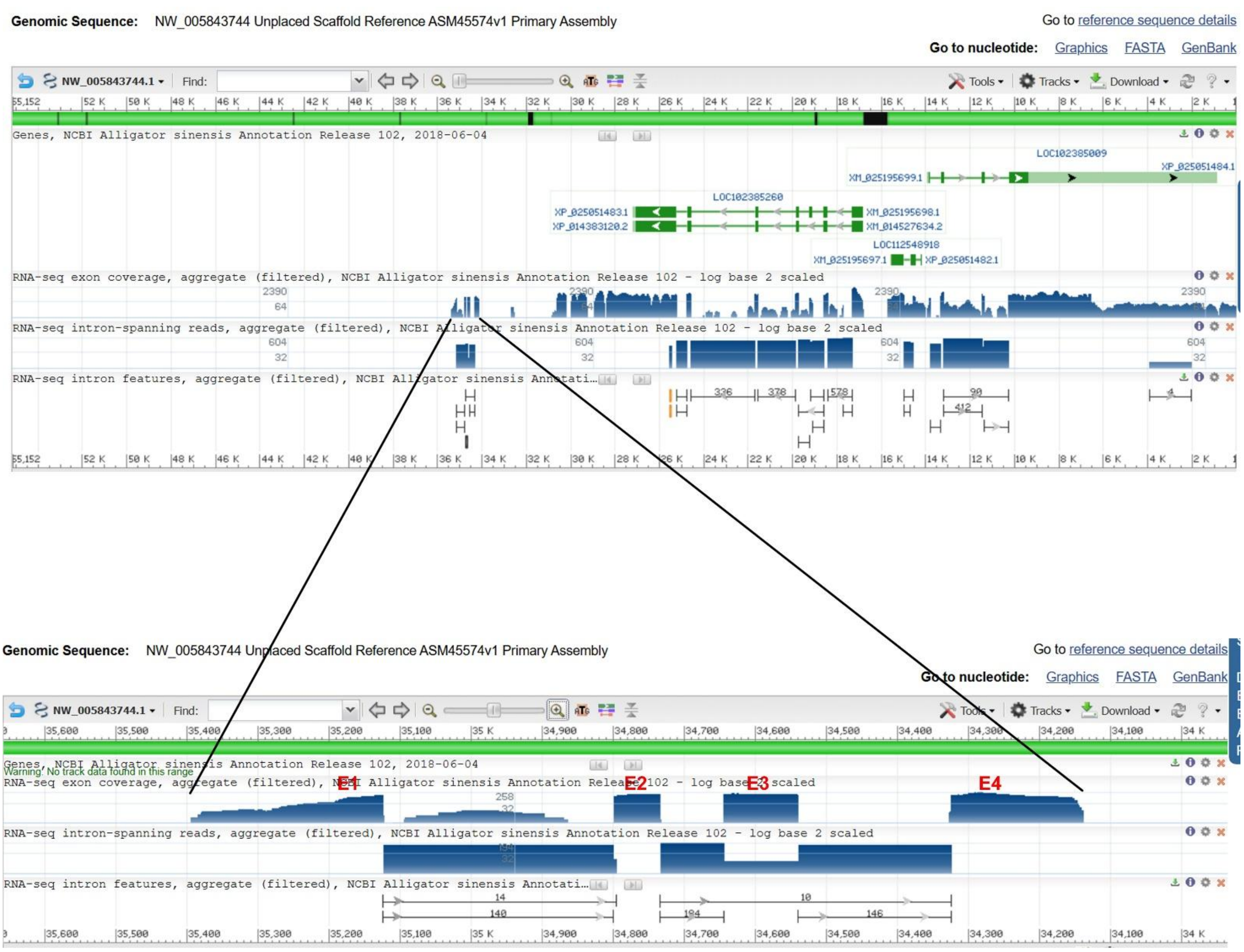


**B:** Sequence of genomic DNA (exons in red and highlighted in yellow, coding region underlined)

Tcatgtttcagtttggtttcacctggcccgggcacaccttctgggaaagcaaattctttgtcacggcaagtgctgggcttggctctgctttgaatcaactctgctttgtttctccttccttctggcacctcttctgggactttccatggggcagccaatgagacctggctgg
gcaggggggaaggcccgcagtgataaagccagggctgtggatgcctctctcagtgccaccaacatggacccacgccttggcttcgtgctgctgctgctggggctgggatcggctgtgaagaatccaggtgagaggggaaggggcccagggaccagctgta
atgtcccaccccagagcgagttgcattgtggcaccaggcaaggggtcctggtatggccaggctccaatatatttcatcccagaggtgcccagctcagcctcaatgcgctgcaccagagccagctgcatctcagggtcctgttatacccagccctgacgtcccat
cccagaacaggctgcatcccagtgtgtcccaccccaaaaccagctgcatcccagcccaggggggagggatccctctctagccccagccccatgccagtgcctaccttgccttctccccttgggcacaggcaggcgtccagcccagaaggaaaatggaggctc
agagacatccctcgacctctgcccatgcgaaggtgagtgtgtcccaccctgaaccatggcagggtgtgagggcaggggggataacccagcccccccctcgccacacacaccgtctcgtttcaggctggttgggcgcctggggccagctcgagggcggcgtct
gggggggggcaggggcacaacagcgcatgcccctgcaggggcatccaccccccacgaggtgatgacaggtgagagccggccaggggtggggccaggggctgggacgggtaagtgggggcactggggagcaggggcactggggacgggtaagtgg
gggcacagcatggggtcagaaggatggaggtagtagggggcactggggagcaggggcgtgggcaaggaatgtgggctgggggagggccacaagggcaccactcacccccttctccccttctccctgcagccccgcaccgccccacacagggccggg
cccagggcaggcccaagaggcaccagcgggtctggcggtgccggcagttcctgcgacggtgccagcagcgcagcctgatgcaggggctgctcctgcccctctaggggctgtggggagggtctccagctccaagctggtcattaaaccccctggaagcac
ctggcgtctgcacatgc

**C:** Sequence of cDNA (coding region in red and highlighted in yellow)

Cccgggcacaccttctgggaaagcaaattctttgtcacggcaagtgctgggcttggctctgctttgaatcaactctgctttgtttctccttccttctggcacctcttctgggactttccatggggcagccaatgagacctggctgggcaggggggaaggcccgcagtg
ataaagccagggctgtggatgcctctctcagtgccaccaacatggacccacgccttggcttcgtgctgctgctgctggggctgggatcggctgtgaagaatccaggcaggcgtccagcccagaaggaaaatggaggctcagagacatccctcgacctctgcc
catgcgaaggctggttgggcgcctggggccagctcgagggcggcgtctgggggggggcaggggcacaacagcgcatgcccctgcaggggcatccaccccccacgaggtgatgacagccccgcaccgccccacacagggccgggcccagggcaggcc
caagaggcaccagcgggtctggcggtgccggcagttcctgcgacggtgccagcagcgcagcctgatgcaggggctgctcctgcccctctaggggctgtggggagggtctccagctccaagctggtcattaaacccccctggaagcac

**D:** Amino acid sequence (signal peptide shaded)

MDPRLGFVLLLLGLGSAVKNPGRRPAQKENGGSETSLDLCPCEGWLGAWGQLEGGVWGGQGHNSACPCRGIHPHEVMTAPHRPTQGRAQGRPKRHQRVWRCRQFLRRCQQRSLMQG
LLLPL

**Fig. S26.** Gene synteny and architecture (**A**), genomic DNA sequence (**B**), cDNA sequence (**C**), and encoded protein sequence (**D**) of the unannotated *cxcl17* gene in the NCBI reference genome of *Alligator sinensis* (Chinese alligator). The information was downloaded from the NCBI gene database (https://www.ncbi.nlm.nih.gov/gene).

### Alignment of avian GPR25 orthologs

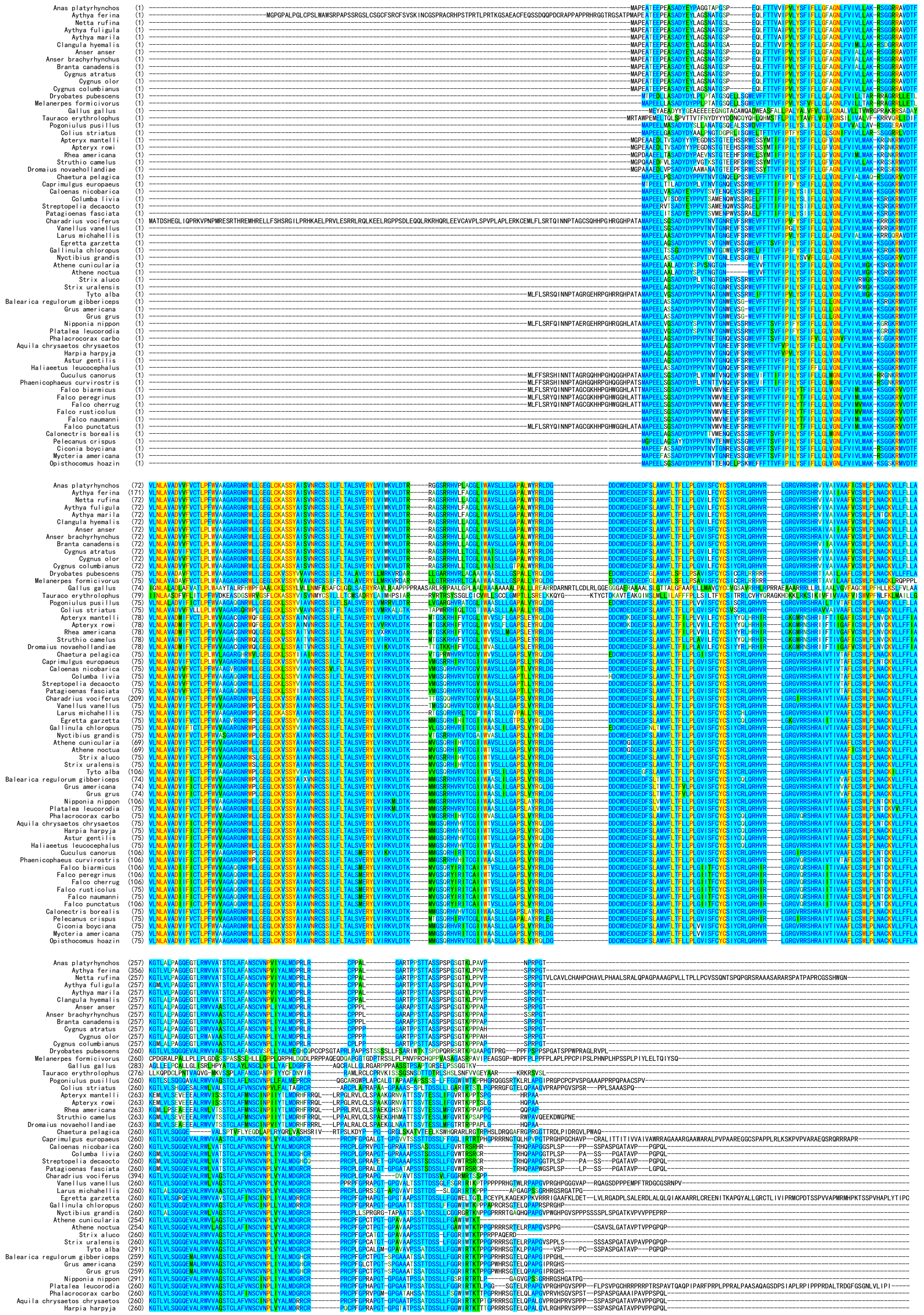

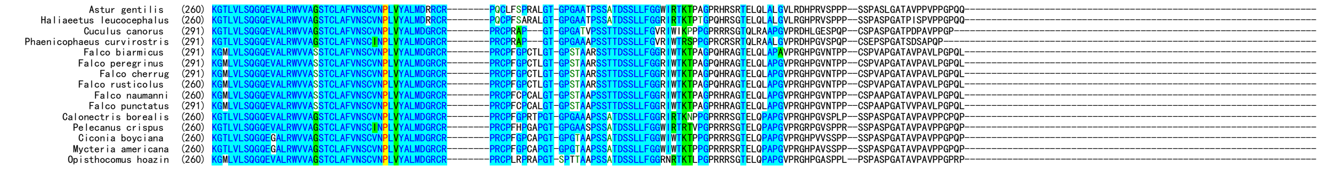

Alignment of reptilian GPR25 orthologs

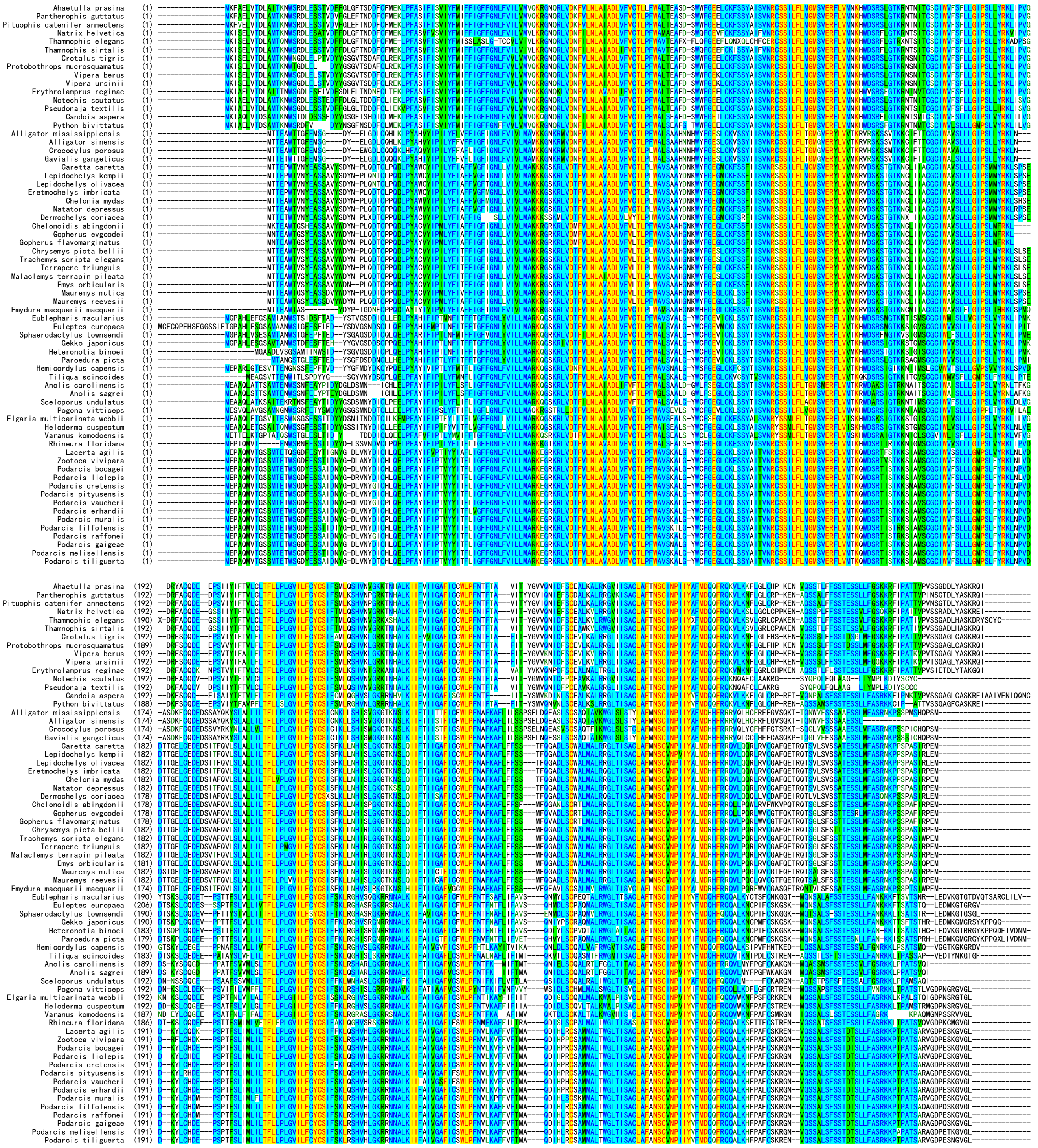

**Fig. S27.** Amino acid sequence alignment of avian and reptilian GPR25 orthologs. Information about these GPR25 orthologs is retrieved from NCBI database and listed in Tabe S3.
